# A system dynamics model to understand the integrated ecological and human dimension aspects of wildlife health and disease management

**DOI:** 10.1101/2025.09.22.677405

**Authors:** Stephanie R. Penk, Christine Anhalt-Depies, Richard E.W. Berl, Thomas S. Fiddaman, Kawika Pierson, Jennifer L. Price Tack, Bryan J. Richards, Erica Rieder, Daniel J. Storm, C. LeAnn White, Daniel P. Walsh

## Abstract

Chronic wasting disease (CWD) presents an ongoing challenge for the management of deer populations and sustaining harvest opportunities across North America. Existing disease models often fail to fully capture the complex interplay between disease dynamics, host ecology, and socio-economic factors. We developed a comprehensive system dynamics (SD) model that integrates demographic, epidemiological, ecological, and socio-economic processes within a single model to more fully characterize the complex network of causal feedbacks throughout the system. The model was calibrated using a Bayesian approach that incorporates prior knowledge to generate biologically interpretable outcomes, even with sparse data. For estimating the joint posterior distribution of model parameters, we leveraged time series of deer abundance, harvest composition, genetic profiles, CWD surveillance, and hunter demographics and behavior. Model outputs reproduced key system behaviors, including observed CWD prevalence trends, deer population dynamics, and hunter license purchasing patterns. Model predictions were most sensitive to parameters governing initial deer population size and recruitment. While model predictions generally aligned with observed data, discrepancies in early CWD detection and overestimation of the reactivation of long-inactive hunters reflect data limitations and modeling challenges. Key results suggest that indirect transmission is necessary to explain observed prevalence, that transmission is moderately density-dependent, and that observed population-level genetic shifts driven by CWD may play a role in transmission and progression. The SD modelling approach enabled estimation of difficult-to-measure parameters and identified potential leverage points for management—such as prioritizing increasing participation in antlerless harvest of existing hunters over the recruitment of new hunters. This integrated modeling approach offers a flexible foundation for adaptive wildlife disease management and emphasizes the value of unifying biological and human dimension processes to better inform effective, evidence-based policy.

## Introduction

Ecology is the study of the interrelationships between organisms and their environment, often with the goal of managing ecosystems amid increasing pressures such as environmental alterations, habitat loss and encroachment, invasive species, and disease (Ellis, 2015)—for the benefit of diverse stakeholders. Ecologists frequently rely on models to describe and predict system dynamics, forming the foundation for many conservation and management strategies. Ecological modeling, by nature, demands a holistic approach (Jørgensen, 2006) to reveal the mechanistic linkages within ecosystems that drive observed patterns and processes (Geary et al., 2020). Despite this need, many ecological studies still adopt reductionist approaches, examining individual components in isolation and often overlooking key endogenous effects, feedback loops, and interactions that shape system trajectories (Evans et al., 2016; Sutherland, 2006). For instance, many important social and anthropogenic pressures –which typically fall under research into the human dimensions of natural resources –such as changes in land use and habitat fragmentation (Haddad et al., 2015) and the use of recreational harvest as a management tool (Berl et al., 2025; Stedman et al., 2004) or as part of conservation interventions (Nilsson et al., 2020; Veríssimo et al., 2025) are widely recognized as having both direct and indirect effects on ecosystems, yet these factors are rarely integrated explicitly into models of ecosystem functioning. The field of human dimensions can provide information for management using applied social science to understand the role of human values, behaviors, and social systems as determinants, participants, and targets of management action (Bennett et al., 2017). Human dimension studies can provide the necessary knowledge, but there remains a pressing need within ecology for methods capable of incorporating the complexity of these social-ecological systems. Such approaches are essential to enable strong inference about the current and future behavior of ecosystems under increasing anthropogenic influence.

Systems thinking is an approach used to evaluate the cause-and-effect relationships among multiple interacting factors and their cumulative impacts on both individual components and the system as a whole (Behl & Ferreira, 2014; Forrester, 1994; Jackson, 2003; Verhoeff et al., 2018). Although systems thinking has been widely advocated in ecological research (e.g., Folke et al., 2021; Jackson, 2003; Mahajan et al., 2019), its application to capture the nonlinearities inherent in ecosystem dynamics—particularly those emerging from interactions with socioeconomic drivers—remains underutilized in modeling efforts. Most current ecological models rely heavily on the use of simple regression-based approaches, low-dimensional models with limited explanatory power, or focused research on single system components, which often lack the generalizability to produce robust predictions for dynamic ecosystems (Evans et al., 2016; Sutherland, 2006). A holistic alternative to traditional methods is system dynamics (SD) modeling, which uses systems thinking insights to create dynamic simulation models from large-scale systems of differential equations that effectively describe the interrelationships within a system of interest and can be used to explore its dynamics through computer simulation (Forrester, 1994). System Dynamics (SD) models are akin to simple ordinary differential equation (ODE) compartment models, in which the compartments are referred to as stocks and the rates at which stocks change are described as flows. Within SD models, causal connections are explicitly mapped as feedback loops so that interdependencies are integrated across multiple ODE systems to create complex combinations of delays, non-linearities, and knock-on effects that drive the system behavior over long time horizons (Forrester, 1994; Sterman, 2000, 2002). In SD models, endogenous variables vary due to casual interactions we include in the model, while exogenous variables are inputs that are not changed by model behavior. These models rely on mechanistic relationships to define the causal structure linking variables, enabling simulations that reflect how system behavior emerges from its underlying dynamics rather than relying on phenomenological relationships which cannot be used to explore novel conditions. This approach allows the merging of advanced quantitative model estimation methods with qualitative knowledge about the structural connections and parameter sensitivities within a system, which can be invaluable for mapping the ecological and socioeconomic complexities that characterize many environmental systems (Elsawah et al., 2017; Treves et al., 2025; Wittmer et al., 2006).

The spread of chronic wasting disease (CWD) in both wild and captive cervids exemplifies the complex interplay between ecological and socioeconomic systems (Figure 1A). Chronic wasting disease is an invariably fatal prion disease affecting deer, elk, moose, and other cervid species (Bishop, 2004; Haley & Hoover, 2015; Vaske et al., 2021). Despite decades of varied control efforts, the disease has continued to increase in both prevalence and geographic range across North America (National Wildlife Health Center, 2025). The causative agent is a misfolded form of the host’s normal prion protein that, unlike most other pathogens, lacks any genetic material (Bartz et al., 2024). Transmission occurs through both direct and indirect pathways. Direct transmission involves contact between infected and susceptible individuals (Dobbin et al., 2023; Haley & Hoover, 2015; Thompson et al., 2024), while indirect transmission occurs when healthy animals are exposed to prions from environments (Almberg et al., 2011; Miller et al., 2004; Vasilyeva et al., 2015) contaminated by infected hosts (Angers et al., 2009; Davenport et al., 2018; Mathiason et al., 2006; Vasilyeva et al., 2015). The difficulty in managing CWD arises in part from the biological complexities of the disease (Haley & Hoover, 2015; C. J. Johnson et al., 2011; Potapov et al., 2013, 2015; Vasilyeva et al., 2015) including the unique characteristics of the prion itself. Prions can persist in the environment for extended periods, accumulating in soil and vegetation (Angers et al., 2009; Mathiason et al., 2006; Nalls et al., 2013; Tamgüney et al., 2009), and remain infectious for years—creating long-lasting environmental reservoirs (Miller et al., 2004). Currently, there are no known treatments for infected animals, and no practical methods exist for decontaminating natural environments (Bartz et al., 2024), presenting a significant challenge to disease mitigation efforts.

**Figure 1.**
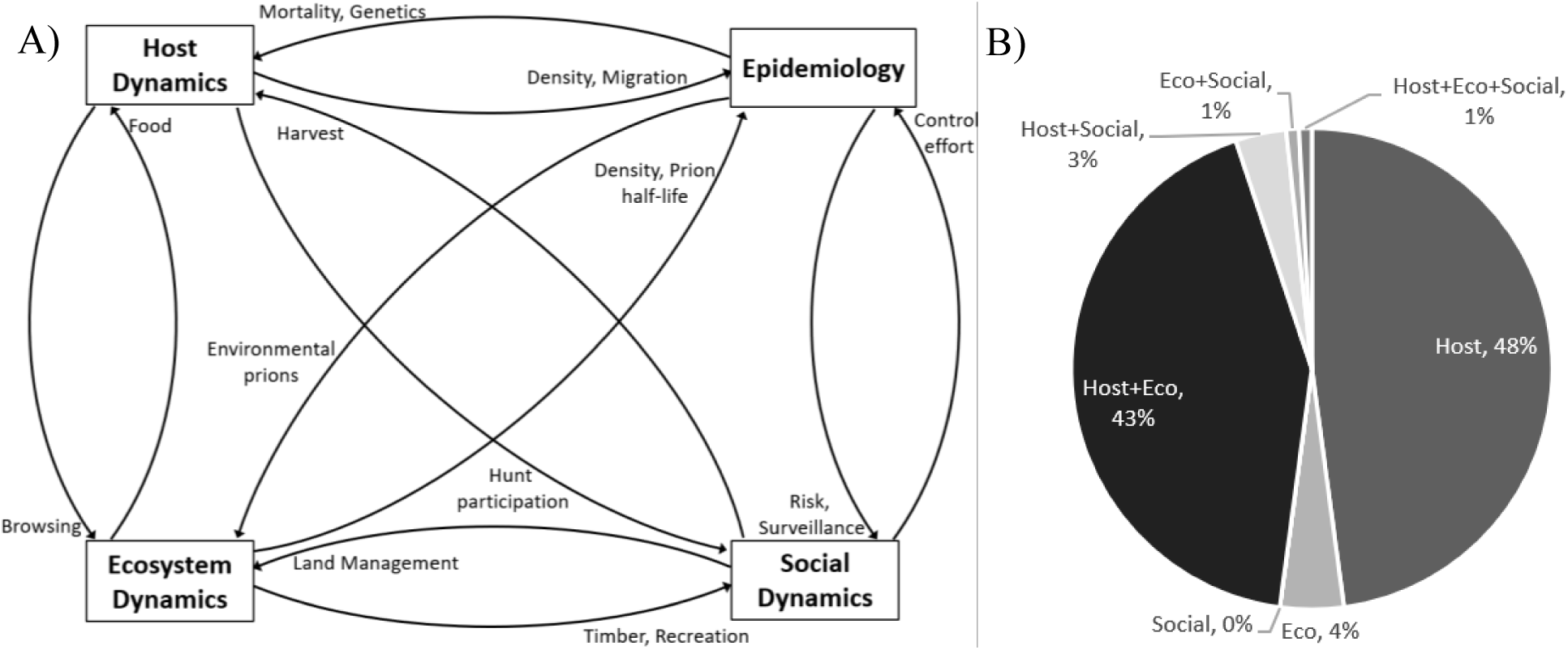
A simplified representation of how systems thinking can be applied to chronic wasting disease (CWD), highlighting selected feedbacks among key system components: CWD epidemiology, host dynamics, ecosystem dynamics, and social dynamics. *Panel A*: Arrows illustrate endogenous feedbacks between the four areas, with representative examples of interactions. Host dynamics include variables at the population level. Ecosystem dynamics encompass ecological factors beyond the individual or population level, as well as the accumulation of environmental prions. Social dynamics capture how humans influence and are influenced by the system through behaviors and socioeconomic factors. *Panel B*: A pie chart summarizes findings from a review of 103 CWD epidemiological models in the literature, models were categorized based on the system components included beyond epidemiology (i.e., host, ecosystem, social dynamics). Models were further categorized based on whether these were incorporated endogenously, exogenously, or mechanistically (refer to Appendix Table S1). This literature review expands on the work of Winter and Escobar (2020) by including studies published between 2018 and 2023.

In addition to the biological complexities of CWD, social and political factors have significantly influenced state [and provincial] wildlife agency responses to the disease, thereby shaping its trajectory over time (Bishop, 2004; Heberlein, 2004; Holsman et al., 2010; Holsman & Petchenik, 2006; Landon et al., 2023; Meeks et al., 2022; Pattison-Williams et al., 2020; Thompson et al., 2023; Vaske, 2010; Vaske et al., 2021). The primary hosts at risk from CWD include high-value game species (e.g., white-tailed deer [Odocoileus virginianus]; Macaulay, 2016), which are not primarily managed to maintain ecological balance but also for their recreational, cultural, and economic importance. As a result, wildlife managers operate under heightened scrutiny from a diverse set of stakeholders with varying perspectives regarding the appropriate management of deer populations (Bradshaw et al., 2021; Cooney & Holsman, 2010; Holsman et al., 2010). Institutional frameworks like the North American Model of Wildlife Conservation have historically reinforced a strong link between wildlife managers and hunters, who were traditionally seen as conservation allies and stewards (Geist et al., 2001; Heffelfinger et al., 2013; Organ et al., 2012). However, as hunter license purchasing and participation in the field declines, their capacity as a management tool continues to diminish (Mohr et al., 2025), while their influence over management decisions remains disproportionately high (Berl et al., 2022; Bruskotter et al., 2022; Morales et al., 2023; Peterson & Nelson, 2017). This imbalance in stakeholder power—where a shrinking demographic continues to exert substantial influence— can marginalize the values and preferences of other stakeholder groups and create pressure on agencies to adopt policies that primarily reflect hunting-related interests with less focus on the ecological health of the system (Decker et al., 1996; Marshall et al., 2007).

To date, most CWD models have concentrated on host population and disease dynamics (Figure 1B). These models have been instrumental in identifying key drivers of disease processes (Belsare et al., 2021; Hefley et al., 2017; Jennelle et al., 2014; Potapov et al., 2013; Storm et al., 2013) and in highlighting population control as a central strategy for reducing disease prevalence and limiting geographic spread (Association of Fish & Wildlife Agencies, 2018; Jennelle et al., 2014; Mysterud & Rolandsen, 2018; Potapov et al., 2016; Storm et al., 2013; Wasserberg et al., 2009). However, based on a review we conducted of the literature (Figure 1; refer to Appendix *Section 1* for methods), only 5% of these models incorporate social or political factors into management decision-making, even though such influences can substantially affect both the implementation and the effectiveness of disease control strategies (Lee et al., 2017; LeJeune et al., 2024; Osi & Ghaffarzadegan, 2025). In cases where human dimensions have been considered (Appendix Table S1), they are typically treated as static inputs—such as fixed harvest rates— failing to reflect the dynamic nature of stakeholder behavior and policy responses. Management interventions are often narrowly framed, manipulating only a few exogenous parameters (e.g., adjusting mortality in specific age or sex classes), without accounting for potential feedbacks such as stakeholder satisfaction, compliance, or political feasibility. As a result, existing models can miss important complexities of the CWD system resulting in management actions that are unfeasible or have unforeseen consequences (Heberlein, 2004). This gap presents a compelling opportunity for the application of systems thinking and SD modeling, which can illuminate the intricate interdependencies among epidemiological, ecological, and human dimensions that shape disease dynamics and management outcomes (Figure 1A).

Here we present a SD model for CWD that integrates a core host-pathogen model with established SD formulations to capture some key human dimensions influencing disease dynamics and management. Model structures were developed using a combination of published literature and a participatory modeling process involving workshops with diverse stakeholder groups and domain experts (CWD Response Plan Review Committee: Input Document, 2022). The model was parameterized using data from the U.S. state of Wisconsin (WI), and model predictions for key metrics and system behaviors were compared to findings from existing studies where available. This holistic approach enabled the integration of both quantitative and qualitative knowledge across ecological, epidemiological, and human dimensions into a unified modeling approach. Notably, the model provides the first estimates for genetic and environmental transmission parameters and reveals feedback loops spanning biological, ecological, and human dimensions. While the model is not designed to prescribe specific policy recommendations, it offers decision-makers a tool to explore the outcomes of various proposed intervention strategies on both disease dynamics and stakeholder-relevant outcomes, thereby supporting the evaluation of potential policy effectiveness and implementation tradeoffs. Furthermore, an extension of this work explicitly integrated the model into a broader decision-making framework to facilitate policy scenario evaluation by state [and provincial] agency managers (CWD Response Plan Review Committee: Input Document, 2022).

## Methods

### Data

The Wisconsin Department of Natural Resources (WDNR) provided time-series data on both white-tailed deer metrics and CWD surveillance, which were used for model estimation. The following data were used in estimating the posterior distributions for model parameters: (1) fawn-to-doe ratios, (2) the ratio of harvested yearling bucks to total harvested bucks, (3) the ratio of harvested yearling does to total harvested does, (4) post-hunt population estimates, (5) the number of CWD-positive deer by age and sex cohort, and (6) the frequency of the *S* allele at the coding polymorphism 286G/A (G96S). WDNR began reporting all deer statistics at the county level in 2017, supplementing the previously-used deer management zones (DMZs) which aggregated multiple counties (Table S16). DMZ-level data were available from 2007 to 2022 for fawn-to-doe ratios and yearling-to-total harvest ratios for both bucks and does. County-level data for these metrics were available from 2017 to 2023 (Appendix *Section 3.2.1.2*). Post-hunt deer population estimates by county were derived from sex-age-kill (SAK) (Roseberry & Woolf, 1991) and accounting models (Rolley, 2013), spanning 2007-2023 (Appendix *Section 3.2.1.3*). CWD surveillance data, including the number of deer testing positive within each age and sex cohort, were also available at the county level from 2002 to 2023 (Appendix *Section 3.2.3.1*). Genotype frequencies at the 286G/A (G96S) polymorphism were available for five years (2004, 2017–2020) and aggregated for Iowa, Dane, and Grant counties—areas corresponding to the original CWD detection zone in WI (Appendix *Section 3.2.1.1*).

County-level time series for antlered and antlerless deer harvest (Appendix *Section 3.2.2*) spanning 2000 to 2023, were also provided by the WDNR. These data, along with the total number of deer tested for CWD in each county, were used to constrain estimates of the proportion of deer harvested and tested annually during model fitting. Additionally, WDNR published a county-level time series for the Winter Severity Index (WSI) from 2000 to 2020, which was used as an environmental input to account for potential impacts on deer recruitment and mortality.

WDNR also provided longitudinal, individual-level data on hunting license purchases from 2005 to 2023. These were aggregated to the county level and used to infer annual hunter activity states and lapses in license purchasing. Following Berl et al. (2025), individuals purchasing a license in a given year were classified as “active” hunters, while those who failed to purchase a license were classified as “inactive” and subdivided further into five states based on the number of consecutive years of inactivity (1, 2, 3, 4, or ≥5 years). These classifications were then used to identify recruitment, retention, and reactivation (R3) events at the level of each individual hunter, based on the year of first license purchase, successive years of license purchasing, or a license purchase by a previously inactive hunter, respectively. R3 events were aggregated as transition probabilities between hunter activity states by county and year and were used alongside counts of active and inactive hunters in model estimation. Recruitment could not be reliably distinguished from reactivation within the first five years of data, and observations of hunter inactivity could only begin accumulating after 2005. Consequently, these variables were treated as missing for the relevant initial time spans (refer to Appendix *Section 3.2.5*). Publicly available data from the U.S. Census Bureau Population Estimates Program (U.S. Census Bureau, 2010, 2020, 2024) were used to construct a time series of estimated total population by county from 2005 to 2023. These data were used as a model constraint to determine the maximum pool of potential hunters that could still be recruited, after subtracting the number of active and inactive hunters (Appendix *Section 3.2.5.3*).

### Model Development

We used a participatory modeling approach by facilitating a series of workshops with stakeholder groups and domain experts (CWD Response Plan Review Committee: Input Document, 2022) to integrate knowledge and data across ecological, epidemiological, and social dimensions influencing CWD dynamics and its management in WI white-tailed deer (Gray et al., 2018). This participatory process was complemented by a review of existing literature to inform the model structure and parameterization. Model development was conducted using Vensim software (Ventana Systems Inc., 2025), a widely used SD tool that facilitates the construction of compartment-based models through graphical representations of stocks (i.e., compartments), flows, and feedback loops. Vensim supports the development of systems of ordinary differential equations that characterize dynamic behavior over time (Eberlein & Peterson, 1992).

Many differential equation-based compartmental models have been developed to capture the progression of CWD infection in cervids (Figure 1B)—from initial exposure to infectious stages and eventual mortality (e.g., Almberg et al., 2011; Miller et al., 2006; Potapov et al., 2013). These models served as valuable references for selecting appropriate functional forms for disease dynamics. By combining this foundation with stakeholder insights and input from wildlife biologists and epidemiologists, we developed a model that endogenously simulates host population dynamics, disease transmission, and human behavioral factors influencing disease outcomes.

Given the complexity and scope of the resulting model, we provide a high-level overview here, with detailed descriptions of the model structure, equations, and parameterization procedures presented in Appendix *Section 2.2*. Model parameters were estimated from WI data, where possible, or informative priors from existing literature and expert knowledge (Appendix *Section 2.2*, Table S17).

### Model Structure

#### Host and Epidemiology

At the core of our model is a *Susceptible–Infected–Clinical* (*SIC*) compartmental structure. This system of differential equations tracks the number of susceptible (i.e., healthy), infected (both exposed and infectious), and clinical (i.e., terminal stage prior to death) individuals within the deer population over time. Infections occur through both direct (animal-to-animal) and indirect (environmental-to-animal) transmission pathways. As mentioned previously, the SD modeling approach refers to compartments as *stocks*, and the rates at which individuals transition between stocks as *flows*. The model tracks the deer population within spatial units termed *regions* (indexed as *j*), defined here as counties (Equation 1). Within each region, individuals in each stock are further stratified by age and sex to allow for cohort-specific dynamics (Figure 2). Age classes (*k*) were defined as follows: 1) fawns (0-1 years), 2) yearlings (one-year olds), 3) two-year olds, 4) three-year olds, 5) four-year olds, 6) five-year olds, or 7) six-year olds and older (6+ years). Sex (*l*) was categorized as either *buck* or *doe* for all age classes, including fawns. This disaggregation allowed the older buck population to be assessed, a key metric for WDNR when considering management options. Later, it permitted comparison of simulated prevalence to age-structured surveillance data.

**Figure 2.**
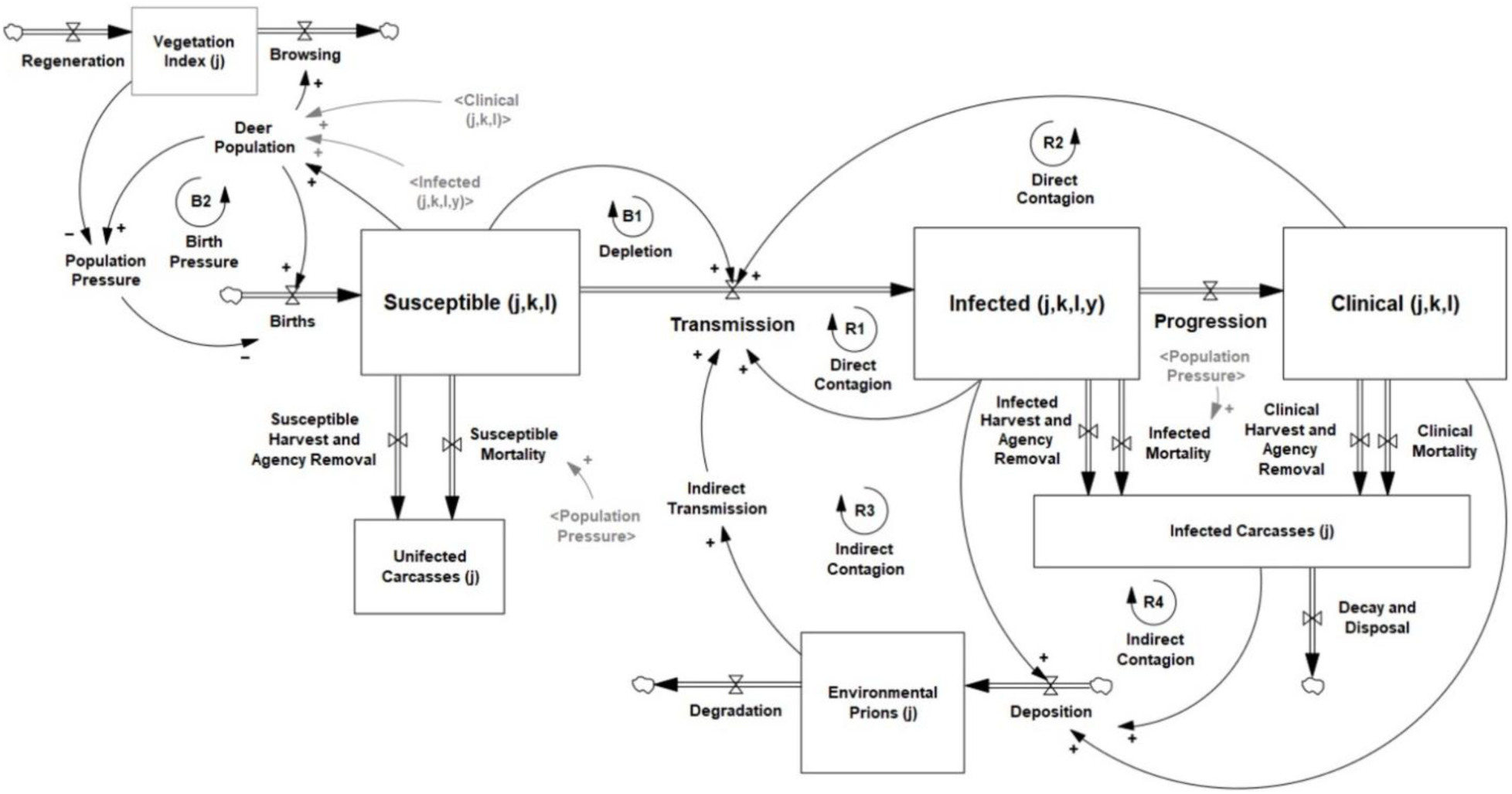
A stock and flow diagram representing the host–disease dynamics at the core of the system dynamics model for chronic wasting disease (CWD) in Wisconsin white-tailed deer. The dynamics that define the susceptible, infected, and clinical deer stocks are shown in addition to the environmental prions and vegetation index. Stocks are depicted with boxes and pipes represent flows either into or out of a stock; clouds represent limitless sources or sinks. Stocks are arrayed by region (j), age class (k), sex (l), infection stage (y). Other causal relationships are represented with either positive or negative arrows from origin to terminal variable. A positive relationship means the same direction of change (i.e. if A increases, B also increases), while a negative relationship means the inverse direction of change (i.e. if A increases, B decreases). Feedback loops are indicated by arrows representing the direction of the loop and labels for either reinforcing (R) or balancing (B) with descriptive name. Variables that are repeated more than once on the diagram for ease of connection to variables located far from the original representation are shown in gray.

To capture disease progression in greater detail, the *Infected* compartment was subdivided into six stages (*y*: 1–6) (Figure 2). The first three stages represent exposed but non-infectious individuals, while the final three stages correspond to prion-shedding, infectious individuals. As supported by prior studies (Almberg et al., 2011; Edmunds et al., 2018; Hamir et al., 2008; Jennelle et al., 2014; Wasserberg et al., 2009; Williams, 2005), individuals transition to the *Clinical* stock shortly after the onset of observable signs of CWD. This final stage is characterized by markedly different mortality rates and behavioral changes, necessitating its separation from the general infected population in the model.

Individuals are removed from each stock through a stock-specific natural mortality rate. A background mortality rate 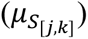 is applied to the *Susceptible* stock and can vary by region and age, while a higher mortality rate (*µ_C_*) is applied to the *Clinical* class to reflect CWD-induced mortality. An additional mortality hazard (*ψ*_*I*_) can be applied to all *Infected* deer to account for pre-clinical CWD-associated mortality (Appendix *Section 2.2.1.1*). Deer are also removed from all stocks via hunter harvest at stock-specific rates (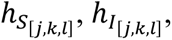 and 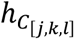 for *Susceptible*, *Infected*, and *Clinical* deer, respectively) or through agency removals (area-specific, targeted removal of deer), with separate rates for *Susceptible* and CWD-positive deer (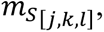 and 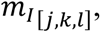 respectively). These harvest and removal rates vary by region, age, and sex (Figure 2; Equation 1; Appendix *Section 2.2.1.3*). We assume hunters generally avoid taking fawns when recognizable and thus scale down the effective harvest rate of fawns relative to older age classes (Appendix *Section 2.2.1.3*; Eq. S15).

Recruitment occurs through the addition of fawns to the *Susceptible* stock, based on the product of a region-specific, density-dependent birth rate (*α*_[*j*]_) and the number of reproductive does (*D*_[*j*]_). Birth is assumed to occur in a pulse at the model year midpoint, aligned with the peak fawning season of June in WI (Jacques et al., 2007), with a 1:1 sex ratio. Individuals age into subsequent classes annually at the beginning of the birth pulse, up to an accumulator class that includes all deer aged six years and older (Appendix *Section 2.2.1.1*). A simplified dispersal mechanism allows deer in any stock to move between regions at an instantaneous rate 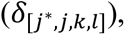 which varies by age, sex, and inter-regional distance (Appendix *Section 2.2.1.2*).

*Susceptible* individuals become *Infected* through both direct (i.e., infected cervid to healthy cervid) and indirect (i.e., environmental CWD prion to healthy cervid) transmission events (Appendix *Section 2.2.3*). These pathways collectively determine the total force of infection (*λ*_[*j*,*k*,*l*]_). Direct and indirect transmission rates are governed by transmission probabilities (*β* parameters), which represent a baseline value for the product of the probability of transmission given contact and contact rate. These *β* parameters are further modulated by density dependence, deer age, sex, and behavioral factors (Appendix *Section 2.2.3.2*). The effect of deer density is decomposed into a cross-sectional effect (reflecting variation in density across regions) and a longitudinal effect (reflecting density variation within each region over time). Decoupling these effects helps mitigate high uncertainty due to measurement limitations in deer population size and range (Appendix *Section 2.2.3.2*). While these effects interact, they have distinct signatures, producing variations in transmission across regions, covariation of transmission with population, and differential responses to antlerless harvest.

After infection, individuals progress through six disease stages in the *Infected* stock according to a stage-specific progression rate (*ρ*_[*j,y*]_). Deer spend approximately 30 weeks in the exposed (non-infectious) stages and 34 weeks in the infectious stages before transitioning into the *Clinical* stock (refer to Appendix *Section 2.2.3.3* for details). All individuals in the final three stages of the *Infected* stock, along with those in the *Clinical* stock, are considered infectious. The total number of infectious individuals in a region (*F*_[*j*]_), contributes to both direct transmission and environmental contamination.

Susceptible and infected deer can disperse between different regions (Appendix *Section 2.2.1.2*); however, we assume that clinical deer do not disperse due to severely impaired motor function (Almberg et al., 2011; Edmunds et al., 2018; Hamir et al., 2008; Jennelle et al., 2014; Wasserberg et al., 2009; Williams, 2005). The influx of susceptible deer into a given region *j* is calculated as the sum across all origin regions *j*⁎ of the product of the instantaneous dispersal rate 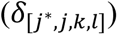 for each age and sex class and the number of susceptible deer 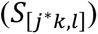 in the origin region (Eq. 1.1). Likewise, the outflow of susceptible deer from region *j* is computed as the sum across all destination regions *j*⁎ of the product of the dispersal rate from region *j* to region 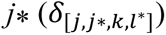 and the number of susceptible deer in the origin region (*S*_[*j*,*k*,*l*]_). Infected deer disperse using the same mechanics, but with the number of infected individuals (*I*_[*j*,*k*,*l*,*y*]_) instead of susceptible individuals (Eq. 1.2).

Environmental prions accumulate in each region (*V*_[*j*]_) through shedding by infectious deer at a per-capita rate (*ε_F_*) and through infectious carcasses left on the landscape 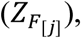 with a multiplier (*ε_Z_*) applied to represent the greater prion load from carcasses compared to live shedding (Figure 2; Equation 1.4; Appendix *Section 2.2.3.4*). Carcasses may degrade naturally due to scavenging and decay (*t_Z_*) or be removed through management actions (*ς*_Ψ_). The environmental prion stock is also depleted via natural degradation or reduction in bioavailability at a constant rate (τ).

The model includes a genetic component based on the *S* allele at the 286G/A (G96S) polymorphism, which has been linked to delayed CWD progression (Denkers et al., 2024; Johnson et al., 2011). The frequency of the *S* allele is treated as a dynamic variable that may influence disease transmission (i.e., probability of infection given contact) and progression rates (i.e., length of time deer live post infection event) via separate parameters. This flexible structure allows the data to inform the impact of host genetics on disease dynamics during model estimation (Appendix *Section 2.2.3.2* and *2.2.3.3*). Note that the estimated genetic effect on transmission rate could be due to a change in the infectiousness of the infected deer or the resistance of the susceptible deer while the effect on the progression rate occurs after transmission occurs and is estimated separately for the exposed and infectious stages.

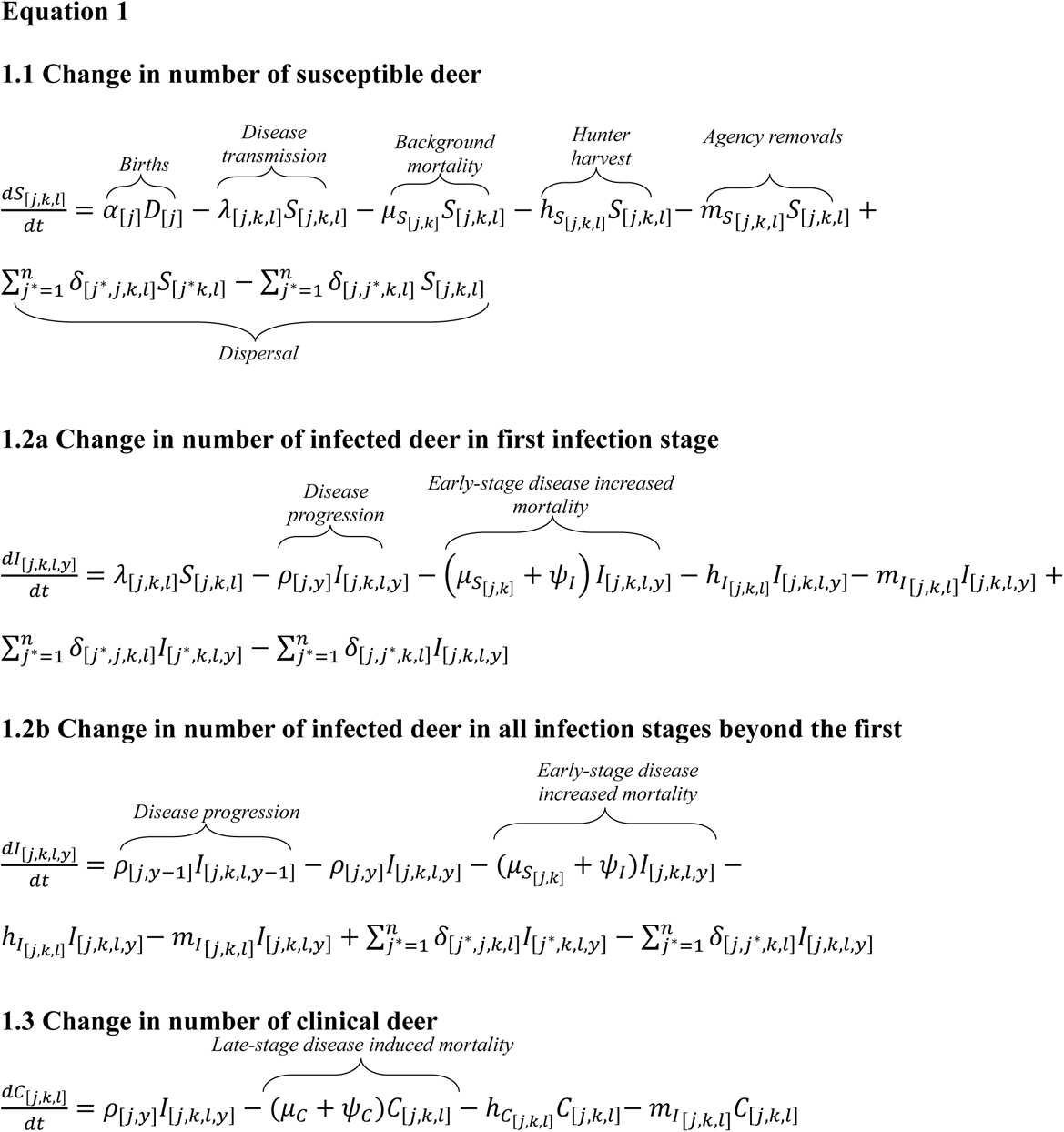

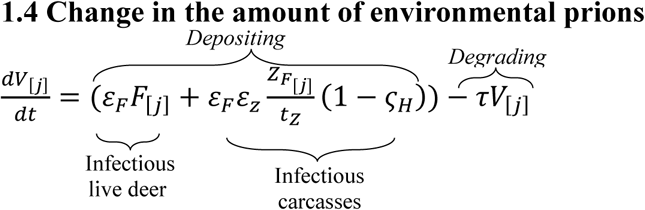

#### Ecosystem

Broader ecosystem-level processes are incorporated into the model through a dynamic vegetation stock variable (*G*_[*j*]_), which represents changes in available habitat for deer relative to initial starting conditions in each region (Appendix *Section 2.2.2*). This is intended to capture in a simple way the carrying capacity limitation of forage adequacy, and the corresponding rebound in vegetation (and later reproduction) that could occur from release of browsing pressure due to density reduction from harvest changes. It also serves as a directional proxy for understory health, which is of interest to some stakeholders.

The vegetation stock increases through seedling regeneration and decreases due to deer browsing (Eq. 2). Regeneration increases the stock at a rate of one over the seedling regeneration time (*t_R_*) however, this growth rate slows as the vegetation stock approaches a theoretical maximum value (*G_max_*) to capture limits on vegetation levels unrelated to deer browsing. Vegetation loss from browsing is modeled as a fraction of the current vegetation level. The fraction is dictated by the difference between the reference population pressure 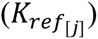 and the pressure on vegetation from the current weighted population 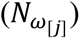 –which adjusts for the lower ecological impact of fawns –as well as the reference per deer browsing rate (*B_ref_*). The reference browsing rate is defined to exactly balance seedling regeneration when the pressure created by the deer population is equal to a region’s reference population pressure. Additionally, external resources such as agricultural browse can supplement natural vegetation, effectively reducing population pressure within a region (Appendix *Section 2.2.1.1*).

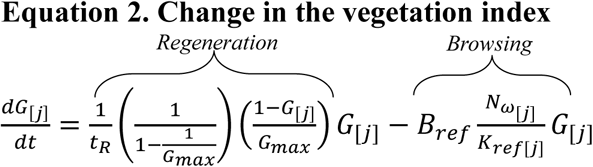

#### Human Dimensions

The human dimensions of the model focus on the dynamics of hunter behavior, who play a central role in the management of CWD. A mechanistic approach was used to represent the hunter population explicitly as a stock within a dedicated submodel (Figure 3), which interacts with the broader *SIC* model. This structure defines an upper bound on achievable harvest and enables evaluation of recruitment, retention, and reactivation (“R3”) strategies as potential levers to influence system outcomes.

**Figure 3.**
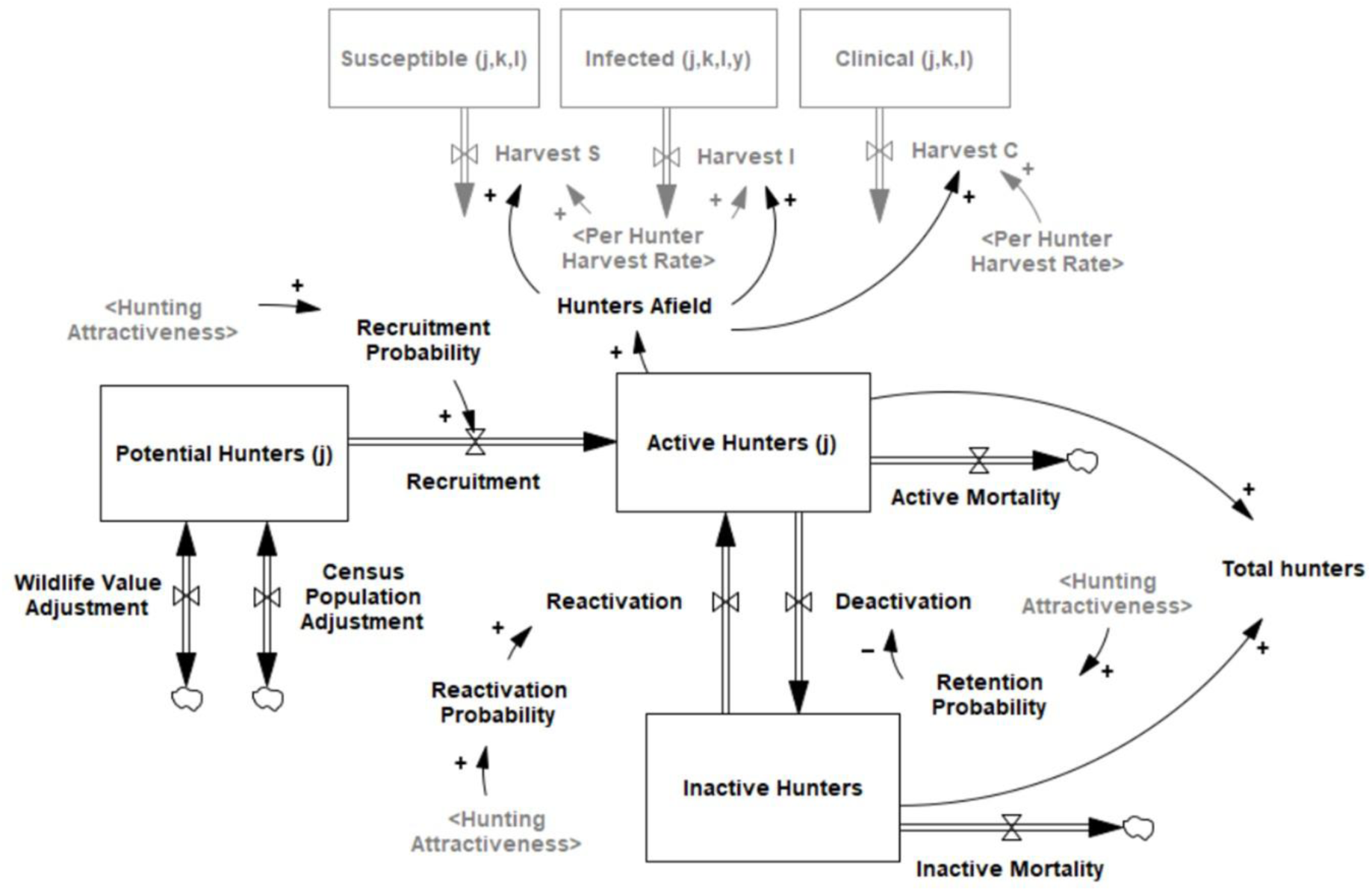
A stock and flow diagram depicting the hunter stock dynamics within the Wisconsin white-tailed deer chronic wasting disease (CWD) system. The dynamics that define the potential, active, and inactive hunter stocks are shown in addition to the connection to the susceptible, infected, and clinical deer stocks. Stocks are depicted with boxes and pipes represent flows either into or out of a stock; clouds represent limitless sources or sinks. Stocks are arrayed by region (j), age class (k), sex (l), infection stage (y). Other causal relationships are represented with either positive or negative arrows from origin to terminal variable. A positive relationship means the same direction of change (i.e. if A increases, B also increases), while a negative relationship means the inverse direction of change (i.e. if A increases, B decreases). Variables that are repeated more than once on the diagram for ease of connection to variables located far from the original representation as well as stocks that are not fully represented with flows are shown in gray.

Within the hunter submodel, *Active hunters* 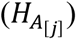 region are defined as residents who have purchased any license type to hunt deer in a region. *Inactive hunters* 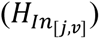 are those who have previously held a license but did not purchase one for the current year. Each year, active hunters can be drawn from any combination of three sources: recruitment from the pool of *Potential hunters* 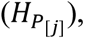 retention of the previous year’s *Active hunters*, or reactivation of individuals from the *Inactive hunter* stock (Equation 3.1). Potential hunters are defined as individuals in the region who have never purchased a hunting license but hold wildlife value orientations supportive of hunting (i.e., traditionalists and pluralists; Manfredo, Teel, et al., 2021). This group is estimated as the total number of residents (*P*_[*j*]_) who would consider hunting based on their wildlife values (*Λ*_[*j*]_; refer to Appendix *Section 2.2.4.3*)—less those already recruited (active + inactive hunters)—with priors drawn from past surveys (Dietsch et al., 2018; Manfredo, Berl, et al., 2021) and fitted to county-level data (Equation 4).

Recruitment into the active hunter stock occurs at a region-specific transition probability 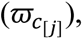 while retention is modeled as the probability that a previously active hunter remains active the following year 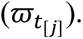 Those who fail to renew their license transition into the first stage of the inactive hunter stock (Equation 3.2a), and since hunters can only move once per year, this transition probability is equal to one minus the retention probability. Reactivation probabilities decline with each consecutive year spent inactive (Hinrichs et al., 2020); thus, the inactive stock is subdivided into five stages (*v*), with stage five capturing individuals who have been inactive for five or more years (Figure 3), beyond which reactivation is highly unlikely (Hinrichs et al., 2020). Finally, hunters in both active and inactive stocks are subject to mortality at continuous rates (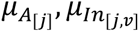 respectively), calculated using the age structure of license holders and CDC-derived age-specific mortality rates (U.S. Department of Health and Human Services, 2024) (refer to Appendix *Section 3.2.5* for details).

Additional social and behavioral factors influencing hunter license purchasing are captured using a formulation for product attractiveness (Sterman, 2000), which combines multiple factors to influence the level of attractiveness of a product (in this case, a hunting license). For the WI context, these factors were identified from literature sources as well as discussions with a broad range of stakeholders including government agencies (e.g., WI DNR, US Department of Agriculture), hunting and policy non-governmental organizations (e.g., WI Wildlife Federation and Sporting Heritage Council), business (e.g., Whitetails of WI), and the interest of Tribal Nations (e.g., Oneida Nation). Formulating the attractiveness of hunting (Å_[*j*]_) to active, inactive, and potential hunters (Appendix *Section 2.2.4.3*) was of particular interest to stakeholders due to its implications for license purchasing, agency revenue, harvest levels, and ultimately, the size of the cervid population and the prevalence of CWD infections (Figure 4).

**Figure 4.**
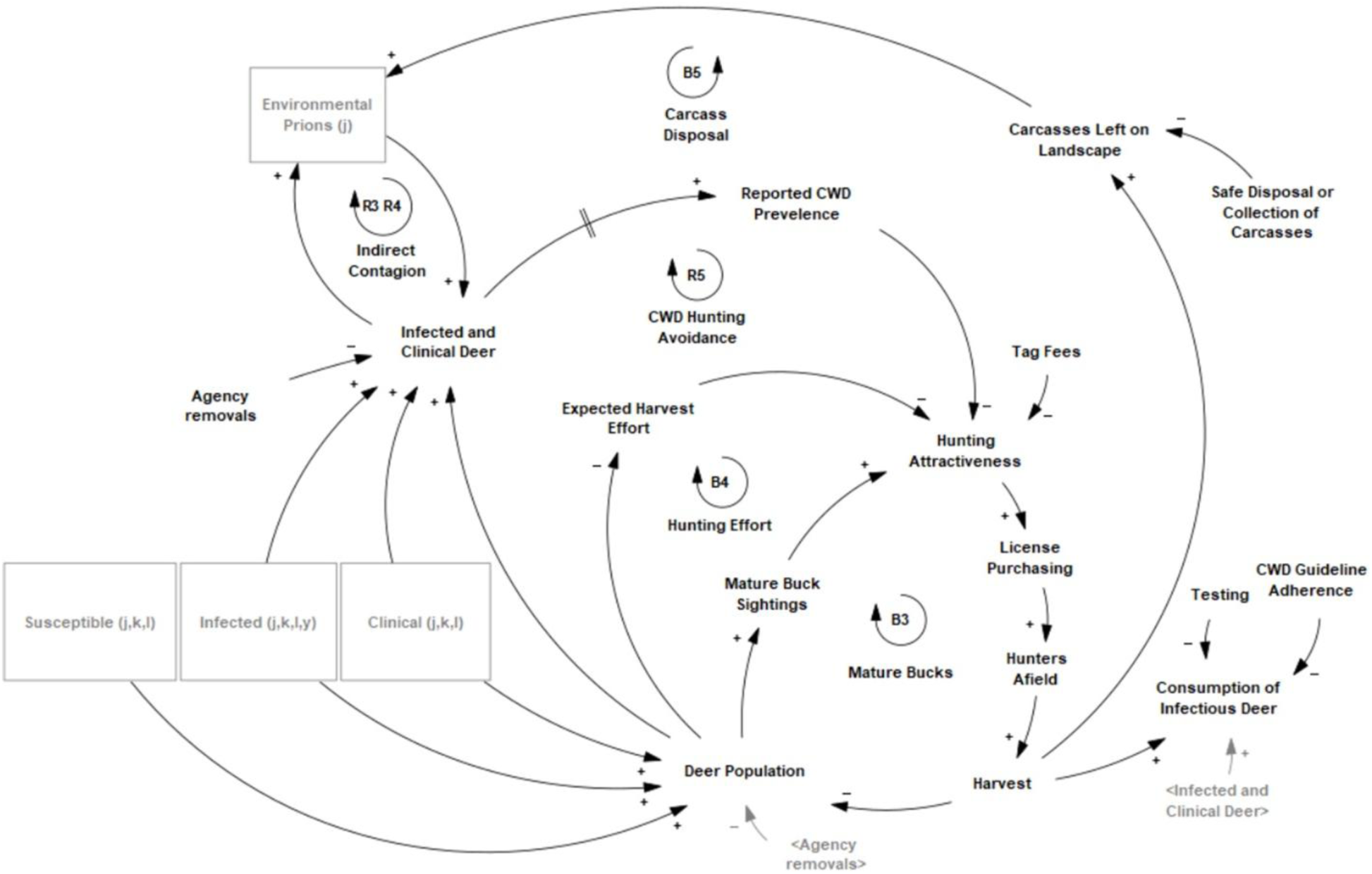
Causal loop diagram illustrating the feedbacks between social dynamics and deer–chronic wasting disease (CWD) processes, along with management interventions that can be modified to assess their influence on the system. Dynamic human dimensions that influence the attractiveness of hunting are shown in relation to the deer population stocks— Susceptible, Infected, and Clinical—and Environmental Prions, as depicted in Figure 2. Stocks are depicted with boxes and pipes represent flows either into or out of a stock. Stocks are arrayed by region (j), age class (k), sex (l), infection stage (y). Other causal relationships are represented with either positive or negative arrows from origin to terminal variable. A positive relationship means the same direction of change (i.e. if A increases, B also increases), while a negative relationship means the inverse direction of change (i.e. if A increases, B decreases). Feedback loops are indicated by arrows representing the direction of the loop and labels for either reinforcing (R) or balancing (B) with descriptive name. Variables that are repeated more than once on the diagram for ease of connection to variables located far from the original representation as well as stocks that are not fully represented with flows are shown in gray.

The R3 transition probabilities governing the number of active hunters—recruitment 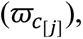 retention 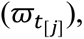 and reactivation 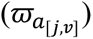— can be modified by the hunting attractiveness within each region (Å_[*j*]_). When Å_[*j*]_ is greater than one, these transition probabilities increase above their model-predicted baseline values, reflecting a higher likelihood of individuals entering or remaining in the active hunter population through purchasing a hunting license.

Conversely, when Å_[*j*]_ is less than one, recruitment, retention, and reactivation rates are proportionally reduced, resulting in fewer hunting licenses purchased. Data from WI hunter surveys indicate that approximately 5% of active hunters do not attempt to harvest deer (Wisconsin Department of Natural Resources, n.d.); therefore, the model assumes that only 95% of active hunters actually participate in field activities (Figure 3). Hunters who do attempt to harvest deer travel from their county of residence to counties where they hunt, at a constant proportion determined by a residence vs. harvest matrix. Hunters then harvest deer at a per-hunter rate specific to that county and stratified by deer age and sex. Both the movement matrix and the per-hunter harvest rates were estimated here from reported license and harvest data (Appendix *Section 3.2.5.1*). This process yields an estimate of the total number of deer harvested under regular hunting licenses (i.e., excluding agricultural damage permits) each season given hunter participation which is a product of estimated number of licenses purchased (i.e., active hunters), their mobilization to the field, and the number of antlered and antlerless deer each take.

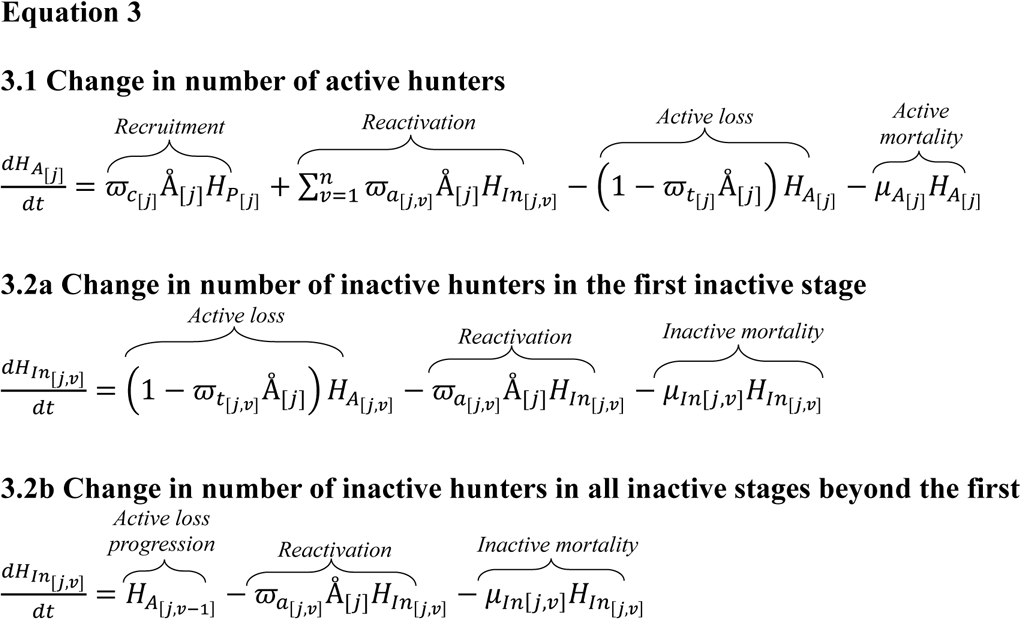

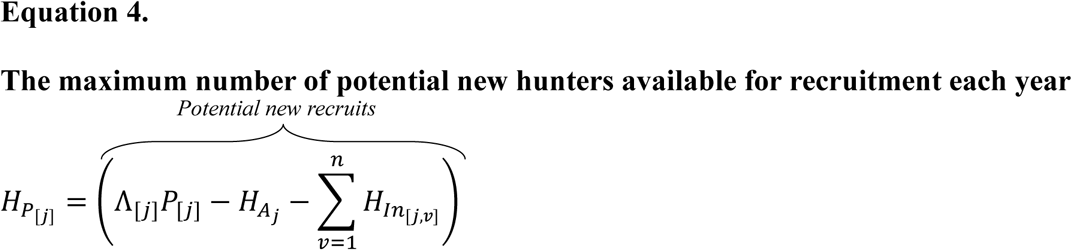

#### Management Policies

Integrating potential management actions into the social dynamics portion of our CWD systems model was crucial for exploring how the system reacts to perturbations during scenario analyses, as well as for facilitating stakeholder engagement when considering effective CWD management strategies. We identified current and past management practices, along with future potential alternatives, to incorporate as variables that can influence the model’s behavior in various ways (Price Tack et al., n.d). Both agency removal and increased hunter harvest (Figure 4) are management strategies aimed at reducing the cervid population to mitigate disease transmission by lowering contacts and decreasing the number of infected deer available. Agency removal is designed to remove a higher proportion of infected deer than would occur through regular hunter harvest. In the model, the proportion of infected deer removed is driven by factors such as the level of effort, true prevalence, and the fraction of the area affected by CWD (Appendix *Section 2.2.4.2*).

Other management actions focus on reducing the accumulation and spread of CWD prions in the environment. These include measures like encouraging safe carcass disposal by hunters and implementing carcass removal by agency personnel (Figure 4). Additionally, disease surveillance (i.e., CWD testing), can be a critical tool in guiding management decisions. The level of surveillance determines how representative the reported CWD prevalence is of the true disease intensity within the population, as well as hunters’ ability to avoid consuming CWD-positive deer (Appendix *Section 2.2.4.1*). Even if a deer is tested, hunters would also need to be aware of and adhere to the Centers for Disease Control and Prevention (CDC) recommendations, which advise that any part of a known-positive deer should not be used or consumed by humans. The availability and awareness of scientific information on CWD and its effects on behavior can potentially be influenced by outreach and education programs, which are represented in the model as a factor impacting hunter adherence with management policies (Figure 4).

### User Implementation

Model users can explore the efficacy and feasibility of various management actions by adjusting key parameters that influence social, ecological, and epidemiological dynamics. Increased hunter harvest can be implemented by directly setting a desired harvest rate (i.e., the percentage of the population to be removed via harvest) or by specifying the number of deer to be harvested per active hunter, which explores the change in harvest rate as a result of changing hunter behavior (Appendix *Section 2.2.4.3*). When the harvest rate is set directly, the model reports the corresponding number of deer harvested per active hunter for review. Agency removal is implemented by defining the percentage of a regional deer population to be removed through agency-directed efforts (Appendix *Section 2.2.4.2*). The total cost of a removal operation is then calculated based on a user-defined cost per deer removed.

Users can also adjust the fraction of harvested deer that are safely disposed of, as well as the fraction of remaining carcasses removed from the landscape, each set independently (Appendix *Section 2.2.3.4*). The number of carcasses tested for CWD can be specified as a target sampling proportion, indicating the share of observed mortalities to be tested. The CWD testing structure in the model allows the apparent prevalence from testing to be estimated as well as the true underlying prevalence within the population, allowing users to compare the divergence of these prevalence estimates under different surveillance regimes. True and apparent prevalence differ due to test insensitivity during the exposed phase. Optionally, the testing structure also permits synthetic generation of projections with measurement errors from sampling effects. Based on the number of tests conducted and a user-defined cost per test, the model calculates the total testing cost. To explore public health implications, users can set the proportion of individuals who disregard present CDC guidelines and consume positive deer, allowing them to evaluate the interaction between testing coverage and adherence to safe consumption practices on the overall risk of consuming infectious deer (Appendix *Section 2.2.4.1*). Additionally, users can specify tag fees required to harvest a deer, which directly influences the attractiveness of hunting within the model (Figure 4). Users can also modify the dispersal of deer between counties to explore how deer movement across counties can impact management outcomes.

### Model Estimation

Our model is complex and nonlinear, necessitating a conservative estimation procedure that (1) incorporates uncertainty, (2) avoids overfitting, and (3) promotes model generalizability (Rahmandad et al., 2021). To achieve this, we employed a hierarchical Bayesian approach for parameter estimation. This estimation approach allows for regional flexibility across counties while preserving the integrity of global mechanisms and incorporating prior knowledge.

Parameter estimation was performed with the Vensim software package (Ventana Systems Inc., 2025) employing the Differential Evolution Adaptive Metropolis (DREAM) sampling algorithm (Vrugt et al., 2009), a Markov chain Monte Carlo (MCMC) method, to explore the entire joint posterior distribution (Appendix *Section 4*). The process seeks to maximize the log of the posterior (minimizing a corresponding loss function), combining prior information with the likelihood of observing the data. Informative priors were applied when relevant information was available, either from published literature or expert judgement, and all priors were truncated at biologically feasible values (Table S17-S18). We estimated the demographic and epidemiological parameters for the *SIC* model (Figure 2) and the region-specific wildlife value orientation and R3 parameters for the hunter submodel (Figure 3) as separate processes.

The *SIC* model was fit using multiple time-series datasets as previously described (*Data* section). For each county *j*, the model expected values were defined as a vector, *i*, at each time, *t*, calculated as a function of the global parameters (*Ġ*) and county-specific (*CS*) parameters being estimated.

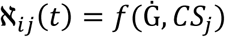

Each element of the vector ℵ_*ij*_(*t*) is derived from the deterministic system of ordinary differential equations (ODEs) specified in Equation 1. For example, post-hunt population estimates are calculated by summing the susceptible, infected, and clinical deer in each county at the time step following harvest. Additional mappings for other model predicted metrics are detailed in Appendix *Section 3.2*. To incorporate measurement error, we assumed an overdispersed Poisson distribution for CWD-positive count data and a lognormal distribution for all other datasets (Appendix *Section 4.1.1*). Model fitting included all counties where CWD has been detected, along with a subset of counties with no detections (Table S14), which helped improve the estimation of demographic parameters across the state.

The same estimation procedure was applied to fit the hunter submodel county-level timeseries data on 1) recruitment counts, 2) reactivation counts, and 3) active hunter numbers. Given the magnitude of the counts, errors in these data were modeled as lognormally distributed (Appendix *Section 4.2.1*). The model expected values can be estimated from the deterministic system of ODEs given in Equation 3. Data from 71 of 72 WI counties were included; Menominee County was excluded due to unique regulatory conditions governing hunting on Tribal lands.

### Model Testing

Model structure and parameter confirmation were conducted throughout the development process to ensure that causal relationships were logically defined and that simulations using estimated parameters produced expected behaviors. All parameter values were informed by literature and expert judgements during the participatory model building workshops. Model structure was reviewed with stakeholders to confirm that the model scope was appropriate and included the key behavioral dynamics and management considerations relevant to their questions. Throughout model construction, feedback loops were examined to verify that causal relationships were properly conserved, and extreme conditions testing was applied iteratively to check for structural errors and behavioral consistency. As an example of this validation process, we assessed the feedback loops illustrated in Figure 2. For the reinforcing loops and the disease balancing loop (B1), we examined the force of infection in relation to varying levels of susceptible deer and either environmental prions (R3, R4) or infectious deer (R1, R2) to ensure that direct and indirect transmission were functioning as intended. The demographic balancing loop (B2) was validated by analyzing the relationship between deer population relative to carrying capacity and vital rates such as birth and background mortality. Unit consistency was confirmed automatically within the Vensim software, and any discrepancies were investigated to ensure dimensional coherence across the system.

To verify that the model structure accurately captured historical trends and to identify potential systematic biases, model predictions were visually compared with observed data across multiple datasets. Quantitative goodness-of-fit metrics, specifically Theil statistics and mean absolute percent error (MAPE), were calculated for regional totals of CWD-positive deer tests and post-hunt population estimates—metrics with sufficiently large sample sizes to reduce the influence of random noise. Theil statistics decompose the total discrepancy between model predictions and observed values into three components: differences in mean (*U_m_*), differences in standard deviation (*U_s_*), and random noise (*U_c_*), providing a comprehensive assessment of model fit. MAPE values represent the average percentage deviation between predicted and observed values and reflect the proportion of variance in the data explained by the model. In social science and large-scale SD models, MAPE values under 20% suggest strong predictive performance, while values between 20% and 50% indicate reasonable forecasting ability.

Model convergence during parameter estimation was assessed using acceptance rates and the univariate Rubin/Brooks-Gelman potential scale reduction factor (PSRF; Brooks & Gelman, 1998). PSRF values approaching 1 indicate successful convergence, whereas values greater than 1.2 suggest non-convergence. For models with more than five free parameters, an average chain acceptance rate between 0.1 and 0.3 is generally indicative of well-functioning MCMC sampling (Ventana Systems, n.d.). Additionally, posterior distributions of estimated parameters were examined for signs of multimodality or boundary effects and were compared to prior distributions to evaluate parameter identifiability and the influence of prior information.

To further explore parameter stability and ensure model robustness, we conducted both local and global sensitivity analyses, as well as an out-of-sample evaluation. Local sensitivity testing was carried out using Vensim’s Sensitivity2All tool on the estimated parameters to determine their influence on model outputs and to evaluate model stability (Appendix *Sections 4.1.3.1 and 4.2.3.1*). This tool perturbs each parameter by ±10% from its mean and calculates the mean absolute deviation (MAD) between the resulting model predictions and the baseline simulation, offering a rapid assessment of the sensitivity of model outputs to changes in individual parameters (Ventana Systems Inc., 2025). Model predictive abilities were tested with an out-of-sample analysis, performed by withholding the final four years of all time-series data from the training process and reserving them for model testing. To account for the influence of parametric uncertainty on model behavior, we conducted a global sensitivity analysis using a Bayesian bootstrap approach (Clyde & Lee, 2001; McGowan et al., 2011). Given the high dimensionality of the model and the covariance among parameters, local sensitivity analysis can sometimes yield misleading inferences—such as changing mortality rates in isolation, ignoring the linkage with birth rate, causing a population collapse. In contrast, global sensitivity analysis allows for parameter variation within the bounds of the posterior distribution while preserving realistic covariation among parameters. This ensures that only behaviorally plausible combinations of parameters—those that would have been accepted during model fitting—are used in simulation, thus avoiding biologically implausible outcomes like deer extinction from unrealistic mortality-recruitment pairs. Therefore, during our simulation experiments we drew from the joint posterior distributions generated by MCMC, allowing the model to capture the implications of parameter uncertainty in a statistically coherent and ecologically meaningful manner.

## Results

### Model Dynamics

The host-epidemiological core of the model contains six feedback loops of interest, four reinforcing loops (i.e., positive feedback loop) and two balancing loops (i.e., negative feedback loop), as shown in Figure 2. The reinforcing loops arise as the pool of infected and clinical individuals increases, which in turn enhances direct transmission (R1 and R2, Figure 2) and indirect transmission through the accumulation of prions in the environment (R3 and R4, Figure 2). Conversely, the disease balancing loop occurs when the available pool of susceptible individuals decreases, leading to a reduction in new infections (B1, Figure 2). The demographics balancing loop (B2, Figure 2) captures how an increase in the deer population results in decreased recruitment and survival as the population approaches the regional maximum capacity (Appendix *Section 2.2.1.1*). This loop is interrelated to the ecosystem layer through the vegetation index, which responds dynamically to population pressure. As more deer consume vegetation (Eq. 2), the availability of resources decreases, limiting reproduction and survival, and in turn impacting the host population (Figure 2). The interactions between ecosystem functions, host population dynamics, disease spread, and human decisions create a feedback system that allows these variables to evolve dynamically within the model (Figures 2–4).

Interdependencies between hunters, the harvest process, and deer populations, introduces an additional three balancing loops and one reinforcing loop (Figure 4). All of these loops operate through their influence on the attractiveness of hunting, which directly influences the number of hunters that purchase licenses and subsequently the number of deer harvested (Eq. 3)– assuming similar mobilization and per hunter harvest. The balancing loops have the potential to limit disease growth by affecting hunting effort, mature buck availability, and carcass disposal (B3-B5, Figure 4). The hunting effort balancing loop (B3, Figure 4) reflects the negative relationship between expected harvest effort and hunting attractiveness. As hunting becomes more attractive, harvest increases, which reduces the deer population, increasing the expected harvest effort and subsequently decreasing hunting attractiveness. Similarly, an increase in harvest results in fewer mature bucks, reducing sightings and the attractiveness of hunting, which in turn decreases harvest—closing the mature buck balancing loop (B4, Figure 4). Here we assume that the intermediary per-hunter factors of participation and success that link license purchase to deer harvest participation remain the same. However, if active hunter participation changes (i.e., less hunters mobilize to go afield, or fewer hunter desire an antlerless deer) or success changes (i.e., more skilled hunters achieving higher per-hunter harvest) then the relationship between number of licenses purchased and number of deer harvested will also change.

The carcass disposal balancing loop (B5, Figure 4) functions through indirect transmission rather than directly through the deer population. Increased hunter harvest leads to more cervid carcasses left on the landscape. Infected carcasses deposit prions into the environment, increasing infection rates and consequently reported CWD prevalence (B5, Figure 4). Increasing safe disposal of hunter harvested deer subsequently decreases carcasses on the landscape, providing a management action with the potential to reduce environmental prion buildup. If hunters are sensitive to the perceived CWD prevalence, this will decrease the attractiveness of hunting and subsequent harvest, which reduces the number of carcasses left on the landscape. The CWD hunting avoidance reinforcing loop (R5, Figure 4) is driven by hunter sensitivity to CWD prevalence. As reported CWD prevalence increases, hunters may choose not to hunt, leading to decreased harvest, increased deer populations, and more infected and clinical deer. This reinforcing feedback loop can accelerate disease growth but will eventually stabilize due to the disease balancing loop (B1, Figure 2) as susceptible deer are depleted.

### Model Testing

The causal relationships between model variables within the infection process loops and demographic balancing loop were preserved as expected. Both direct (Figure S7) and indirect (Figure S8) forces of infection decline to zero in the absence of infectious deer or environmental prions and reach a maximum as the number of susceptible deer diminished. Birth rates (Figure S9) and background survival probabilities (Figure S10) increased and decreased, respectively, when deer populations were below and above population pressure, a metric of deer population relative to carrying capacity. The posterior distributions for all estimated parameters approximated normal distributions, making summary statistics such as means and standard deviations appropriate for reporting in both the *SIC* model (Tables 1, S17; Figures S1-6) and hunter submodel (Tables 1, S18; Figures S17-20). All estimated parameters and model loss function achieved PSRF values below 1.2 (Tables 1, S17, S18), and mean chain acceptance rates remained below 0.3 in the hunter submodel (Figure S24). Mean chain acceptance rates were slightly higher in the *SIC* model (0.4; Figure S14) but still far from 0 (indicates algorithm is not generating viable points) or 1 (indicates steps are too conservative) and deemed acceptable given the high number of free parameters. Cumulative distribution plots for estimated parameters showed minimal variation across MCMC chains (Figure S15), with mean values stabilizing in later iterations. These diagnostics collectively indicate no evidence of nonconvergence.

Qualitative assessments indicate that model predictions generally match trends observed in the time series data used to calibrate both the *SIC* (Figures 5a–b, S11) and hunter submodel components (Figures 5c, S21), with no evident systematic bias when all available data are used for model fitting. One exception is the tendency for the model predicted post-hunt deer populations to diverge from sharp changes in time series data (Figure 5a). Additionally, the model overpredicts the number of hunters reactivating after five or more years of inactivity (Figure S21f). Theil statistics for both the *SIC* and hunter submodel attribute most of the deviation between model predictions and observed data to random noise, with smaller contributions from differences in mean and standard deviation—indicative of a good overall model fit (Figures S12, S22). Mean Absolute Percent Error (MAPE) values are generally below 20% for both models (Figures S13, S23), indicating strong forecasting accuracy, as average absolute differences between predictions and observations remain within acceptable thresholds. However, the median MAPE for reactivation counts in the fifth inactive hunter stage exceeds 20%, suggesting predictive capabilities for this subcomponent are suboptimal.

**Figure 5.**
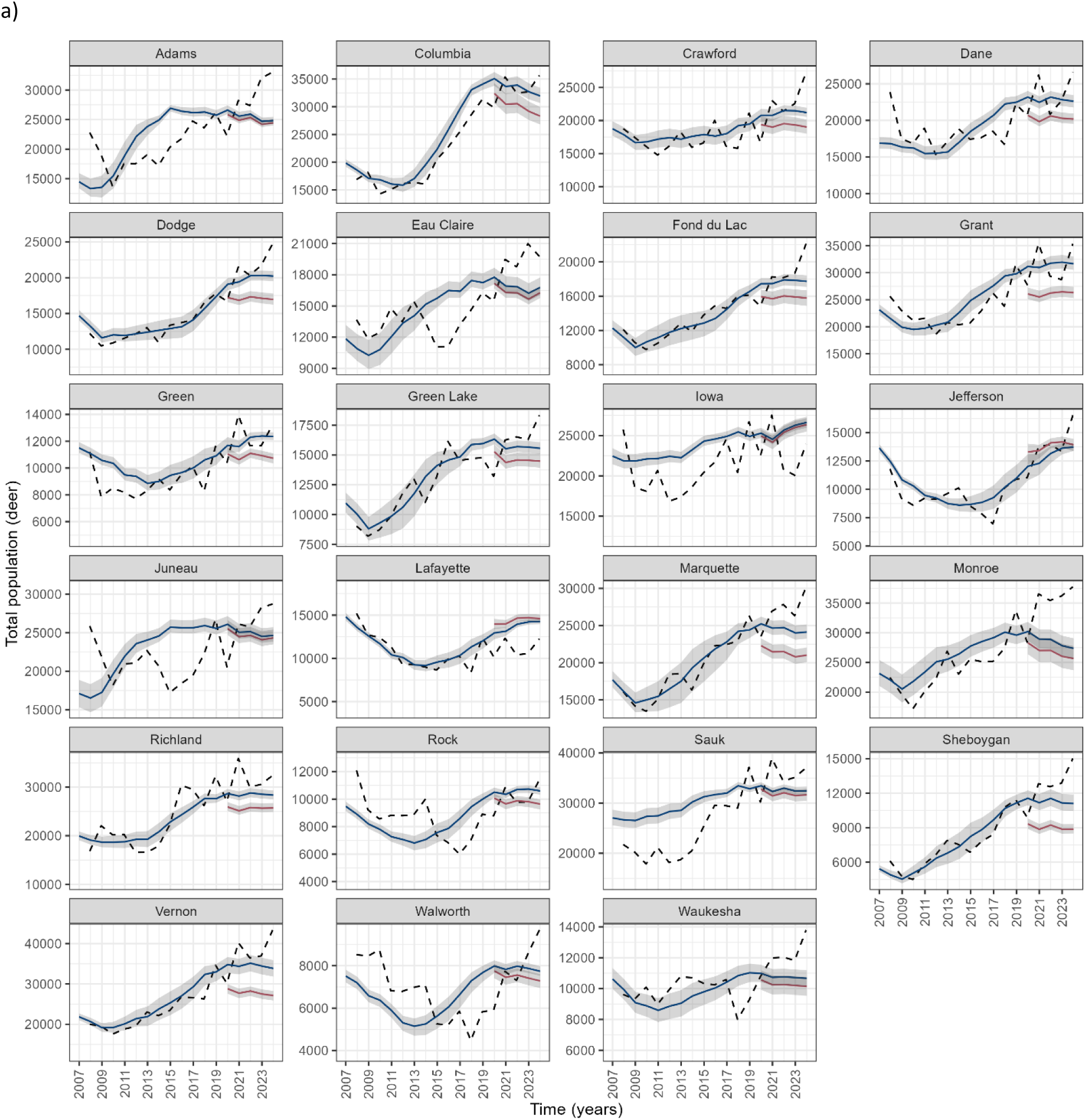
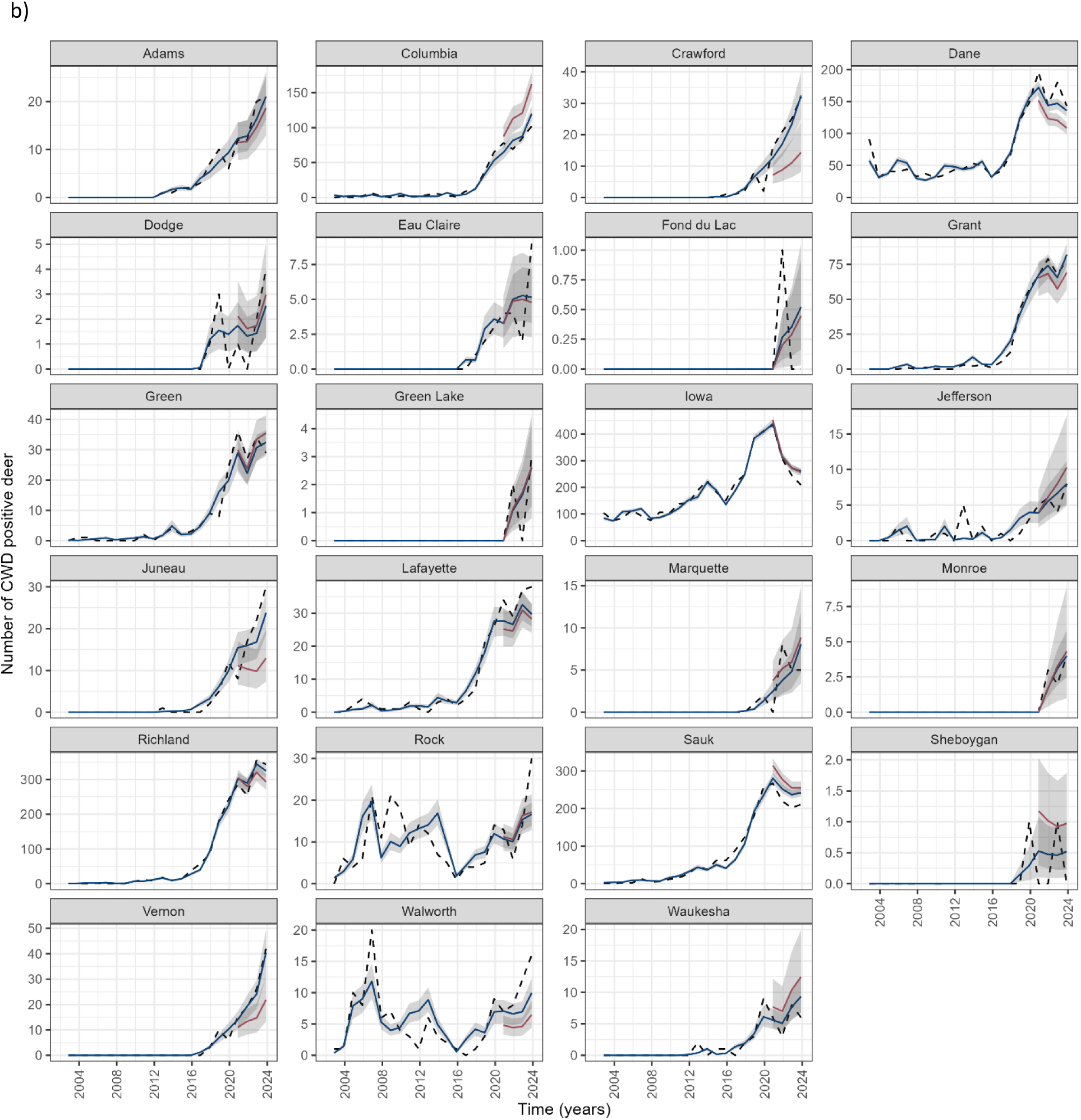
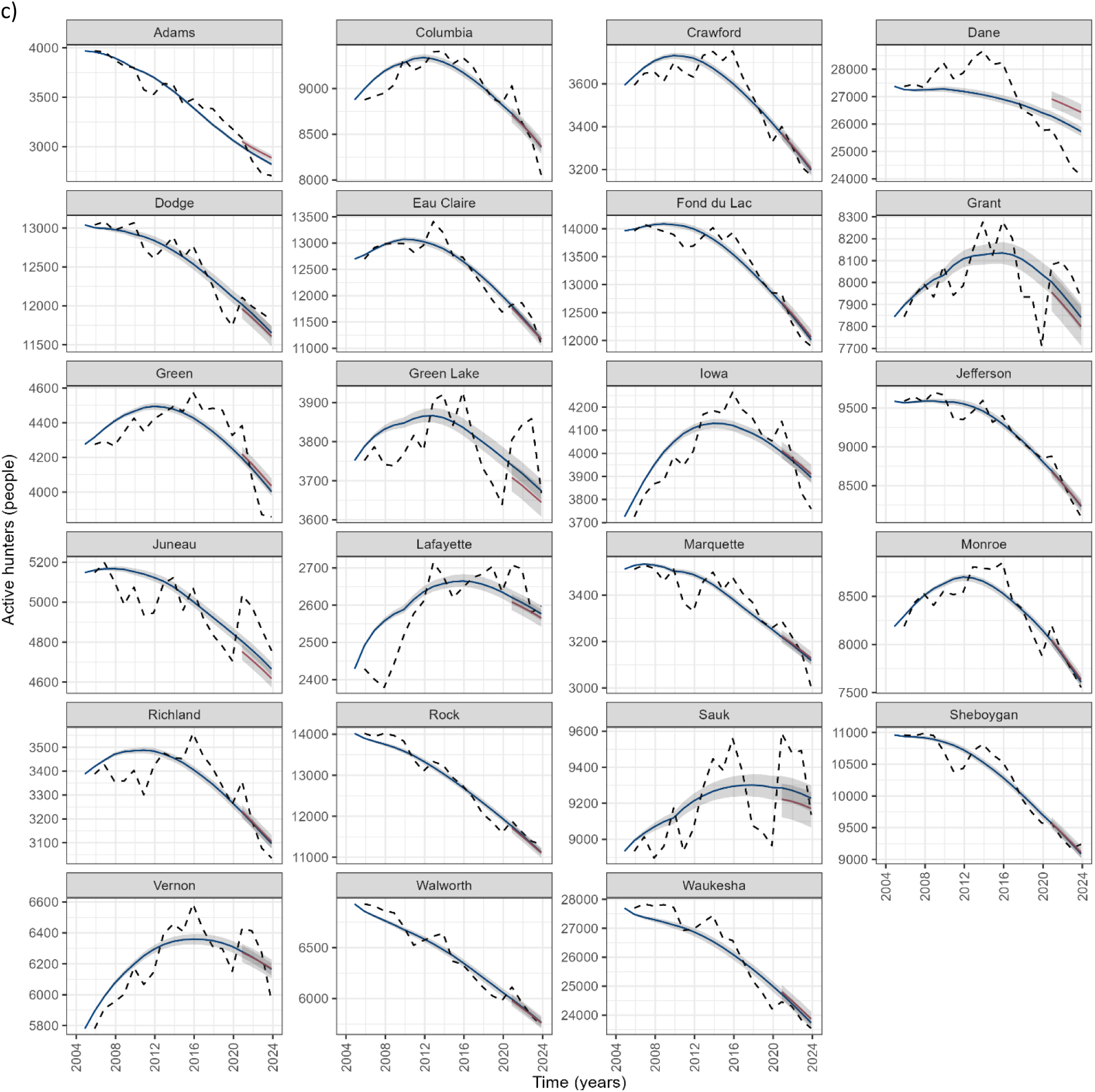
Predictions from the susceptible-infected-clinical (*SIC*) model of chronic wasting disease (CWD) in white-tailed deer and the hunter submodel, fitted to both the full dataset (blue) and a restricted out-of-sample dataset (red), compared to historical time series data (dashed lines) by county provided by the Wisconsin Department of Natural Resources. Model predictions, including the mean (solid line) and 95% credible intervals (shaded grey), were generated over 1000 parameter combination draws from the joint posterior, thus incorporating parameter uncertainty and measurement errors in the data. The panels display: (a) post-hunt total deer population, (b) number of CWD-positive deer sampled, and (c) number of active hunters for the 23 states included in the *SIC* model. Predictions for the testing data (2020-2024) are presented for the out-of-sample fit which was trained on data prior to 2020.

In general, all *SIC* model predictions of the datasets showed sensitivity to local changes in parameters that modify initial conditions, recruitment rates, and infection rates (Appendix *Section 4.1.3.2*). The parameters that allow for the initial population (*init pop adj[Region]*; refer to Appendix *Section 3.2.1.3* for details and Figure S5 and Table S17 for estimates) and population pressure (*Relative CC[Region]*; refer to Appendix *Section 2.2.1.1* for details and Figure S2 and Table S17 for estimates) to be adjusted away from values estimated directly from abundance and deer range data had a strong influence on all predictions. The recruitment rate is a result of birth rates and fawn mortality, and all predictions were strongly influenced by the reference birth rate (Table 1) and, to an extent, the effective harvest rate age weight of fawns (Table 1) –which impacts the number of fawns that are harvested and thus their mortality—in addition to the reference background and early fawn mortality rates themselves (Table 1) which directly impact recruitment and older deer mortality. Given that CWD is invariably fatal, the infection parameters that strongly influenced the number of expected positives (Appendix *Section 4.1.3.2*), namely the transmission coefficient for direct transmission and the duration of the exposed and infectious stages (Table 1), also influenced model predictions of other data sources such as the fawn-doe and yearling harvest ratios. In the hunter submodel, predictions were generally sensitive to the retention rate and wildlife value orientation (Table 1). For more detailed information on the magnitude and ranking of parameter sensitivities for each data source used to fit the *SIC* and hunter submodel, refer to Appendix *Section 4.1.3.2*. and *4.2.3.1*, respectively.

**Table 1.**
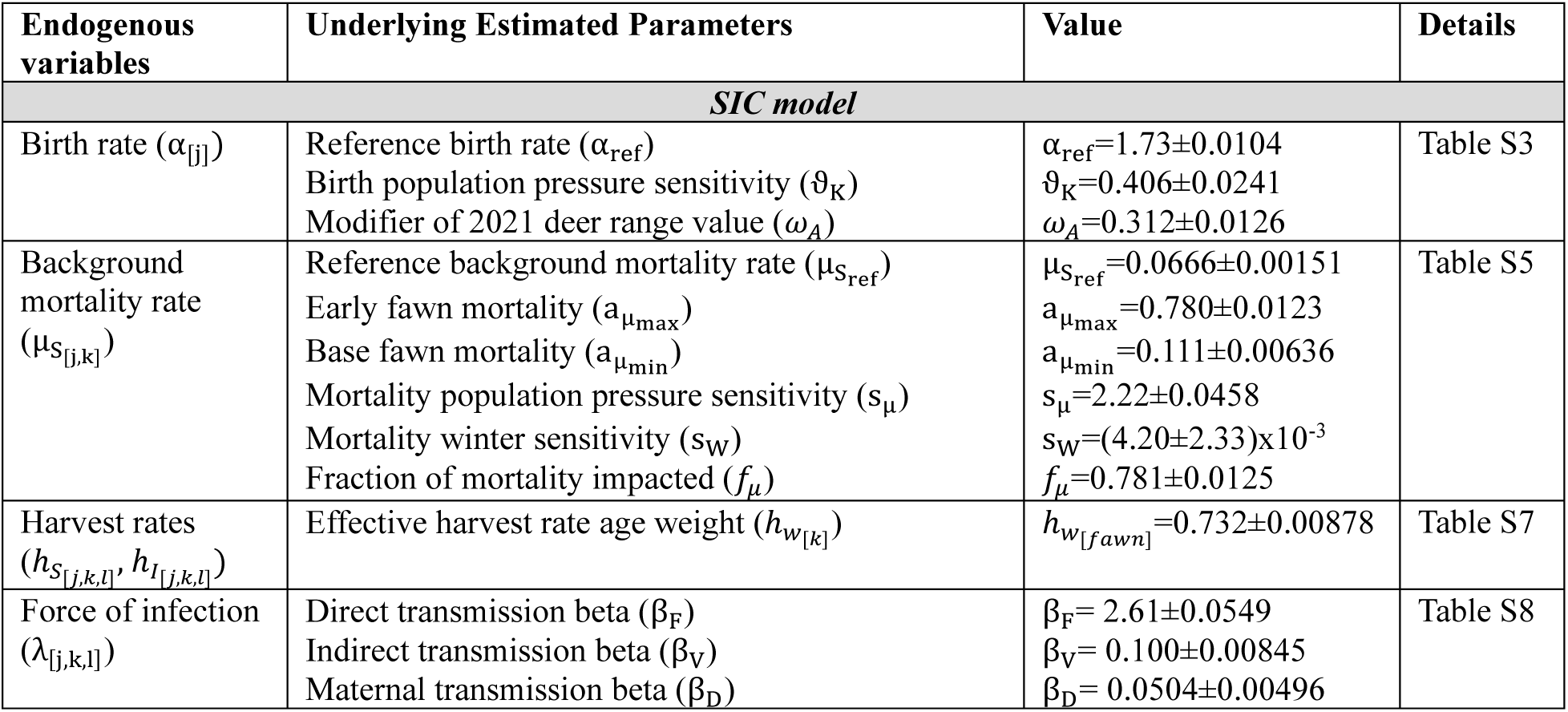
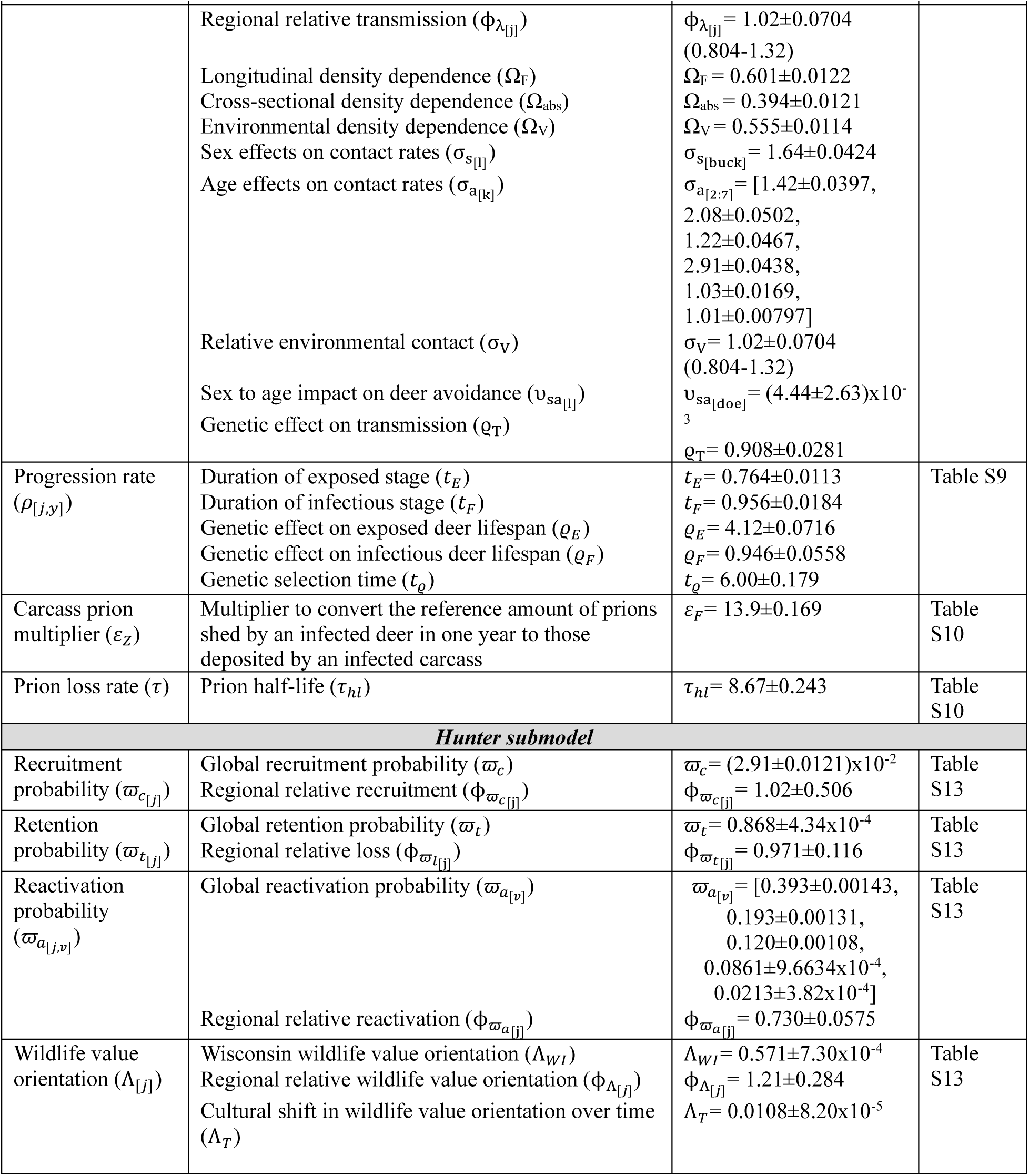
Values for epidemiological and demographic Susceptible-Infected-Clinical (*SIC*) model parameters and parameters driving active hunters in the hunter submodel estimated during the model fitting process. The table presents endogenous variables, as described in the model methods, alongside the underlying parameters from which they are derived. Parameter estimates are reported as mean values and standard deviations from the joint posterior distribution. For hierarchical parameters (i.e., county-level modifiers), we report the mean and standard deviation across all counties, as well as the minimum and maximum values. A complete accounting of county-specific estimates is provided in Appendix Figures S1-6. Appendix tables corresponding to each endogenous variable that detail the functional relationships between these variables and their underlying parameters are referenced.

When parameters were fit to a restricted dataset (i.e., the last four years of data withheld), the resulting estimates remained generally normally distributed, but the mean shifted for several parameters related to infection and background deer survival. The age effect on contacts increased for all ages but for four-year-olds, which decreased along with the sex effects on buck contacts. The estimated prion half-life and carcass prion multiplier decreased while the environmental contact increased. The reference background mortality parameter increased, as did the bias against fawn harvest. When compared to the testing data, the model predicted deer populations (2020-2024) were generally lower than the data (Figure 5a). Credible intervals of out-of-sample predictions for CWD-positive deer in surveillance generally overlapped with the data and tracked the trend (Figure 5b). Similarly, predictions for fawn-to-doe (Figure S2a) and yearling buck ratios (Figure S2c) were generally higher than those observed in the data, with overlap between predictions at certain timepoints. The credible intervals for the predicted frequency of the *S* allele in 2020 overlapped with the data (Figure S2b) as did the predictions for the yearling doe share of the harvest in most counties (Figure S2d). The *SIC* model’s qualitative measures of fit remained strong, Theil statistics showed a large portion of the error in predicted metrics were attributed to random error (Figure S12) and the associated MAPE values suggest strong predictive capabilities from the model fit (Figure S13). In the hunter submodel, neither parameter estimates, nor model predictions were influenced by fitting to a restricted dataset (Figure 5c, S21-23).

The resulting *SIC* model joint posterior estimates indicate that the transmission coefficients (i.e., betas –product of the contact rate and probability of infection given contact) for indirect and maternal transmission are non-zero (Table 1). Density dependence parameters for both direct and indirect transmission are of a similar magnitude (Table 1) and take intermediate values between zero (indicates no density effect) and one (indicates strong density effects). Parameters describing the *S* allele effect at codon 96 on the transmission and infectious stage progression of CWD are near one indicating minimal evidence for impacts on the transmission of CWD or the progression of the disease during the infectious period. However, the *S* allele effect on the exposed stage progression is estimated to be approximately four (Table 1) suggesting this allele significantly impacts disease progression. Lastly, county-specific annual per-hunter, antlered and antlerless harvest rates (Figures S27-32; Table S19) provided evidence for a slightly positive temporal trend (Time coefficient = 0.0027, 95% CI: 0.0014, 0.0041) and a negative trend across time (Time coefficient = –0.0093, 95% CI: –0.0116, –0.0071) for the per-capita hunter harvest of antlered and antlerless deer, respectively.

## Discussion

### Model Performance and Validation

The model structure developed in this study effectively captures the expected dynamic relationships among demographic, epidemiological, ecosystem, and human social components of the system. It preserves key causal relationships between variables, and the emergent metrics for host, ecosystem, disease, and social dynamics—calculated endogenously from the calibrated model—closely align with reported values and trends in the literature. For example, modeled birth rates for fawns, yearlings, and older does fall within the ranges reported for WI (McCaffery et al., 1998), and the inverse relationship between birth rates and deer density (Keyser et al., 2005; McCaffery et al., 1998; Swihart et al., 1998) is preserved (Figure S9). Age-specific background mortality rates align with published survival probabilities (Figure S10) in the absence of harvest (Rohm et al., 2007; Verme et al., 1968; Vreeland et al., 2004; Wasserberg et al., 2009).

Within the CWD management zone, the model estimates apparent prevalence to be approximately 24% lower than true prevalence, consistent with findings by Viljugrien et al. (2019) after adjusting for test sensitivity (Haley et al., 2012). In Wisconsin’s endemic regions, disease saturation at the county level scale becomes evident at 25% prevalence in the surveillance data, with a projected peak of 36% by 2050—closely matching historical trends reported by the WDNR (Wisconsin Department of Natural Resources, 2023). Additionally, the proportion of infections attributed to indirect transmission increases gradually over time, reflecting findings by Almberg et al. (2011) and Thompson et al. (2024).

The dynamics of the human dimensions of the model are also consistent with prior research. The model estimates that traditionalist or pluralist wildlife value orientations—common among potential hunters—are more prevalent in rural than in urban counties (Figure S20), in line with literature linking urbanization to increasing mutualist values (Manfredo, Berl, et al., 2021; Manfredo, Teel, et al., 2021). Furthermore, after accounting for hunter movement between counties, the model replicates the reported decline in license purchasing (Mohr et al., 2025; Figure 5c; Riley et al., 2016; Winkler & Warnke, 2013). Collectively, these results demonstrate that the model structure reliably captures system behaviors despite the large number of parameters to be estimated.

System Dynamics models, and ODE models more broadly, are mechanistic a priori structures that combine diverse information sources to constrain admissible solutions. Model results must conform not only to formal data and priors, but also to dimensional consistency, conservation laws, and face validity of processes described to stakeholders in nonmathematical terms.

Additionally, the mechanistic representation is tested for robustness against extreme conditions outside the historical range of data. The model structure that was derived by incorporating prior knowledge and known biological constraints within our system permits more reliable estimation of model parameters with biologically interpretable outcomes, even with sparse data.

One can think of this as a process of regularization—which seeks to prevent overfitting— combining priors that promote consistency with the underlying system of equations to penalize variety in parameters that does not align with the structure of the model dictated by the various physical processes operating within the system. For the host-disease component, data (6,509 points, mostly surveillance) is fairly abundant relative to the parameter set (173 in total, substantially nuisance parameters like population initial conditions), so key features are well-constrained despite limitations like a lack of direct data for population. To the extent that parameter combinations are poorly constrained, we at least expect this to be revealed in the joint posterior distribution.

Some posterior distributions for parameters—particularly those related to genetic and environmental prion dynamics— are difficult to precisely estimate due to limited data and considerable uncertainty about underlying mechanisms. For example, the genetic shift in the *S* allele frequency can result from multiple plausible combinations of genetic effects on disease transmission and progression. However, regularization through model structure and the integration of multiple datasets results in the joint posterior distribution being bounded by biologically feasible parameter ranges (Hossain et al., 2024; Karniadakis et al., 2021; Kosmatopoulos et al., 1995), demonstrating a range of plausible values for even those parameters with identifiability issues and providing novel insights into the underlying epidemiological processes (Table 1).

This approach parallels advances in physics-informed neural networks (Cuomo et al., 2022; Kharazmi et al., 2021; Raissi et al., 2019; Secci et al., 2024; Zhu et al., 2021), and integrated population models (Riecke et al., 2019), both of which have gained traction in modeling complex biological systems (Christian et al., 2024; Gamelon et al., 2021; Kashinath et al., 2021; Kissas et al., 2020; Weldy et al., 2023; Zipkin et al., 2023). Developing our model using regularization techniques ensured it reflected the most current scientific understanding of the host-pathogen system and broader hunter dynamics, allowing for estimation of previously unmeasured parameters and hypothesis testing around indirect transmission, density dependence, and genetic shifts.

### Key Insights and Limitations

Model results suggest indirect transmission (Table 1) must occur within the system at some level –given realistic values of direct transmission and observed prevalence – which aligns with the growing body of literature suggesting the relevance of an indirect transmission loop within the CWD system (Almberg et al., 2011; Plummer et al., 2018; Thompson et al., 2024). Despite the comparatively smaller magnitude of the indirect beta (Table 1), the persistence of prions on the landscape leads to a buildup in the later phases of an epidemic, eventually resulting in a greater proportion of new infections stemming from the indirect loop. These dynamics delay the contribution of the indirect loop to transmission, suggesting that early intervention is conducive to effective management. Transmission was determined to have an intermediate dependence on the density within a population (Table 1) rather than being solely frequency or density dependent (Habib et al., 2011; Joly et al., 2006; Kjær & Schauber, 2022; Storm et al., 2013) – further supporting the utility of density reduction as a CWD management tool.

Results suggest that the genetic shift towards the *S* allele causes a delayed progression of the disease, rather than impacting transmission (Table 1). These results align with experimental infections studies that span longer time periods (Denkers et al., 2024; C. Johnson et al., 2006), indicating that individuals with *S* alleles are still susceptible to CWD, but undergo longer incubation periods (i.e., exposed stage). Although a longer incubation period could result in a higher lifetime reproductive value for infected individuals, it is key to consider that these genetic differences do not appear to reduce transmission and, thus, are unlikely to prevent eventual CWD spread. Furthermore, our understanding of the difference in lifetime shedding rates between these genotypes is lacking, leading to a dearth of data regarding comparative contributions to environmental prions and subsequent indirect transmission (Denkers et al., 2024).

Our mechanistic model of active hunters allows managers to estimate the per-hunter harvest required for a given management scenario, providing an important social feasibility check when evaluating potential management scenarios. By incorporating hunters and deer within the same model we were able to estimate historic county-specific per-hunter harvest rates, revealing increasing antlered (Figures S27-28; Table S19) and decreasing antlerless (Figures S30-31; Table S19) deer per-hunter harvest rates over time. This change in harvest behavior is indicative of the importance not only of license sales, but of hunter behavior after purchase, suggesting that R3 programs must look beyond the metric of license sales to disentangle the expected impact of hunters on deer populations.

To address uncertainty in deer population estimates, the model uses a reference value to evaluate changes in longitudinal density rather than relying on absolute population size— countering the variability and potential bias in estimates used to assess density dependence in typical regressions. Because our model incorporates mechanistic demographic processes and observed harvest data, it can provide a check on data streams that are themselves a product of statistical modeling efforts. For example, deer population data estimated from the SAK and accounting models can be biased by harvest changes, leading to misleading trends that inform estimates of both deer populations and the harvest rates of those populations (Millspaugh et al., 2006; Rolley, 2013; R. E. Rolley, retired WDNR biologist, written comm., February 9, 2012). Our model predictions diverged from the data at the beginning of the time series when earn-a-buck legislation was present — which modified antlered harvest— suggesting a higher deer population. Since our model is constrained by mechanistic representations of deer demographics and actual harvest numbers, these estimates suggest that the rapid population growth depicted in the data is unlikely. Given that counties have varied surveillance histories as well as different antlerless harvest and population abundance trajectories, we expect the divergence of data from model predictions to also vary by county based on data quality and availability. Similarly, the model predicted values for the reactivation of hunters from the final inactive stage are consistently higher than suggested by the data, which is not unexpected given that this stock should include recruitment events prior to the beginning of our dataset and excludes unobserved hunter mortality events—both of which are unobservable and only estimated using available proxies.

Given limited resources, insights from our sensitivity analyses can help to (a) prioritize data collection efforts by identifying variables with the greatest influence on disease dynamics to inform future decision making (i.e., “value of information”; Runge et al., 2011), and (b) identify leverage points that can yield the greatest impact through management intervention (Restif et al., 2012). While transmission parameters remain important, our model highlights the dominant role of the initial deer population and recruitment rates in shaping CWD outbreak trajectories (Thompson et al., 2024). These results suggest that increasing accuracy of the population metrics (i.e., abundance, sex ratio, age structure) and demographic rates will significantly improve understanding of disease dynamics. Alternatively, experimental studies on CWD can allow us to inform individual parameters such as the prion half-life or maternal transmission coefficient, which can improve accuracy and precision of these parameter estimates.

Given the importance of hunter harvest in controlling the spread of CWD, we consider both the number of hunters and the number of deer harvested per hunter. The hunter submodel suggests efforts to increase recruitment and reactivation have been insufficient to offset declines in license purchasing under the continuation of the ongoing cultural changes represented by a shift away from traditionalist wildlife values. Given the difficulty of countering this widespread change, retaining existing hunters has a greater continued impact on overall license purchasing than recruitment. The need to balance the mortality rates of an aging hunter pool underscores the need to sustain (and marginally increase) some level of recruitment, even under perfect retention scenarios. Importantly, our model shows that modifying the behavior of current hunters to increase harvest per hunter may offer a practical lever for achieving harvest goals. For example, in Iowa County WI, over the last three years hunters harvested an average of 0.25 antlerless deer per hunter per year (Figure S32). Therefore, if just 50% of existing Iowa County hunters committed to harvesting one antlerless deer each year it would double present antlerless harvest rates—leading to reduced deer density and, subsequently, lower rates of CWD transmission. Although increasing harvest is not a novel concept for controlling CWD, the SD modelling approach makes it possible to integrate ecological and human dimensions within a single model, resulting in reliable estimates of present conditions and allowing for defined management targets and the specific level of intervention needed to reach them.

### Conclusions and Future Directions

Wildlife managers are increasingly confronted with threats and management challenges from emerging wildlife diseases (Henke et al., 2007), yet response frameworks akin to those used in human and agricultural health systems remain largely absent (Langwig et al., 2015; Wilkinson et al., 2024). A holistic approach that can encompass the biological uncertainty, conflicting stakeholder objectives, and imbalanced power dynamics (Decker et al., 1996; Dressel et al., 2018; Henke et al., 2007; McEachran et al., 2024) can help avoid paralysis caused by the substantial complexity of natural systems. The SD model presented here offers an integrated view of the interactions among deer, CWD, vegetation, and hunters—factors that dynamically influence real management outcomes. The model has been used to explore the impact of individual actions or simultaneous implementation of actions as part of portfolios representing different response strategies (Price Tack et al., n.d.). The SD model can be used to explore counterfactual histories (i.e., what would have happened if alternative management had been employed) and to explore uncertainties in system mechanisms through hypotheses testing (e.g., transmission pathways; see Penk et al., 2021 for example).

The model has been intentionally designed for iterative refinement as new data and insights emerge. For example, the human dimensions components could be expanded to include additional policy or economic feedbacks to enhance the ability to explore cost and benefits of different management strategies beyond the current capabilities based solely on tag fees, test costs, and agency removal costs. In addition to supporting rigorous exploration of quantitative uncertainty in model parameters and structure (Amorocho-Daza et al., 2025; Elsawah et al., 2020; Moallemi et al., 2020; Naugle et al., 2024; Schwaninger & Groesser, 2009; Yue et al., 2018), the SD modeling process also addresses qualitative uncertainties, such as competing objectives or limited transparency within the social and policy domains (Amorocho-Daza et al., 2025; Elsawah et al., 2015; Király & Miskolczi, 2019). When combined with a participatory modeling approach, SD facilitates meaningful stakeholder engagement and communication, enhancing shared understanding and trust in the process and model outcomes (Amorocho-Daza et al., 2025; Elsawah et al., 2017; Maier et al., 2016). These attributes make SD models particularly well-suited for informing wildlife disease management that requires coordination between science and the social processes that shape its application.

Ultimately, the complexity of wildlife disease systems like CWD—where host population, epidemiological, ecological, and human dimensions interact dynamically over time—benefit from a systems thinking approach and SD modeling. The approach is exemplified by the principles of *One Health*, where human, wildlife, and environmental well-being are managed as an integrated whole in acknowledgement of the interconnectedness between each element (Cunningham et al., 2017). SD models like the one presented here can provide an essential tool for managers trying to balance these factors when making management decisions in the face of nonlinear and varied dynamics across the system facets and over time. Moreover, SD models can be integrated with other decision-making approaches (e.g., structured decision making; Runge, 2020) or Resist-Accept-Direct frameworks (Lynch et al., 2021)—which assist managers deciding whether to resist, accept or direct ecological change—and determine optimal strategies. Where applied, SD models reveal leverage points, forecast unintended consequences, and allow managers to test strategies via computer simulations before implementation. By striving to make sense of uncertainty, the SD modelling approach empowers agencies to move beyond reactive decision-making. Instead of inaction in the face of complex, nonlinear systems, managers can proactively guide interventions that are robust, adaptive, and ultimately beneficial for both humans and ecosystems.

## Acknowledgements

We would like to thank the stakeholders and system experts who participated in our model building meetings as well as Emily Almberg and Melia Devivo for their thoughtful review of our manuscript and model. Any use of trade, firm, or product names is for descriptive purposed only and does not imply endorsement by the U.S. Government.

## Data Availability Statement

No data were collected for this study.

