## Appendix S1 for "A system dynamics model to understand the integrated ecological and human dimension aspects of wildlife health and disease management"

This information product has been peer reviewed and approved for publication as a preprint by the U.S. Geological Survey.

### A holistic approach to assessing and addressing wildlife health issues using system dynamics modeling.

Stephanie R. Penk[1*]; https://orcid.org/0000-0002-8027-4372

Christine Anhalt-Depies[2]; https://orcid.org/0000-0002-1295-3154

Richard E.W. Berl[3,4†]; https://orcid.org/0000-0002-4154-1319

Thomas S. Fiddaman[5]; https://orcid.org/0000-0003-1112-3199

Kawika Pierson[6]; https://orcid.org/0000-0002-1662-8746

Jennifer L. Price Tack[2]; https://orcid.org/0000-0002-5378-8183

Bryan J. Richards[3]; https://orcid.org/0000-0001-9955-2523

Erica Rieder[1]; https://orcid.org/0000-0001-6499-3580

Daniel J. Storm[2]

C. LeAnn White[3]; https://orcid.org/0000-0002-5004-5165

Daniel P. Walsh[7]; https://orcid.org/0000-0002-7772-2445

[1]Montana Cooperative Wildlife Research Unit, Wildlife Biology Program, University of Montana, Missoula, MT 59812, U.S.A

[2]Wisconsin Department of Natural Resources, 107 Sutliff Avenue, Rhinelander, WI 54501, USA

[3]U.S. Geological Survey, National Wildlife Health Center, 6006 Schroeder Road, Madison, WI 53711, U.S.A

[4]U.S. Geological Survey, Eastern Ecological Science Center at the Patuxent Research Refuge, 12100 Beech Forest Road, Laurel, MD 20708, U.S.A

[5] Ventana Systems Inc., 8105 SE Nelson Road, Olalla, Washington 98539, U.S.A.

[6] Willamette University, Atkinson Graduate School of Management, U.S.A

[7]U.S. Geological Survey, Montana Cooperative Wildlife Research Unit, Wildlife Biology Program, University of Montana, Missoula, MT 59812, U.S.A

[†]Current address: 12100 Beech Forest Road, Laurel, MD 20708, USA

#### Appendix S1

Any use of trade, firm, or product names is for descriptive purposes only and does not imply endorsement by the U.S. Government.

### 1 Literature search

**Table S1. Statistical and mathematical models of chronic wasting disease (CWD) are categorized by the system facets they incorporate—host dynamics, social dynamics, and ecosystem dynamics—and by how these facets are represented in the model: endogenously, exogenously, or mechanistically. All included models explicitly account for CWD epidemiology, such as prevalence, disease-associated mortality, or transmission.** *Host dynamics* include variables related to sex and age structure, healthy vital rates, dispersal, genetics, and other population-level biological traits. *Social dynamics* refer to human influences on the system (e.g., hunting, carcass transport, supplemental feeding) and economic components such as management costs or revenue. *Ecosystem dynamics* include higher-order ecological factors, such as predator effects, vegetation dynamics, and indirect effects on non-deer species; environmental prion accumulation is also considered an ecosystem dynamic. *Endogenous (“Endo”)* variables are determined by interactions within the model, akin to dependent variables in statistical models. *Exogenous (“Exo”)* variables are externally determined and serve as independent variables. *Mechanistic (“Mech”)* variables are based on mathematical representations of underlying processes, rather than empirical data patterns alone. Here we build on the Web of Science literature search in Winter and Escobar (2020), removing models that include only epidemiology. We extended their review to include relevant studies published between 2018 and the end of 2023 and classified each model based on the facets addressed and how they were integrated.

| Manuscript Citation | Epidemiology | | | Host Dynamics | | | Ecosystem Dynamics | | | Social Dynamics | | |
|---|---|---|---|---|---|---|---|---|---|---|---|---|
| | *Endo* | *Exo* | *Mech* | *Endo* | *Exo* | *Mech* | *Endo* | *Exo* | *Mech* | *Endo* | *Exo* | *Mech* |
| (Miller et al., 2000) | X | | X | X | | X | | | | | | |
| (Conner et al., 2000) | | X | | | X | | | | | | | |
| (Gross and Miller, 2001) | X | | X | X | | X | | | | | | |
| (Joly et al., 2003) | X | | | | X | | | | | | | |
| (Diefenbach et al., 2004) | | X | | | X | | | | | X | | |
| (Conner and Miller, 2004) | X | X | X | X | | | | | | | | |
| (Farnsworth et al., 2005) | X | | | | X | | | X | | | | |
| (Miller and Conner, 2005) | X | | | | X | | | | | | | |
| (Krumm et al., 2005) | X | | | | X | | | | | | | |
| (Grear et al., 2006) | X | | | | X | | | | | | | |
| (Johns and Mehl, 2006) | X | | X | | X | | X | | | | | |
| (Farnsworth et al., 2006) | X | | | | X | | | X | | | | |
| (Joly et al., 2006) | X | | | | X | | | X | | | | |
| (Miller et al., 2006) | X | X | X | X | X | X | X | X | | | | |
| (Conner et al., 2007) | | X | | | X | | | | | X | | |
| (Farnsworth et al., 2007) | X | | | | X | | | X | | | | |
| (Nusser et al., 2008) | | X | | | X | | | | | | | |
| (Miller et al., 2008) | | X | | X | | | | | | | | |
| (Blanchong et al., 2008) | X | | | X | X | | | X | | | | |
| (Joly et al., 2009) | | X | | | X | | | | | | | |
| (Wasserberg et al., 2009) | X | X | X | X | | X | | | | | | |
| (Osnas et al., 2009) | X | | | | X | | | | | | | |

| Manuscript Citation | Epidemiology | | | Host Dynamics | | | Ecosystem Dynamics | | | Social Dynamics | | |
|---|---|---|---|---|---|---|---|---|---|---|---|---|
| | *Endo* | *Exo* | *Mech* | *Endo* | *Exo* | *Mech* | *Endo* | *Exo* | *Mech* | *Endo* | *Exo* | *Mech* |
| (Song and Lawson, 2009) | X | X | X | X | X | | | | | | | |
| (Heisey et al., 2010) | X | | | | X | | | | | | | |
| (Grear et al., 2010) | X | | | | X | | | | | | | |
| (Dulberger et al., 2010) | | X | | X | | | | | | | | |
| (Lawson and Song, 2010) | X | | | | X | | | | | | | |
| (Walsh and Miller, 2010) | | X | | | X | | | | | | | |
| (Catherine I. Cullingham et al., 2011) | | X | | | X | | | | | | | |
| (C. I. Cullingham et al., 2011) | | X | | X | X | | | X | | | | |
| (Wild et al., 2011) | X | | X | X | | X | | X | | | | |
| (Almberg et al., 2011) | X | X | X | X | | X | | X | | | | |
| (Sharp and Pastor, 2011) | X | X | X | X | X | X | X | X | X | | | |
| (Robinson et al., 2012) | X | X | X | X | X | X | | | | | | |
| (Al-Arydah et al., 2012) | X | X | X | X | X | X | | | | | | |
| (Blanchong et al., 2012) | | X | | X | X | | | | | | | |
| (Rees et al., 2012) | | X | | | X | | | X | | | | |
| (Potapov et al., 2012) | X | X | X | X | X | X | | | | | | |
| (Storm et al., 2013) | X | X | X | | X | | | X | | | | |
| (Matsumoto et al., 2013) | | X | | | X | | | | | | | |
| (Cortez and Weitz, 2013) | X | X | X | X | X | X | X | X | X | | | |
| (Potapov et al., 2013) | X | X | X | X | X | X | X | X | X | | | |
| (Mateus-Pinilla et al., 2013) | X | | | | X | | | X | | | | |
| (O'Hara Ruiz et al., 2013) | X | | | X | | | | X | | | | |
| (Robinson et al., 2013) | X | | | | | | | X | | | | |
| (Manjerovic et al., 2014) | X | | | | | | | X | | | | |
| (Garlick et al., 2014) | X | X | X | X | X | X | | X | X | | | |
| (Oraby et al., 2014) | X | | | X | X | X | | X | | | | |
| (Monello et al., 2014) | X | X | | X | X | | | | | | | |
| (Jennelle et al., 2014) | X | X | X | X | X | X | | | | | | |
| (Heisey et al., 2014) | X | | | | X | | | | | | | |
| (Williams et al., 2014) | | X | | X | X | X | | | | | | |
| (Garlick et al., 2014) | X | X | X | X | X | X | X | X | X | | | |
| (Vasilyeva et al., 2015) | X | X | X | X | X | X | X | X | X | | | |
| (Potapov et al., 2015) | X | X | X | | X | | | | | | | |
| (Geremia et al., 2015) | X | X | X | X | X | X | | | | | | |

| Manuscript Citation | Epidemiology | | | Host Dynamics | | | Ecosystem Dynamics | | | Social Dynamics | | |
|---|---|---|---|---|---|---|---|---|---|---|---|---|
| | *Endo* | *Exo* | *Mech* | *Endo* | *Exo* | *Mech* | *Endo* | *Exo* | *Mech* | *Endo* | *Exo* | *Mech* |
| (Sun et al., 2015) | X | X | X | X | X | X | X | X | X | | | |
| (Al-arydah et al., 2016) | X | X | X | X | X | X | | | | | | |
| (Edmunds et al., 2016) | | X | | X | X | X | | | | | | |
| (Evans et al., 2016) | X | | | X | | | | X | | | | |
| (Mejía Salazar et al., 2016) | | X | | X | X | | | X | | | | |
| (Nobert et al., 2016) | X | | | | X | | | | | | | |
| (Samuel and Storm, 2016) | X | X | | | X | | | | | | | |
| (Mejía-Salazar et al., 2017) | | X | | X | X | | | X | | | | |
| (Hefley et al., 2017b) | X | | X | | X | | | X | | | | |
| (Galloway et al., 2017) | | X | | | X | X | | | | | | |
| (Monello et al., 2017) | X | | | | X | | | | | | | |
| (Hefley et al., 2017a) | X | | X | | X | | | X | | | | |
| (DeVivo et al., 2017) | | X | | X | X | X | | | | | | |
| (Edmunds et al., 2018) | | X | | | X | | X | | | | | |
| (Davenport et al., 2018) | X | X | | | X | | | | | | | |
| (Maji et al., 2018) | X | X | X | X | X | X | X | X | X | | | |
| (Wolfe et al., 2018) | X | | | | X | | | | | | | |
| (Jennelle et al., 2018) | X | X | | | X | | | | | | | |
| (Walker et al., 2020) | X | | | | X | | | X | | | | |
| (Belsare and Stewart, 2020) | X | X | X | X | X | X | | X | | | | |
| (Belsare et al., 2020) | X | X | X | X | X | X | | X | | | | |
| (Makau et al., 2020) | X | | | | X | X | | | | | | |
| (Maloney et al., 2020) | X | X | X | X | X | X | X | X | X | X | X | X |
| (Mysterud et al., 2020) | X | X | X | X | X | X | | | | | | |
| (Belsare et al., 2021) | X | X | X | X | X | X | | X | | | | |
| (Galloway et al., 2021) | X | X | X | X | X | X | | | | | | |
| (Lacava et al., 2021) | X | X | | X | X | | | | | | | |
| (Ott-Conn et al., 2021) | X | | | | X | | | | | | | |
| (Mysterud et al., 2021) | X | X | X | X | X | X | | X | | | | |
| (Sargeant et al., 2021) | X | | | | X | | | | | | | |
| (Smolko et al., 2021) | X | | | | X | | | X | | | | |
| (Winter et al., 2021) | X | | | | | | | X | | | | |
| (Brandell et al., 2022) | X | X | X | X | X | X | X | X | X | | | |
| (Cook et al., 2022) | X | X | X | | | | | X | | | | |

| Manuscript Citation | Epidemiology | | | Host Dynamics | | | Ecosystem Dynamics | | | Social Dynamics | | |
|---|---|---|---|---|---|---|---|---|---|---|---|---|
| | *Endo* | *Exo* | *Mech* | *Endo* | *Exo* | *Mech* | *Endo* | *Exo* | *Mech* | *Endo* | *Exo* | *Mech* |
| (Hanley et al., 2022) | X | X | X | X | X | X | X | X | X | | | |
| (Kjær and Schauber, 2022) | X | X | X | X | X | X | | X | | | | |
| (Ketz et al., 2022) | X | | | | X | | | | | | | |
| (Rogers et al., 2022) | X | X | X | X | X | X | | | | | | |
| (Seabury et al., 2022) | X | | | | X | | | | | | | |
| (Xu et al., 2022) | X | X | X | X | X | X | X | X | X | | | |
| (Tian et al., 2022) | X | | | | X | | | X | | | | |
| (Samuel, 2023) | X | | | | X | | | X | | | | |
| (Cook et al., 2023) | X | | X | | X | | | | | | | |
| (Mysterud et al., 2023) | X | X | | | X | | | | | | | |
| (Dobbin et al., 2023) | X | | | X | X | | | X | | | | |
| (Hearst et al., 2023) | X | | | | X | | | | | | | |
| (Strasburg and Christensen, 2024) | X | X | X | X | X | X | | X | | | | |
| (Dugovich et al., 2024) | | X | | | X | | | | | | | |
| (Thompson et al., 2024) | X | | X | X | X | X | | X | | | | |
| (Walter et al., 2024) | | X | | | X | | | X | | | | |
| (Ahmed et al., 2024) | X | | | | | | | X | | | X | |
| (Hoar et al., 2024) | | X | | X | | | | | | | | |
| (Booth et al., 2024) | | X | | X | | | | | | | | |
| (Davis et al., 2024) | X | | | | | | | X | | | | |
| (McClure and Powell, 2025a) | X | | X | X | X | X | X | X | X | | | |
| (McClure and Powell, 2025b) | X | | X | X | X | X | X | | X | | | |
| (Mysterud et al., 2025) | X | X | X | X | X | X | | | | | | |
| (Schuler et al., 2025) | | X | | | X | | | | | | X | |
| (Moss et al., 2025) | X | | | | X | | | | | | X | |
| (Mandujano Reyes et al., 2025) | X | | X | | | | | X | | | | |
| (Kanzinger and Abbott, 2025) | X | X | X | X | X | X | X | X | X | | | |

**Table S2. A summary table of the number of statistical and mathematical models of chronic wasting disease (CWD) that incorporate each combination of system facets—host dynamics, social dynamics, and ecosystem dynamics—alongside epidemiology. All included models explicitly account for CWD epidemiology (e.g., prevalence, disease-associated mortality, or transmission).** *Host dynamics* include variables related to sex and age structure, healthy vital rates, dispersal, genetics, and other population-level biological traits. *Social dynamics* refer to human influences on the system (e.g., hunting, carcass transport, supplemental feeding) and economic components such as management costs or revenue. *Ecosystem dynamics* include higher-order ecological factors, such as predator effects, vegetation dynamics, and indirect effects on non-deer species; environmental prion accumulation is also considered an ecosystem dynamic. Here we build on the Web of Science literature search in Winter and Escobar (2020), removing models that include only epidemiology. We extended their review to include relevant studies published between 2018 and September 2025 and classified each model based on the facets addressed and how they were integrated.

| | Epidemiology and Host Dynamics | Epidemiology and Ecosystem Dynamics | Epidemiology and Social Dynamics | Epidemiology, Host, and Ecosystem Dynamics | Epidemiology, Host, and Social Dynamics | Epidemiology, Ecosystem, and Social Dynamics | Epidemiology, Host, Ecosystem, and Social Dynamics |
|---|---|---|---|---|---|---|---|
| **Manuscript Count** | 56 | 5 | 0 | 50 | 4 | 1 | 1 |

### 2 Model description

#### 2.1 Purpose

This model was developed to simulate the interactions among host-pathogen dynamics, social behavior, and ecosystem processes within a unified, holistic framework. In its current application, the model focuses on white-tailed deer (*Odocoileus virginianus*) populations in Wisconsin affected by chronic wasting disease (CWD). The overarching goal is to evaluate and inform management strategies that not only mitigate disease spread but also remain viable under real-world social constraints, including stakeholder preferences and political considerations.

#### 2.2 Model structure and key formulations

At its core, the model uses a susceptible-infected compartmental framework to track healthy and diseased individuals over time. The deer population is stratified by age and sex, with infections arising through both direct (animal-to-animal) and indirect (environmental prion) transmission pathways. The model is disaggregated by region which is equivalent here to county (corresponding with deer management jurisdictions) but does not include spatial dynamics among counties.

The model is implemented in Vensim® (Ventana Systems Inc., 2025) as a system of differential equations that govern the flow of individuals between compartments—referred to as stocks—including susceptible, infected (subdivided into exposed and infectious stages), and clinical individuals (i.e., those in the final phase of disease prior to death). Additionally, the model includes a stock for environmental prion accumulation within each region (refer to Eq. S1). We refer to this as the susceptible-infected-clinical (*SIC*) model of chronic wasting disease (CWD) in white-tailed deer.

While disease dynamics and background deer mortality are continuous, a few processes are (optionally, active by default) represented as discrete events. Harvest occurs during a single time step in the fall, and births occur at a single time at the end of spring. We have not observed substantial changes in the dynamics when estimating continuous vs. discrete versions of the system, but the discrete events simplify several model-data comparisons, because the data are collected at various times during the year, over which there can be large variations. Normally deer are a continuous quantity, but the model can also be run with integer deer and stochastic transitions. This is primarily of interest for exploring low prevalence rates, where phenomena like stochastic extinction of disease are possible. This is less relevant for estimation, because we focus on counties with significant reporting of infected deer. Inclusion of stochastic effects would broaden credible intervals but is computationally prohibitive.

**Equation S1**

**S1.1 Change in number of susceptible deer**

$$\frac{dS_{[j,k,l]}}{dt} = \overbrace{\alpha_{[j]}D_{[j]}}^{\textit{Births}} - \overbrace{\lambda_{[j,k,l]}S_{[j,k,l]}}^{\textit{Disease transmission}} - \overbrace{\mu_{S_{[j,k]}}S_{[j,k,l]}}^{\textit{Background mortality}} - \overbrace{h_{S_{[j,k,l]}}S_{[j,k,l]}}^{\textit{Hunter harvest}} - \overbrace{m_{S_{[j,k,l]}}S_{[j,k,l]}}^{\textit{Agency removals}} + \overbrace{\sum_{j^*=1}^{n}\delta_{[j^*,j,k,l]}S_{[j^*k,l]}}^{\textit{Dispersal}} - \sum_{j^*=1}^{n}\delta_{[j,j^*,k,l]}\,S_{[j,k,l]}$$

**S1.2a Change in number of infected deer in first infection stage**

$$\frac{dI_{[j,k,l,y]}}{dt} = \lambda_{[j,k,l]}S_{[j,k,l]} - \overbrace{\rho_{[j,y]}I_{[j,k,l,y]}}^{\textit{Disease progression}} - \overbrace{(\mu_{S_{[j,k]}} + \psi_I)I_{[j,k,l,y]}}^{\textit{Early-stage disease increased mortality}} - h_{I_{[j,k,l]}}I_{[j,k,l,y]} - m_{I_{[j,k,l]}}I_{[j,k,l,y]} + \sum_{j^*=1}^{n}\delta_{[j^*,j,k,l]}I_{[j^*,k,l,y]} - \sum_{j^*=1}^{n}\delta_{[j,j^*,k,l]}I_{[j,k,l,y]} \qquad y = 1$$

**S1.2b Change in number of infected deer in all infection stages beyond the first**

$$\frac{dI_{[j,k,l,y]}}{dt} = \overbrace{\rho_{[j,y-1]}I_{[j,k,l,y-1]} - \rho_{[j,y]}I_{[j,k,l,y]}}^{\textit{Disease progression}} - \overbrace{(\mu_{S_{[j,k]}} + \psi_I)I_{[j,k,l,y]}}^{\textit{Early-stage disease increased mortality}} - h_{I_{[j,k,l]}}I_{[j,k,l,y]} - m_{I_{[j,k,l]}}I_{[j,k,l,y]} + \sum_{j^*=1}^{n}\delta_{[j^*,j,k,l]}I_{[j^*,k,l,y]} - \sum_{j^*=1}^{n}\delta_{[j,j^*,k,l]}I_{[j,k,l,y]} \qquad y > 1$$

**S1.3 Change in number of clinical deer**

$$\frac{dC_{[j,k,l]}}{dt} = \rho_{[j,y]}I_{[j,k,l,y]} - \overbrace{(\mu_C + \psi_C)C_{[j,k,l]}}^{\textit{Late-stage disease induced mortality}} - h_{C_{[j,k,l]}}C_{[j,k,l]} - m_{I_{[j,k,l]}}C_{[j,k,l]}$$

**S1.4 Change in amount of environmental prions**

$$\frac{dV_{[j]}}{dt} = \overbrace{(\underbrace{\varepsilon_F F_{[j]}}_{\textit{Infectious live deer}} + \underbrace{\varepsilon_F \varepsilon_z \frac{z_{F[j]}}{t_Z}(1 - \varsigma_H)}_{\textit{Infectious carcasses}})}^{\textit{Depositing}} - \overbrace{\tau V_{[j]}}^{\textit{Degrading}}$$

A separate population is tracked for each modeled region (*j*), with individuals in each of the three states—Susceptible (S), Infected (I), and Clinical (C)—further disaggregated by age class (*k*) and sex ($l$) to capture cohort-specific dynamics. Age classes include: (1) fawns (0–1 years), (2) yearlings (1–2 years), (3) two-year-olds, (4) three-year-olds, (5) four-year-olds, (6) five-year-olds, and (7) individuals six years and older. Sex is tracked as either male (*buck*) or female (*doe*). The infected stock is also subdivided into six infection stages ($y$), with the first three stages considered non-infectious and the final three shedding prions. Individuals progress through these stages over time following initial infection.

Because CWD is invariably fatal (Aguzzi et al., 2007; Collinge and Clarke, 2007; Williams et al., 2002), with death following shortly after the appearance of clinical symptoms (Almberg et al., 2011; Edmunds et al., 2018; Hamir et al., 2008; Jennelle et al., 2014; Wasserberg et al., 2009; Williams, 2005), the model does not include a recovered class. Infected individuals who progress to the final stage of infection are transitioned to a separate clinical stock to reflect the markedly different mortality risks and behavior exhibited by symptomatic deer (Edmunds et al., 2016; Williams et al., 2002). Disaggregation by age and sex increases model dimensionality substantially: within each region, 14 susceptible stocks, 14 clinical stocks, and 84 infected stocks are tracked to fully capture demographic and disease progression heterogeneity. Descriptions of model parameters in Equation S1 follow in subsequent sections.

**Table S3. Mapping between the full variable names used to construct the system dynamics model in Vensim (*Full Variable Name*) and the abbreviated forms (*Short Form*) used in the manuscript to represent infection states and birth processes.** The table includes each variable's units, the corresponding equation used to calculate it, or the source from which its value was derived.

| Short Form | Full Variable Name | Units | Equation/Source |
|---|---|---|---|
| $S_{[j,k,l]}$ | Healthy[Region, age, sex] | deer | Equation S1.1 |
| $I_{[j,k,l,y]}$ | Infected IE[Region, age, sex, stage] | deer | Equation S1.2 |
| $C_{[j,k,l]}$ | Clinical[Region, age, sex] | deer | Equation S1.3 |
| $\alpha_{[j]}$ | D Birth Rate[Region] | fraction/year | Equation S2.1 |
| $\alpha_{ref}$ | Ref Birth Rate | fraction/year | Estimated (Table S17) |
| $\alpha_{max}$ | Max Birth Rate Ratio | dimensionless | $\frac{2.2}{\alpha_{ref}}$ |
| $\alpha_{min}$ | Min Birth Rate Ratio | dimensionless | 0 |
| $\vartheta_K$ | Birth CC sensitivity | dimensionless | Estimated (Table S17) |
| $b_{C_{[j]}}$ | Buck coverage effect[Region] | dimensionless | $\frac{1}{\left(1+e^{4\left(1-\frac{b_{f_{[j]}}}{b_{min}}\right)}\right)}$ |
| $b_{f_{[j]}}$ | Buck fraction[Region] | fraction | $\frac{Q_{[j]}}{(Q_{[j]}+D_{[j]})}$ |
| $b_{min}$ | Min buck fraction | fraction | 0.1 (Milner et al., 2007) |
| $N_{[j]}$ | Population[Region] | deer | Equation S3.2 |
| $N_{\omega_{[j]}}$ | Weighted adult population[Region] | deer | Equation S3.1 |
| $K_{[j]}$ | Carrying Capacity[Region] | deer | Equation S4 |
| $D_{[j]}$ | Reproductive population[Region, does] | deer | Equation S2.2a |
| $Q_{[j]}$ | Reproductive population[Region, bucks] | deer | Equation S2.2b |
| $\omega_G$ | Fawn pop weight | dimensionless | 0.5 |
| $a_{\alpha_{[k]}}$ | age effect repro[age] | dimensionless | [0.3,0.66,1,1,1] (Green et al., 2017) |
| $K_{ref_{[j]}}$ | Ref CC[Region] | deer | Equation S5 |

| Short Form | Full Variable Name | Units | Equation/Source |
|---|---|---|---|
| $K_{\Delta_{[j]}}$ | Relative CC[Region] | dimensionless | Estimated (Table S17) |
| $N_{ref_{[j]}}$ | Ref population[Region] | deer | Data (Section 3.2.1.3) |
| $Y_{ref_{[j]}}$ | Ref density | deer/mile$^2$ | Equation S6 |

#### 2.2.1 Host dynamics

##### *2.2.1.1 Births and background mortality*

The model operates on a yearly cycle beginning January 1st, with each year divided into 16 time steps ($t_{step}$: 1–16), each representing roughly 3.25 weeks. Reproduction occurs as a single birth pulse during timestep 9, aligning with peak fawning season in Wisconsin (mid-May to July) (Jacques et al., 2007). Fawns are added to the first age class ($k$ = 1) based on the product of the density-dependent per-capita birth rate ($\alpha_{[j]}$) and the number of reproductive does ($D_{[j]}$) in each region (Eq. S1.1). Births are assumed to occur at a 1:1 sex ratio (Green et al., 2017; Verme, 1983), and all fawns are initially assigned to the susceptible stock; maternal transmission accounts for any increased infection risk to fawns given infectious doe levels.

**Equation S2**

$$\alpha_{[j]} = \begin{cases} \alpha_{ref}\left[\frac{1}{1+e^{4\left(1-\frac{b_{f_{[j]}}}{b_{min}}\right)}}\right]\left[\alpha_{max} + \frac{(\alpha_{min}-\alpha_{max})}{1+e^{\left(-4\vartheta_K\left(\frac{N_{\omega_{[j]}}}{K_{[j]}}-1\right)\right)}}\right] & t_{step} = 8 \\ 0 & t_{step} \neq 8 \end{cases}$$

**Equation S2.1. Regional per-capita birth rate**

$$D_{[j]} = \sum_{k=1}^{n}\sum_{l=2}^{2} S_{[j,k,l]}a_{\alpha_{[k]}} + \sum_{k=1}^{n}\sum_{l=2}^{2}\sum_{y=1}^{n} I_{[j,k,l,y]}a_{\alpha_{[k]}}$$

**Equation S2.2a. Regional total of reproductive does**

$$Q_{[j]} = \sum_{k=1}^{n}\sum_{l=1}^{1} S_{[j,k,l]}a_{\alpha_{[k]}} + \sum_{k=1}^{n}\sum_{l=1}^{1}\sum_{y=1}^{n} I_{[j,k,l,y]}a_{\alpha_{[k]}}$$

**Equation S2.2a. Regional total of reproductive bucks**

The per-capita birth rate ($\alpha_{[j]}$; Eq. S2.1) is derived by modifying a global reference birth rate ($\alpha_{ref}$; Table S3) using two region-specific multipliers: one capturing buck coverage effects (Milner et al., 2007) and another capturing density-dependent suppression of reproduction (Keyser et al., 2005; Swihart et al., 1998). Extremely low buck to doe ratios must occur to impact fertilization in ungulates (Milner et al., 2007), thus, buck coverage ([second term]; Eq. S2.1) is modeled as a logistic function based on the current fraction of bucks in the reproductive population ($b_{f_{[j]}}$) and a threshold buck fraction ($b_{min}$). The structure creates a steep increase towards 100% and very little reproduction below $b_{min}$ (Table S3). In practice buck fractions are normally adequate and the coverage effect is inactive, but it is provided for robustness in extreme conditions. Density dependence ([third term]; Equation. S2.1) is modeled using logistic scaling between user-defined upper ($\alpha_{max}$) and lower ($\alpha_{min}$) bounds on per-capita reproduction, with scaling based on the difference between the weighted population ($N_{\omega_{[j]}}$) and population pressure level ($K_{[j]}$; Equation. S4) and birth rate sensitivity to it ($\vartheta_K$; Table S3). While only $\alpha_{ref}$ is calibrated directly, priors ensure that $\alpha_{[j]}$ remains within biologically plausible limits (Table S15).

Reproductive does and bucks (Eq. S2.2a, b) are calculated by summing all susceptible and infected individuals by sex, weighted by age-specific reproductive probabilities ($a_{\alpha_{[l]}}$), which reflect reduced reproduction in fawns and yearlings (Green et al., 2017). Clinical deer are excluded from reproduction due to advanced disease effects (Almberg et al., 2011; Edmunds et al., 2018; Hamir et al., 2008; Jennelle et al., 2014; Wasserberg et al., 2009; Williams, 2005).

**Equation S3**

$$N_{\omega_{[j]}} = \left( \frac{\sum_{k=2}^{n}\sum_{l=1}^{n} S_{[j]} + \sum_{k=2}^{n}\sum_{l=1}^{n}\sum_{y=1}^{n} I_{[j]} + \sum_{k=2}^{n}\sum_{l=1}^{n} C_{[j]}}{\omega_G\left(\sum_{k=1}^{1}\sum_{l=1}^{n} S_{[r]} + \sum_{k=1}^{1}\sum_{l=1}^{n}\sum_{y=1}^{n} I_{[j]} + \sum_{k=1}^{1}\sum_{l=1}^{n} C_{[j]}\right)} \right)$$ **Equation S3.1 Total population within each region weighted for population pressure**

$$N_{[j]} = \sum_{k=1}^{n}\sum_{l=1}^{n} S_{[j]} + \sum_{k=1}^{n}\sum_{l=1}^{n}\sum_{y=1}^{n} I_{[j]} + \sum_{k=1}^{n}\sum_{l=1}^{n} C_{[j]}$$ **Equation S3.2 Total population within each region**

The weighted population size ($N_{\omega_{[j]}}$; Table S3.1) adjusts total abundance to reflect the lower resource demand of fawns (Equation S3), combining total adult population and the scaled contribution of fawns (via $\omega_G$). Total regional deer population size ($N_{[j]}$) is calculated by summing all deer across the susceptible ($S_{[j]}$), infected ($I_{[j]}$), and clinical ($C_{[j]}$) stocks of a given region (Equation. S3.2).

$$K_{[j]} = K_{ref_{[j]}}\left(X + (1 - X)G_{[j]}\right)$$ **Equation S4. Population pressure on deer within each region**

Population pressure ($K_{[j]}$; Eq. S4) reflects ecological limits on recruitment and mortality, akin to a traditional carrying capacity, and is computed as the product of the reference population pressure ($K_{ref_{[j]}}$) and the availability of both external (e.g., agriculture, food plots) and natural resources tracked as a vegetation index ($G_{[j]}$). In our model, population pressure is nominally 1 when population is at the halfway point between zero and the abundance at which population size is so large that recruitment is essentially zero and mortality is maximum. When external resources ($X$; Table S4) are zero, deer rely solely on natural vegetation; when X = 1, external resources fully determine $K_{[j]}$. The regional vegetation index ($G_{[j]}$) is dynamically modeled (refer to *Ecosystem dynamics*).

The reference population pressure for each region ($K_{ref_{[j]}}$; Equation. S5) is calculated as the product of the reference total population size ($N_{ref_{[j]}}$) and a region-specific pressure coefficient ($K_{\Delta_{[j]}}$), an estimated parameter which indicates whether the reference total population was at a level where population pressure was occurring. This sets a baseline at the simulation's start where the vegetation index is one and population pressure varies directly with food availability around the established reference. The reference total population size is derived from observed abundance estimates and model-fitted adjustments (refer to *Population estimates*). The reference deer density ($Y_{ref_{[j]}}$; Eq. S6) is calculated using the reference habitat area ($A_{ref_{[j]}}$; Eq. S7), which combines known deer range ($A_{N_{[j]}}$; Table S4) and a scaled contribution of surrounding landscape based on the habitat suitability parameter ($\omega_A$; Table S4) estimated during model fitting. Once the reference values are established during model optimization, they remain constant and allow us to see how relative changes from the start point influence the system.

$K_{ref_{[j]}} = N_{ref_{[j]}} K_{\Delta_{[j]}}$ **Equation S5. Reference population pressure for deer within each region**

$Y_{ref_{[j]}} = N_{ref_{[j]}} A_{ref_{[j]}}$ **Equation S6. Reference population of deer within each region**

$A_{ref_{[j]}} = A_{N_{[j]}} + \omega_A \left(A_{[j]} - A_{N_{[j]}}\right)$ **Equation S7. Reference area of deer habitat within each region**

**Table S4. Mapping between the full variable names used to construct the system dynamics model in Vensim (*Full Variable Name*) and the abbreviated forms (*Short Form*) used in the manuscript to represent the determination of population pressure in the system dynamics model.** The table includes each variable's units, the corresponding equation used to calculate it, or the source from which its value was derived.

| Short Form | Full Variable Name | Units | Equation/Source |
|---|---|---|---|
| $X$ | External resources | dimensionless | 0.5 |
| $G_{[j]}$ | Vegetation Index | dimensionless | Equation S16 |
| $A_{ref_{[j]}}$ | Ref area[Region] | mile$^2$ | Equation S7 |
| $A_{N_{[j]}}$ | Deer range 2021[Region] | mile$^2$ | Data (refer to *Data and processing* below) |
| $A_{[j]}$ | county land area[Region] | mile$^2$ | Data (refer to *Data and processing* below) |
| $\omega_A$ | nonrange weight | dimensionless | Estimated (Table S17) |
| $G_{max}$ | Max Veg Index | dimensionless | 1.5 |
| $t_R$ | Vegetation Regen Time | years | 20 |
| $B_{ref}$ | reference browsing rate | years$^{-1}$ | $t_R^{-1}$ |

Age advancement occurs once annually during the birth pulse ($t_{step}$ = 8), with individuals moving to the next age class. Those in the oldest cohort ($k$=7) remain until removed by mortality. Discrete aging minimizes the age-dispersion effects that would occur with a continuous process. Background mortality is modeled as continuous and includes causes such as senescence, predation, vehicle accidents, poaching, starvation, and non-CWD diseases. The background mortality rate for susceptible deer ($\mu_{S_{[j,k]}}$; Eq. S8) is calculated as the sum of a baseline mortality rate ($\mu_{S_{ref}}$) and an age-specific modifier ($a_{\mu_{[k]}}$), scaled by population pressure effects ($\varsigma_{\mu[j]}$) and winter severity ($W_\mu$) (Verme et al., 1968). Mortality is constrained above zero for senescence by applying a factor that limits the proportion of environmentally influenced variation ($f_\mu$).

$\mu_{S_{[j,k]}} = f_\mu \left(\mu_{S_{ref}} + a_{\mu_{[k]}}\right) \varsigma_{\mu_{[j]}} W_\mu$ **Equation S8. Background mortality rate of healthy deer**

**Equation S9**

$$a_{\mu_{[k]}} = \begin{cases} z_\mu & k = 1 \\ 0 & k > 1 \end{cases}$$ **Equation S9.1. Effect of age on deer mortality**

$$\varsigma_{\mu_{[j]}} = \left(\frac{N_{\omega_{[j]}}}{K_{[j]}}\right)^{s_\mu}$$ **Equation S9.2. Effect of population pressure on deer mortality**

$$W_\mu = \begin{cases} 1 & 3 \le t_{step} < 15 \\ 1 + s_W\left(\frac{W_c}{W_m}\right) & All\ other\ values \end{cases}$$ **Equation S9.3. Effect of winter severity on deer mortality**

The effect of age on deer mortality (Eq. S9.1) is only applied to fawns ($k$=1). The fawn excess mortality ($z_\mu$; Table S5) follows a two-phase model based on age: elevated mortality during the first three months post-birth ($a_{\mu_{max}}$), and a step-function to drop to a minimum thereafter ($a_{\mu_{min}}$), a distinction made based on deer survival studies (Rohm et al., 2007; Verme et al., 1968; Vreeland et al., 2004). The details of fawn survival have little practical effect on model dynamics but make comparisons with measured fawn-doe ratios more satisfying. The effect of population pressure on mortality ($\varsigma_{\mu_{[j]}}$; Eq. S9.2) increases proportionally to the weighted population size ($N_{\omega_{[j]}}$) relative to the population pressure ($K_{[j]}$), scaled by a sensitivity parameter ($s_\mu$; Table S5). Winter severity effects (Eq. S9.3) apply only during relevant timesteps (i.e., those associated with winter months) and increase mortality in proportion to the deviation of current conditions ($W_C$; Table S5) from the 2000–2020 average ($W_M$; Table S5), scaled by winter sensitivity ($s_W$; Table S5).

**Table S5. Mapping between the full variable names used to construct the system dynamics model in Vensim (*Full Variable Name*) and the abbreviated forms (*Short Form*) used in the manuscript to represent background mortality of deer and additional mortality due to chronic wasting disease in the system dynamics model.** The table includes each variable's units, the corresponding equation used to calculate it, or the source from which its value was derived.

| Short Form | Full Variable Name | Units | Equation/Source |
|---|---|---|---|
| $\mu_{S_{[j,k]}}$ | background mortality rate[Region,age] | fraction/year | Equation S8 |
| $\mu_{S_{ref}}$ | ref background mortality rate | fraction/year | Estimated (Table S17) |
| $f_\mu$ | frac mortality variable | Fraction/year | Estimated (Table S17) |
| $a_{\mu_{[k]}}$ | age excess mortality[age] | fraction/year | Equation S9.1 |
| $z_\mu$ | Fawn excess mortality | fraction/year | $= \begin{cases} a_{\mu_{max}} & 9 \le t_{step} < 14 \\ a_{\mu_{min}} & All\ other\ values \end{cases}$ |
| $\varsigma_{\mu[r]}$ | CC effect mortality[Region] | dimensionless | Equation S9.2 |
| $W_\mu$ | WSI effect mortality | dimensionless | Equation S9.3 |
| $a_{\mu_{max}}$ | early fawn mortality | fraction/year | Estimated (Table S17) |
| $a_{\mu_{min}}$ | Base fawn mortality | fraction/year | Estimated (Table S17) |
| $s_\mu$ | mortality CC sensitivity | dimensionless | Estimated (Table S17) |
| $s_W$ | mortality WSI sensitivity | dimensionless | Estimated (Table S17) |
| $W_c$ | time centered WSI | dimensionless | Data (refer to Data and processing below) |
| $W_m$ | mean WSI 2000 2020 | dimensionless | Data (refer to Data and processing below) |
| $\psi_I$ | infected excess mortality hazard | fraction/year | 0 |
| $\mu_C$ | CWD mortality rate | fraction/year | 2.5 |
| $\psi_C$ | clinical excess predation | fraction/year | 0 |

For mortality of infected deer (Eq. S1.2), mortality rate is the sum of background mortality rate ($\mu_{S_{[j,k]}}$; Eq. S8) and an added hazard from early disease-related impairments ($\psi_I$; Table S5) (Conner et al., 2000; DeVivo et al., 2017; Edmunds, 2013; Krumm et al., 2005). For clinical deer, mortality (Eq. S1.3) is governed by a fixed CWD mortality rate ($\mu_C$; Table S5), uniform across age due to the overwhelming lethality of the disease once clinical symptoms appear, and an additional hazard for clinical symptoms ($\psi_C$; Table S5). These rates are applied to the respective compartment populations to determine mortality-driven losses.

Additional mortality sources, including hunter harvest and agency removals (area-specific, targeted removal of deer), are described in the following section.

##### 2.2.1.2 Dispersal

The model structure permits susceptible and infected deer to disperse between regions; however, this effect is not presently active for estimation of the model. We assume that clinical deer do not disperse due to severely impaired motor function (Almberg et al., 2011; Edmunds et al., 2018; Hamir et al., 2008; Jennelle et al., 2014; Wasserberg et al., 2009; Williams, 2005). The influx of susceptible deer into a given region $j$ is calculated as the sum across all origin regions $j*$ of the product of the instantaneous dispersal rate ($\delta_{[j^*,j,k,l]}$) for each age and sex class and the number of susceptible deer ($S_{[j^*k,l]}$) in the origin region (Eq. S1.1). Likewise, the outflow of susceptible deer from region $j$ is computed as the sum across all destination regions $j*$ of the product of the dispersal rate from region $j$ to region $j*$ ($\delta_{[j,j*,k,l^*]}$) and the number of susceptible deer in the origin region ($S_{[j,k,l]}$). Infected deer disperse using the same framework, but with the number of infected individuals ($I_{[j,k,l,y]}$) instead of susceptible individuals (Eq. S1.2).

$$\delta_{[j^*,j,k,l]} = \delta_{ref}\delta_{\Im_{[k,l]}}\delta_{R_{[j*,j]}}$$ **Equation S10. Dispersal rate between regions**

$$\delta_{R_{[j^*,j]}} = \begin{cases} \frac{1}{\delta_{d_{[j*,j]}}^{\ 2}} & j^* \neq j \\ 0 & j^* = j \end{cases}$$ **Equation S11. The propensity of deer dispersal given distance between regions**

The dispersal rate ($\delta_{[j^*,j,k,l]}$; Eq. S10) is modeled as the product of three factors: the reference dispersal rate ($\delta_{ref}$; Table S6), the age- and sex-specific dispersal modifier ($\delta_{\Im_{[k,l]}}$; Table S6), and the propensity of deer to disperse as a function of relative distance between regions ($\delta_{R_{[j^*,j]}}$). We assume that females do not disperse between regions due to strong matriarchal social structure, and that only male yearlings disperse, typically over short distances (Long et al., 2008; Nixon et al., 2007; Skuldt et al., 2008). The propensity of yearling males to disperse is inversely proportional to the square of the relative distance between origin and destination regions (Eq. S11). The relative distance between regions ($\delta_{d_{[j*,j]}}$) is defined as the Euclidean distance between the centroid of region, $j$, to that of region, $j^*$. Dispersal does not occur within the same region (Table S6).

**Table S6. Mapping between the full variable names used to construct the system dynamics model in Vensim (*Full Variable Name*) and the abbreviated forms (*Short Form*) used in the manuscript to represent the dispersal of deer between regions in the system dynamics model.** The table includes each variable's units, the corresponding equation used to calculate it, or the source from which its value was derived.

| Short Form | Full Variable Name | Units | Equation/Source |
|---|---|---|---|
| $\delta_{[j^*,j,k,l]}$ | migration dispersal rate[FromRegion,ToRegion,age,sex] | deer/year | Equation S10 |
| $\delta_{ref}$ | ref migration dispersal rate | fraction/year | **user defined value between 0-0.1** |
| $\delta_{\Im_{[k,l]}}$ | age sex mobility[age, sex] | dimensionless | $= \begin{cases} 0 & l \neq 1 \\ 1 & k = 1 \ and \ l = 1 \end{cases}$ |
| $\delta_{R_{[j^*,j]}}$ | relative migration dispersal rate[FromRegion,ToRegion] | fraction | Equation S11 |
| $\delta_{d_{[j*,j]}}$ | relative distance[FromRegion,ToRegion] | dimensionless | Data (refer to *Data and processing* below) |

##### *2.2.1.3 Human caused mortality*

The entire harvest occurs in a single annual pulse at the timestep corresponding to mid-November through the second week of December ($t_{step}$=15), which aligns with the primary harvest period in Wisconsin (Wisconsin Department of Natural Resources, n.d.). Harvest within each stock is governed by an instantaneous harvest rate, which dictates the fraction of deer removed from each age and sex class. Harvest is calculated separately for three age-sex groupings: older bucks [$k$=5,6,7; $l$=1], young bucks [$k$= 2,3,4; $l$=1], all antlerless deer which includes does [$k$=1-7; $l$=2] and fawn bucks [$k$=1; $l$=1].

Total harvest accounts for deer lost to wounding, which adds an additional 7% above the registered harvest (Aebischer et al., 2014; Ditchkoff et al., n.d.; Fuller, 1990; Jennings et al., 2014; Pedersen et al., 2008; Wallingford et al., 2017). The number of deer removed via hunter harvest from each stock (Eq. S1.1, Eq. S1.2, Eq. S1.3) is calculated as the product of the number of deer in the stock ($S_{[j,k,l]}$); $I_{[j,k,l,y]}$; $C_{[j,k,l]}$) and the respective instantaneous harvest rate ($h_{S_{[j,k,l]}}$; $h_{I_{[j,k,l]}}$; $h_{C_{[j,k,l]}}$).

$$h_{S_{[j,k,l]}} = h_{[j,k,l]}$$ **Equation S12. The instantaneous harvest rate of susceptible deer**

$$h_{I_{[j,k,l]}} = h_{[j,k,l]} + h_{F_{[j,k,l]}}$$ **Equation S13. The instantaneous harvest rate of infected deer**

$$h_{C_{[j,k,l]}} = h_{F_{[j,k,l]}}$$ **Equation S14. The instantaneous harvest rate of clinical deer**

The instantaneous susceptible harvest rate (Eq. S12) is equal to the base instantaneous harvest rate ($h_{[j,k,l]}$). The instantaneous infected harvest rate (Eq. S13) can be elevated when infection-targeted harvest is active, through the addition of a user-defined rate ($h_{F_{[j,k,l]}}$; Table S7). Clinical deer are assumed to be visibly ill and therefore typically avoided during standard hunting activity. As such, clinical deer are not removed via hunter harvest unless an active infection-targeted harvest is implemented (Eq. S14), in which case the harvest rate matches the infection-targeted harvest rate ($h_{F_{[j,k,l]}}$).

The base harvest rate ($h_{[j,k,l]}$) is determined either by harvest data (over history) or by a user-defined target harvest rate (in the future or counterfactual scenarios, Eq. S15). Historical harvest rates ($h_{H_{[j,k,l]}}$) are derived from annual time series data on antlered and antlerless harvests (categories 1 and 2 for antlered, category 3 for antlerless) and demographic data (refer to *Data processing*). The user may optionally specify a target harvest rate ($h_{g_{[j,k,l]}}$) between zero and one for any age and sex category (Table S7). The applied harvest rate for each year depends on the intervention switch ($\Psi_{[j]}$), a binary variable where a value of one indicates that a management intervention has occurred and a value of zero indicates no intervention. When $\Psi_{[j]} = 0$, the model uses the historical harvest rate, when $\Psi_{[j]}$=1, the target harvest rate is applied (Eq. S15). If no target rate is specified but the simulation is beyond the existing time series for the historical rate, then the historical rate smoothed over the three most recent years is used.

$$h_{[j,k,l]} = \begin{cases} h_{H_{[j,k,l]}} h_{w_{[a]}} p_{H_{[j]}} & \Psi = 0 \\ h_{g_{[j,k,l]}} h_{w_{[a]}} p_{H_{[j]}} & \Psi = 1 \end{cases}$$

**Equation S15. The instantaneous harvest rate for each age and sex category of deer**

To better reflect hunter preferences, the base harvest rate ($h_{H_{[j,k,l]}}$ or $h_{g_{[j,k,l]}}$) is further modified by an age-specific weight ($h_{w_{[k]}}$), which reduces the likelihood of fawn harvest (Wojcik and Stenglein, n.d.).

**Table S7. Mapping between the full variable names used to construct the system dynamics model in Vensim (*Full Variable Name*) and the abbreviated forms (*Short Form*) used in the manuscript to represent the harvest of deer by hunters in the system dynamics model.** The table includes each variable's units, the corresponding equation used to calculate it, or the source from which its value was derived.

| Short Form | Full Variable Name | Units | Equation/Source |
|---|---|---|---|
| $h_{S_{[j,k,l]}}$ | healthy harvest rate[Region,age,sex] | fraction/year | Equation S12 |
| $h_{I_{[j,k,l]}}$ | infected harvest rate[Region,age,sex] | fraction/year | Equation S13 |
| $h_{C_{[j,k,l]}}$ | clinical harvest rate[Region,age,sex] | fraction/year | Equation S14 |
| $h_{[j,k,l]}$ | Harvest Rate[Region,age,sex] | fraction/year | Equation S15 |
| $h_{F_{[j,k,l]}}$ | active infection targeted harvest rate[Region] | fraction/year | $= \begin{cases} 0 & \Psi_{[r]} = 0 \\ * & \Psi_{[r]} = 1 \end{cases}$ ***user defined value between 0-1** |
| $\Psi_{[j]}$ | Intervention switch[Region] | dimensionless | **user defined value of 0 or 1** |
| $h_{\omega_{[k]}}$ | age harvest weight[age] | dimensionless | [Estimated Table S17, 1, 1, 1,1,1,1] |
| $h_{H_{[j,k,l]}}$ | historic buck harvest rate[Region] for *k*=2,3,4,5,6,7 and *l*=1<br><br>historic antlerless harvest rate[Region] for *k*=1,2,3,4,5,6,7 and *l*=2 OR *k*=1; *l*=1 | fraction/year | *Historic harvest rates* |
| $h_{g_{[j,k,l]}}$ | final young buck harvest rate[Region] for *k*=2,3,4 and *l*=1<br><br>final older buck harvest rate[Region] for *k*=5,6,7 and *l*=1<br><br>final antlerless harvest rate[Region] for *k*=1,2,3,4,5,6,7 and *l*=2 OR *k*=1 and *l*=1 | fraction/year | *Historic harvest rates* |

When management intervention is active in a region ($\Psi_{[j]}$=1), additional mortality may result from agency-directed agency removals, which occur in at a user-defined target rate in a pulse at timestep 3 during the Winter. The number of susceptible deer removed by agency removals (Eq. S1.1) is calculated as the product of the instantaneous agency removals rate for susceptible deer ($m_{S[j,k,l]}$) and the total number of susceptible deer ($S_{[j,k,l]}$). The number of infected and clinical deer removed via agency removals is calculated as the product of the instantaneous infected agency removals rate ($m_{I[j,k,l]}$) and the total number of infected ($I_{[j,k,l,y]}$) or clinical ($C_{[j,k,l]}$) deer respectively (Eq. S1.2, S1.3). The instantaneous agency removals rates are informed by the agency's target effort, the age and sex structure of the population and specific prevalence, and the size of the management region (refer to *Agency removals*).

##### 2.2.2 Ecosystem dynamics

The vegetation index is a dimensionless stock representing the availability of natural deer browse, normalized to condition at the start of the simulations when it evaluates to one. Its dynamics are governed by the balance between deer browsing and vegetation regeneration within each region (Eq. S16). Regeneration is modeled as a return of the vegetation index toward a region-specific maximum value in the absence of browsing ($G_{max}$; Table S4). The rate of this return depends on the average vegetation regeneration time ($t_R$; Table S4) and the current vegetation index. Specifically, the fractional regeneration rate decreases linearly from one when the index is equal to one, to zero when the index reaches $G_{max}$, such that regeneration slows as the system approaches the vegetation carrying capacity.

$$\frac{dG_{[j]}}{dt} = \overbrace{\frac{1}{t_R}\left(\frac{1}{1-\frac{1}{G_{max}}}\right)\left(\frac{1-G_{[j]}}{G_{max}}\right)G_{[j]}}^{Regeneration} - \overbrace{B_{ref}\frac{N_{\omega[j]}}{K_{ref[j]}}G_{[j]}}^{Browsing}$$

**Equation S16. Change in the vegetation index**

The browsing rate within each region is defined as the product of the current vegetation index, the reference browsing rate ($B_{ref}$), and the ratio of the weighted deer population ($N_{\omega_{[j]}}$; Table S3) to the reference population pressure ($K_{ref}$; Eq. S5). Here, $B_{ref}$ is equivalent to the inverse of the vegetation regeneration time and reflects the browsing pressure per deer when the vegetation index is one. The weighted deer population incorporates age-specific impacts, reducing the contribution of fawns to total browsing pressure. The reference population pressure ($K_{ref}$) represents the deer population size at which vegetation remains stable (i.e., the vegetation index is maintained at one). When the current population exceeds this reference, browsing surpasses regeneration and vegetation declines; when it is below the reference, regeneration outpaces browsing and the vegetation index increases.

##### 2.2.3 Epidemiology

###### *2.2.3.1 Transmission pathways*

Susceptible deer transition to the infected class through a disease transmission flow defined as the product of the force of infection ($\lambda_{[j,k,l]}$) and the number of susceptible deer ($S_{[j,k,l]}$) within each age and sex class in each region (Eq. S1.1, S1.2). The force of infection is an endogenous variable that varies by region, age class, and sex (Eq. S17), and consists of three components: horizontal transmission, maternal transmission, and indirect (environmental) transmission.

$$\lambda_{[j,k,l]} = \lambda_{F_{[j,k,l]}} + \lambda_{V_{[j,k,l]}} + \lambda_{D_{[j,k,l]}}$$

**Equation S17 Total force of infection**

The horizontal force of infection ($\lambda_{F_{[j,k,l]}}$; Eq. S18.1) and the indirect force of infection ($\lambda_{V_{[j,k,l]}}$; Eq. S18.2) represent instantaneous rates of infection via contact with infectious deer (Mathiason et al., 2006; Tamgüney et al., 2009) and contact with infectious prions in the environment (Almberg et al., 2011; Miller et al., 2006, 2004), respectively, within each demographic cohort and region. The maternal force of infection ($\lambda_{D_{[j,k,l]}}$; Eq. S18.3) captures the rate at which infectious does transmit the disease to their offspring at birth (Miller et al., 2000).

**Equation S18**

$$\lambda_{F_{[j,k,l]}} = \beta_F c_{F_{[j,k,l]}} \frac{F_{[j]}}{N_{[j]}} \left( \varrho_{\lambda_{[j]}} \phi_{\lambda_{[j]}} \right)$$ **S18.1 Force of infection from direct horizontal transmission**

$$\lambda_{V_{[j,k,l]}} = \beta_V c_{V_{[j,k,l]}} \frac{V_{[j]}}{V_{ref_{[j]}}} \left( \varrho_{\lambda_{[j]}} \phi_{\lambda_{[j]}} \right)$$ **S18.2 Force of infection from indirect environmental transmission**

$$\lambda_{D_{[j,k,l]}} = \begin{cases} \beta_D \frac{D_{F_{[j]}}}{D_{[j]}} & k = 1 \\ 0 & k > 1 \end{cases}$$ **S18.3 Force of infection from direct maternal transmission**

Horizontal transmission is partially driven by the proportion of infectious individuals ($F_{[j]}$; Eq. S29) within the total regional population ($N_{[j]}$), multiplied by the base horizontal transmission rate ($\beta_F$; Table S8) -defined as the product of contact rate between infectious and susceptible individuals and the probability of infection given that contact- and an age- and sex-specific modifier of the effective contact rate ($c_{F_{[j,k,l]}}$). This formulation allows the force of infection to vary across age and sex classes within each region and to respond dynamically to changing population structure and disease prevalence. Indirect transmission is driven by the environmental prion load ($V_{[j]}$), normalized by the reference prion load at the start of the simulation ($V_{ref_{[j]}}$). This normalization allows the model to express environmental prion pressure as a relative index without requiring explicit units. The resulting prion load ratio is multiplied by the indirect transmission rate ($\beta_V$; Table S8) -defined as the product of contact rate between infectious prions and susceptible individuals and the probability of infection given that contact- and an age- and sex-specific modifier of the effective environmental contact rate ($c_{V_{[j,k,l]}}$).Maternal transmission occurs only within the fawn age class ($k$=1). Fawns are born to the susceptible stock but face an elevated exposure risk beyond the other age classes equivalent to the product of the general maternal transmission ($\beta_D$) and the regional prevalence of disease within the reproductive doe population ($\frac{D_{F_{[j]}}}{D_{[j]}}$; Table S8). Additional parameters allow both the direct and indirect components of the force of infection to be modulated by host behavior, disease characteristics, ecosystem features, or social dynamics.

###### *2.2.3.2 Modifiers of transmission*

Relative contact rate parameters $c_{F_{[j,k,l]}}$ and $c_{V_{[j,k,l]}}$ are embedded within the direct (Eq. 18.1) and indirect (Eq. S18.2) force of infection rates respectively. These modifiers are endogenously modeled (Eq. S19) as a product of the host, disease, and human-related factors that influence contact rates, and thus shape the force of infection.

**Equation S19**

$$c_{F[j,k,l]} = \left[\kappa_{F_{[k,l]}} \frac{n_k n_l}{\left(\sum_{k=1}^{n}\sum_{l=1}^{n}\kappa_{F_{[k,l]}}\right)}\right]\left[1-\varphi_{F_{[j]}}\right]\left[\frac{N_{[j]}}{\Omega_F N_{ref_{[j]}}+(1-\Omega_F)N_{[j]}}\right]\left[\frac{N_{ref_{[j]}}}{\left(\Omega_{abs}A_{ref_{[j]}}Y_{ref}+(1-\Omega_{abs})N_{ref_{[j]}}\right)}\right]$$ **S19.1 Relative direct contact rate between deer**

$$c_{V[j,k,l]} = \kappa_{V_{[k,l]}}\left[1-\varphi_{V_{[j]}}\right]\left[\frac{N_{[j]}}{\Omega_V N_{ref_{[j]}}+(1-\Omega_V)N_{[j]}}\right]\left[\frac{N_{ref_{[j]}}}{\left(\Omega_{abs}A_{ref_{[j]}}Y_{ref}+(1-\Omega_{abs})N_{ref_{[j]}}\right)}\right]$$ **S19.2 Relative indirect contact rate between deer and environment**

Age- and sex-related heterogeneities in horizontal (deer-to-deer) and environmental (deer-to-prion) effective contact rates are incorporated through age-sex contact effect parameters: $\kappa_{F_{[k,l]}}$ for direct transmission (Eq. S19.1) and $\kappa_{V_{[k,l]}}$ for indirect transmission (Eq. S19.2). These parameters account for behavioral patterns such as increased female-female contacts within matriarchal groups (Grear et al., 2006; Osnas et al., 2009; Potapov et al., 2013) and potential size-related differences in environmental exposure. To maintain comparability across cohorts, the age-sex contact effects are centered around one by scaling each parameter by the ratio of the product of total number of sex ($n_l$) and age ($n_k$) classes to the sum of all cohort-specific contact effects (Eq. S19.1 [first term]).

Human-mediated influences, such as baiting or supplemental feeding (Sorensen et al., 2014), are captured through regional social dynamic modifiers in both direct ($\varphi_{F_{[j]}}$) and indirect ($\varphi_{V_{[j]}}$) transmission. These modifiers activate only under specific management scenarios ($\Psi_{[j]}$=1) and allow scenario testing of human behaviors that increase or decrease contact rates. A value of zero implies no effect, while values above or below one increase or reduce transmission risk, respectively (Table S8). The effect size of policies such as a baiting and feeding ban is presently unknown, confounded by unobserved compliance, and can't be estimated with available data.

The effect of deer population density on transmission is divided into two components: relative density and absolute density. Relative density captures longitudinal changes within each region (third term, Eq. S19.1, S19.2), modeled as the ratio of the current deer population ($N_{[j]}$) to the regional reference population ($N_{ref_{[j]}}$; Eq. S6). Its influence is modulated by global deer density-dependence coefficients ($\Omega_F$ for direct and $\Omega_V$ for indirect transmission), which are estimated during calibration (refer to *Estimation method*). These coefficients allow the model to flexibly estimate the transmission mode between a fully frequency-dependent system (value of 0) and a fully density dependent system (value of 1) to explore the full range of findings in the literature (Almberg et al., 2011; Jennelle et al., 2014; Joly et al., 2003; Wasserberg et al., 2009).

Absolute density effects are fixed at the first time-step and remain constant throughout the simulation (fourth term, Eq. S19.1, S19.2). This accounts for the use of reference values, which can mask cross-sectional variation in regional density. It is derived from the deviation of the empirically estimated reference population ($N_{ref_{[j]}}$) from a baseline computed as the product of the reference area ($A_{ref_{[j]}}$)) and reference density ($Y_{ref}$), scaled by the global absolute density dependence parameter ($\Omega_{abs}$).

**Equation S20**

$$\kappa_{F_{[k,l]}} = \begin{cases} \sigma_{a_{[k]}} & l = 1 \\ \sigma_{s_{[l]}}\left(1 + \left(\sigma_{a_{[k]}} - 1\right)\upsilon_{sa_{[l]}}\right) & l \neq 1 \end{cases}$$ **Equation S20.2 The effect of age and sex on deer to deer contact rates**

$$\kappa_{V_{[k,l]}} = (1 - \sigma_V) + \sigma_V\left[\kappa_{F_{[k,l]}}\frac{n_k n_l}{(\sum_{k=1}^{n}\sum_{l=1}^{n}\kappa_{F_{[k,l]}})}\right]$$ **Equation S20.1 The effect of age and sex on deer to environment contact rates**

The age-sex contact effects on transmission are endogenously computed (Eq. S20). The effect of age and sex on direct effective contacts, $\kappa_{F_{[k,l]}}$, is calculated from the specific marginal effects of age ($\sigma_{a_{[k]}}$) and sex ($\sigma_{s_{[l]}}$), along with a parameter that represents the relative importance of sex versus age in shaping intraspecific avoidance behavior ($\upsilon_{sa_{[l]}}$). This structure allows sex to mediate the impact of age across cohorts. Fawns are assumed to show no sex-based differences in contact rates (Eq. S20.1). For all other ages, the equation is formulated so that as the relative impact of sex moves towards zero, the effect of age on contacts and disease risk is ignored by the model (Eq. S20.1). Adult bucks serve as the reference group (i.e., $\upsilon_{sa_{[l=1]}}$=1), and the comparative effect in does is estimated during model calibration (Table S8).

For environmental contacts, the age-sex effect ($\kappa_{V_{[k,l]}}$) is calculated as a weighted average that shifts between the constant sex effects and the constant age effects based on the dispersion of sexes and ages within each age-sex cohort. We estimate the difference in sex-age effects on environmental contacts compared to direct contacts ($\sigma_V$) such that a value of one would result in the same age-sex effect on indirect contacts as in direct (Eq. S20.2).

Potential influences on the probability of infection given contact due to genetic variation (Blanchong et al., 2009; Johnson et al., 2006)—specifically, the frequency of the *S* allele at the 286G/A (G96S) coding polymorphism—are captured by the genetic transmission effect modifier ($\varrho_{\lambda_{[j]}}$; Eq. 21). Note that the effects could be either on the infectiousness of the infected deer or the resistance of the susceptible deer.

$$\varrho_{\lambda_{[j]}} = \varrho_{[j]}\varrho_T + (1 - \varrho_{[j]})$$ **Equation S21. The effect of genetics on transmission**

The genetic effect on transmission (Eq. S21) is governed by the regional frequency of the *S* allele ($\varrho_{[j]}$; Eq. S25) and the strength of the genetic effect ($\varrho_T$; Table S8). The effect of genetics on transmission rates is high when the frequency of the favored allele is low and reaches a minimum equivalent to the strength of the genetic effect on transmission when all deer within a population have the allele (Eq. S21).

We incorporate a hierarchical approach by including a relative transmission modifier ($\phi_{\lambda_{[j]}}$) in the force of infection equations (Eq. S18.1, S18.2), enabling the model to account for unexplained inter-regional variation in transmission dynamics, analogous to "random effects" in statistical models.

**Table S8. Mapping between the full variable names used to construct the system dynamics model in Vensim (*Full Variable Name*) and the abbreviated forms (*Short Form*) used in the manuscript to represent the transmission of chronic wasting disease in the system dynamics model.** The table includes each variable's units, the corresponding equation used to calculate it, or the source from which its value was derived.

| Short Form | Full Variable Name | Units | Equation/Source |
|---|---|---|---|
| $\lambda_{[j,k,l]}$ | Infection Rate FOI[Region,age,sex] | fraction/year | Equation S17 |
| $\lambda_{F_{[j,k,l]}}$ | Direct Infection Rate[Region,age,sex] | fraction/year | Equation S18.1 |
| $\lambda_{V_{[j,k,l]}}$ | Indirect Infection Rate[Region,age,sex] | fraction/year | Equation S18.2 |
| $\lambda_{D_{[j,k,l]}}$ | Vertical Infection Rate[Region,age,sex] | fraction/year | Equation S18.3 |
| $\beta_F$ | Direct beta | 1/year | Estimated (Table S17) |
| $c_{F_{[j,k,l]}}$ | relative contact rate[Region,age,sex] | dimensionless | Equation S19.1 |
| $\varrho_{\lambda_{[j]}}$ | gen eff transmission rate[Region] | dimensionless | Equation S21 |
| $\phi_{\lambda_{[j]}}$ | Relative transmission[Region] | dimensionless | Estimated (Table S17) |
| $\beta_V$ | Indirect beta | 1/year | Estimated (Table S17) |
| $c_{V_{[j,k,l]}}$ | envir relative contact rate[Region,age,sex] | dimensionless | Equation S19.2 |
| $\beta_D$ | Vertical transmission beta | 1/year | Estimated (Table S17) |
| $D_{F_{[j]}}$ | infectious reproductive population[Region, does] | deer | $\sum_{k=1}^{n}\sum_{l=}^{2}\sum_{y=1}^{n} I_{[j,k,l,y]}a_{\alpha_{[k]}}$ |
| $\kappa_{F_{[k,l]}}$ | age sex effect contacts[age,sex] | dimensionless | Equation S20.1 |
| $\kappa_{V_{[k,l]}}$ | envir age sex effect contacts[age,sex] | dimensionless | Equation S20.2 |
| $n_k$ | ELMCOUNT(age) | dimensionless | 7 |
| $n_l$ | ELMCOUNT(sex) | dimensionless | 2 |
| $\Omega_F$ | density dependence | dimensionless | Estimated (Table S17) |
| $\Omega_{abs}$ | abs density dependence | dimensionless | Estimated (Table S17) |
| $\varphi_{F_{[j]}}$ | contact reduction[Region] | dimensionless | $= \begin{cases} 0 & \Psi_{[j]} = 0 \\ * & \Psi_{[j]} = 1 \end{cases}$ ***user defined value between -1 to 1** |
| $\Omega_V$ | envir density dependence | dimensionless | Estimated (Table S17) |
| $\varphi_{V_{[j]}}$ | envir contact reduction[Region] | dimensionless | $= \begin{cases} 0 & \Psi_{[j]} = 0 \\ * & \Psi_{[j]} = 1 \end{cases}$ ***user defined value between -1 to 1** |
| $\sigma_{s_{[l]}}$ | sex eff contacts[sex] | dimensionless | Estimated (Table S17) |
| $\sigma_{a_{[k]}}$ | age eff contacts[age] | dimensionless | Estimated (Table S17) |
| $\sigma_V$ | relative envir contact diversity | dimensionless | Estimated (Table S17) |
| $\upsilon_{sa_{[l]}}$ | sex rel age dispersion[sex] | dimensionless | $= \begin{cases} 1 & l = 1 \\ Estimated\ (Table\ S15) & l = 2 \end{cases}$ |
| $\varrho_T$ | gen effect transmission | dimensionless | Estimated (Table S17) |

##### 2.2.3.3 *Disease progression*

Once infected, deer progress through six stages of disease within the infected compartment, ultimately transitioning to the clinical compartment after the final stage ($y$=6). The transition of deer between stages is calculated as the product of the progression rate ($\rho_{[j,y]}$) and number of infected individuals ($I_{[j,k,l,y]}$) in each cohort and infection stage within a region (Eq. S1.2, S1.3). The instantaneous progression rate (Eq. S22) is defined as the quotient of the reference progression rate for each stage ($\rho_{ref_{[y]}}$) and the genetic progression effect modifier ($\varrho_{\rho_{[j,y]}}$). The genetic progression effect captures how the frequency of the $S$ allele at codon 286 (G96S) of the prion protein (PRNP) gene influences progression through the exposed and infectious stages (Table S9) after infection, and ultimately affects the lifespan of CWD-positive deer (Johnson et al., 2011).

$$\rho_{[j,y]} = \frac{\rho_{ref[y]}}{\varrho_{\rho[j,y]}}$$ **Equation S22. The progression rate of deer through the infected stages to the clinical stock**

The reference progression rate (Eq. S23) is calculated separately for the exposed and infectious stages by evenly dividing the total duration of each stage across the number of sub-stages in each ($n_E$, $n_F$; Table S9). The total durations are based on the expected durations between exposure and infectiousness ($t_E$; Table S9) (Almberg et al., 2011; Hanley et al., 2022; Kjær and Schauber, 2022; Wasserberg et al., 2009), and the expected amount of time between becoming infectious and progressing to the clinical stage ($t_F$; Table S9) (Al-arydah et al., 2016; Hamir et al., 2008; Mathiason, 2023; Oraby et al., 2014) respectively.

$$\rho_{ref[y]} = \begin{cases} \frac{n_E}{t_E} & y = 1,2,3 \\ \frac{n_F}{t_F} & y = 4,5,6 \end{cases}$$ **Equation S23. The reference progression rate of infected deer through each stage of infection**

The regional genetic progression modifier ($\varrho_{\rho_{[j,y]}}$) is computed using the regional frequency of the *S* allele ($\varrho_{[j]}$) and the estimated genetic impact on lifespan in the exposed ($\varrho_E$) and infectious ($\varrho_F$) stages (Eq. S24.1). Studies show that deer with the *S* allele tend to have longer survival after CWD exposure (Hamir et al., 2008; Hoover et al., 2017; Johnson et al., 2011; Robinson et al., 2012). The total genetic effect on total lifespan ($\varrho_L$; Eq. S24.2) is calculated using these stage-specific modifiers and the initial model-based proportion of time spent in each the exposed and infectious stages ($f_E$). A prior is placed on the magnitude of this genetic effect to integrate both data and expert knowledge during model calibration (refer to *Estimation method*).

**Equation S24**

$$\varrho_{\rho_{[j,y]}} = \begin{cases} \varrho_{[j]}\varrho_E + (1 - \varrho_{[j]}) & y = 1,2,3 \\ \varrho_{[j]}\varrho_F + (1 - \varrho_{[j]}) & y = 4,5,6 \end{cases}$$ **Equation S24.1 The genetic effect progression regional modifier**

$$\varrho_L = f_E\varrho_E + (1 - f_E)\varrho_F$$ **Equation S24.2 The genetic effect on total infected lifespan**

The *S* allele frequency in each region ($\varrho_{[j]}$) is modeled as a stock (Eq. S25) which moves towards a disease mediated equilibrium frequency ($\varrho_{Q_{[j]}}$). The change is proportional to the product of the current allele frequency and the difference between the current and equilibrium values, divided by the time scale for natural selection ($t_\varrho$; Table S9). The initial proportion of time infected deer spent in the exposed stage ($f_E$; Eq. S26) is calculated as the implied time duration spent in all exposed sub-stages divided by the implied time duration spent across all infected sub-stages during model initialization (Table S9).

$$\frac{d\varrho_{[j]}}{dt} = \frac{\varrho_{[j]}(\varrho_{Q_{[j]}} - \varrho_{[j]})}{t_\varrho}$$ **Equation S25. Change in the frequency of the *S* allele**

$$f_E = \frac{t_E}{(t_E + t_F)}$$ **Equation S26. The initial proportion of time infected deer spend as exposed rather than infectious**

The equilibrium frequency ($\varrho_{Q[j]}$; Eq. S27) is an endogenous variable reflecting the *S* allele frequency that natural selection favors, based on pre-CWD base frequency ($\varrho_{base}$; Table S9) and the current infection level in the region ($\zeta_{[j]}$). The equilibrium frequency increases as a function of prevalence and the difference between the base and biologically constrained maximum frequency ($\varrho_{max}$=1), scaled by a threshold prevalence ($\zeta_T$).

$$\varrho_{Q_{[j]}} = \varrho_{base} + (\varrho_{max} - \varrho_{base})\frac{\zeta_{[j]}}{(\zeta_T+\zeta_{[j]})}$$

**Equation S27. The desired natural equilibrium frequency of *S* allele in a population**

True prevalence within a region is calculated as the total number of infected deer divided by the total population (Eq. S28). This differs from the reported prevalence, which is based on positive test results. While true prevalence cannot be directly observed in natural systems due to sampling limitations and imperfect diagnostics (Mysterud et al., 2023; Viljugrein et al., 2021, 2019), modeling enables its estimation, offering valuable comparisons with the reported prevalence generated by surveillance data.

$$\zeta_{[j]} = \frac{\left(\sum_{k=1}^{n}\sum_{l=1}^{n}\sum_{y=1}^{n} I_{[r]}+\sum_{k=1}^{n}\sum_{l=}^{n} C_{[j]}\right)}{N_{[j]}}$$

**Equation S28. The true prevalence of chronic wasting disease in a population**

**Table S9. Mapping between the full variable names used to construct the system dynamics model in Vensim (*Full Variable Name*) and the abbreviated forms (*Short Form*) used in the manuscript to represent the progression of chronic wasting disease in infected deer in the system dynamics model.** The table includes each variable's units, the corresponding equation used to calculate it, or the source from which its value was derived.

| Short Form | Full Variable Name | Units | Equation/Source |
|---|---|---|---|
| $\rho_{[j,y]}$ | Progression rate[region, stage] | fraction/year | Equation S22 |
| $\rho_{ref_{[y]}}$ | ref Progression rate[stage] | fraction/year | Equation S23 |
| $\varrho_{\rho_{[j,y]}}$ | gen eff I lifespan[region] | dimensionless | Equation S24.1 *y*=1,2,3 |
| | gen eff E lifespan[region] | dimensionless | Equation S24.1 *y*=4,5,6 |
| $\varrho_E$ | genetic effect lifespan E | dimensionless | Estimated (Table S17) |
| $\varrho_F$ | genetic effect lifespan I | dimensionless | Estimated (Table S17) |
| $n_E$ | E stage | dimensionless | 3 |
| $n_F$ | I stage | dimensionless | 3 |
| $t_E$ | Exposed duration | years | Estimated (Table S17) |
| $t_F$ | Infectious duration | years | Estimated (Table S17) |
| $\varrho_L$ | gen eff lifespan | dimensionless | Equation S24.2 |
| $\varrho_{[j]}$ | Resistance Freq[Region] | fraction | Equation S25 |
| $f_E$ | E share initial duration | fraction | Equation S26 |
| $t_\varrho$ | selection time | years | Estimated (Table S17) |
| $\varrho_{Q_{[j]}}$ | equil resistance freq[Region] | fraction | Equation S27 |
| $\varrho_{base}$ | base frequency | fraction | 0.15 |
| $\varrho_{max}$ | max resistance freq | fraction | 1 (Biological limit) |
| $\zeta_{[j]}$ | true prevalence[Region] | fraction | Equation S28 |
| $\zeta_T$ | threshold prevalence for resistance | fraction | 0.1 |

##### 2.2.3.4 *Environmental prion dynamics*

The stock of environmental prions in each region ($V_{[j]}$) represents the total infectious prions present on the landscape (Eq. S1.4). Prions are deposited into the environment through two primary processes: shedding by live, infectious deer and decomposition of infectious carcasses. The deposition rate is governed by the annual amount of prions shed by each infectious deer ($\varepsilon_F$;

Table S10), and this is modified by the relative prion contribution of decomposing carcasses ($\varepsilon_Z$; Table S10). The total number of infectious deer in each region ($F_{[j]}$) is calculated as the sum of all clinical deer and all infectious deer (Eq. S29), those in the final three stages of the infected stock ($y$=4,5,6). The total amount of infectious carcasses on the landscape ($Z_{F_{[j]}}$) is further scaled by scavenging or decay ($t_Z$; Table S10), and management actions to remove them ($\varsigma_\Psi$; Table S10). The stock of environmental prions is depleted by the degradation of prions in the environment ($V_{[j]}$) at a rate dictated by the average prion infectious lifespan ($\tau$; Eq. S30). This process is first-order, though in reality prion deposition, transport and availability may involve multiple stages.

$$\frac{dV_{[j]}}{dt} = \overbrace{\Big(\underbrace{\varepsilon_F F_{[j]}}_{\text{Infectious live deer}} + \underbrace{\varepsilon_F \varepsilon_Z \frac{Z_{F_{[j]}}}{t_Z}(1-\varsigma_\Psi)}_{\text{Infectious decaying carcasses}}\Big)}^{\textit{Depositing}} - \overbrace{\tau V_{[j]}}^{\textit{Degrading}}$$

**Equation S1.4 Change in amount of environmental prions**

Environmental prions degrade over time and lose their infectiousness at a rate determined by the average prion lifespan ($\tau$). The prion lifespan is calculated as the prion half-life ($\tau_{hl}$; Table S10) over the natural logarithm of two, accounting for exponential decay (Eq. S30). This relationship ensures that prion decay is modeled with biologically appropriate timing and decay behavior.

$$F_{[j]} = \sum_{k=1}^{n}\sum_{l=1}^{n}\sum_{y=4}^{n} I_{[j]} + \sum_{k=1}^{n}\sum_{l=1}^{n} C_{[j]}$$

**Equation S29. The total number of live, infectious deer within each region**

$$\tau = \frac{\tau_{hl}}{\ln 2}$$

**Equation S30. The lifespan of a viable, infectious prion on the landscape**

The number of infectious carcasses on the landscape in each region ($Z_{F_{[j]}}$) is tracked dynamically (Eq. S31). This stock increases due to carcasses from infectious deer that died naturally or through harvest when carcasses remain on the landscape. The infectious deer left on the landscape due to natural mortality (Eq. S31 line 1) is the total number of the infectious deer within the last three stages of the infected stock ($y$=4,5,6) multiplied by their total natural mortality, summed with the total number of clinical deer multiplied by their natural mortality rate (refer to *Births and background mortality*). The infectious deer left on the landscape due to harvest (Eq. S31 line 2) is derived from the total number of harvested infectious deer, adjusted according to how they are handled post-harvest.

Infectious carcasses ($Z_{F_{[j]}}$) are removed from the landscape through natural decay (Eq. S31 line 4), or management intervention (Eq. S31 line 3). The rate of decay is based on the average carcass lifetime on the landscape ($t_Z$; Table S10), while the rate of carcass collection is determined by the proportion collected through active management ($\varsigma_\Psi$; Table S10). This proportion is user-defined and can be used to explore management impacts. Additionally, some carcasses are removed quickly if they result from specific types of non-harvest mortality, such as vehicular collisions. This is modeled using a separate collection parameter ($\varsigma_U$; Table S10).

$$\frac{dZ_{F_{[j]}}}{dt} = \overbrace{\left[\sum_{k=1}^{n}\sum_{l=1}^{n}\sum_{y=4}^{n}(\mu_{S_{[j,k]}} + \psi_I)I_{[j,k,l,y]} + \sum_{k=1}^{n}\sum_{l=1}^{n}(\mu_C + \psi_C)C_{[j,k,l]}\right]}^{\textit{Infectious deer background mortality}}$$

$$\overbrace{+[\underbrace{\frac{z_{H_{[j]}}}{t_H}}_{\textit{Carcass handling}} - \underbrace{\frac{z_{H_{[j]}}}{t_H}\varsigma_{T_{[j]}}}_{\textit{Carcass transport}} - \underbrace{\frac{z_{H_{[j]}}}{t_H}\varsigma_{\iota_{[j]}}\left(1 - \varsigma_{T_{[j]}}\right)}_{\textit{Safe carcass disposal}} + \underbrace{\mho\left(\textstyle\sum_{k=1}^{n}\sum_{l=1}^{n} h_{S[j,k,l]}S_{[j,k,l]} + \sum_{k=1}^{n}\sum_{l=1}^{n}\sum_{y=4}^{n} h_{I[j,k,l]}I_{[j,k,l,y]}\right)}_{\textit{Carcass left on landscape after field dressing}}]}^{\textit{Harvested infectious deer left on the landscape}}$$

$$\overbrace{-[\varsigma_\Psi \frac{z_{F_{[j]}}}{t_Z} + \left(\textstyle\sum_{k=1}^{n}\sum_{l=1}^{n}\sum_{y=4}^{n}(\mu_{S_{[j,k]}} + \psi_I)I_{[j,k,l,y]} + \sum_{k=1}^{n}\sum_{l=1}^{n}(\mu_C + \psi_C)C_{[j,k,l]}\right)\varsigma_U]}^{\textit{Infectious deer carcasses collected off the landscape}}$$

$$\overbrace{-\frac{z_{F_{[j]}}}{t_Z}(1 - \varsigma_\Psi)}^{\textit{Infectious carcass decay}}$$

**Equation S31. Change in total infectious carcasses on landscape**

When an infectious deer is harvested ($Z_{H_{[j]}}$), its carcass will either be left on the landscape, safely disposed of within the region, or exported out of a region safely (Eq. S31 line 2). Once harvested, carcasses are assumed to be processed immediately -within a time step- based on a relatively short field dressing time ($t_H$; Table S10). This processing time is very short compared to the time scales we are interested in, thus carcasses are sorted into one of three outcome stocks during each timestep. A constant proportion of the processed carcasses is transported outside of the region ($\varsigma_{T_{[j]}}$; Table S10), while another proportion is safely disposed of within the region ($\varsigma_{\iota_{[j]}}$; Table S10). The safe disposal rate only applies to carcasses not already transported. Any remaining carcasses are considered left on the landscape, which includes a constant fraction of all harvested deer representing gut piles left behind during field dressing ($\mho$; Table S10).

The final fate of all carcasses—whether from harvest or natural mortality—is governed by the same decay and collection dynamics. We track both healthy and infected carcasses, because the proportions of each affect the workload for carcass management policies. Approximately 15% of annual mortalities are attributed to non-harvest mortalities such as harvest wounding, poaching, predation, or vehicular collisions (Wisconsin Department of Natural Resources, personal communication). Of this, 7% is attributed to harvest wounding (refer to *Human caused mortality*) and is accounted for in our harvest mortality. The remaining 8% is attributed to other background mortality sources, including 6% from vehicle collisions (Fuller, 1990; Jennings et al., 2014). Since collection of road-killed carcasses is often impractical due to location and accessibility, we assume a maximum of 75% of non-harvest carcasses could be recovered through management action.

Harvested infectious carcasses ($Z_{H_{[j]}}$) are tracked in a similar manner (Eq. S32). The number of infectious deer harvested is (Eq. S32 line 1) the total number of the infectious deer within the last three stages of the infected stock ($y$=4,5,6) multiplied by the infected harvest rates, summed with the total number of clinical deer multiplied by their harvest rate (refer to *Human caused mortality*). The harvested infectious deer carcasses are then either exported to another region (Eq. S32 line 2), left on the landscape (Eq. S32 line 3), or safely disposed of (Eq. S32 line 4) according to the same structures and parameters described above for these processes.

$$\frac{dZ_{H[j]}}{dt} = \overbrace{[\textstyle\sum_{k=1}^{n}\sum_{l=1}^{n}\sum_{y=4}^{n} h_{I[j,k,l]} I_{[j,k,l,y]} + \sum_{k=1}^{n}\sum_{l=1}^{n} h_{C[j,k,l]} C_{[j,k,l]}]}^{\textit{Infectious deer harvest}}$$

$$-\overbrace{[\frac{Z_{H[j]}}{t_H} \varsigma_{T[j]}]}^{\textit{Carcass transport}}$$

$$\overbrace{-[\underbrace{\frac{Z_{H[j]}}{t_H}}_{\textit{Carcass handling}} - \underbrace{\frac{Z_{H[j]}}{t_H}\varsigma_{T[j]}}_{\textit{Carcass transport}} - \underbrace{\frac{Z_{H[j]}}{t_H}\varsigma_{\iota[j]}\left(1-\varsigma_{T[j]}\right)}_{\textit{Safe carcass disposal}} + \underbrace{\text{Ʊ}(\textstyle\sum_{k=1}^{n}\sum_{l=1}^{n} h_{S[j,k,l]} S_{[j,k,l]} + \sum_{k=1}^{n}\sum_{l=1}^{n}\sum_{y=4}^{n} h_{I[j,k,l]} I_{[j,k,l,y]})}_{\textit{Carcass left on landscape after field dressing}})]}^{\textit{Harvested infectious deer left on the landscape}}$$

$$-\overbrace{[\frac{Z_{H[j]}}{t_H}\varsigma_{\iota[j]}\left(1-\varsigma_{T[j]}\right)]}^{\textit{Safe carcass disposal}}$$

**Equation S32. Change in total infectious carcasses harvested**

**Table S10. Mapping between the full variable names used to construct the system dynamics model in Vensim (*Full Variable Name*) and the abbreviated forms (*Short Form*) used in the manuscript to represent the accumulation of environmental chronic wasting disease prions in the system dynamics model.** The table includes each variable's units, the corresponding equation used to calculate it, or the source from which its value was derived.

| Short Form | Full Variable Name | Units | Equation/Source |
|---|---|---|---|
| $V_{[j]}$ | Environmental Prions[Region] | prions | Equation S1.4 |
| $Z_{F_{[j]}}$ | Carcasses on Landscape[Region,type] | deer | Equation S31 *type*= c infected |
| $\varepsilon_F$ | Prions per deer year | prions/deer/year | 1 |
| $\varepsilon_Z$ | Carcass Deer Year Equivalent | year | Estimated (Table S17) |
| $t_Z$ | carcass landscape lifetime | year | 0.25 |
| $\varsigma_\Psi$ | carcass fraction collected | fraction | **user defined value between 0 to 1** |
| $F_{[j]}$ | Infectious population | deer | Equation S29 |

| Short Form | Full Variable Name | Units | Equation/Source |
|---|---|---|---|
| $\tau$ | Prion Life | year | Equation S30 |
| $\tau_{hl}$ | Prion Half Life | year | Estimated (Table S17) |
| $Z_{H_{[j]}}$ | Hunter Carcasses[Region,type] | deer | Equation S32 *type*= c infected |
| $t_H$ | carcass processing time | year | 0.0625; Time step of the model |
| $\varsigma_{T_{[j]}}$ | carcass fraction transported[Region] | fraction | 0.2 |
| $\varsigma_{\iota_{[j]}}$ | carcass safe disposal fraction[Region] | fraction | $= \begin{cases} \varsigma_{init_{[j]}} & \Psi = 0 \\ \varsigma_g & \Psi = 1 \end{cases}$ |
| $\mho$ | gut pile fraction | fraction | 0.8 |
| $\varsigma_U$ | other mortality collected | fraction | **user defined value between 0 to 0.75** |
| $\varsigma_{init_{[j]}}$ | initial carcass safe disposal fraction[Region] | fraction | 0.2 |
| $\varsigma_g$ | future carcass safe disposal fraction | fraction | **user defined value between 0 to 1** |

##### 2.2.4 Social dynamics

Our model extends beyond biological processes to endogenously incorporate key social and institutional processes that influence the chronic wasting disease (CWD) system. One such process is disease surveillance, a key factor as CWD testing regimes shape stakeholder perceptions of disease distribution and prevalence. To capture this, the model treats testing as an endogenous and (optionally) stochastic process, which determines the reported prevalence and spatial spread of CWD based on both the underlying true disease dynamics and the testing regime implemented. Inclusion of stochastic effects in testing is important because the variance in forecasts depends on the aleatory uncertainty of the measurement process as well as the epistemic uncertainty of the underlying system dynamics.

Targeted management interventions such as agency removals—intensive culling within known disease hotspots—are also incorporated. Agency removals preferentially target sick and infected deer and is implemented in the model through infection state– and cohort-specific harvest rates, which can be based on historical values or set by the user. The effectiveness of agency removals depends on both the spatial extent of the targeted area and the local disease prevalence. Together, these factors shape the ability of agency personnel to encounter and remove infected individuals, influencing differential removal outcomes across age and sex classes.

Recreational deer harvest by hunters is modeled as a function of both hunter license purchasing and hunter behavior—in terms of mobilization to the field and the per-hunter harvest rates of antlered and antlerless deer. The number of hunters that purchase a license is endogenously generated within the model and calibrated using historical license sales data (refer to Hunter submodule). The attractiveness of hunting is modeled endogenously, driven by system attributes such as perceived deer abundance and the presence of mature bucks, which influence hunter effort and thus harvest pressure over time. Here we only allow the hunting attractiveness to modify license purchasing behavior, not hunter mobilization (i.e., proportion of those that purchase that go afield), or the per-hunter harvest rates.

##### *2.2.4.1 Testing*

Historic surveillance data do not distinguish between four- and five-year-old deer. To align the model with available data, we grouped these age classes together. As such, model age classes used for surveillance-based calculations are: (1) fawns [0–1 years], (2) yearlings [1–2 years], (3) two-year-olds [2–3 years], (4) three-year-olds [3–4 years], (5) four- and five-year-olds [4–6 years], and (6) six years and older [6+ years] (denoted $k^*$). This grouping is used to calculate both the predicted observed fraction of positive tests and the number of samples per year (refer to *Data and processing*).

Sampling effort directly influences the probability of detecting CWD in a region. Increased sampling increases the likelihood of early detection when prevalence is low, enabling more timely management responses. By incorporating historical sampling dynamics, the model estimates the discrepancy between observed (reported) prevalence and true, underlying prevalence. We define the time to first detection of CWD in a region as the difference between the time of the first infected individual's appearance and the first positive test result. The reported prevalence, based on regional testing, is updated annually in the model. This reflects the concentrated burst of test information available during the harvest and the slower pace of public information dissemination relative to the more immediate access available to agency personnel interacting with surveillance data.

Testing is modeled as a stochastic process. Recent positive tests ($R_{P_{[j]}}$) are accumulated for reporting and are removed one year after entry (Eq. S33.1). The flow of positive tests ($\Gamma_{P_{[j,k^*,l]}}$) at each time ($t$) includes contributions from carcasses originating from harvest and natural mortality (collectively termed "observed mortalities") as well as from agency removals. Each test outcome is a draw from a binomial distribution (Eq. S33.2), with the number of trials equal to the number of sampled carcasses ($\Gamma_{N_{[j,k^*,l]}}$ and $\Gamma_{M_{[j,k,l]}}$), and the probability of success equal to the proportion of those that are in a detectable stage of infection ($f_{P_{[j,k^*,l]}}$ and $f_{Pm_{[j,k,l]}}$; Table S11). The structure accounts for the sampling error around the surveillance data used to constrain the model during estimation.

**Equation S33.**

$$\frac{dR_{P_{[j]}}}{dt} = \sum_{k=1}^{n} \sum_{l=1}^{n} (\Gamma_{P_{[j,k,l,t]}} - \Gamma_{P_{[j,k,l,t-1]}})$$ **Equation S33.1 Change in recent CWD positive deer carcasses tested**

$$\Gamma_{P_{[j,k,l]}} = Bin\left(\Gamma_{N_{[j,k^*,l]}}, f_{P_{[j,k^*,l]}}\right) + Bin\left(\Gamma_{M_{[j,k,l]}}, f_{Pm_{[j,k,l]}}\right)$$ **S33.2 CWD positive deer carcasses**

The number of deer tested through harvest surveillance ($\Gamma_{N_{[j,k^*,l]}}$) depends on whether management interventions are in place (Eq. S34.1). If not ($\psi$=0), historic sampling targets ($\Pi_{[j,k^*,l]}$) are used. If interventions are active ($\psi$=1), sampling is determined as the product of the observed mortalities ($U_{[j,k,l]}$) and a user-defined regional sampling rate ($\Pi_{\Gamma_{[j]}}$). The total number of sharpshot deer ($\Gamma_{M_{[j,k,l]}}$; Eq. S34.2) is the number of susceptible and infected individuals culled, calculated as the product of each stock weighted by their respective agency removals rates ($m_{S[j,k,l]}$ and $m_{I[j,k,l]}$).

**Equation 34.**

$$\Gamma_{N_{[j,k^*,l]}} = \begin{cases} \Pi_{[j,k^*,l]} & \Psi = 0 \\ U_{[j,k,l]}\Pi_{\Gamma_{[j]}} & \Psi = 1 \end{cases}$$ **Equation S34.1 Target number of CWD test for harvest surveillance**

$$\Gamma_{M_{[j,k,l]}} = m_{S_{[j,k,l]}}S_{[j,k,l]} + m_{I_{[j,k,l]}}(\textstyle\sum_{y=1}^{n} I_{[j,k,l,y]} + C_{[j,k,l]})$$ **Equation S35.2 Total number of deer that are culled via agency removals**

The probability that a carcass will test positive depends on whether it is in a detectable stage of infection. For harvest and background mortality ($f_{P_{[j,k^*,l]}}$; Eq. S35.1), this is the proportion of observed carcasses in detectable stages ($\frac{U_{d_{[j,k,l]}}}{U_{[j,k,l]}}$). For agency removals ($f_{Pm_{[j,k,l]}}$; Eq. S35.2), this is the proportion of total culls in a detectable stage ($\frac{M_{d_{[j,k,l]}}}{\Gamma_{M_{[j,k,l]}}}$). If no deer were culled, the probability for agency removals is set to zero to avoid division by zero.

**Equation 35**.

$$f_{P_{[j,k^*,l]}} = \frac{U_{d_{[j,k,l]}}}{U_{[j,k,l]}}$$ **Equation S35.1 The fraction of observed mortalities that can test positive for CWD**

$$f_{Pm_{[j,k,l]}} = \begin{cases} \frac{M_{d_{[j,k,l]}}}{\Gamma_{M_{[j,k,l]}}} & \Gamma_{M_{[j,k,l]}} \neq 0 \\ 0 & \Gamma_{M_{[j,k,l]}} = 0 \end{cases}$$ **Equation S35.2 The fraction of agency removals culls that can test positive for CWD**

The total number of observed mortalities ($U_{[j,k,l]}$) includes all harvested deer and a fraction ($f_{\Gamma_\mu}$) of background mortalities that are collected (e.g., roadkill) and available for testing (Eq. S36). The number in a detectable stage ($U_{d_{[j,k,l]}}$) is calculated by removing deer that have not progressed far enough in the disease to be detected. This includes all clinical animals and a fraction of infected individuals in earlier stages, weighted by stage-specific detectability ($d_{[y]}$; Eq. S37.1). Similarly, detectable agency removals culls ($M_{d_{[j,k,l]}}$) are estimated based on clinical cases and stage-specific detectability (Eq. S37.2).

$$U_{[j,k,l]} = \left[h_{S_{[j,k,l]}}S_{[j,k,l]} + \textstyle\sum_{y=1}^{n} h_{I_{[j,k,l]}}I_{[j,k,l,y]} + h_{C_{[j,k,l]}}C_{[j,k,l]}\right] + f_{\Gamma_\mu}\left[\mu_{S_{[j,k]}}S_{[j,k,l]} + \textstyle\sum_{y=1}^{n}(\mu_{S_{[j,k]}} + \psi_I)I_{[j,k,l,y]} + (\mu_C + \psi_C)C_{[j,k,l]}\right]$$ **Equation S36. The total number of deer carcasses available for CWD surveillance testing**

**Equation 37.**

$$U_{d_{[j,k,l]}} = h_{C_{[j,k,l]}}C_{[j,k,l]} + f_{\Gamma_\mu}(\mu_C + \psi_C)C_{[j,k,l]} + d_{[y]}\left(\textstyle\sum_{y=1}^{n} h_{I_{[j,k,l,y]}}I_{[j,k,l,y]} + \textstyle\sum_{y=1}^{n} f_{\Gamma_\mu}(\mu_{S_{[j,k]}} + \psi_I)I_{[j,k,l,y]}\right)$$ **Equation S37.1 The total number of deer carcasses in a detectable infection stage and available for CWD surveillance testing**

$$M_{d_{[j,k,l]}} = m_{I_{[j,k,l]}}C_{[j,k,l]} + d_{[y]}\left(\textstyle\sum_{y=1}^{n} m_{I_{[j,k,l]}}I_{[j,k,l,y]}\right)$$ **Equation S37.2 The total number of agency removals culls in a detectable infection stage**

We assume that deer in the clinical stage always test positive. Detectability for infected individuals (Eq. S38) increases with disease progression (Sigurdson et al., 1999), approaching the test sensitivity limit ($\vartheta_d$; Table S11). Detectability for each infection stage is determined by the ratio of the reference time since exposure ($t_{refE_{[y]}}$) to the detection time threshold ($t_d$), with a maximum of one (Eq. S38). The reference time (Table S11) is initialized at model setup prior to any future genetic shifts (refer to *Disease progression*).

$d_{[y]} = Min(1, \frac{t_{refE_{[y]}}}{t_d})\vartheta_d$ **Equation S38. The ability of testing to detect CWD within the different sub-stages of the Infected stock**

The number of infectious deer consumed by humans ($F_{consumed_{[j]}}$) depends on testing coverage and human behavior. We assume that a proportion of individuals knowingly consume infected meat ($f_{Pc}$), while others avoid it. Infectious deer in detectable stages that are not tested (or whose test results are ignored) contribute to potential human exposure. If no testing occurs, all harvested infectious deer are assumed to be consumed. If full testing occurs and all positive results are avoided, only undetectable infections contribute to human exposure (Eq. S39). Note that humans don't harvest or consume clinically sick deer ($h_{C_{[j,k,l]}} = 0$) but we include the structure for completeness.

**Equation S39. Estimate of the amount of infectious harvest that is consumed by humans.**

$$F_{consumed_{[j]}} = \underbrace{\sum_{k=1}^{n}\sum_{l=1}^{n}\left(h_{C_{[j,k,l]}}C_{[j,k,l]} + \left(\sum_{y=4}^{n} h_{I_{[j,k,l,y]}}I_{[j,k,l,y]}\right)\right)}_{\text{Annual infectious harvest}} - \Pi_{\Gamma_{[j]}}(1 - f_{Pc})\underbrace{\sum_{k=1}^{n}\sum_{l=1}^{n}\left(h_{C_{[j,k,l]}}C_{[j,k,l]} + \sum_{y=4}^{n} d_{[y]}\left(h_{I_{[j,k,l,y]}}I_{[j,k,l,y]}\right)\right)}_{\text{Annual detectable infectious harvest}}$$

**Table S11. Mapping between the full variable names used to construct the system dynamics model in Vensim (*Full Variable Name*) and the abbreviated forms (*Short Form*) used in the manuscript to represent the surveillance testing regime of chronic wasting disease in the system dynamics model.** The table includes each variable's units, the corresponding equation used to calculate it, or the source from which its value was derived.

| Short Form | Full Variable Name | Units | Equation/Source |
|---|---|---|---|
| $R_{P_{[j]}}$ | YTD Positive[Region] | deer | Equation S33.1 |
| $\Gamma_{P_{[j,k^*,l]}}$ | Positive Sampling[Region,age data,sex] | deer/year | Equation S33.2 |
| $\Gamma_{N_{[j,k^*,l]}}$ | Sampling[Region,age data,sex] | deer/year | Equation S34.1 |
| $\Gamma_{M_{[j,k,l]}}$ | Agency removals Sampling[Region,age,sex] | deer/year | Equation S34.2 |
| $\Pi_{[j,k^*,l]}$ | data surv N samples[Region,age data,sex] | deer/year | Data (refer to *Data and processing* below) |
| $\Gamma_{g_{[j]}}$ | Target sampling[Region] | deer/year | **user defined value between 20 to 500** |
| $\Pi_{\Gamma_{[j]}}$ | Target sampling rate[Region] | fraction | $\frac{\Gamma_{g_{[j]}}}{\sum_{k=1}^{n}\sum_{l=1}^{n} U_{[j,k,l]}}$ |
| $f_{P_{[j,k^*,l]}}$ | observed fraction positive Age Data[Region,age data,sex] | fraction | Equation 35.1 |
| $f_{Pm_{[j,k,l]}}$ | agency removals frac positive[Region,age,sex] | fraction | Equation 35.2 |
| $U_{[j,k,l]}$ | observed mortality[Region,age,sex] | deer | Equation S36 |
| $f_{\Gamma_\mu}$ | fraction non-harvest mortality captured in surveillance | fraction | 0.1 (Fuller, 1990; Jennings et al., 2014; Wallingford et al., 2017) |
| $U_{d_{[j,k,l]}}$ | observed detectable infected mortality[Region,age,sex] | deer | Equation 37.1 |
| $M_{d_{[j,k,l]}}$ | Agency removals Detectable[Region,age,sex] | deer/year | Equation 37.2 |
| $d_{[y]}$ | stage detectability[stage] | fraction | Equation 38 |
| $t_d$ | detection threshold time | year | 1 (Haley et al., 2012) |

| Short Form | Full Variable Name | Units | Equation/Source |
| --- | --- | --- | --- |
| $t_{refE_{[y]}}$ | ref stage time from exposure[stage] | year | $= \begin{cases} \frac{1}{\rho_{ref_{[y]}}} & y = 1 \\ \frac{(t_{refE_{[y=1]}}+1)}{\rho_{ref_{[y]}}} & y \neq 1 \end{cases}$ |
| $\vartheta_d$ | max sensitivity | fraction | 0.96 (Hibler et al., 2003; Viljugrein et al., 2021) |
| $f_{Pc}$ | fraction positive consumed | fraction | 0.8 (Bradshaw et al., 2021) or user dictated value between 0-1 |

##### *2.2.4.2 Agency removals*

Agency-directed deer removals are modeled as a discrete, annual intervention occurring in a single pulse each year ($t_{step}$= 3), allowing its demographic and epidemiological impacts to be accurately captured. The number of deer removed from each stock due to agency removals was calculated as the product of the cohort-specific agency removal rate and the number of individuals in that stock (Eq. S1.1, S1.2, S1.3). Agency removal rates varied by infection status and were defined separately for susceptible ($m_{S_{[j,k,l]}}$) and infected/clinical ($m_{I_{[j,k,l]}}$) deer in each region, age, and sex class (Eq. S40.1 and S40.2, respectively). Here, the targeted agency removal rates for susceptible ($m_{gS_{[j]}}$) and infected deer ($m_{gI_{[j]}}$) were modulated by a cohort-specific enrichment factor ($\epsilon_{\Im_{[j,k,l]}}$), which represents the targeting bias toward certain age and sex classes based on observed infection patterns and agency personnel ability. The primary mechanism for this enrichment is spatial, i.e. agency personnel focus on areas within a county known to have higher-than-average prevalence.

**Equation S40**

$$m_{S_{[j,k,l]}} = m_{gS_{[j]}} \epsilon_{\Im_{[j,k,l]}}$$ **S40.1 Agency removal rate of susceptible individuals**

$$m_{I_{[j,k,l]}} = m_{gI_{[j]}} \epsilon_{\Im_{[j,k,l]}}$$ **S40.2 Agency removal rate of infected and clinical individuals**

The cohort specific enrichment (Eq. S41) is modeled as an exponential function of the relative cohort-specific prevalence ($\phi_{\zeta_{[j,k,l]}}$) and a user-defined cohort discernment parameter ($\Theta_{\Im_{[j]}}$), which describes the ability of agency personnel to distinguish among cohorts in the field (Table S12). The relative cohort-specific prevalence (Eq. S42) is calculated as the ratio of the true infection prevalence within a given cohort ($\zeta_{\Im_{[j,k,l]}}$) to the overall regional prevalence ($\zeta_{[j]}$), with a value of zero assigned where infection is absent.

$$\epsilon_{\Im_{[j,k,l]}} = \phi_{\zeta_{[j,k,l]}}{}^{\Theta_{\Im_{[j]}}}$$ **S41. Cohort specific enrichment of deer agency removals**

$$\phi_{\zeta_{[j,k,l]}} = \begin{cases} \frac{\zeta_{\Im_{[j,k,l]}}}{\zeta_{[j]}} & \zeta_{\Im_{[j,k,l]}} \neq 0 \\ 0 & \zeta_{\Im_{[j,k,l]}} = 0 \end{cases}$$ **S42. Relative cohort-specific prevalence**

The regional targeted agency removal rates ($m_{gS[j]}$, $m_{gI[j]}$) are dynamically allocated between susceptible and infected individuals based on the total desired agency removal rate ($m_{[j]}$), the infection discernment ability of agency personnel($\Theta_{I_{[j]}}$), the regional prevalence ($\zeta_{[j]}$), and the spatial aggregation of infected deer near the outbreak epicenter ($\zeta_{P_{[j]}}$) as follows (Eq. S43):

**Equation S43**

$m_{gS[j]} = m_{[j]}(\frac{(1-\left(\Theta_{I_{[j]}}\zeta_{P_{[j]}}+\left(1-\Theta_{I_{[j]}}\zeta_{[j]}\right)\right))}{(1-\zeta_{[j]})})$ **S43.1. Targeted agency removal rate of susceptible individuals**

$m_{gI[j]} = m_{[j]}(\frac{\left(\Theta_{I_{[j]}}\zeta_{P_{[j]}}+\left(1-\Theta_{I_{[j]}}\zeta_{[j]}\right)\right)}{\zeta_{[j]}})$ **S43.2. Targeted agency removal rate of infected and clinical individuals**

Ability to identify infected deer is very limited during the early stages of disease but infections become more recognizable as deer progress through the stages of the disease such that clinical deer are readily recognized. Agency personnel infection discernment ($\Theta_{I_{[j]}}$) is modeled as a user-controlled parameter ranging from 0 (no ability to distinguish infection) to 1 (perfect recognition), with a baseline of 0.5 (Table S12). These discernment parameters (cohort, infection) are intended to reflect management tradeoffs—e.g., if investing in training improves targeting precision, does it translate to a significant improvement in agency removals results. The total desired agency removal rate ($m_{[j]}$) was based either on historical patterns ($m_{m_{[j]}}$) or a user-defined management scenario ($m_{g_{[j]}}$), depending on whether active management intervention was triggered during the simulation (Eq. S44).

$m_{[j]} = \begin{cases} m_{m_{[j]}} & \Psi = 0 \\ m_{g_{[j]}} & \Psi = 1 \end{cases}$ **S44. Management desired rate of agency removals**

To further account for spatial heterogeneity in infection, we calculate the relative clumping of infected deer ($\zeta_{P_{[j]}}$), defined as a function of maximum prevalence ($\zeta_{max_{[j]}}$), true prevalence ($\zeta_{[j]}$), and the fraction of the region affected by CWD ($f_{A_{I_{[j]}}}$) and under agency removals ($f_{A_{m_{[j]}}}$). This is informed by observation of the relationship between prevalence and spatial extent at the township level within Wisconsin counties. The effect should be regarded as a rough approximation, because one would expect the spatial distribution of infected deer to change in response to extensive application of sharpshooting.

This is calculated as the product of two terms (Eq. S45) where the first results in the maximum prevalence or the true prevalence when the fraction of the area that is infected relative to the agency removals area is small or large respectively. The second term captures the ratio of areas with infected deer to the area where personnel are looking for infected deer, representing both the increasing challenge of locating infected deer as CWD spreads and the decreasing spatial precision of management as effort intensifies. If the maximum between the area infected and the agency removals area is zero, then the second term in the full equation becomes zero to avoid division by zero (Eq. S45).

**Equation S45. The relative clumping of infected deer at outbreak epicenter**

$$\zeta_{P_{[j]}} = \frac{\left(2\zeta_{max_{[j]}}-\left(\zeta_{max_{[j]}}-\zeta_{min_{[j]}}\right)Min(f_{A_{I_{[j]}}},f_{A_{m_{[j]}}})\right)}{2} \times \frac{f_{A_{I_{[j]}}}}{Max(f_{A_{I_{[j]}}},f_{A_{m_{[j]}}})}$$

Maximum prevalence (Eq. S46.1) was constrained as the minimum of either: (1) the greater of the saturation prevalence ($\zeta_{sat}$) or the true prevalence ($\zeta_{[j]}$), and (2) a scaling function ($\xi_{[j]}$) representing potential localized concentration. Taking the maximum between these two terms captures the idea that, if the true prevalence in a region is less than the saturation prevalence, then no spatial distribution of deer will create a greater prevalence within the region. The scaling function is calculated as double the true prevalence divided by the fraction of the area that is affected by CWD within the region and set to zero if the area is unaffected (Eq. S46.2). The saturation prevalence establishes an upper bound on the possible CWD prevalence in a population which is limited by disease and host dynamics. The minimum prevalence of CWD in the deer population within a region is given by the true prevalence and the maximum prevalence that can be found across the landscape within a region (Eq. S46.3).

**Equation S46**

$\zeta_{max_{[j]}} = Min(Max(\zeta_{sat}, \zeta_{[j]}), \xi_{[j]})$ **S46.1 The maximum prevalence possible within a region**

$$\xi_{[j]} = \begin{cases} \frac{2\zeta_{[j]}}{f_{A_{I_{[j]}}}} & f_{A_{I_{[j]}}} \neq 0 \\ 2\zeta_{[j]} & f_{A_{I_{[j]}}} = 0 \end{cases}$$ **S46.2 The potential localized concentration of infected deer**

$$\zeta_{min_{[j]}} = \begin{cases} 2\zeta_{[j]} - \zeta_{max_{[j]}} & \zeta_{max_{[j]}} < 2\zeta_{[j]} \\ 0 & \zeta_{max_{[j]}} > 2\zeta_{[j]} \end{cases}$$ **S46.3 The minimum prevalence possible within a region**

The fraction of a region affected by CWD ($f_{A_{I_{[j]}}}$) was modeled as an exponential function of true prevalence (Eq. S47), calibrated using empirical township-level data from Wisconsin (refer to *Data and processing*).

$$f_{A_{I_{[r]}}} = 1 - e^{(\frac{-\zeta_{[j]}}{0.07})}$$ **S47. The fraction of area in a region that is affected by chronic wasting disease**

Similarly, the fraction of the region where agency removals is conducted ($f_{A_{m_{[j]}}}$) scaled with the management agency removals rate (Eq. S46), reflecting effort diffusion over space.

$$f_{A_{m_{[j]}}} = 1 - e^{(\frac{-m_{[j]}}{m_{max}})}$$ **S48. The fraction of area in a region where agency removals are occurring**

The result of these model structures is that at the start of an epidemic (i.e., low true prevalence) the infected deer are much more likely to be clumped at the epicenter, allowing for a higher targeted agency removals rate for infected deer compared to susceptible ones (Eq. S43). Including the impact of disease diffusion allows our model to capture the change in the effectiveness of agency removals given the stage of the epidemic.

**Table S12. Mapping between the full variable names used to construct the system dynamics model in Vensim (*Full Variable Name*) and the abbreviated forms (*Short Form*) used in the manuscript to represent the impacts of agency mandated agency removals on the deer and disease dynamics in the system dynamics model.** The table includes each variable's units, the corresponding equation used to calculate it, or the source from which its value was derived.

| Short Form | Full Variable Name | Units | Equation/Source |
|---|---|---|---|
| $m_{S_{[j,k,l]}}$ | Agency removals rate[Region,age,sex,c infected] | fraction/year | Equation S40.1 |
| $m_{I_{[j,k,l]}}$ | Agency removals rate[Region,age,sex,c non infected] | fraction/year | Equation S40.2 |
| $\epsilon_{\Im_{[j,k,l]}}$ | age sex enrichment[Region,age,sex] | dimensionless | Equation S41 |
| $\Theta_{\Im_{[j]}}$ | age sex targeting sensitivity[Region] | dimensionless | **user defined value between 0-1** |
| $\phi_{\zeta_{[j,k,l]}}$ | relative prevalence[Region,age,sex] | dimensionless | Equation S42 |
| $\zeta_{\Im_{[j,k,l]}}$ | prevalence by age sex[Region,age,sex] | dimensionless | $\frac{(\sum_{y=1}^{n} I_{[j,k,l,u]}+C_{[j,k,l,y]})}{(S_{[j,k,l]}+\sum_{y=1}^{n} I_{[j,k,l,y]}+C_{[j,k,l]})}$ |
| $m_{gS[j,k,l]}$ | targeted agency removals rate[Region, c non infected] | fraction/year | Equation S43.1 |
| $m_{gI[j,k,l]}$ | targeted agency removals rate[Region, c infected] | fraction/year | Equation S43.2 |
| $m_{[j]}$ | desired agency removals rate[Region] | fraction/year | Equation S44 |
| $m_{g[j]}$ | future agency removals rate[Region] | fraction/year | **user defined value between 0-0.1** |
| $m_{m[j]}$ | historic agency removals rate[Region] | fraction/year | 0 |
| $\Theta_{I_{[j]}}$ | target correlation[Region] | dimensionless | 0.5 |
| $\zeta_{P_{[j]}}$ | perfect targeted prevalence[Region] | fraction | Equation S45 |
| $\zeta_{max_{[j]}}$ | max prevalence[Region] | fraction | Equation S46.1 |
| $\zeta_{min_{[j]}}$ | min prevalence[Region] | fraction | Equation S46.3 |
| $\zeta_{sat}$ | saturation prevalence | fraction | 0.6 |
| $\xi_{[j]}$ | *within the equation for 'max prevalence[Region]' | fraction | Equation S46.2 |
| $f_{A_{I_{[j]}}}$ | fraction area affected[Region] | fraction | Equation S47 |
| $m_{max[j]}$ | max SS treatment rate | fraction/year | 0.5 |
| $f_{A_{m_{[j]}}}$ | agency removals target area[Region] | fraction | Equation S48 |

##### *2.2.4.3 Hunter participation*

The hunter participation sector determines the propensity of hunters to seek a license in the coming year. We tracked a population of deer hunters for each region ($j$), categorizing individuals into one of two stocks: *Active hunters* and *Inactive hunters* (Eq. S49). All movements between hunter stocks occur in a discrete pulse ($t_{step}$=14), just prior to the harvest pulse. Active hunters are defined as those who have purchased one or more deer hunting licenses in the current year (Eq. S49.1), whereas inactive hunters are those who previously held a license but did not purchase one during the current year (Eq. S49.2). Each year, the number of active hunters ($H_{A_{[j]}}$) may increase through recruitment from a pool of potential hunters ($H_{P_{[j]}}$) with a regional recruitment probability ($\varpi_{c_{[j]}}$), or through reactivation of individuals from the inactive stock ($H_{In_{[j,v]}}$) at a regional stage-specific reactivation probability ($\varpi_{a_{[j,v]}}$). Active hunters who do not purchase a license transition into the first stage of inactivity ($H_{In_{[j,1]}}$) at a probability equal to one minus the regional probability of retention ($\varpi_{t_{[j]}}$). Mortality also reduces the number of active hunters at a region-specific rate ($\mu_{A_{[j]}}$).

To enable hierarchical parameter estimation across regions, we defined each of the R3 (Recruitment-Retention-Reactivation) transition probabilities as the product of a global base rate

($\varpi_c$, $\varpi_t$, $\varpi_{a_{[j,v]}}$) and a region-specific modifier ($\phi_{\varpi_{c[j]}}$, $\phi_{\varpi_{l[j]}}$, $\phi_{\varpi_{a[j]}}$; Table S13). Given that retention probabilities tend to exceed 0.8, we applied regional modifiers to the *loss* value (i.e., 1-$\varpi_t$) to avoid parameter estimates approaching or exceeding 1 during optimization, thereby improving model stability during estimation. These R3 transitions can be further modified by the perceived attractiveness of hunting in the region ($Å_{[j]}$) given, for example, the abundance of mature bucks, which influences an individual's decision to purchase a hunting license.

To better reflect declining reactivation probability over time, we disaggregated the inactive hunter stock into five sequential annual stages of inactivity (*v*; Eq. S49.2), following Hinrichs et al. (2020). Individuals who fail to renew their license enter the first year of inactivity ($H_{In_{[j,1]}}$; Eq. S49.2a), and in subsequent years, non-reactivated individuals advance one stage at each time step (Eq. S49.2b), up to stage five. The final stage aggregates all individuals inactive for five or more years, after which reactivation is considered highly unlikely (Table S13). Inactive hunters are also subject to mortality, with rates specific to both region and stage ($\mu_{In_{[j,v]}}$).

**Equation S49**

**S49.1 Change in number of active hunters**

$$\frac{dH_{A_{[j]}}}{dt} = \overbrace{\varpi_{c_{[j]}} Å_{[j]} H_{P_{[j]}}}^{\textit{Recruitment}} + \overbrace{\textstyle\sum_{v=1}^{n} \varpi_{a_{[j,v]}} Å_{[j]} H_{In_{[j,v]}}}^{\textit{Reactivation}} - \overbrace{\left(1 - \varpi_{t_{[j]}} Å_{[j]}\right) H_{A_{[j]}}}^{\textit{Active loss}} - \overbrace{\mu_{A_{[j]}} H_{A_{[j]}}}^{\textit{Active mortality}}$$

**S49.2a Change in number of inactive hunters in the first inactive stage**

$$\frac{dH_{In_{[j,v]}}}{dt} = \overbrace{\left(1 - \varpi_{t_{[j,v]}} Å_{[j]}\right) H_{A_{[j,v]}}}^{\textit{Active loss}} - \overbrace{\varpi_{a_{[j,v]}} Å_{[j]} H_{In_{[j,v]}}}^{\textit{Reactivation}} - \overbrace{\mu_{In[j,v]} H_{In_{[j,v]}}}^{\textit{Inactive mortality}} \quad v = 1$$

**S49.2b Change in number of inactive hunters in all inactive stages beyond the first**

$$\frac{dH_{In_{[j,v]}}}{dt} = \overbrace{H_{A_{[j,v-1]}}}^{\textit{Active loss progression}} - \overbrace{\varpi_{a_{[j,v]}} Å_{[r]} H_{In_{[j,v]}}}^{\textit{Reactivation}} - \overbrace{\mu_{In[j,v]} H_{In_{[j,v]}}}^{\textit{Inactive mortality}} \quad v > 1$$

Active hunters are recruited from a pool of *potential hunters* ($H_{P[j]}$), which includes all residents in a county who may consider hunting but have never purchased a deer license. We estimate this pool by multiplying the county's total population ($P_{[j]}$) by the proportion of individuals who would ever consider hunting, based on their wildlife value orientation ($\Lambda_{[j]}$) given the trend of social values towards wildlife and hunting (Manfredo et al., 2021b). We then subtract all individuals already active or previously recruited (i.e., in inactive stages) to estimate the remaining recruitable pool (Eq. S50). The processes of births, deaths, and aging are implicitly accounted for within the census data dictating county populations, thus, we simply update the maximum potential hunters within a region each year given the number of active and inactive hunters that year and the trend in wildlife value orientation.

**Equation S50. The maximum number of potential new hunters available for recruitment each year**

$$H_{P_{[j]}} = \left(\Lambda_{[j]}P_{[j]} - H_{A_{[j]}} - \sum_{v=1}^{n} H_{In_{[j,v]}}\right)$$

We accounted for shifts in societal attitudes toward hunting by allowing the county-specific wildlife value orientation (WVO) to change over time (Eq. S51). We initialized this value using the statewide estimate for Wisconsin in 2018 ($\Lambda_{WI}$), and applied a linear decline based on survey data from multiple states showing a downward trend in hunting interest when polled in 2004 and again in 2017-2018 (Manfredo et al., 2021b). A region-specific modifier ($\phi_{\Lambda_{[j]}}$) was applied to the statewide baseline, and the overall value was updated annually by a fixed slope ($t_{\Lambda}$) given the simulation year ($t$).

**Equation S51. The county specific wildlife value orientation over time**

$$\Lambda_{[j]} = \left(\Lambda_{WI}\phi_{\Lambda_{[j]}} - \Lambda_{T}(t - t_{\Lambda})\right)$$

**Table S13. Mapping between the full variable names used to construct the system dynamics model in Vensim (*Full Variable Name*) and the abbreviated forms (*Short Form*) used in the manuscript to represent the active hunter numbers.** The table includes each variable's units, the corresponding equation used to calculate it, or the source from which its value was derived.

| Short Form | Full Variable Name | Units | Equation/Source |
|---|---|---|---|
| $H_{A_{[j]}}$ | A Hunters[Region] | people | Equation S49.1 |
| $H_{In_{[j,v]}}$ | I Hunters[Region] | people | Equation S49.2 |
| $H_{P_{[j]}}$ | Pmax Hunters[Region] | people | Equation S50 |
| $\varpi_{c_{[j]}}$ | rec regional[Region] | people/year | $\varpi_{c}\phi_{\varpi_{c_{[j]}}}$ |
| $\varpi_{c}$ | recruitment | people/year | Estimated (Table S16) |
| $\phi_{\varpi_{c_{[j]}}}$ | Relative regional recruitment[Region] | dimensionless | Estimated (Table S16) |
| $\varpi_{t_{[j]}}$ | ret regional[Region] | people/year | $1-(1-\varpi_{t})\phi_{\varpi_{l_{[j]}}}$ |
| $\varpi_{t}$ | retention | people/year | Estimated (Table S16) |
| $\phi_{\varpi_{l_{[j]}}}$ | Relative regional loss[Region] | dimensionless | Estimated (Table S16) |
| $\varpi_{a_{[j,v]}}$ | react regional[Region, Inactive] | people/year | $\varpi_{a_{[v]}}\phi_{\varpi_{a_{[j]}}}$ |
| $\varpi_{a_{[v]}}$ | reactivation[Inactive] | people/year | Estimated (Table S16) |
| $\phi_{\varpi_{a_{[j]}}}$ | relative regional reactivation[Region] | dimensionless | Estimated (Table S16) |
| $\mu_{A_{[j]}}$ | A mort rate[Region] | people/year | Data (refer to *Data and processing* below) |
| $\mu_{In_{[j,v]}}$ | I mort rate[Region] | people/year | Data (refer to *Data and processing* below) |
| $P_{[j]}$ | County Pop[Region] | people | Data (refer to *Data and processing* below) |
| $\Lambda_{[j]}$ | regional WVO[Region] | dimensionless | Equation S51 |
| $\Lambda_{WI}$ | WI WVO | fraction | Estimated (Table S16) |
| $\phi_{\Lambda_{[j]}}$ | relative regional WVO[Region] | dimensionless | Estimated (Figure S) |
| $\Lambda_{T}$ | cultural shift in WVO | 1/year | Estimated (Table S16) |
| $t$ | Time | year | Model time variable |
| $t_{\Lambda}$ | WI WOV datapoint | year | 2018 |

##### 2.2.4.3.1 Hunter harvest

The number of hunters interacts with the core susceptible-infected-clinical (*SIC*) model of chronic wasting disease (CWD) in white-tailed deer through the number of deer harvested. To determine the number of active hunters afield in each region ($H_{F_{[j]}}$; Table S14), we calculate the product of the total number of active hunters that are residents in each region ($H_{A_{[j]}}$) and the hunter mobilization probability ($\Delta$). Not all hunters harvest deer in their county of residence. Thus, the number of hunters attempting to harvest deer in region $j$ ($H_{H_{[j]}}$; Eq. S52) is calculated by summing across all origin regions (i.e., county of residence), $j^*$, the product of the number of hunters afield from region $j^*$, and the proportion of those hunters that travel to region $j$ to hunt, as dictated by the hunter travel matrix ($\delta_{H_{[j^*,j]}}$).

**Equation S52. The number of hunters that are attempting to harvest a deer in each region.**

$$H_{H_{[j]}} = \sum_{j^*}^{n} \delta_{H_{[j^*,j]}} H_{F_{[j^*]}}$$

Once the number of hunters in each region is established, we estimate the number of deer harvested per hunter in two possible ways. First, when a harvest rate is given (refer to *Human caused mortality*), we calculate the number of deer that are required to be harvested per hunter to achieve that harvest rate ($R_{fawn_{[j,l]}}$, $R_{older_{[j,l]}}$, $R_{antlerless_{[j]}}$, $R_{antlered_{[j]}}$) by subtracting the number of deer taken through non-regular licenses ($h_{Ag_{[j,k,l]}}$; e.g., agency-directed harvest) from the total number of deer harvested, and dividing the result by the number of hunters afield in that region. Second, when the average harvest per hunter is specified directly, we estimate total regular harvest of each type ($E_{fawn_{[j,l]}}$, $E_{older_{[j,l]}}$, $E_{antlerless_{[j]}}$, $E_{antlered_{[j]}}$), by multiplying the rate by the number of hunters harvesting in each region. We present calculations for fawns and older deer, the estimates for antlerless and antlered harvest under each scenario are simply calculated at the sum of all does and fawns and all yearling and older bucks respectively.

These per-hunter harvest estimates provide an important social feasibility check when evaluating potential management scenarios. For instance, strategies that implicitly require hunters to double or triple their harvests may be impractical to implement, especially in the absence of strong incentives.

When harvest rates are dictated, the average number of fawns and of older deer taken per hunter are given by Equation S53.1 and S53.2 respectively:

**Equation S53. The number of deer that must be taken per hunter through regular licenses given a certain harvest rate**

$$R_{fawn_{[j,l]}} = \frac{\left(\sum_{k=1}^{1} h_{S_{[j,k,l]}} S_{[j,k,l]} + \sum_{k=1}^{1}\sum_{y=1}^{n} h_{I_{[j,k,l]}} I_{[j,k,l,y]} - \sum_{k=1}^{1} h_{Ag_{[j,k,l]}}\right)}{H_{H_{[j]}}}$$

**S53.1 The number of fawns**

$$R_{older_{[j,l]}} = \frac{\left(\sum_{k=2}^{n} h_{S_{[j,k,l]}} S_{[j,k,l]} + \sum_{k=2}^{n}\sum_{y=1}^{n} h_{I_{[j,k,l]}} I_{[j,k,l,y]} - \sum_{k=2}^{n} h_{Ag_{[j,k,l]}}\right)}{H_{H_{[j]}}}$$

**S53.2 The number of older deer**

Alternatively, if the per-hunter harvest rate is specified, the total number of deer harvested with regular licenses is computed by multiplying this rate by the number of hunters afield (Eq. S54). These rates are cohort specific with the historic values ($\mathrm{P}_{fawn_{[j,l]}}$, $\mathrm{P}_{older_{[j,l]}}$) estimated during hunter submodel fitting. These values may be modified by user-specified adjustment factors

($\Upsilon_{fawn_{[j,l]}}$, $\Upsilon_{older_{[j,l]}}$). When this factor is set to zero, the per-hunter harvest rate equals the model-estimated value. These per hunter harvest rates are averaged across all hunters that attempted to harvest a deer, including those who did not kill a deer.

**Equation S54. The estimated number of deer that are harvested through regular license harvest given the per hunter harvest and total number of hunters**

$E_{fawn_{[j,l]}} = (\mathrm{P}_{fawn_{[j,l]}} + \Upsilon_{fawn_{[j,l]}})H_{H_{[j]}}$ **S54.1 The total number of fawns**

$E_{older_{[j,l]}} = (\mathrm{P}_{older_{[j,l]}} + \Upsilon_{older_{[j,l]}})H_{H_{[j]}}$ **S54.2 The total number of older deer**

These estimates reflect only deer harvested through regular license opportunities and do not include deer that are wounded and unrecovered, nor those taken under non-standard permits such as nuisance or depredation licenses. To account for additional mortality due to wounding, all estimates of harvested deer are multiplied by 1.07, assuming a 7% wounding loss rate (refer to *Historic harvest rates*). These adjusted values are then summed with mortalities from non-standard permits ($h_{Ag_{[j,k,l]}}$ ) and divided by the total number of deer in the relevant cohort to derive the effective hunter harvest rate at harvest time that can be used in simulations.

**Table S14. Mapping between the full variable names used to construct the system dynamics model in Vensim (*Full Variable Name*) and the abbreviated forms (*Short Form*) used in the manuscript to represent the estimating of hunter harvest given hunter numbers and participation behaviors.** The table includes each variable's units, the corresponding equation used to calculate it, or the source from which its value was derived.

| Short Form | Full Variable Name | Units | Equation/Source |
|---|---|---|---|
| $\Delta$ | mobilization | dimensionless | 0.95 (Hunter survey report data) |
| $H_{F_{[j]}}$ | Hunters afield | people | $\Delta H_{A_{[j]}}$ |
| $H_{H_{[j]}}$ | Total hunters harvesting[Region] | people | $\sum_{j^*}^{n} \delta_{H_{[j^*,j]}} \Delta H_{A_{[j]}}$ |
| $\delta_{H_{[j^*,j]}}$ | Hunter harvest region[FromRegion, ToRegion] | dimensionless | *Active and inactive hunters* |
| $h_{Ag_{[j,k,l]}}$ | Data agriculture permit fawn harvest[Region,sex]<br>*k*=1 | deer/year | *Regular license harvest* |
| $h_{Ag_{[j,k,l]}}$ | Data agriculture permit older deer harvest[Region,sex]<br>*k*=2,3,4,5,6,7 | deer/year | *Regular license harvest* |
| $R_{fawn_{[j,l]}}$ | Required fawns harvested per hunter[Region,sex] | deer/people/year | Equation S53.1 |
| $R_{older_{[j,l]}}$ | Required older deer harvested per hunter[Region,sex] | deer/people/year | Equation S53.1 |
| $\mathrm{P}_{fawn_{[j,l]}}$ | Fawns harvested per hunter[sex] | deer/people/year | Estimated |
| $\mathrm{P}_{older_{[j,l]}}$ | Older deer harvested per hunter[Region,sex] | deer/people/year | Estimated |
| $\Upsilon_{fawn_{[j,l]}}$ | target change in fawn per hunter harvest[sex] | deer/people/year | User dictated value between 0 to 2 |
| $\Upsilon_{older_{[j,l=2]}}$ | target change in older doe per hunter harvest | deer/people/year | User dictated value between 0 to 2 |
| $\Upsilon_{older_{[j,l=1]}}$ | target change in older buck per hunter harvest | deer/people/year | User dictated value between 0 to 2 |
| $E_{fawn_{[j,l]}}$ | Predicted fawn harvested[Region, sex] | deer/people/year | Equation S54.1 |
| $E_{older_{[j,l]}}$ | Predicted older deer harvested[Region, sex] | deer/people/year | Equation S54.1 |

##### 2.2.4.3.2 Hunting attractiveness

The number of active hunters—and, all else equal, the number of deer harvested—can be dynamically adjusted through user-defined changes to the recruitment ($\varpi_{c_{[j]}}$), retention ($\varpi_{t_{[j]}}$), and reactivation ($\varpi_{a_{[j,v]}}$) probabilities. These rates are modulated by the perceived attractiveness of the hunting experience in each region ($Å_{[j]}$), which serves as a scalar on license purchasing behavior. When perceived attractiveness exceeds one, more licenses are purchased than expected based on historic rates; when it is below one, license purchasing declines. The dynamics of this perceived attractiveness are governed by the differential between current hunting attractiveness ($Å_{h_{[j]}}$) – scaled by the reference attractiveness ($Å_{ref_{[j]}}$) – and the perceived relative attractiveness region ($Å_{[j]}$), all scaled by a user-specified temporal delay parameter ($t_{Å}$ ) that captures the lag between changes in actual hunt conditions and hunter recognition of those changes (Eq. S55.1). This formulation introduces delay into hunter behavior, ensuring that responses to changes in the system do not occur instantaneously but unfold over time.

The reference attractiveness (Eq. S55.2) depends on whether hunter licensing intervention ($\psi_{R3}$) is active. If no intervention is occurring ($\psi_{R3} = 0$), the reference attractiveness equals the current attractiveness, thereby fixing $Å_{[j]}$ at one and removing its influence from hunter behavior (refer to Eq. S47). If an intervention is active ($\psi_{R3} =1$), the reference attractiveness is locked to the value at the onset of intervention, allowing perceived attractiveness to evolve relative to this benchmark. This structure ensures that the influence of perceived attractiveness on hunter recruitment, retention, and reactivation is explicitly controlled by the user.

To calculate the hunting attractiveness (Eq. S55.3), we adapted the framework of Sterman's logit choice model (Sterman, 2000), which posits that various attributes of a product—or in this case, a hunting experience—determine its market share or appeal. In our model, five factors contribute multiplicatively to the hunting attractiveness: expected hunt yield (i.e., the number of deer likely to be harvested given effort), cost of hunting, perceived CWD prevalence, perceived deer abundance, and perceived mature buck presence. We make the simplifying assumption that each R3 rate is impacted by all factors dictating attractiveness in the same manner.

**Equation S55**

$$\frac{dÅ_{[j]}}{dt} = \left(\frac{Å_{h_{[j]}}}{Å_{ref_{[j]}}} - Å_{[j]}\right) \Big/ t_{Å}$$ **S55.1 The change in the perceived hunting attractiveness**

$$Å_{ref_{[j]}} = \begin{cases} Å_{h_{[j]}} & \psi_{R3} = 0 \\ Å_{h_{[j]}} \, at\ time\ when\ \psi_{R3}\ became\ 1 & \psi_{R3} = 1 \end{cases}$$ **S55.2 The reference hunting attractiveness**

$$Å_{h_{[j]}} = e_{mb_{[j]}} e_{sight_{[j]}} e_{cost_{[j]}} e_{CWD_{[j]}} e_{yield_{[j]}}$$ **S55.3 The current hunting attractiveness**

With the exception of the effect of reported CWD, each of these components is normalized relative to its initial value (signified by INITIAL in equations). Each is raised to a user-defined exponent that represents hunter sensitivity to that factor (Table S15). Absolute values of exponents greater than one indicate hypersensitivity, while values less than one reflect reduced sensitivity approaching no effect as they approach zero.

The effect of mature bucks ($e_{mb_{[j]}}$; Eq. S56.1) is calculated as the current proportion of mature bucks in the harvest relative to the initial proportion, scaled to the hunter sensitivity to mature bucks ($s_{mb}$). The effect of deer sightings ($e_{sight_{[j]}}$; Eq. S56.2) is derived by comparing current total deer abundance to its initial value, reflecting the assumption that higher deer numbers increase sighting rates, scaled by the sensitivity to sightings ($s_{sight}$). The effect of cost ($e_{cost_{[j]}}$; Eq. S56.3) is calculated as the current net hunting cost relative to its initial value, raised to the hunter cost sensitivity ($s_{cost}$).

The direct effect of CWD on hunting attractiveness ($e_{CWD_{[j]}}$; Eq. S56.4) is modeled as an exponential decay function based on the reported CWD prevalence in harvested deer. The decay is scaled by a negative sensitivity parameter ($s_{CWD}$), such that higher prevalence reduces attractiveness. Reported prevalence is calculated as the number of CWD-positive tests ($R_{P_{[j]}}$; Table S11) divided by the total number of deer tested ($\sum_{l=1}^{n}\left(\sum_{k=1}^{n}\Gamma_{M_{[j,k,l]}} + \sum_{k*=1}^{n}\Gamma_{N_{[j,k*,l]}}\right)$; see *Testing*). If no testing has occurred, the prevalence is set to zero and has no effect.

The effect of expected hunt yield ($e_{yield_{[j]}}$; Eq. S56.5) is calculated as the inverse of the expected effort required to harvest a deer ($h_{effort_{[j]}}$), adjusted for the perceived disutility of harvesting a CWD-positive animal ($p_{CWD_{[j]}}$), and scaled by hunter sensitivity to yield ($s_{yield}$).

**Equation S56**

$$e_{mb_{[j]}} = \frac{\left. \sum_{k=5}^{n}\sum_{l=1}^{n=1}\left(h_{S_{[j,k,l]}}S_{[j,k,l]}+\sum_{y=1}^{n}h_{I_{[j,k,l]}}I_{[j,k,l,y]}\right) \middle/ \sum_{k=1}^{n}\sum_{l=1}^{n}\left(h_{S_{[j,k,l]}}S_{[j,k,l]}+\sum_{y=1}^{n}h_{I_{[j,k,l]}}I_{[j,k,l,y]}\right) \right.}{INTIAL\left[\left. \sum_{k=5}^{n}\sum_{l=1}^{n=1}\left(h_{S_{[j,k,l]}}S_{[j,k,l]}+\sum_{y=1}^{n}h_{I_{[j,k,l]}}I_{[j,k,l,y]}\right) \middle/ \sum_{k=1}^{n}\sum_{l=1}^{n}\left(h_{S_{[j,k,l]}}S_{[j,k,l]}+\sum_{y=1}^{n}h_{I_{[j,k,l]}}I_{[j,k,l,y]}\right) \right.\right]}^{s_{mb}}$$

**S56.1 Mature buck effect**

$$e_{sight_{[j]}} = \frac{N_{[j]}}{INTIAL\left[N_{[j]}\right]}^{s_{sight}}$$

**S56.2 The effect of sighting deer**

$$e_{cost_{[j]}} = \frac{H_{cost}}{INTIAL[H_{cost}]}^{s_{cost}}$$

**S56.3 The effect of costs associated with hunting**

$$e_{CWD_{[j]}} = e^{-s_{CWD}\left[ R_{P_{[j]}} \middle/ \sum_{l=1}^{n}\left(\sum_{k=1}^{n}\Gamma_{M_{[j,k,l]}}+\sum_{k*=1}^{n}\Gamma_{N_{[j,k*,l]}}\right) \right]}$$

**S56.4 The direct effect of chronic wasting disease prevalence**

$$e_{yield_{[j]}} = \left[\frac{(1-p_{CWD_{[j]}})}{h_{effort_{[j]}}}\right]^{s_{yield}}$$

**S56.5 The effect of expected effort for desired yield**

The perceived disutility of positives itself (Eq. S57) reflects two components: hunter awareness of CWD, calculated as the ratio of positive deer ($R_{P_{[j]}}$;Table S11) to active hunters ($H_{A_{[j]}}$; Table S13) in a region, and the salience of CWD ($\epsilon_{CWD}$), a user-defined constant ranging from zero (no perceived importance) to one (maximum importance). Following empirical evidence that many hunters continue to consume deer even after receiving a positive CWD test (Bradshaw et al.,

2021), we use a default value of 0.5 (Table S15). If no testing occurs, the perceived positive effect is set to zero.

**Equation S57. The perceived disutility of harvesting a CWD-positive deer**

$$p_{CWD_{[j]}} = \frac{R_{P_{[j]}}}{H_{A_{[j]}}} \epsilon_{CWD}$$

The expected harvest effort ($h_{effort_{[j]}}$) changes over time according to both a fixed baseline time component ($t_x$) and the perceived difficulty of harvesting a desired deer (Eq. S58). The perceived difficulty is based on the current perceived abundance ($N_{p_{[j]}}$) relative to its initial value, with the rate of change modulated by an information delay ($t_{effort}$) representing how quickly hunters perceive shifts in deer availability. Because travel time constitutes a substantial portion of overall effort, we set the baseline effort contribution at 0.75. This configuration assumes that underlying deer population dynamics influence only 25% of the total harvest effort, thereby reflecting that much of the time investment required for hunting is relatively invariant to fluctuations in local abundance.

**Equation S58. The change in the expected harvest effort**

$$\frac{dh_{effort_{[j]}}}{dt} = \frac{\left(t_x + (1 - t_x) INTIAL\left[N_{p_{[j]}}\right] / N_{p_{[j]}}\right) - h_{effort_{[j]}}}{t_{effort}}$$

Perceived deer abundance ($N_{p_{[j]}}$; Eq. S59) is computed as a weighted sum of desired available deer across cohorts. Preferences are encoded through weights for mature bucks ($\omega_{mb}$), any buck ($\omega_b$), and all nonclinical deer at all ($\omega_a$). The result is normalized to the initial population size ($N_{ref_{[j]}}$; Table S3), providing a relative index of how favorable the current population is to hunters based on their stated preferences.

**Equation S59. The perceived abundance of deer desired by hunters**

$$N_{p_{[j]}} = \frac{\omega_{mb} \sum_{k=5}^{n} \sum_{l=1}^{n=1} \left(S_{[j,k,l]} + \sum_{y=1}^{n} I_{[j,k,l,y]}\right) + \omega_b \sum_{k=2}^{n} \sum_{l=1}^{n=1} \left(S_{[j,k,l]} + \sum_{y=1}^{n} I_{[j,k,l,y]}\right) + \omega_a \sum_{k=1}^{n} \sum_{l=1}^{n} \left(S_{[j,k,l]} + \sum_{y=1}^{n} I_{[j,k,l,y]}\right)}{N_{ref_{[j]}}}$$

It is important to note that the relative influence of each factor contributing to hunting attractiveness is determined by user-defined constants, reflecting primarily the user's beliefs about the system with only a subset of parameters reflecting empirical data. This framework is not calibrated to observed hunter behavior but is instead designed to facilitate exploration of potential system feedbacks under varying assumptions about hunter sensitivities. In addition to modifying sensitivity parameters, users can adjust the underlying metrics themselves based on the specific questions being investigated. For instance, the net cost of hunting can be modeled endogenously as the difference between anticipated direct expenses—such as license fees and travel costs—and any incentives that offset these expenses, including pay-for-performance (P4P) programs or other cost-reducing interventions. This structure provides flexibility to simulate a wide range of hypothetical policy or behavioral scenarios within a consistent modeling framework.

**Table S15. Mapping between the full variable names used to construct the system dynamics model in Vensim (*Full Variable Name*) and the abbreviated forms (*Short Form*) used in the manuscript to represent the feedbacks between the deer-disease facets and hunting attractiveness.** The table includes each variable's units, the corresponding equation used to calculate it, or the source from which its value was derived.

| Short Form | Full Variable Name | Units | Equation/Source |
|---|---|---|---|
| $Å_{[j]}$ | Perceived Relative Attractiveness[Region] | dimensionless | Equation S55.1 |
| $Å_{ref_{[j]}}$ | Reference Attractiveness[Region] | dimensionless | Equation S55.2 |
| $Å_{h_{[j]}}$ | Hunting attractiveness[Region] | dimensionless | Equation S55.3 |
| $t_{Å}$ | Purchasing adjustment time | year | 1 |
| $e_{mb_{[j]}}$ | Effect of mature buck value[Region] | dimensionless | Equation S56.1 |
| $e_{sight_{[j]}}$ | Effect of sightings[Region] | dimensionless | Equation S56.2 |
| $e_{cost_{[j]}}$ | Effect of cost[Region] | dimensionless | Equation S56.3 |
| $e_{CWD_{[j]}}$ | Effect of CWD[Region] | dimensionless | Equation S56.4 |
| $e_{yield_{[j]}}$ | Effect of yield[Region] | dimensionless | Equation S56.5 |
| $s_{mb}$ | Sensitivity to mature bucks | dimensionless | **User dictated between 0 and 2** |
| $s_{sight}$ | Sensitivity to sightings | dimensionless | **User dictated between 0 and 2** |
| $s_{cost}$ | Sensitivity to cost | dimensionless | **User dictated between -2 and 0** |
| $s_{CWD}$ | Sensitivity to CWD | dimensionless | **User dictated between 0 and 2** |
| $s_{yield}$ | Sensitivity to yield | dimensionless | **User dictated between 0 and 2** |
| $h_{cost}$ | Hunter net cost[Region] | dollars/deer | **User dictated** |
| $p_{CWD_{[j]}}$ | Perceived positive effect[Region] | fraction | Equation S57 |
| $\epsilon_{CWD}$ | Relative valuation to positives | fraction | 0.5<br>***user dictated between 0 and 1** |
| $h_{effort_{[j]}}$ | Expected harvest effort[Region] | dimensionless | Equation S58 |
| $t_{effort}$ | Effort adj time | years | 3<br>***user dictated greater than 0** |
| $t_x$ | Frac time fixed | fraction | 0.75<br>***user dictated between 0 and 1** |
| $N_{p_{[j]}}$ | Perceived abundance[Region] | dimensionless | Equation S59 |
| $\omega_{mb}$ | Weight to mature bucks | fraction | 0.2<br>***user dictated between 0 and 1** |
| $\omega_b$ | Weight to bucks | fraction | 0.4<br>***user dictated between 0 and 1** |
| $\omega_a$ | Weight to any deer | fraction | (1-$\omega_{mb}$-$\omega_b$) |

### 3 Data and processing

#### 3.1 Summary

Publicly available data on white-tailed deer in Wisconsin were obtained from the Wisconsin Department of Natural Resources (WIDNR), both through their website and via direct provision. These data were compiled into Excel or csv files and included time series that were either used directly in our model or processed to generate cleaned and organized datasets, accounting for missing values prior to use as model variables.

The compiled data included deer population estimates at the county level from 2002 to 2023 and at the deer management zone (DMZ) level from 2002 to 2022. Information on deer hunting license purchases by county residents and nonresidents was available from 2005 to 2023, and annual county-level harvest data were provided from 2000 to 2023. Additional metrics indicating population age structure were provided, including fawn-to-doe ratios and yearling buck percentages at the DMZ level from 2007 to 2022, and at the county level from 2017 to 2023, along with yearling doe percentages at the county level from 2017 to 2023. County-level chronic wasting disease (CWD) surveillance data by age and sex were provided from 2001 to 2023. Genotype frequencies at codon 96 of the prion protein gene (PRNP), associated with CWD susceptibility, were available at five time points within the original infection area. Additional supporting data included the statewide Winter Severity Index from 2000 to 2021, the estimated land area and deer range within each county as of 2021, and estimated distances between county centers calculated using publicly available maps and the open-source software QGIS.

Because deer management practices in Wisconsin have changed over time—particularly in terms of the size and boundaries of management units—it was necessary to reconcile differences in spatial scale across datasets. Historically, deer management units (DMUs) were grouped into broader deer management zones (DMZs) based on ecological and demographic characteristics, whereas current management is conducted with DMUs that correspond with counties. The geography of CWD and deer management zones has varied over time. To integrate historical data into our model despite these changes, we mapped each county to its corresponding DMZ, allowing us to align spatially heterogeneous data sources. Four DMZs are recognized in WIDNR records: Central Farmland, Central Forest, Northern Forest, and Southern Farmland. Each DMZ comprises multiple DMUs, and in some cases, further subdivision into DMZ groups is used to account for population-level differences in deer demography and habitat conditions.

Our model operates primarily at the county level, which we define as the spatial unit of analysis (region $j$), but we include subscripts to associate counties with their respective DMZs or DMZ groups when higher-order aggregation is needed to match data sources. Table S16 presents the mapping of counties to DMZs, their associated model subscripts, and identifies which counties were used in the model. Although several counties fall within a given DMZ, only a subset was selected for inclusion based on data completeness and relevance to the modeled dynamics.

**Table S16. Deer management in Wisconsin has been conducted at varying spatial scales over the past three decades.** While each county is now treated as an individual deer management unit (DMU), historical management grouped counties into broader deer management zones (DMZs) based on shared habitat characteristics and deer demographic patterns. Some deer demographic metrics continue to be reported by subsets of these DMZs, referred to as DMZ Groups, to reflect differences in subpopulation dynamics. Each of the nine DMZ Groups is nested within one of four primary DMZs. A single DMU may be associated with more than one DMZ. The model described in this study uses the county level as the primary spatial unit of analysis (Region $j$) but incorporates additional subscripts to aggregate counties into DMZs or DMZ Groups where necessary to match data structure. This table lists the DMZs and DMZ Groups, the associated model subscript names, the DMUs included in each group, and the subset of counties actually used in the model.

| Deer Management Zone (DMZ/DMZ Group) | Model Subscript Name | Deer Management Units (DMU) | Counties Used in Model |
|---|---|---|---|
| Central Farmland/C Farmland | CFarm/G Cfarm | Clark, Green Lake, Marathon, Marinette, Marquette, Oconto, Outagamie, Portage, Shawano, Waupaca, Waushara, Wood | Marquette, Green Lake |
| Central Forest/ C Forest | CFor/G Cfor | Adams, Clark, Eau Claire, Jackson, Juneau, Monroe, Wood | Adams, Juneau |

| | | | |
|---|---|---|---|
| Northern Forest/NC Forest | Not used in model | Ashland, Iron, Langlade, Lincoln, Oneida, Price, Taylor, Vilas | N/A |
| Northern Forest/NE Forest | Not used in model | Florence, Forest, Marinette, Menominee, Oconto | N/A |
| Northern Forest/NW Forest | Not used in model | Bayfield, Burnett, Chippewa, Douglas, Rusk, Sawyer, Washburn | N/A |
| Southern Farmland/SE Farmland | SFarm/G SEfarm | Dane, Dodge, Green, Jefferson, Kenosha, Milwaukee, Ozaukee, Racine, Rock, Walworth, Washington, Waukesha | Dane, Dodge, Green, Jefferson, Rock, Walworth, Waukesha |
| Southern Farmland/SW Farmland | SFarm/G SWfarm | Columbia, Crawford, Grant, Iowa, Lafayette, Richland, Sauk, Vernon | Columbia, Crawford, Grant, Iowa, Lafayette, Richland, Sauk, Vernon |
| Central Farmlands/W Farmland | CFarm/G Wfarm | Adams, Barron, Buffalo, Chippewa, Dunn, Eau Claire, Jackson, Juneau, La Crosse, Monroe, Pepin, Pierce, Polk, St. Croix, Trempealeau | Eau Claire, Monroe |
| Central Farmland/Lake Michigan Farmland | CFarm/G Mfarm | Brown, Calumet, Door, Fond du Lac, Kewaunee, Manitowoc, Sheboygan, Winnebago | Sheboygan, Fond du Lac |

#### 3.2 Data sources

##### 3.2.1 Deer demographics

We used estimates of genotype frequencies, total population size, fawn-to-doe ratios, the proportion of yearling bucks in the antlered harvest, and the proportion of yearling does in the doe harvest to calibrate allele frequencies, population size, and the age and sex structure of deer within each region of the model.

###### *3.2.1.1 Genetics*

Time series data estimating genotype frequencies at the coding polymorphism 286G/A (G96S) in white-tailed deer were provided by the Wisconsin Department of Natural Resources (WIDNR) across five time periods (2004, 2017, 2018, 2019, 2020) near the original CWD detection area. The 2004 sample was collected shortly after the first CWD detection in Iowa and Dane counties; later samples were obtained as part of the Southwest Wisconsin Deer Study, including Iowa, Dane, and Grant counties. These data were used to calculate the frequency of the S allele in the Southwest (SW) region of Wisconsin and stored in the Vensim data variable *S Allele Freq Data[SW region]*. This frequency was used directly during model calibration to compare the model-predicted S allele frequency (*S Freq[SW region]*) with empirical values averaged across the three counties.

###### *3.2.1.2 Recruitment and age structure*

WIDNR began reporting deer statistics at the county level in 2017; before that, data were reported by larger management units (Table S14). Nine counties (Adams, Chippewa, Clark, Eau Claire, Jackson, Juneau, Oconto, Marinette, Wood) are split between two DMZs (forest and farmland), and deer metrics were estimated separately for each. Because the standard deviations for the two zones within each county overlapped for nearly all data points, we used data from the zone representing the larger area in each county, which is typically the farmland zone, except for Adams County.

WIDNR provided annual estimates (2017–2022) of the fawn-to-doe ratio (FDR), the proportion of harvested bucks that are yearlings (YBP), and the proportion of harvested does that are yearlings (YDP). These data were time-shifted to match the corresponding model time steps and stored as *data FDR county time match[Region]*, *data YBP county time match[Region]*, and *data YDP county time match[Region]*, respectively. These were used directly in model calibration to compare with model-predicted values: *sim fawn doe ratio[Region]*, *yearling harvest share[Region, buck]*, and *yearling harvest share[Region, doe]*.

The county level FDR, YBP, and YDP were used directly during model calibration. Model-predicted FDR (*sim fawn doe ratio[Region]*) was calculated as the ratio of nonclinical fawns to nonclinical does older than fawns (age classes 2–7), while model-predicted yearling harvest shares (*yearling harvest share[Region, buck]* and (*yearling harvest share[Region, doe]*) were the proportion of yearlings among harvested adults (age classes 3–7) for each sex. Model-predicted FDR was calculated at timestep 13 (Summer) to match the Summer Deer Observation survey period (August–September; Summer Deer Observations for DNR Staff and Cooperators | | Wisconsin DNR), while model-predicted yearling harvest shares were calculated at timestep 15 (Fall), corresponding with peak harvest season.

WIDNR also provided FDR and YBP metrics at the DMZ group level from 2007–2022. These were calculated for the Central Forest (*[G CFor]*) region, the Southeastern Farmland (*[G SEfarm]*; Dane, Green, Jefferson, Rock, Walworth, Waukesha, Dodge), the Southwestern Farmland (*[G SWfarm]*; Columbia, Crawford, Grant, Iowa, Lafayette, Richland, Sauk, Vernon), the Western Central Farmland (*[G Wfarm]*; Eau Claire, Monroe), and the Central Farmland (*[G Cfarm]*; Marquette, Green Lake) (refer to Table S14). These DMZ group metrics were also time-shifted to match the appropriate sampling periods, converted to proportions, and saved as *data DMZ FDR time match[DMZ group]* and *DMZ YBP time match[DMZ group]*. They were used in model calibration to compare model-predicted FDR (*sim DMZ fawn doe ratio[DMZ group]*) and YBP (*sim DMZ yearling buck frac[DMZ group]*) calculated using the same procedure as for counties. These data supplemented the county level FDR and YBP metrics, which only exist for 2017-2024, during calibration.

###### *3.2.1.3 Population estimates*

County-level post-harvest deer population estimates from accounting and sex-age-kill (SAK) models were provided by WIDNR annually from 2007–2020. These values were time-shifted to timestep 16 (post-harvest period at the end of the year) and stored in the Vensim variable *Post Hunt Deer Population Data time match[Region]*. They were compared directly to the model-predicted post-hunt population (*sim post hunt population[Region]*) during calibration. Estimates are not age-sex structured, limiting their utility for model initialization. The SAK estimates assume stationary harvest behavior, an assumption which was likely violated by large changes in antlerless harvest policy between 2003 and 2010.

Independent, non-SAK population estimates from the CWD management zone were available from 2002-2013 (Rolley, 2014). We scaled these data by the 2007-2010 average value to estimate a post-hunt index for the time series. We assumed the same scale over time to interpolate estimates for missing years of county-level abundance and saved the values as *scaled population from DMZ interp[Region]*. These data helped initialize population size (Eq. S60) and structure (Eq. S61) prior to the availability of county-level estimates.

**Equation S60. Estimation of the region-specific initial deer abundance**

$$initial\ population_{[j]} = init\ pop\ adj_{[j]} scaled\ population\ from\ DMZ\ interp_{[j]}$$

Each county's initial estimated population size at model start (year 2000) was multiplied by a free parameter *init pop adj[Region]* to account for uncertainty and saved as *reference population[Region]* (Eq. S61). To initialize age and sex structure, the reference population was multiplied by the variable *initial share[Region, age, sex]*, representing the proportion of individuals in each cohort, and saved as *reference population struc[Region, age, sex]*. This structure is used to initialize the susceptible stock and the reference aggregate harvest rate (*ref agg harvest rate[Region]*), which subsequently determined initial environmental prion levels (*Ref Prions[Region]*).

**Equation S61. Estimation of the region-specific initial age and sex distribution of deer**

$$reference\ population_{[j,k,l]} = discrete\ initial\ share_{[j,k,l]} initial\ population_{[j]}$$

Initial mortality and birth rates, along with aging rates, were used to estimate the initial share (*discrete initial share[Region, age, sex]*) in each age-sex cohort in accordance with a modified version of Little's Law (Stock = Inflow × Residence Time), adapted for discrete age groups (Eq. S62). Because survival modifies the number of deer reaching each age class, we incorporate the end-of-year survival rates for each cohort to estimate the expected number of deer per birth in each age and sex class at each time of the year in the discrete model (*survival to age at TOY[Region, age, sex, TOY]*). The initial share for each cohort is then calculated as the product of the yearly surviving population per birth—at the first timestep—and the relative initial age distribution, normalized by the sum across all cohorts

**Equation S62. Estimation of the initial share with consideration of the discretization of biological processes**

$$discrete\ initial\ share_{[j,k,l]} = \frac{survival\ to\ age\ at\ TOY_{[j,k,l]} rel\ init\ pop_{[j,k,l]}}{\sum_{k=1}^{n} \sum_{l=1}^{n} survival\ to\ age\ at\ TOY_{[j,k,l]}\ rel\ init\ pop_{[j,k,l]}}$$

The variable *rel init pop[Region,age,doe]* captured the relative size of each cohort at model initialization (Eq. S63) using two free parameters: *init age adj[Region]* (to adjust cohort growth rates) and *init buck sex ratio adj[Region]* (to adjust sex ratios). The initial age structure assumes a rapidly growing population with age distribution modelled as an exponential decay function of cohort age (*cohort age[age]*) scaled by a region-specific correction factor (*init age adj[Region]*). To estimate the initial relative population of bucks and does, the age distribution is adjusted by the initial buck sex share (*init buck sex ratio[Region]*)— multiplying for bucks and dividing for does.

**Equation S63. Estimation of the abundance and distribution of deer across cohorts at model initialization**

$$rel\ init\ pop_{[j,k,l]} = \begin{cases} e^{\left(init\ age\ adj_{[j]}\left(cohort\ age_{[j]} - \frac{cohort\ age_{[k=7]}}{2}\right)\right)} init\ buck\ sex\ ratio_{[j]} & l = 1 \\ \dfrac{e^{\left(init\ age\ adj_{[j]}\left(cohort\ age_{[k]} - \frac{cohort\ age_{[k=7]}}{2}\right)\right)}}{init\ buck\ sex\ ratio_{[j]}} & l = 2 \end{cases}$$

#### 3.2.2 Hunter harvest

Annual hunter harvest data were used as a model constraint during estimation to allow harvest rate estimation based on observed values.

##### *3.2.2.1 Harvest data*

County-level data from 2000 to 2023 were directly assigned to Vensim variables to reflect the number of antlered (*data antlered harvest[Region]*) and antlerless (*data antlerless harvest[Region]*) deer harvested each year. The total annual harvest (*data total harvest[Region]*) was calculated as the sum of both harvest types within each county. To align with the model's time step for simulated harvest ($t_{step}$=15), which corresponds to Fall (when most harvest occurs), the antlered and antlerless harvest data were time-shifted and saved as *data antlered harvest time match[Region]* and *data antlerless harvest time match[Region]* respectively. These adjusted values are used to estimate historical harvest rates (refer to *Historic harvest rates*) which in turn inform harvest rates applied within the model's mortality structure (refer to *Human caused mortality*).

##### *3.2.2.2 Historic harvest rates*

Historic harvest rates for both antlered and antlerless deer were estimated using the time-aligned harvest data during model calibration. This approach allows for uncertainty in reported population numbers and provides a foundation for harvest-based scenario modeling. The historic harvest rates (Eq. S64) are calculated as the proportion of the weighted population removed through harvest each year and saved as the Vensim variables *Indicated historic buck harvest rate[Region]* and *Indicated historic antlerless harvest rate[Region]*. To account for wounding loss—deer that are shot but not recovered—we adjust the reported harvest numbers upward by 7% (*unreported wounding mortality*), a correction based on literature values (Aebischer et al., 2014; Ditchkoff et al., n.d.; Fuller, 1990; Jennings et al., 2014; Pedersen et al., 2008; Wallingford et al., 2017). This adjustment reflects unreported mortality during the hunting season, while other forms of background unreported mortality (e.g., poaching) are handled separately through background mortality rates.

**Equation S64. Estimation of historic harvest rates**

$$Indicated\ historic\ antlered\ harvest\ rate_{[j]} = \frac{data\ antlered\ harvest\ time\ match_{[j]}(1 + unreported\ wounding\ mortlaity)}{weighted\ nonclinical\ buck\ population_{[j]}} \quad \textbf{S64.1 Antlered deer}$$

$$Indicated\ historic\ antlerless\ harvest\ rate_{[j]} = \frac{data\ antlerless\ harvest\ time\ match_{[j]}(1 + unreported\ wounding\ mortlaity)}{weighted\ nonclinical\ antlerless\ population_{[j]}} \quad \textbf{S64.1 Antlerless deer}$$

The weighted populations exclude clinical (symptomatic) deer, under the assumption that hunters actively avoid visibly sick individuals. The weighted buck population includes all bucks yearling and older. Fawns—which are classified as antlerless regardless of sex—are included in the weighted antlerless population but are downweighted using age-specific harvest weights (*age harvest weight[age]*) to reflect the lower likelihood of fawn harvest.

To prevent overharvest leading to extinction, the resulting harvest rates are capped at 0.8/year, based on empirical analysis of observed maximum county-level harvest-to-population ratios. This effect primarily applies to future scenario experiments rather than the historical period.

For initialization, the denominators are calculated as the product of the initial population (Eq. S60), initial sex share (assuming 30% bucks), and an initial harvest rate adjustment (both set to 1 here).

The antlerless harvest rate variable includes additional functionality to allow user-defined adjustments during scenario testing. This enables simulation of regulatory interventions such as Earn-a-Buck policies or targeted reductions in antlerless harvest pressure, rather than relying solely on historic data-derived rates.

##### 3.2.3 Chronic wasting disease

We used chronic wasting disease (CWD) surveillance data to calibrate disease dynamics within the model and estimated region-specific parameters to dictate the magnitude of initial disease introduction, assuming introduction was one year prior to first detection.

###### *3.2.3.1 CWD surveillance data*

County-level CWD surveillance data from 2002 to 2023 were used to determine the total number of tests performed and the number of positive detections within each age-sex cohort and region. The surveillance data did not distinguish between four- and five-year-old deer, so we loaded the data using an alternative age classification. We did not attempt to correct for potential aging inaccuracies that may arise from dentition-based age estimation (Adamsid and Blanchong, 2020). Instead, we used the age classes as assigned by Wisconsin DNR biologists at the time of data collection.

To reconcile differences between the model's age structure *k* (*age*: fawn, yearling, 2,3,4,5,6+) and the surveillance data structure *k** (*age data*: fawn, yearling, 2, 3, 4&5, 6+), we grouped model-predicted CWD prevalence and positives accordingly, enabling a direct comparison between model predictions and surveillance data. If testing was conducted within a given cohort and region but no positives were detected, a value of zero was recorded, indicating a true zero. In contrast, if no testing was conducted, the value was marked as missing (NA), distinguishing it from an observed negative result.

Although the CWD sampling year spans April 1 – March 31, given that most samples originate from hunter-harvested deer,we time-shifted the data to match the model's harvest time step ($t_{step}$=15, corresponding to December). The total number of CWD tests, stratified by region, age, and sex, were saved in the Vensim variable *data surv N samples w0s time match[Region, age data, sex]*, while the number of positive tests was stored as *data surv Positive w0s time match[Region, age data, sex]*.

These data informed the model's annual sampling rate and assisted in calibrating the expected number of positive tests predicted by the model for each cohort, enabling us to account for discrepancies between observed prevalence and true underlying prevalence. The total number of samples drive the regional sampling rate data variable (*Sampling[Region, age data, sex]*) and the number of observed positives is compared to those predicted by the model for each cohort (*expected positive[Region, age data, sex]*). We also calculated the total number of positives per region to support regional-scale model fitting to provide a larger sample size for evaluating model fit (refer to Estimation method).

###### 3.2.3.2 Disease introduction

To simulate realistic disease dynamics, CWD was introduced into the model in counties where the disease had been detected prior to 2024. Each affected county was assigned a unique pulse time (*Pulse time[Region]*), denoting the year when a proportion of susceptible individuals were transitioned into the infected class. We assumed that CWD was present at least one year prior to the first documented positive case, a conservative estimate given the disease's long incubation period and typical delay between introduction and detection.

Initial infections were assumed to be evenly distributed across infection stages. The fraction of susceptible individuals converted to infected during the pulse was estimated from the surveillance data (*Pulse infections[Region]*), allowing the model to match observed patterns by adjusting the intensity of disease seeding (refer to Estimation method). These pulse values were fixed for subsequent simulations following model calibration.

##### 3.2.4 Landscape values

We incorporated several landscape-level variables to aid in model calibration, including inter-regional distances, historical winter severity, and deer habitat area.

###### 3.2.4.1 Distance matrix

Distances between Wisconsin counties were estimated using publicly available county shapefiles in QGIS. We calculated the Euclidean distance between county centroids and normalized these values by dividing by the shortest observed distance, resulting in a relative distance matrix for spatial analyses.

###### 3.2.4.2 Winter severity index

Annual winter severity index (WSI) data for each county and the statewide average were obtained from the WIDNR (Deer Statistics (wi.gov)). We used statewide average values from 2000–2024 and stored these in Vensim as *winter severity index*. To synchronize the reported WSI values with model timing, we shifted each value back by half a year (e.g., WSI for 2021 was applied from model year 2021.5 to 2022.5) and saved the time-aligned data as *time centered WSI*. The mean WSI over the period was stored as *mean WSI 2000 2024*.

###### 3.2.4.3 Land area

Total county area and deer habitat area were sourced from U.S. Census and WIDNR datasets for 2021, respectively. These were assigned to the Vensim data variables *county land area[Region]* and *deer range 2021[Region]*. We assumed that both the total and deer-suitable land area remained constant throughout the 2000–2024 simulation period.

##### 3.2.5 Hunter license data

Hunter license purchasing behavior and demographic information were integrated into the model using license and harvest data provided by WIDNR, supplemented with publicly available census data. Data from all 72 Wisconsin counties were included in hunter submodel fitting.

###### *3.2.5.1 Regular license harvest*

We defined "regular license" harvest as deer taken under licenses that allow a single individual to legally kill a set number of deer during the standard hunting season. These are distinct from agriculture or farmland nuisance permits that allow a much larger number of deer to be taken under a single permit. To estimate harvest per individual hunter, we filtered data to include only deer reported under regular license harvest.

Antlered and antlerless harvests were recorded by sex and age (fawn or older) for most years. For 2006–2008, harvests were only classified into antlered and antlerless categories. Separate time series of regular license and special permit harvests were prepared for each county from 2005–2023.

###### *3.2.5.2 Active and inactive hunters*

License data were available from 2005-2023 at the county level (i.e., the hunter's county of residence) and included a count of those hunters that lived in-state (residents) or out-of-state (non-residents). Active hunters were defined as unique individuals who purchased any deer-hunting license in a given year. These values were saved in Vensim as *A[Region]*, then time-shifted to align with harvest (*A opt[Region]* at timestep = 15). We used WI DNR survey data (Wisconsin Wildlife Reports | | Wisconsin DNR) to estimate that, on average, 95% of licensed hunters attempt to harvest a deer, which we defined as the mobilization rate (*mobilization rate*).

To estimate county-to-county hunter movements, we used data on each hunter's county of residence and county of harvest. The proportion of hunters from each county harvesting in other counties (including their own) was stored in the Vensim variable *Hunter harvest region[FromRegion, ToRegion]*. These proportions showed little variation over time, thus they were assumed to be constant and applied to all hunters, including those who did not successfully harvest. These proportions dictate the patterns of travel for harvest and thus the total number of mobilized hunters harvesting in each county as the product of the number of active hunters in a county and the movement matrix.

We tracked hunter inactivity using license purchase records beginning in 2006, as the first observations of active hunters were in 2005. Hunters who skipped a year were assigned to an "inactive" class, with additional years of inactivity progressing them through a series of five inactive states (1–5+ years). Individuals with five or more years of inactivity were pooled in the final stage. Because mortalities are unobserved and indistinguishable from long-term inactivity in the dataset, deaths also accumulate within this fifth stage. Similarly, customers that first purchased a license prior to 2005 and remained inactive for the observed period should be in this fifth stage but are unobserved and thus unaccounted for, leading to us allowing for larger error surrounding this stage. These values were saved as *I[Region, Inactive]* and time-shifted to *I opt[Region, Inactive]* to align with the harvest time step. Given the uncertainty of real mortality and unobserved inactive hunter numbers, these data are not used in model fitting but rather as a qualitative check on results.

Reactivation events were tracked for each inactive class beginning with the third year of data. For example, reactivation from one year of inactivity could first be observed in 2007, two consecutive years of inactivity in 2008, and so on. These values were saved as *Ireacts opt[Region, Inactive]*. These count data contain less measurement error than the total number of inactive hunters since they are concretely observed and do not include unobserved mortalities. However, uncertainty remains in the reactivation of those who have been inactive for up to the first five years of the dataset since customers who had been active prior to our dataset would be misclassified as recruitment. Recruitment of new hunters was estimated from 2010–2023 using the established inactive hunter pool, with counts stored as *PRecruit opt[Region]*. Reactivation and recruitment events were time-shifted to $t_{step}$ = 14 (0.8125) and divided by the step size (0.0625) to align with the single pulse structure in the model.

###### 3.2.5.3 Potential hunters and mortality

County-level population estimates from the U.S. Census (U.S. Census Bureau, 2024, 2020, 2010) were used to represent the total potential hunter pool. These values included all residents regardless of age or health and served as an upper bound on possible recruitment. Population estimates were saved as *P[Region]* and time-shifted to *County Pop[Region]* for alignment with the timestep prior to harvest ($t_{step}$=14). Given that the census data already includes mortality and births, we simply update the total population each year.

We estimated county-specific mortality rates for the hunting population by weighting age- and sex-specific rates from the U.S. CDC WONDER database (U.S. Department of Health and Human Services, 2024) to reflect the demographics of active and inactive hunters (stage-specific), who were assumed to be aged >5 years and primarily white, non-Hispanic males. Mean mortality rates for active and inactive hunters were saved as *A mort mean[Region]* and *I mort mean[Region, Inactive]*, respectively.

###### 3.2.5.4 Wildlife value orientation

Following Manfredo et al. (2008) and the America's Wildlife Values project, we incorporated wildlife value orientations (WVO) as a determinant of participation in hunting wild game. Approximately 36% of Wisconsin residents were categorized as "traditionalists," indicating alignment with extractive wildlife use, and 21% as "pluralists," who hold these values alongside those typical of "mutualists," who value increased rights or protections for wildlife (Dietsch et al., 2018). We used their sum (57%) as a conservative upper bound on the proportion of the population that might ever consider engaging in hunting, regardless of other social or demographic characteristics.

To account for temporal shifts in public attitudes, we applied a statewide annual decline of -0.5% (SD = 0.3%) in this combined proportion, consistent with recent findings across Western states

(Manfredo et al., 2021b). The slope was used to project forward and backward from the existing WI 2018 datapoint of 57% (Dietsch et al., 2018). During model fitting, the WI WVO baseline and rate of decline were allowed to vary as free parameters, using the stated values as priors, within a hierarchical framework to accommodate variation across individual counties in Wisconsin (refer to *Hunter submodule*).

### 4 Estimation method

#### 4.1 Susceptible-Infected model

##### 4.1.1 Estimation procedure

We estimated model parameters by simulation of the system of differential equations at the county level (Region = County), incorporating age, sex, and infection-status disaggregation as described in Section 2 (*Model structure and key formulations*). All features of the disease-host system are estimated simultaneously in a Bayesian hierarchical approach, with extensive use of priors from literature and subject matter expertise.

Calibration of the SIC core of the model relies primarily on time-series data of white-tailed deer populations and CWD surveillance provided by the Wisconsin Department of Natural Resources (refer to *Data sources*) to estimate demographic and epidemiological parameters (Table S17). Six key metrics were used in model fitting: 1) fawn to doe ratio, 2) ratio of yearling bucks to total harvested bucks, 3) ratio of yearling does to total harvested does, 4) post-hunt population estimate, 5) the frequency of the *S* allele, and 6) the number of tested CWD positive deer within each age and sex cohort. To extend the datasets of fawn to doe ratio and ratio of yearling bucks to total harvested bucks, we fit the model to both the DMZ level and county-level data.

A log-normal error distribution was assumed for all datasets, except the CWD-positive count data, which were modeled using an overdispersed Poisson distribution to account for variance inflation typical in epidemiological count data. The model was constrained by the annual county-level hunter harvest, used as an exogenous driver.

The model was initialized with a fully Susceptible deer population. Infected individuals were introduced via a one-time pulse in each county one year prior to the first documented CWD detection (Table S17; *Pulse Infections[Region]*).

To account for heterogeneity across counties not explicitly captured by the core model structure, we implemented a hierarchical estimation approach. This allowed county-level variation in transmission dynamics and demographic characteristics to be absorbed by random-effect-like parameters (Table S17), including:

- *relative transmission[Region]* (county-specific transmission modifier)
- *Relative CC[Region]* (initial population pressure modifier)
- *init pop adj[Region]* (initial population adjustment factor)

- *init buck sex ratio adj[Region]* (initial male-to-female ratio modifier)
- *init age adj[Region]* (initial age structure adjustment factor)

We conducted two rounds of model fitting. A full fit, using the complete time series of all six datasets, and a restricted fit, using a dataset that excluded the final four years of the time series datasets (i.e., truncated at 2019) to test the model's generalizability and forecasting ability.

#### 4.1.2 Estimation results

##### *4.1.2.1 Parameters*

**Table S17. Summary of the epidemiological and demographic parameters estimated during calibration of the susceptible-infected-clinical (*SIC*) model of chronic wasting disease (CWD) in white-tailed deer, including their associated priors and posterior distributions.** Each variable is reported using its internal Vensim name for transparency and reproducibility. Priors were specified directly for each parameter where data or expert judgment permitted, and in some cases, priors were defined for a product of multiple parameters (e.g., realized birth rate). To prevent the optimization algorithm from exploring biologically implausible regions of the parameter space (e.g., negative birth rates), we imposed hard bounds (minimum and maximum values) on each parameter. Where possible, citations for prior values informed by the empirical literature are provided. Final parameter estimates are reported as the posterior mean ± standard deviation, derived from the joint posterior distribution sampled during the fitting process along with their associated univariate Rubin/Brooks-Gelman potential scale reduction factor (PSRF; Brooks and Gelman, 1998). PSRF values approaching 1 indicate successful convergence, whereas values greater than 1.2 suggest non-convergence. Maximum PSRF values are reported for region specific parameters.

| Variable | Definition | Informative Prior [Min, Max] | Estimated Values | PSRF | Literature |
|---|---|---|---|---|---|
| *Chronic Wasting Disease* | | | | | |
| Direct beta | The base instantaneous rate of disease transmission through direct contact between deer. This rate is the product of the average per capita deer-deer contact rate and the probability of infection given contact between a susceptible and an infected deer. | *Lognormal*(1.2, 0.2) [1x10$^{-6}$, 3] | 2.61±0.0549 | 1.00 | (Jennelle et al., 2014; Miller et al., 2000; Potapov et al., 2013; Wasserberg et al., 2009) |
| Indirect beta | The base instantaneous rate of disease transmission through contact between deer and CWD prions in the environment. This rate is the product of the average per capita deer-environment contact rate and the probability of infection given contact between a susceptible deer and an infected environment. | *Lognormal*(0.1, 0.2) [1x10$^{-6}$, 3] | 0.100±0.00845 | 1.00 | (Sun et al., 2015) |
| Maternal transmission beta | The instantaneous rate of disease transmission through contact between fawns and dams. This rate is the product of the average per capita fawn-dam contact rate and the probability of infection given contact between a susceptible fawn and an infectious doe. | *Lognormal*(0.05, 0.2) [1x10$^{-6}$, 1] | 0.0504±0.00496 | 1.00 | (Miller et al., 2000) |
| density dependence & | A scaling parameter describing how transmission rates (direct and indirect) respond to temporal changes in deer density within a | *Normal*(0.5, 0.25) [0,1] | 0.601±0.0122 & 0.555±0.0114 | 1.00 & 1.00 | (Storm et al., 2013) |

| Variable | Definition | Informative Prior [Min, Max] | Estimated Values | PSRF | Literature |
|---|---|---|---|---|---|
| envir density dependence | county. A value of one implies classic density-dependent transmission; a value of zero implies frequency-dependent transmission; intermediate values reflect partial density-dependence. | | | | |
| abs density dependence | A modifier capturing the regional (i.e., cross-sectional) variation in transmission associated with absolute differences in deer density between counties. | *Uniform*(0,1) | 0.394±0.0121 | 1.01 | |
| prion half life | The number of years that CWD prions persist in the environment before degrading by half. | [1,16] | 8.67±0.243 | 1.00 | (Almberg et al., 2011; Kjær and Schauber, 2022; Maloney et al., 2020; Miller et al., 2006; Potapov et al., 2013; Sun et al., 2015) |
| carcass deer year equivalent | A scaling factor representing the proportionate contribution of infectious carcasses to environmental prion load relative to living infected deer. | *Uniform*(0.1, 10) | 13.9±0.169 | 1.00 | (Davenport et al., 2018; Miller et al., 2004) |
| Exposed duration | The average number of years a deer remains in the exposed (non-infectious) stage after CWD transmission. | *Uniform*(0.3, 1.2) | 0.764±0.0113 | 1.00 | 0.6 (Heisey et al., 2014; Mathiason, 2023; Robinson et al., 2012; Wasserberg et al., 2009) |
| Infectious duration | The average number of years a deer remains in the infectious stage before progressing to the clinical phase. | *Uniform*(0.35,1.4) | 0.956±0.0184 | 1.00 | (Heisey et al., 2014; Mathiason, 2023; Robinson et al., 2012; Wasserberg et al., 2009) |
| genetic effect lifespan E & genetic effect lifespan I | The multiplicative effect of the *S* allele on survival time in the exposed (E) and infectious (I) stages of CWD after infection. A value of one indicates no effect; values greater than one increase lifespan. | *Lognormal*(0.5, 1.3) [0.2, 5] | 4.12±0.0716 & 0.946±0.0558 | 1.00 & 1.00 | Johnson 2011, Hoover 2017 |
| genetic effect transmission | The multiplicative effect of the *S* allele on transmission (i.e., the probability of infection given contact). A value of one indicates no reduction in transmission due to the allele. | *Lognormal*(1, 0.5) [0, 2] | 0.908±0.0281 | 1.01 | (Denkers et al., 2024) |

| Variable | Definition | Informative Prior [Min, Max] | Estimated Values | PSRF | Literature |
|---|---|---|---|---|---|
| selection time | The number of years after CWD introduction that must pass before genetic selection pressure begins to shift G96S allele frequencies in the population. | *Uniform*(1,20) | 6.00±0.179 | 1.00 | |
| Pulse infections[Region] | The estimated proportion of deer infected one year prior to the first CWD-positive test in a region. | *Uniform*($10^{-6}$,0.1) | $(4.84 \pm 4.23)\times10^{-3}$<br><br>($7.54\times10^{-6}$, 0.0202)<br>*mean±SD across all regions, min,max mean | 1.00 | |
| relative transmission[Region] | A constant multiplier that adjusts transmission rates to account for average regional variation in direct and indirect transmission. | *Uniform*(0.5, 2) | 1.02±0.0704<br><br>(0.804, 1.32)<br>*mean±SD across all regions, min, max mean | ≤1.01 | Centered around the actual estimate and allowed to halve or double. |
| age eff contacts[older] | A scaling factor for the relative difference in contact or probability of infection given contact by age class, expressed relative to the youngest age class (*fawns*)<br>*older: yearling, two, three, four, five, old* | *Uniform*(1, 4) | 1.42±0.0397<br>2.08±0.0502<br>1.22±0.0467<br>2.91±0.0438<br>1.03±0.0169<br>1.01±0.00797 | ≤1.01 | |
| sex eff contacts[buck] | A scaling factor indicating the relative difference in contact or probability of infection given contact by sex. A value of one means equal risk, and values greater than one reflect higher risk in bucks. | *Uniform*(1, 4) | 1.64±0.0424 | 1.00 | |
| relative envir contact diversity | A parameter allowing environmental contact rates to share the same normalization framework as deer-deer contacts while varying in magnitude. | *Uniform*(0, 1) | $(3.59 \pm 2.20)\times10^{-3}$ | 1.00 | |
| positive overdispersion | The variance term for the count data of CWD-positive deer, modeled with a Poisson distribution with overdispersion. | *Uniform*(0.01, 1) | 0.0455±0.0149 | 1.00 | |
| CWD MZ bias | A correction factor that accounts for any additional biases in population estimates specific to the original CWD management zones. | *Uniform*(0.5, 2) | 1.14±0.0237 | 1.01 | Centered around the actual estimate and allowed to halve or double. |
| *Deer Demographics* | | | | | |
| Ref Birth Rate | The instantaneous rate of the baseline per capita births for reproductive does, invariant of population pressure. Bounds are imposed on the reference birth rate, but a prior is imposed on the realized birth rate (which includes population pressure effects) to | *Uniform*(1, 2.5) on reference birth rate<br><br>*Lognormal*(1.8, 0.25) on the realized birth rate | 1.73±0.0104 | 1.01 | (McCaffery et al., 1998) |

| Variable | Definition | Informative Prior [Min, Max] | Estimated Values | PSRF | Literature |
|---|---|---|---|---|---|
| | avoid biologically implausible values. | | | | |
| Birth CC sensitivity | Governs how quickly the birth rate responds to deviations between the current population size and the level at which population pressure begins to occur. This log-normal prior is jointly optimized with the *mortality CC sensitivity*. | *Lognormal*(1, 0.1) [0,4] | 0.406±0.0241 | 1.00 | (McCaffery et al., 1998) |
| mortality CC sensitivity | Governs how quickly the natural mortality rate responds to deviations between the current population size and the level at which population pressure begins to occur. This log-normal prior is jointly optimized with the *Birth CC sensitivity*. | *Lognormal*(1, 0.1) [0,2] | 2.22±0.0458 | 1.00 | (Carstensen et al., 2009; Vreeland et al., 2004) |
| frac mort variable | The proportion of natural mortality that varies seasonally or with population pressure. The proportion that is not variable accounts for senescence (i.e., age-related physiological decline). | *Lognormal*(0.5, 0.1) [0.01,0.99] | 0.781±0.0125 | 1.00 | |
| base fawn mortality | The additional annual natural mortality rate for fawns beyond the first few weeks of life, attributed specifically to age-related hazards. Jointly estimated with *early fawn mortality*. | *Lognormal*(0.2, 0.25) [0.001,0.5] | 0.111±0.00636 | 1.00 | (Carstensen et al., 2009; Vreeland et al., 2004) |
| early fawn mortality | The additional annual natural mortality rate for fawns within the first few weeks of life, attributed specifically to age-related hazards. Jointly estimated with *base fawn mortality*. | *Lognormal*(0.4, 0.25) [0.1,1] | 0.780±0.0123 | 1.00 | (Carstensen et al., 2009; Vreeland et al., 2004) |
| Relative CC[Region] | A region-specific constant that modifies the difference between the initial reference population and the initial population pressure level. A value of one indicates equivalence; less than/greater than one indicates that the reference population was above/below the population pressure level, respectively. | *Uniform*(0.5, 2) [0.5,2] | 1.16±0.196<br><br>(0.822, 1.65) *mean±SD across all regions, min, max mean | ≤1.01 | Centered around the actual estimate and allowed to halve or double. |
| sex rel age dispersion[doe] | A scaling factor describing the relative impact of sex to age on contacts. Fixed at one for bucks and adjusted for does. As the value approaches zero, the impact of age on doe contacts is removed. | *Uniform*(0, 1) | $(4.44 \pm 2.63) \times 10^{-3}$ | 1.00 | (DIEFENBACH et al., 2008; Long et al., 2008; Lutz et al., 2015; Nixon et al., 2007) |

| Variable | Definition | Informative Prior [Min, Max] | Estimated Values | PSRF | Literature |
|---|---|---|---|---|---|
| init pop adj[Region] | A regional adjustment factor to account for uncertainty in the estimated population size at model initialization. | *Uniform*(0.5, 2) | 0.978±0.139 (0.708, 1.34) *mean±SD across all regions, min,max mean | ≤1.02 | Centered around the actual estimate and allowed to halve or double. |
| init buck sex ratio adj[Region] | A regional adjustment factor to account for intraregional variation in sex ratios at model initialization. | *Uniform*(0.5, 2) | 0.958±0.232 (0.500, 1.57) *mean±SD across all regions, min, max mean | ≤1.01 | Centered around the actual estimate and allowed to halve or double. |
| init age adj[Region] | A regional adjustment factor to account for intraregional variation in age structure at model initialization. | *Uniform*(-1, 1) | -0.0364±0.220 (-0.905, 0.469) *mean±SD across all regions, min, max mean | ≤1.01 | Centered around the actual estimate and allowed to halve or double. |
| mortality WSI sensitivity | Governs how quickly the mortality rate responds to deviations away from the average winter severity value. | *Uniform*(0,1) | $(4.20±2.33)x10^{-3}$ | 1.01 | |
| ref background mortality rate | The baseline annual mortality rate for non-clinical deer in the absence of population pressure. | *Lognormal*(0.1, 0.2) [0.001, 0.15] | 0.0666±0.00151 | 1.01 | (Al-Arydah et al., 2012; Jennings et al., 2014) |
| age harvest weight [fawn] | A scaling parameter that reduces the effective harvest rate of fawns relative to older age classes. | *Uniform*(0,1) | 0.732±0.00878 | 1.00 | Full range of possibilities (Wojcik and Stenglein, n.d.) |
| nonrange weight | A scaling factor comparing the county land area to the defined deer range area. Low values suggest that the defined deer range closely approximates true deer land area use. | *Uniform*(0, 1) | 0.312±0.0126 | 1.00 | Full range of possibilities |

##### *4.1.2.2 Maps of region-specific parameter*

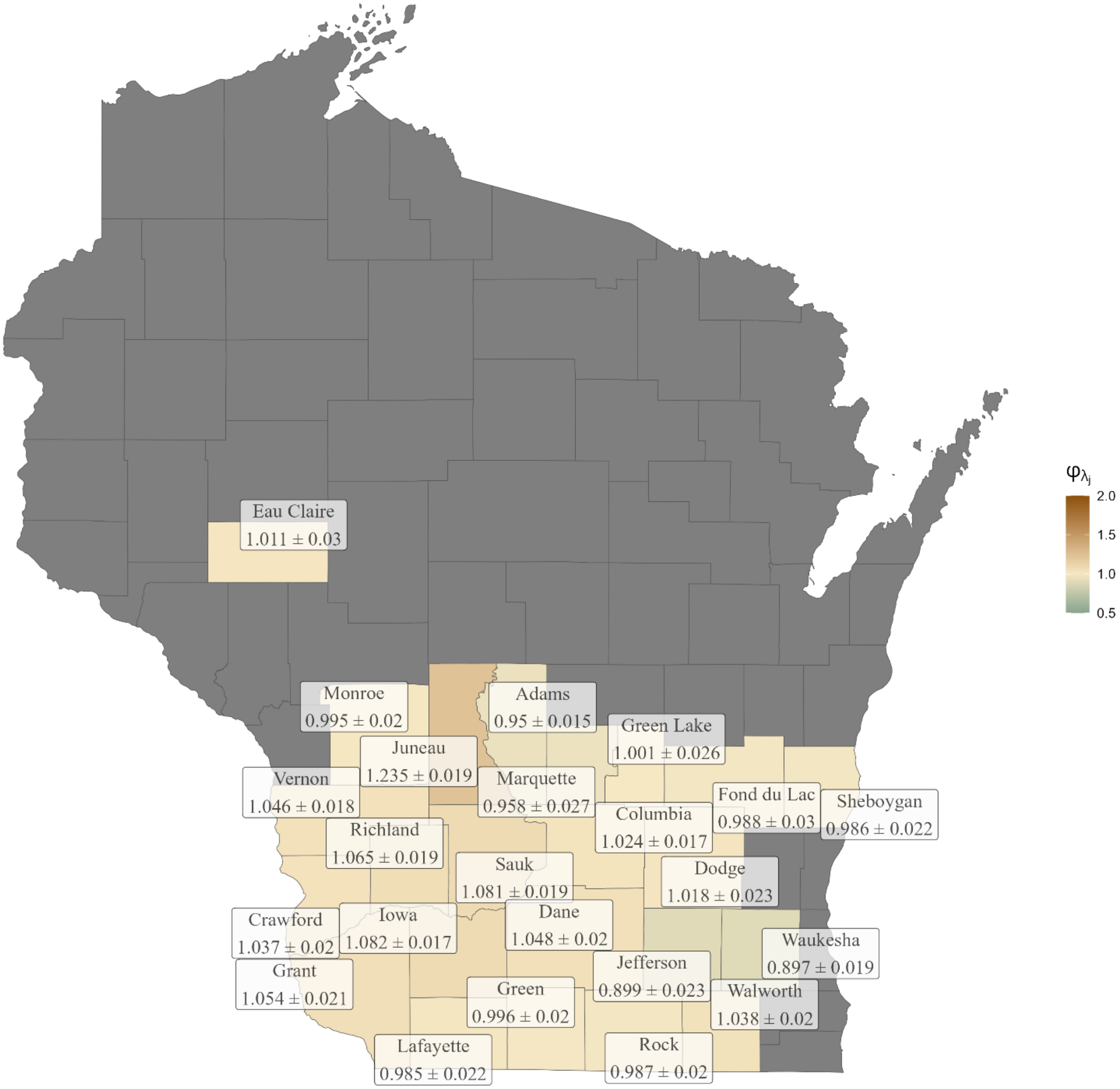


**Figure S1. County specific mean and ± 1 standard deviation of regional relative transmission estimated from the susceptible-infected-clinical model of chronic wasting disease in white-tailed deer in Wisconsin, fit to the full dataset.** A value of one indicates that a county's transmission rate matches the global mean; values greater than and less than one indicate higher and lower transmission than the global mean respectively. Variation can be interpreted as capturing a variety of unmodeled features, including uncertainty in density dependent effects due to imperfect or time-varying deer range and unobserved habitat effects on deer mobility.

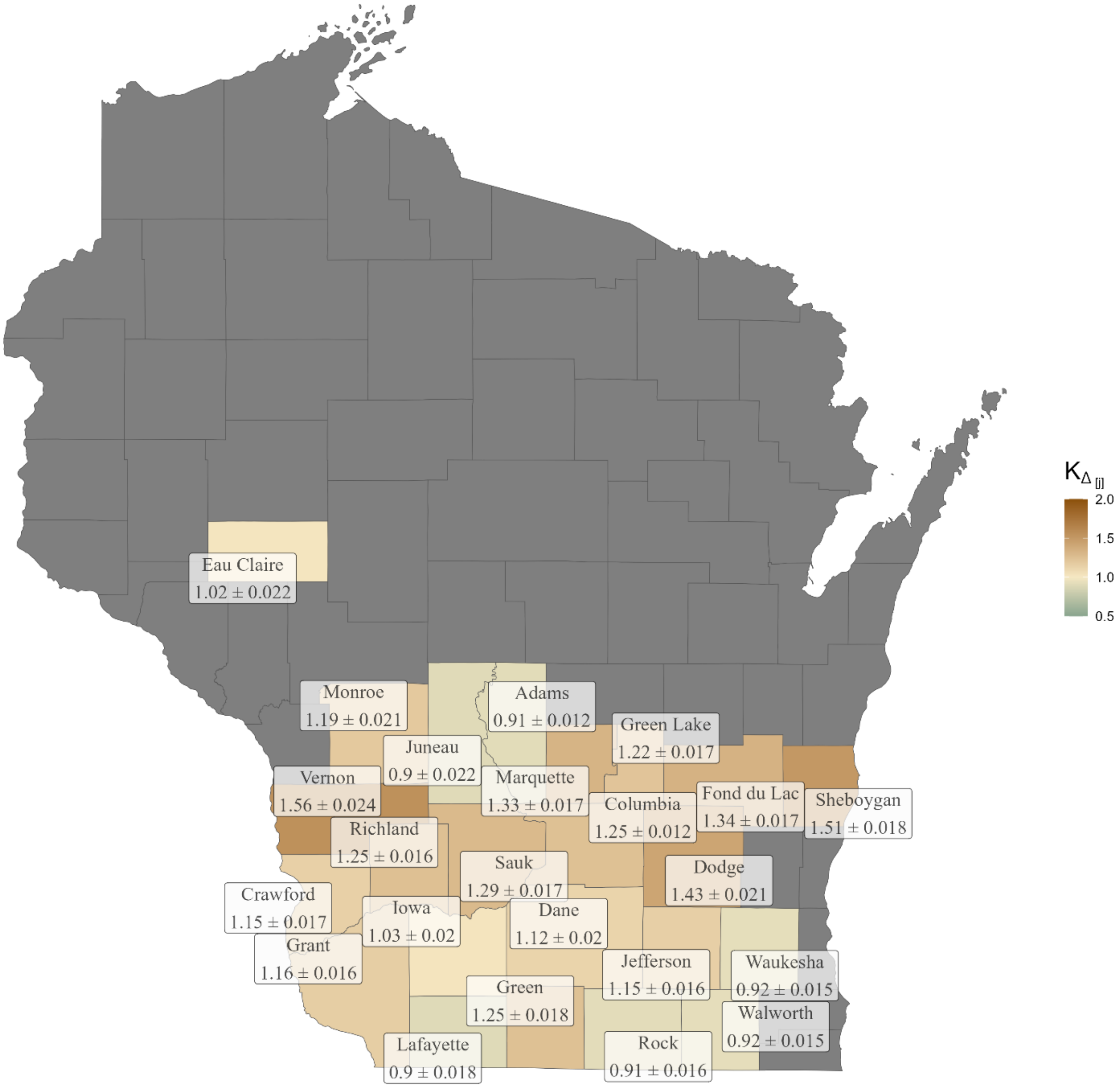


**Figure S2. County specific mean and ± 1 standard deviation of regional relative population pressure estimated from the susceptible-infected-clinical model of chronic wasting disease in white-tailed deer in Wisconsin, fit to the full dataset.** Population pressure reflects ecological limits on recruitment and mortality, akin to a traditional carrying capacity. A value of one for the relative population pressure indicates that a county's initial reference population was exactly equal to the initial population pressure level; values greater than and less than one indicate initial reference population is higher and lower than initial population pressure level respectively. These effects are difficult to interpret due to the low fidelity of population data used.

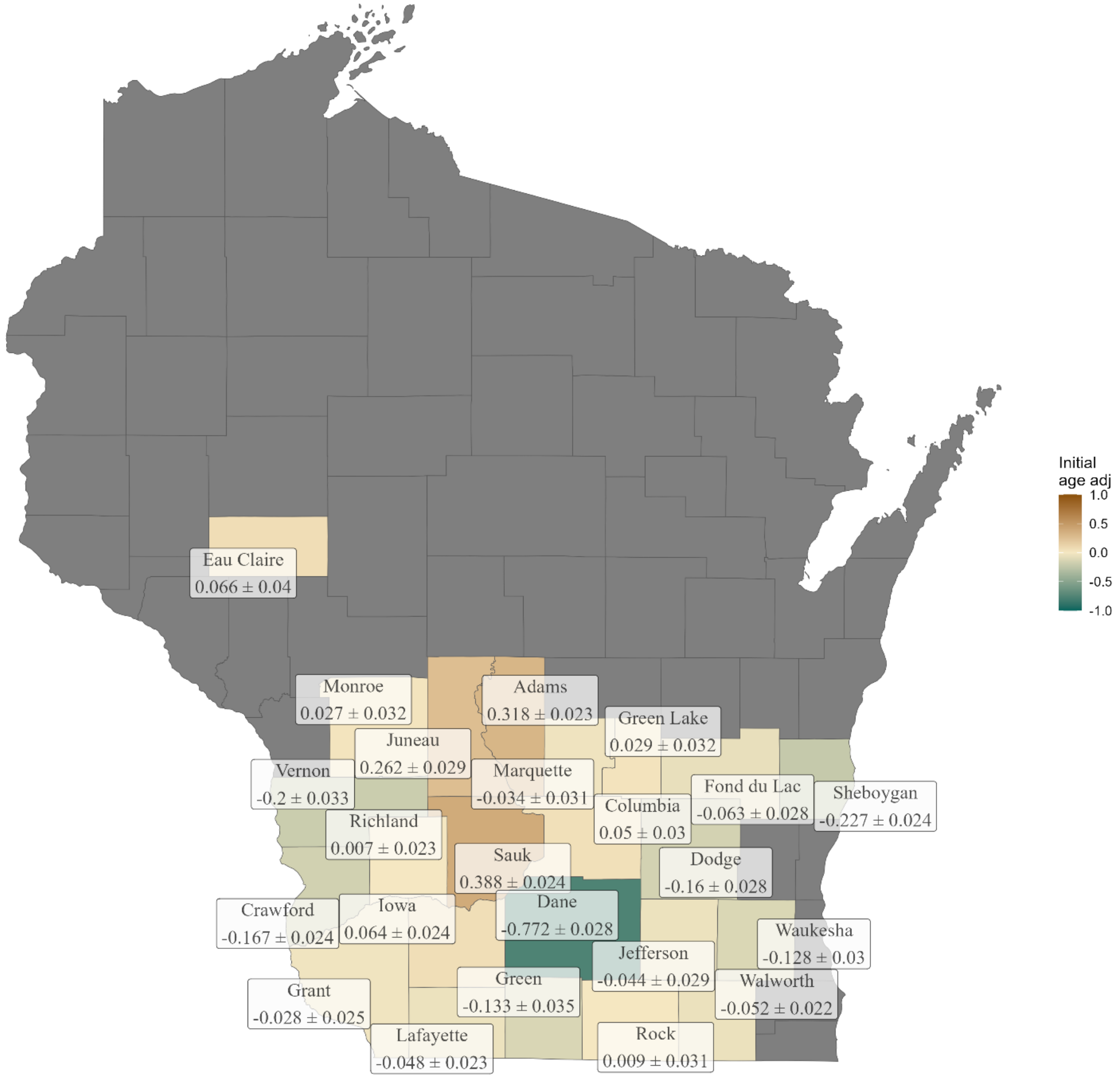


**Figure S3. County specific mean and ± 1 standard deviation for region-specific adjustment of the initial age structure of the deer population estimated from the susceptible-infected-clinical model of chronic wasting disease in white-tailed deer in Wisconsin, fit to the full dataset.** A value of zero indicates that a county's initial age structure matches the global mean dictated by age specific survival equilibrium; values above and below indicate initial age structure is older and younger, respectively, than the global mean. This affects only model initialization, and therefore has little influence after 1 or 2 generations of deer turnover.

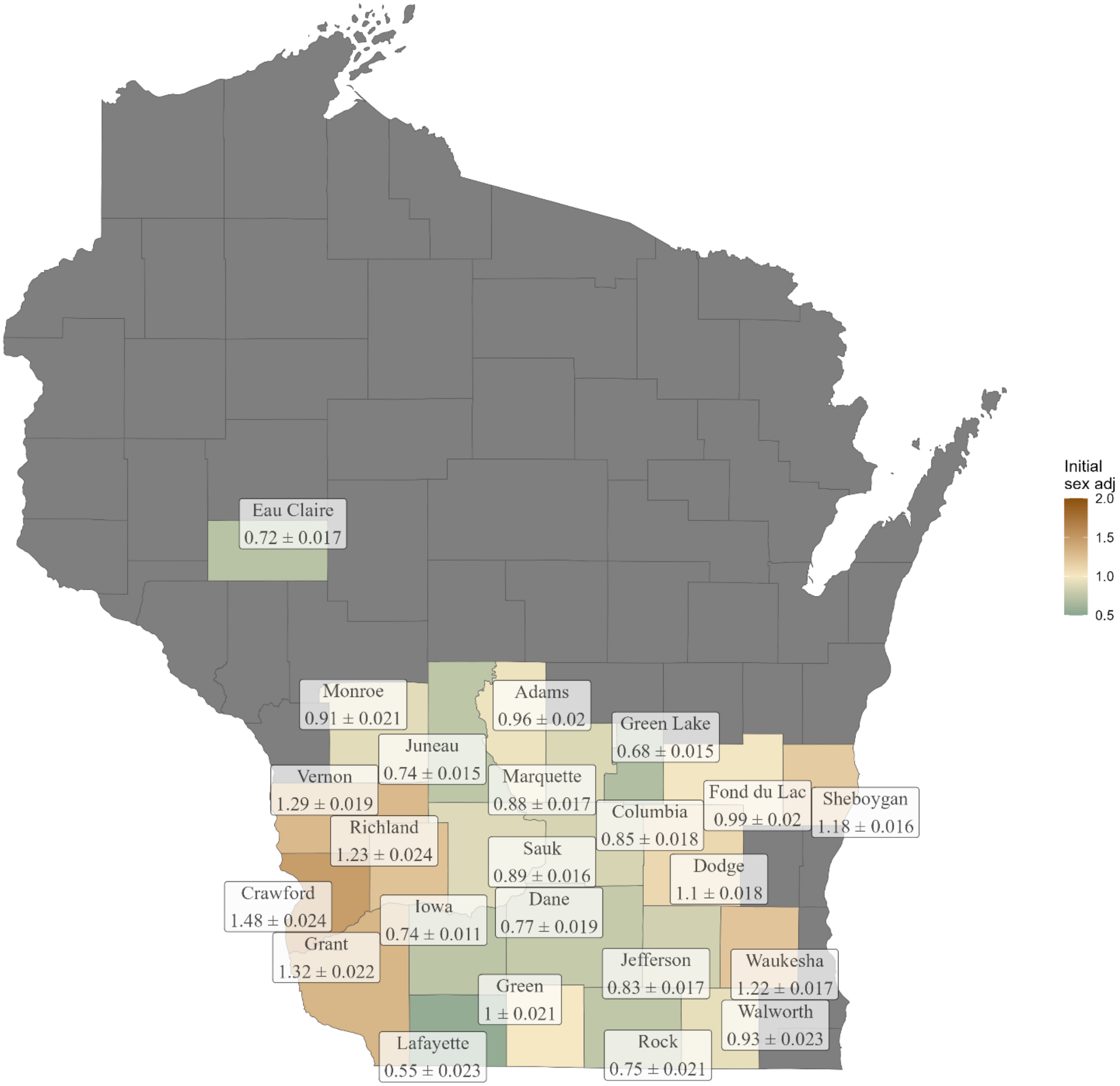


**Figure S4. County specific mean and ± 1 standard deviation for region-specific adjustment of the initial sex ratio of the deer population estimated from the susceptible-infected-clinical model of chronic wasting disease in white-tailed deer in Wisconsin, fit to the full dataset.** A value of one indicates that a county's initial sex ratio matches the global mean dictated by sex specific survival equilibrium; values above and below indicate a county's ratio of bucks is higher and lower respectively than the global mean. This affects only model initialization, and therefore has little influence after 1 or 2 generations of deer turnover.

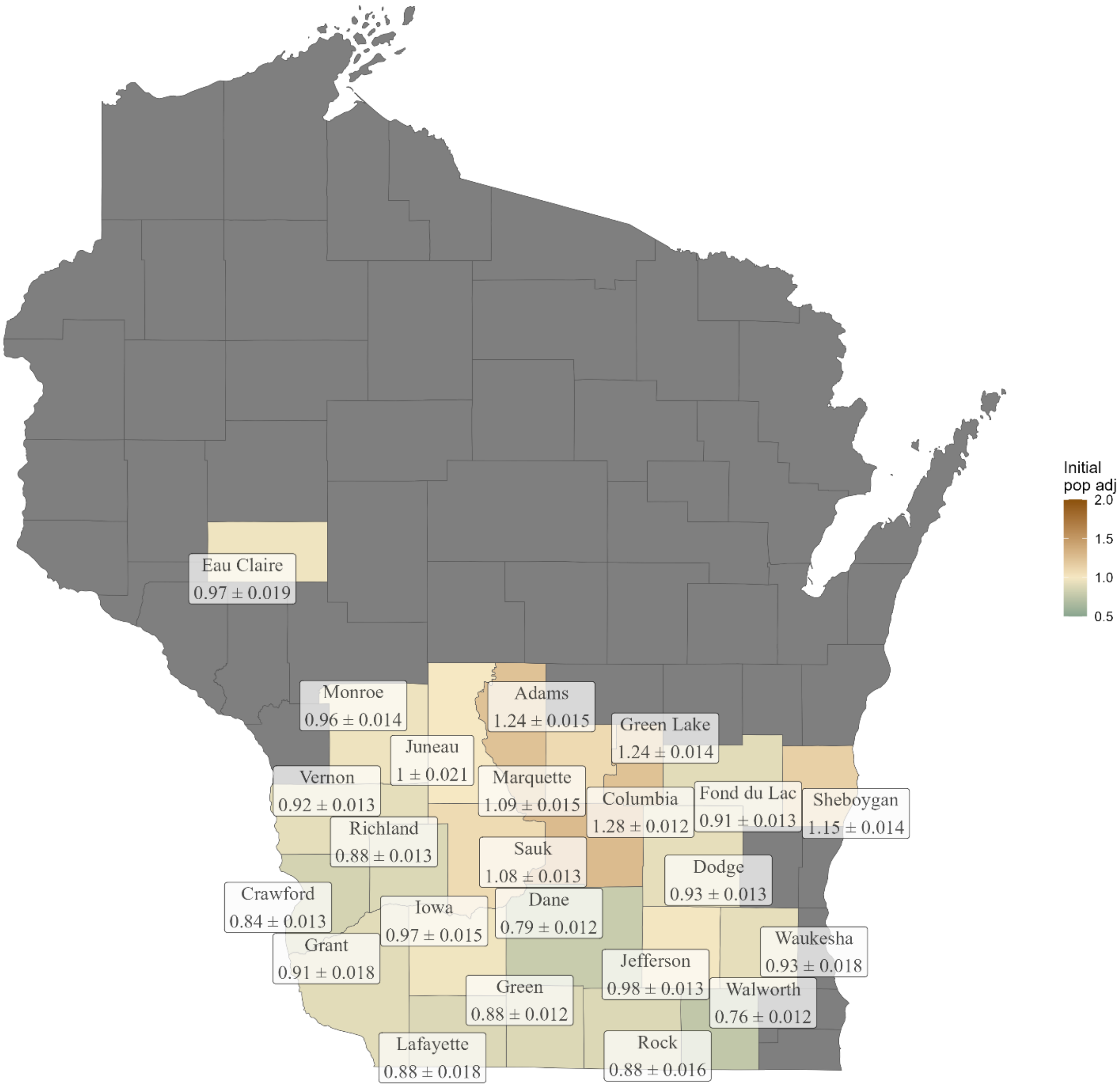


**Figure S5. County specific mean and ± 1 standard deviation for region-specific adjustment of the initial deer abundance estimated from the susceptible-infected-clinical model of chronic wasting disease in white-tailed deer in Wisconsin, fit to the full dataset.** A value of one indicates that a county's initial abundance matches the values indicated by historic data; values above and below indicate a county's initial deer abundance is higher and lower respectively. This affects only model initialization, and therefore has little influence after 1 or 2 generations of deer turnover.

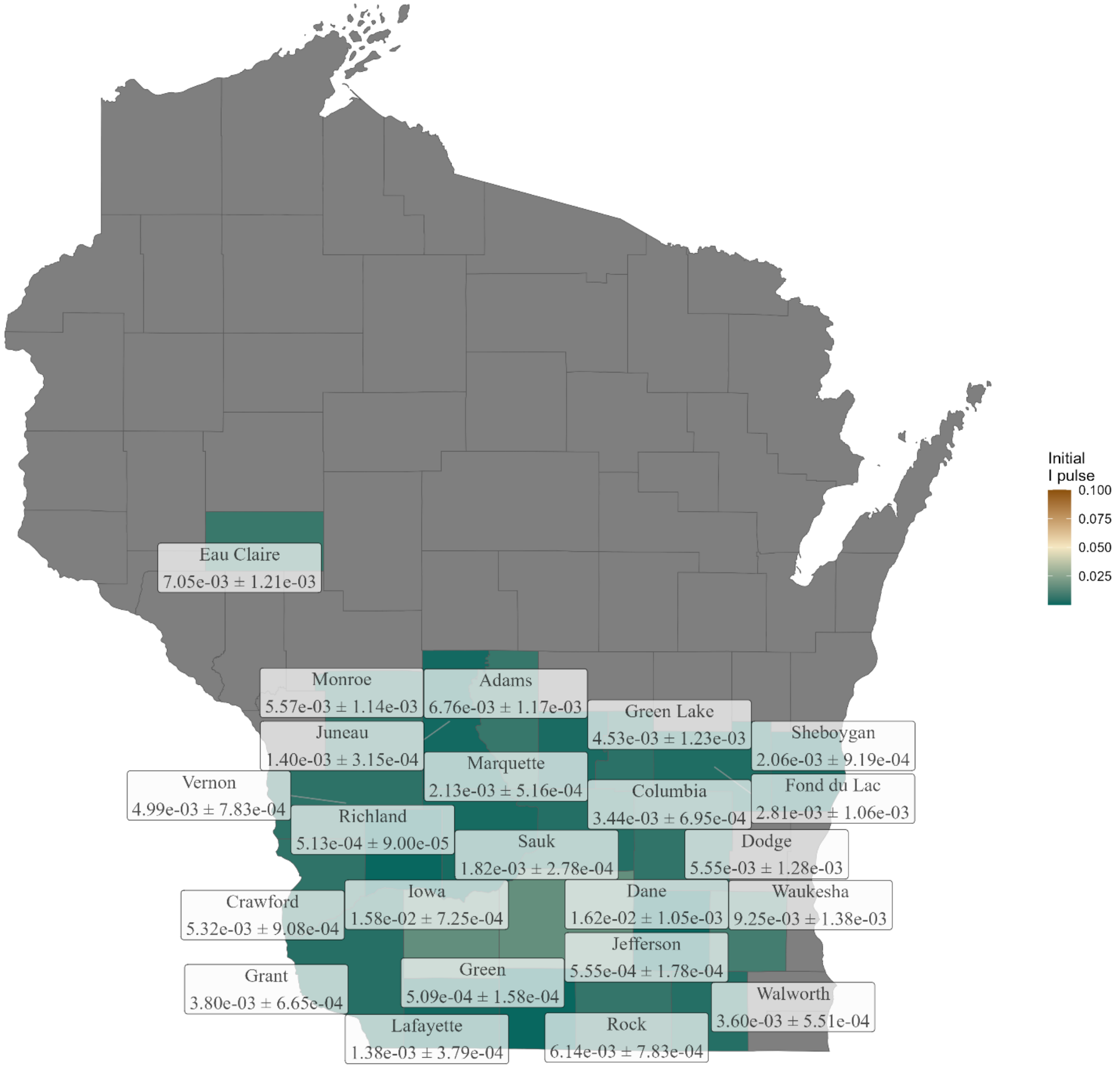


**Figure S6. County specific mean and ± 1 standard deviation for region-specific initial pulse of infected deer estimated from the susceptible-infected-clinical model of chronic wasting disease in white-tailed deer in Wisconsin, fit to the full dataset.** A higher value indicates that a higher fraction of the existing susceptibles in a county are moved to the infected class the year prior to the first detection of chronic wasting disease.

##### *4.1.2.3 Model structure confirmations*

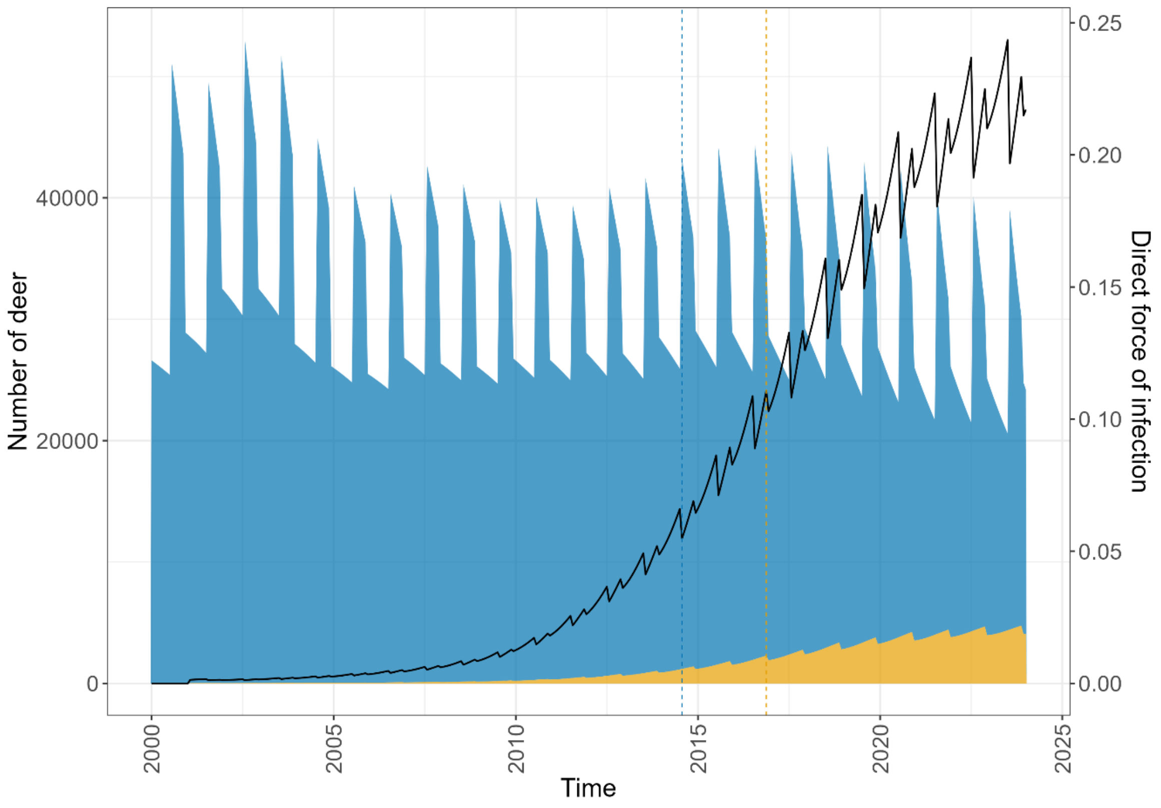


**Figure S7. Estimated force of direct infection for female white-tailed deer aged six years and older (black), given the number of infectious (orange) and healthy (blue) deer present on the landscape during a model simulation in Sauk county, Wisconsin.** This illustrates the modeled impact of the ratio of healthy to infectious population on transmission dynamics, highlighting the disease balancing loop as susceptible deer are depleted and the direct transmission reinforcing loop as infectious deer accumulate. The vertical lines indicate points in time when a decrease in the number of healthy (blue) or infectious (orange) deer on the landscape elicited a drop in the force of infection. Only points where total deer density was between 48-53 deer/$m^2$ are plotted to restrict the influence of density dependence.

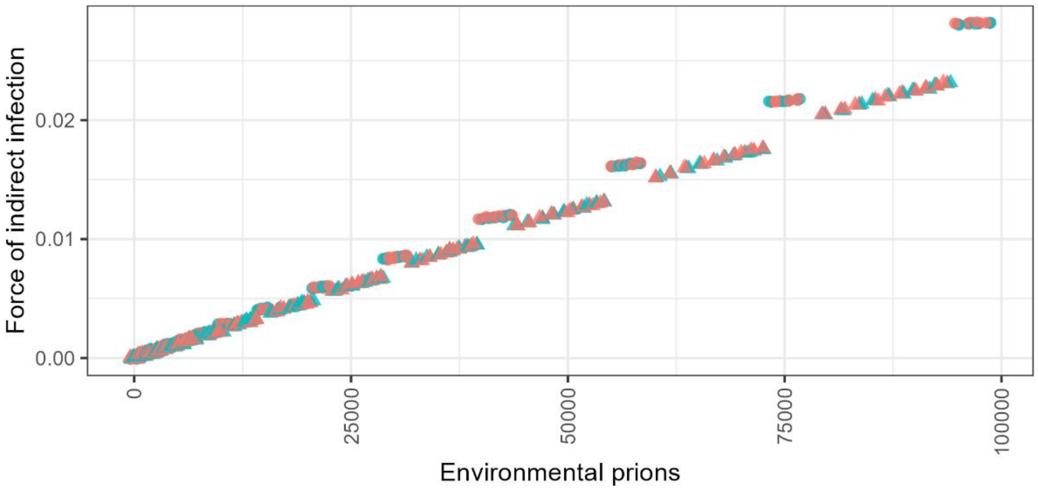


**Figure S8. Estimated force of indirect infection for female (blue) and male (red) white-tailed deer aged six years and older, as a function of the number of infectious environmental prions at low (triangle) and high (circle) numbers of healthy deer present on the landscape during model simulations in Sauk county, Wisconsin.** This illustrates the modelled relationship between environmental contamination and indirect transmission risk, highlighting the disease balancing loop as susceptible deer are depleted and the indirect transmission reinforcing loop as environmental prions accumulate. Low and high numbers of healthy deer range between 24,000-27,000 and 35,000-37,000 deer respectively. Note that indirect transmission hazard is nearly identical across all age and sex cohorts, such that values for males and females overlap and displayed here with a small, horizontal offset (jitter).

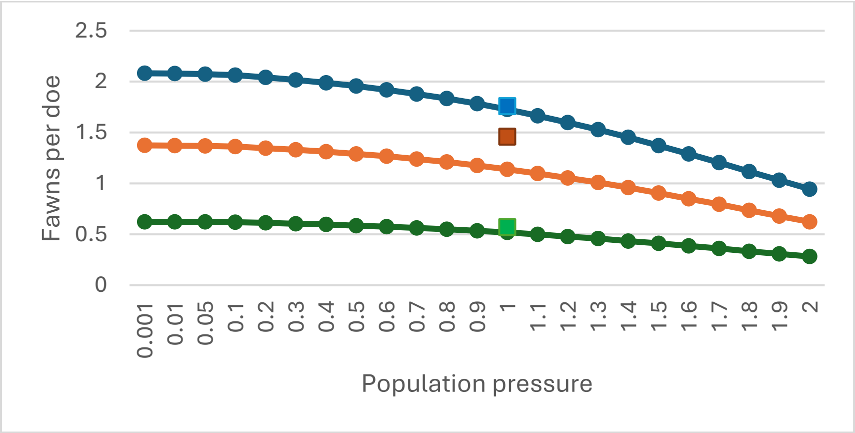


**Figure S9. Relationship between deer birth rates and deer population pressure –a metric of ecological limits on recruitment and mortality, akin to a traditional carrying capacity.** The fawn (green), yearling (orange), and adult (blue) birth rates are shown as the number of fawns born per doe per year, which accounts for pregnancy rates and the number of fetuses. Birth rates for Southern Wisconsin deer range from McCaffery et al. (1998) are shown as square points.

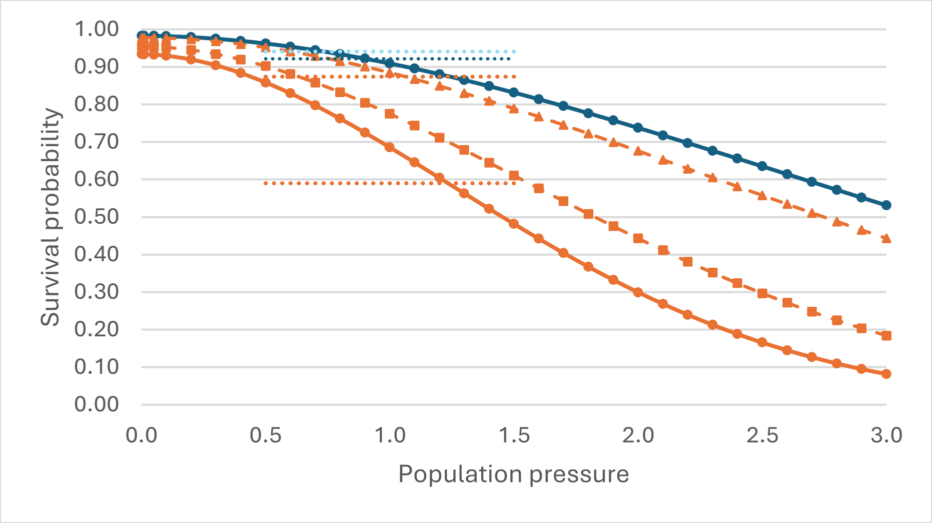


**Figure S10. Background survival probabilities for fawns (orange) and for all yearling and older deer (blue) across a range of deer population pressure –a metric of ecological limits on recruitment and mortality, akin to a traditional carrying capacity.** Survival probabilities are calculated as $e^{(-(background\ mortality\ rate)t)}$ where the background mortality rates are the model estimated parameters and *t* is the number of years survival is calculated over (i.e., *t* =1 for annual survival). Annual survival estimated here for fawns and older deer are depicted with solid lines and circle markers, early fawn mortality (first 12 weeks of life) is depicted by dashed line and square marker, late fawn mortality (13-52 weeks of life) is depicted by dashed line and triangle markers. Rates were calculated in the absence of winter mortality, which caused a negligible decrease across all survival probabilities. Horizontal dotted lines represent a minimum (Rohm et al., 2007) and maximum (Wasserberg et al., 2009) annual values for fawn annual survival probabilities in southern IL and WI respectively, adult females (light blue) and adult males (dark blue) annual survival probabilities (Wasserberg et al., 2009) from the literature.

##### *4.1.2.4 Model predictions*

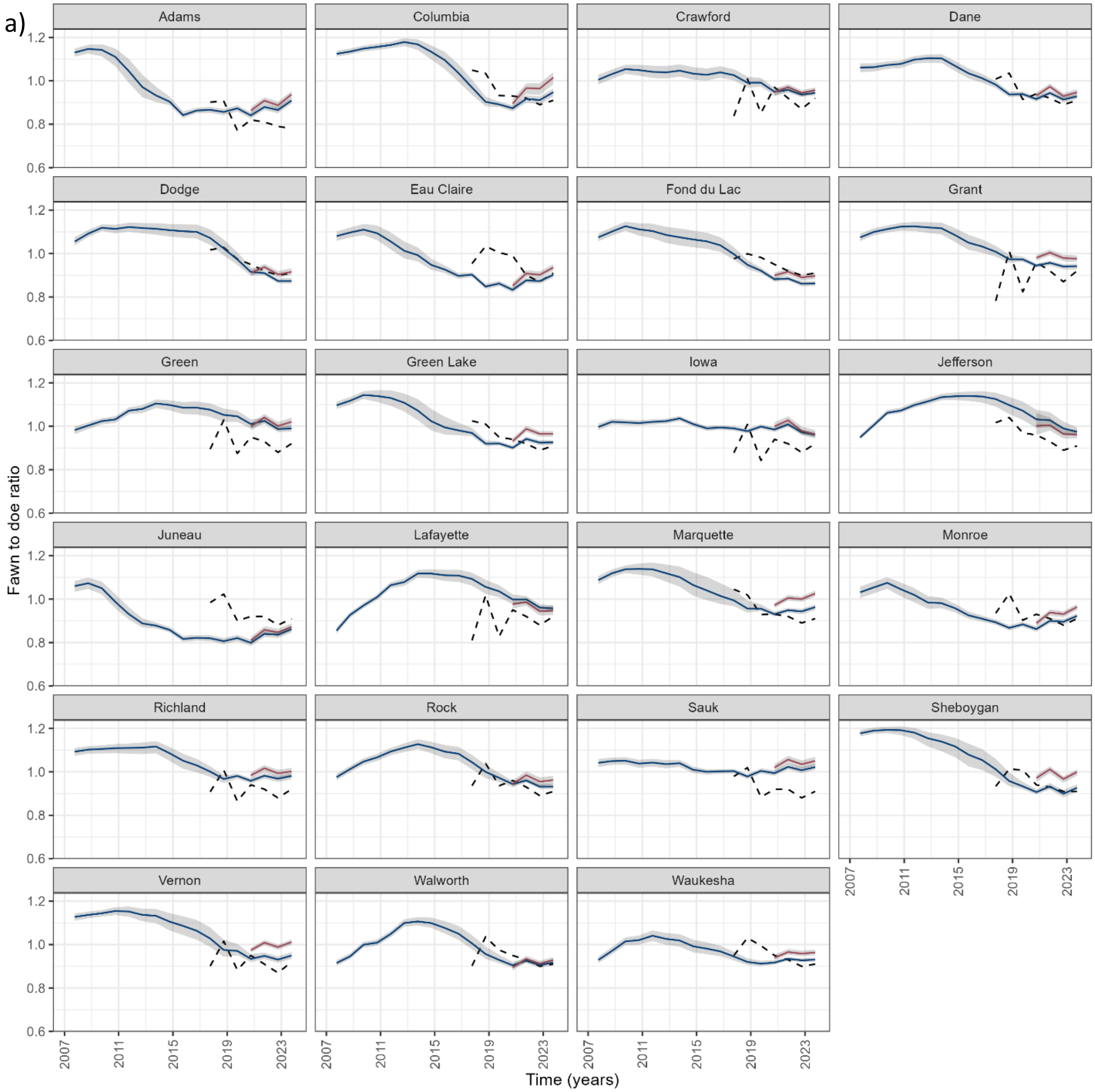


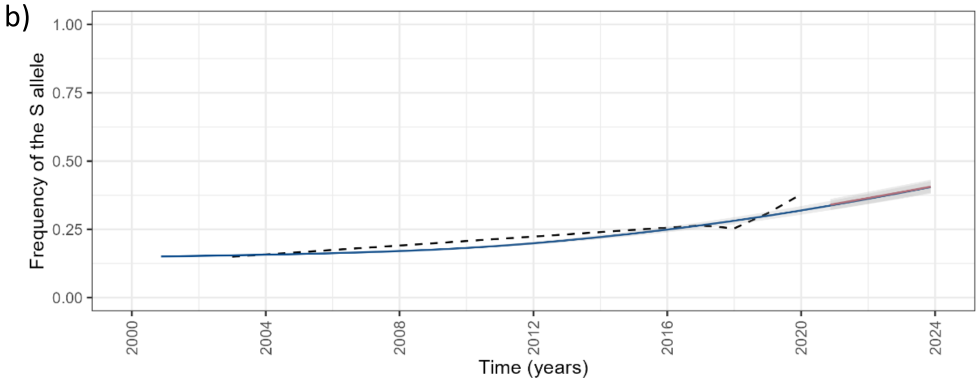

c)

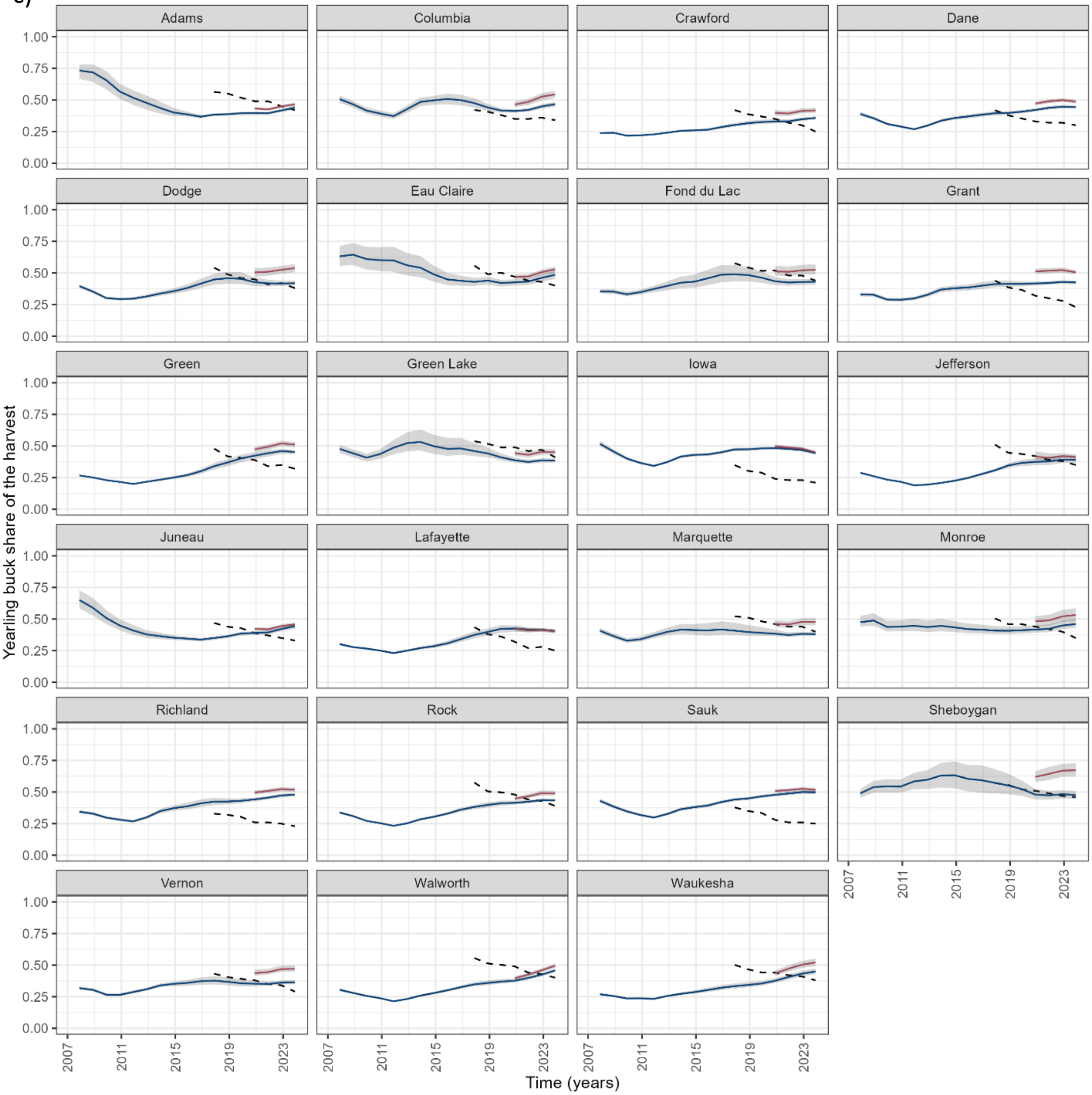

Adams
Columbia
Crawford
Dane
Dodge
Eau Claire
Fond du Lac
Grant
Green
Green Lake
Iowa
Jefferson
Juneau
Lafayette
Marquette
Monroe
Richland
Rock
Sauk
Sheboygan
Vernon
Walworth
Waukesha
Yearling buck share of the harvest
0.00
0.25
0.50
0.75
1.00
2007
2011
2015
2019
2023
Time (years)

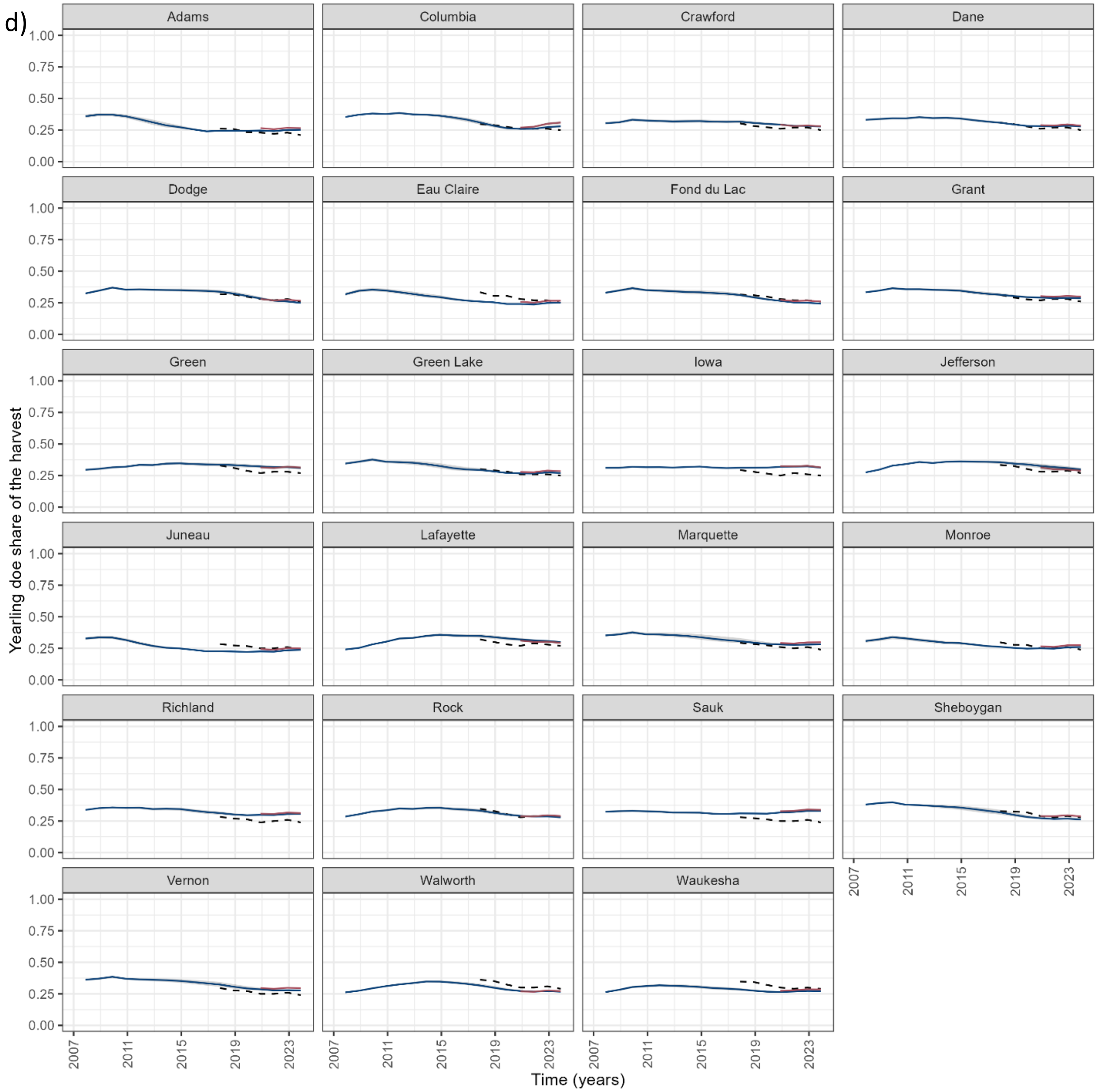


**Figure S11. Model predictions from the susceptible-infected-clinical model of chronic wasting disease (CWD) in white-tailed deer in Wisconsin, fit to the full dataset (blue) and a restricted dataset truncated at 2019 (red), compared against historical time-series data (dashed).** Solid lines represent posterior mean predictions, with 95% credible intervals shaded in grey, calculated across 1,000 parameter sets sampled from the joint posterior to reflect both parameter uncertainty and data measurement error. Panels display: (a) the fawn-to-doe ratio from summer observations, (b) the frequency of the *S* allele in the core CWD area (averaged across Dane, Grant, and Iowa counties), (c) the yearling share of the antlered harvest, and (d) the yearling share of the antlerless harvest. Predictions for the testing data (2020-2024) are presented for the out-of-sample fit, which was trained on data prior to 2020.

a)

b)

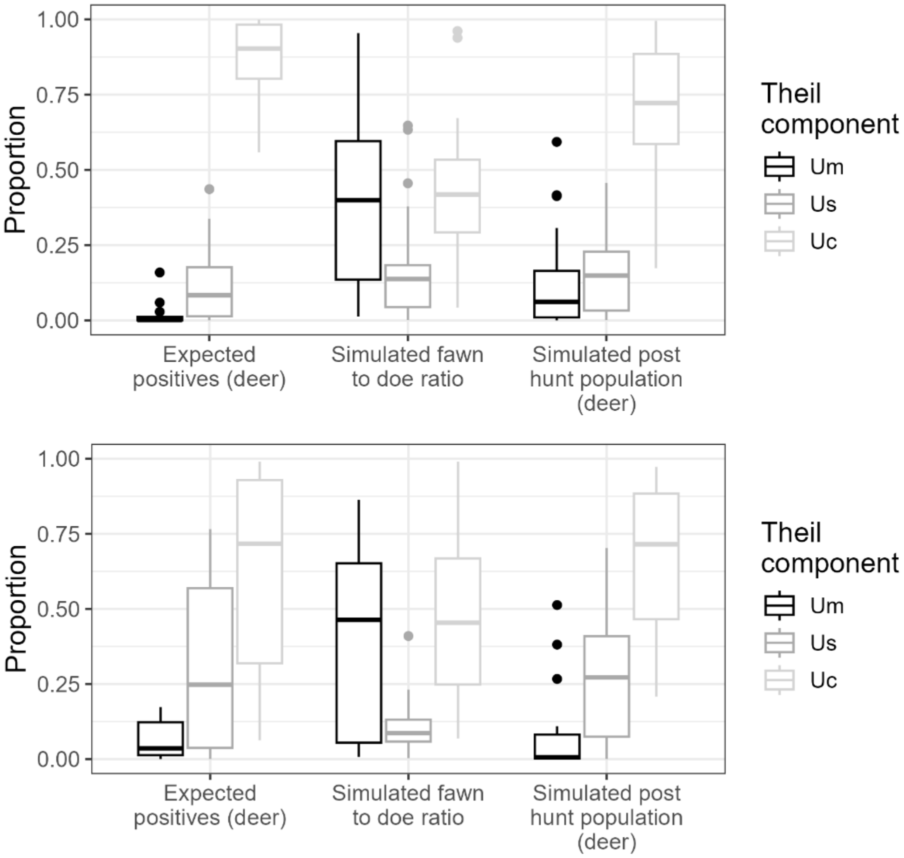


**Figure S12. Theil statistics comparing historical time-series data from the Wisconsin Department of Natural Resources to the simulated estimates from the susceptible-infected-clinical model of chronic wasting disease (CWD) in white-tailed deer fit to (a) the full dataset and (b) a restricted dataset truncated at 2019.** Theil statistics decompose the total error between model predictions and observed data into components attributed to: (Um) systematic bias in the mean, (Us) differences in variance, and (Uc) unexplained error (random noise). The three components sum to one. Boxplots represent Theil statistics across regions for three key metrics: the total number of expected positive CWD tests, the fawn-to-doe ratio, and the post-hunt deer population. Boxes denote the interquartile range (IQR), the line indicates the median, whiskers extend to 1.5×IQR, and points represent outliers. Due to problematic data, we expect more systematic variation (Um, Us) in population metrics, which we observe.

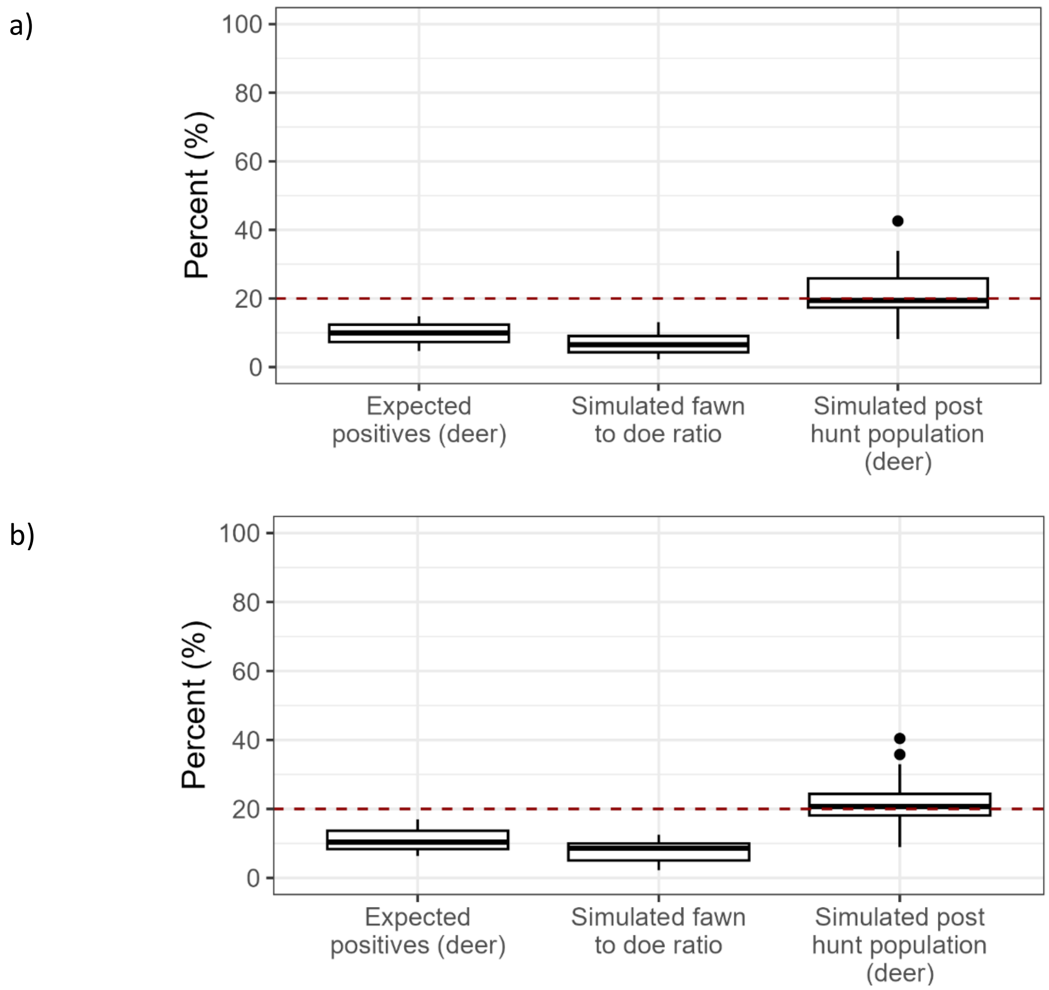


**Figure S13. Mean absolute percentage error (MAPE) comparing historical time-series data from the Wisconsin Department of Natural Resources to the simulated estimates from the susceptible-infected-clinical model of chronic wasting disease (CWD) in white-tailed deer fit to (a) the full dataset and (b) a restricted dataset truncated at 2019.** MAPE is a unitless measure of model accuracy, representing the proportion of variance in the data explained by the model. In the context of large-scale system dynamics models, MAPE values below 20% and 50% are typically considered indicative of good and reasonable forecasting performance, respectively. Boxplots display MAPE across regions for three metrics: the total number of expected positive CWD tests, the fawn-to-doe ratio, and the post-hunt total deer population. Boxes show the interquartile range, with the median as a central line, whiskers extending to 1.5×IQR, and outliers shown as points.

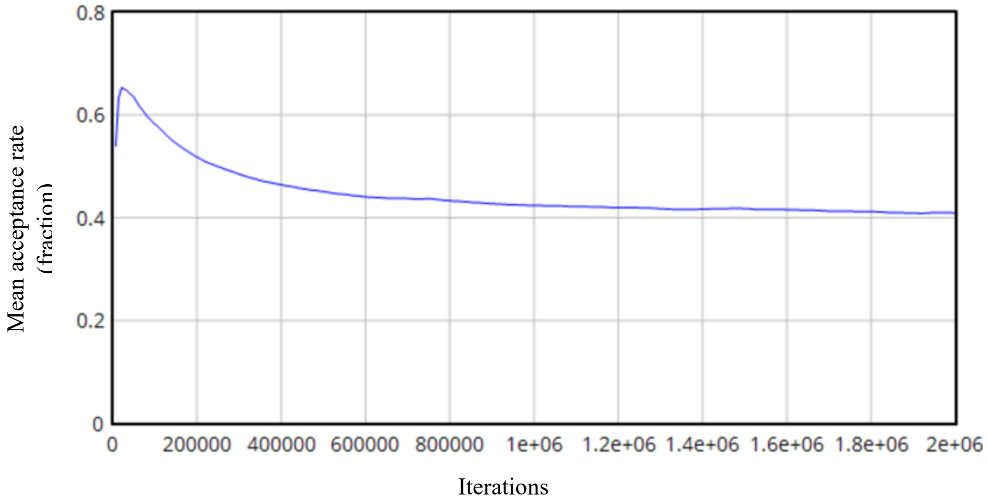


**Figure S14. Mean acceptance rate across Markov chain Monte Carlo chains (reflected as a proportion) from the posterior distribution of the disaggregated susceptible-infected-clinical model of chronic wasting disease in white-tailed deer in Wisconsin**. Acceptance rates reflect the efficiency of the sampling process and indicate good convergence when rates are neither too low (indicating poor mixing) nor too high (indicating overly conservative proposal steps).

a)

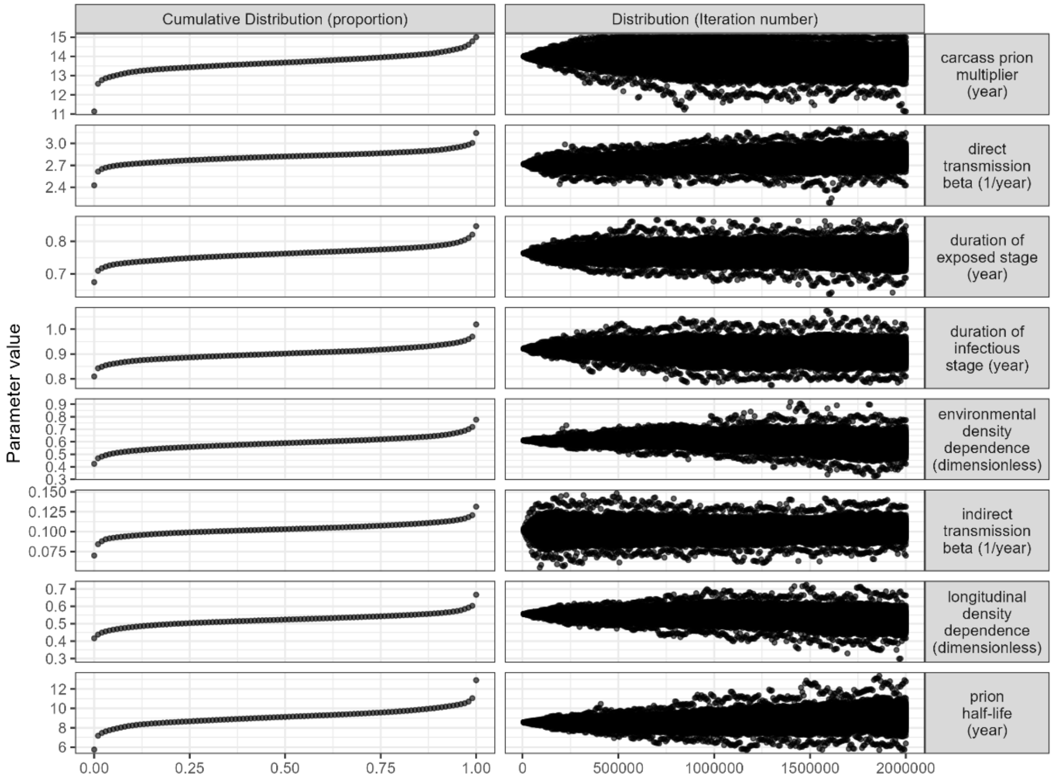

b)

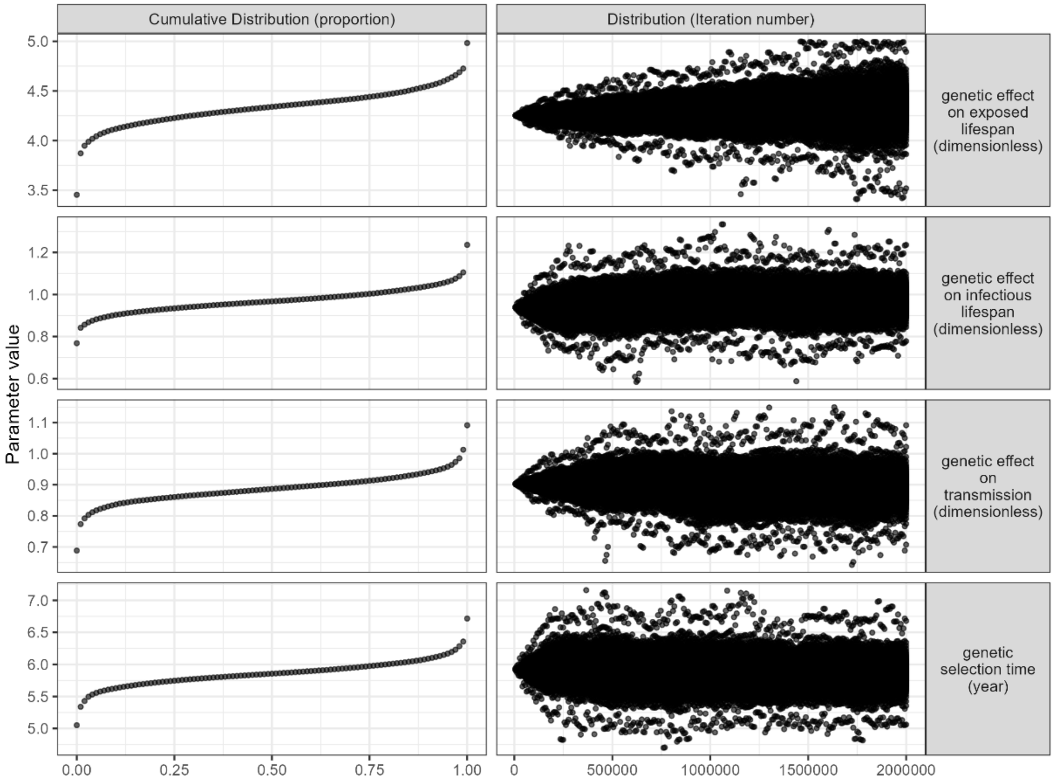
Cumulative Distribution (proportion)
Distribution (Iteration number)
Parameter value
genetic effect on exposed lifespan (dimensionless)
genetic effect on infectious lifespan (dimensionless)
genetic effect on transmission (dimensionless)
genetic selection time (year)
0.00
0.25
0.50
0.75
1.00
0
500000
1000000
1500000
2000000

c)

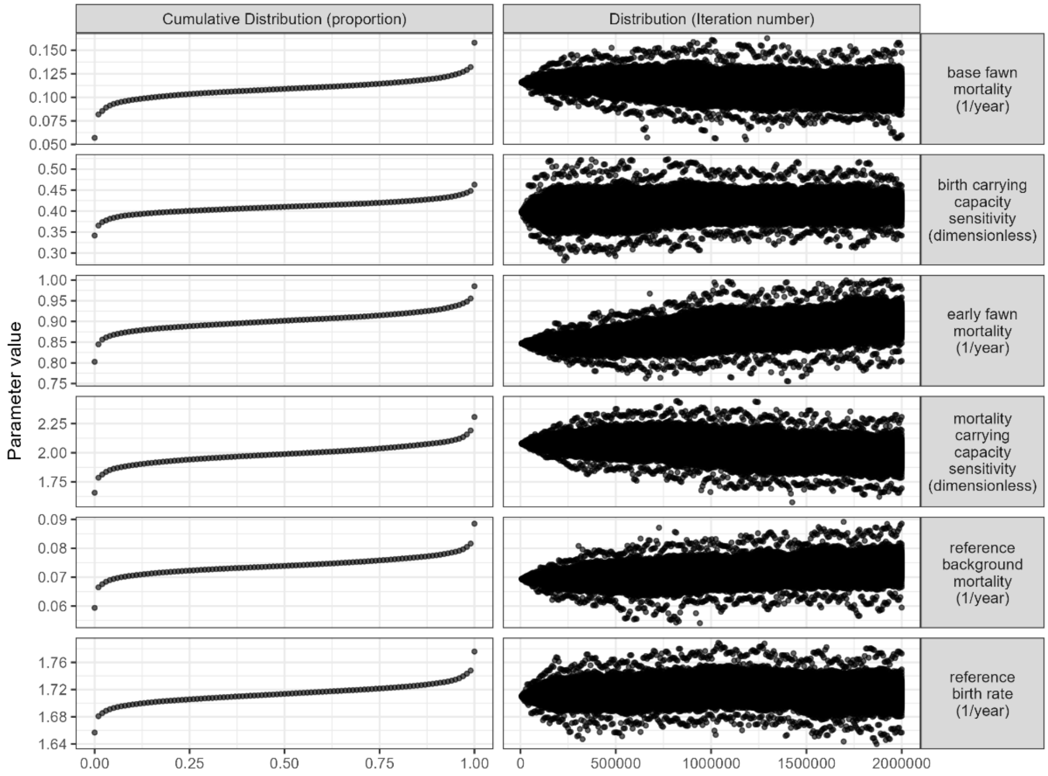
Cumulative Distribution (proportion)
Distribution (Iteration number)
Parameter value
base fawn mortality (1/year)
birth carrying capacity sensitivity (dimensionless)
early fawn mortality (1/year)
mortality carrying capacity sensitivity (dimensionless)
reference background mortality (1/year)
reference birth rate (1/year)
0.00
0.25
0.50
0.75
1.00
0
500000
1000000
1500000
2000000

**Figure S15. Posterior parameter distributions across Markov chain Monte Carlo (MCMC) chains from the posterior distribution of the disaggregated susceptible-infected-clinical model of chronic wasting disease in white-tailed deer in Wisconsin**. The left column shows cumulative distributions, which summarize the range and relative probability of parameter values across all retained samples. The right column shows trace plots for parameter values by each iteration, illustrating mixing and convergence of the chains as well as variability across draws. Together, these views provide complementary perspectives on both the stationary distributions of parameters and the sampling behavior of the MCMC algorithm. The first panel depicts key disease parameters (a), the second panel contains parameters specific to genetics (b) and demographic parameters are given in the third (c).

##### 4.1.3 Sensitivity analyses

###### *4.1.3.1 Local parameter sensitivity*

a)

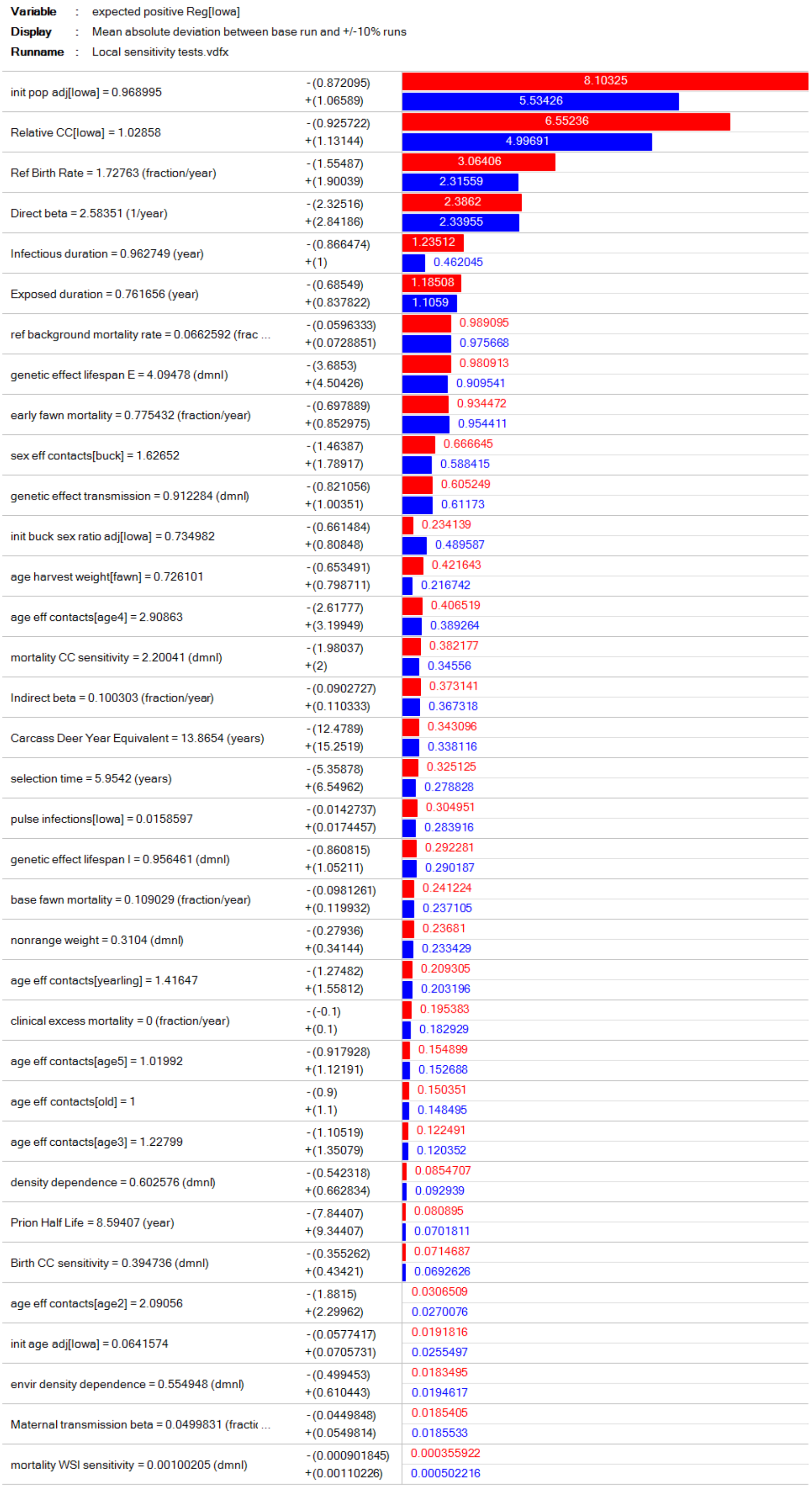
Variable : expected positive Reg[Iowa]
Display : Mean absolute deviation between base run and +/-10% runs
Runname : Local sensitivity tests.vdfx
init pop adj[Iowa] = 0.968995
-(0.872095)
+(1.06589)
8.10325
5.53426
Relative CC[Iowa] = 1.02858
-(0.925722)
+(1.13144)
6.55236
4.99691
Ref Birth Rate = 1.72763 (fraction/year)
-(1.55487)
+(1.90039)
3.06406
2.31559
Direct beta = 2.58351 (1/year)
-(2.32516)
+(2.84186)
2.3862
2.33955
Infectious duration = 0.962749 (year)
-(0.866474)
+(1)
1.23512
0.462045
Exposed duration = 0.761656 (year)
-(0.68549)
+(0.837822)
1.18508
1.1059
ref background mortality rate = 0.0662592 (frac ...
-(0.0596333)
+(0.0728851)
0.989095
0.975668
genetic effect lifespan E = 4.09478 (dmnl)
-(3.6853)
+(4.50426)
0.980913
0.909541
early fawn mortality = 0.775432 (fraction/year)
-(0.697889)
+(0.852975)
0.934472
0.954411
sex eff contacts[buck] = 1.62652
-(1.46387)
+(1.78917)
0.666645
0.588415
genetic effect transmission = 0.912284 (dmnl)
-(0.821056)
+(1.00351)
0.605249
0.61173
init buck sex ratio adj[Iowa] = 0.734982
-(0.661484)
+(0.80848)
0.234139
0.489587
age harvest weight[fawn] = 0.726101
-(0.653491)
+(0.798711)
0.421643
0.216742
age eff contacts[age4] = 2.90863
-(2.61777)
+(3.19949)
0.406519
0.389264
mortality CC sensitivity = 2.20041 (dmnl)
-(1.98037)
+(2)
0.382177
0.34556
Indirect beta = 0.100303 (fraction/year)
-(0.0902727)
+(0.110333)
0.373141
0.367318
Carcass Deer Year Equivalent = 13.8654 (years)
-(12.4789)
+(15.2519)
0.343096
0.338116
selection time = 5.9542 (years)
-(5.35878)
+(6.54962)
0.325125
0.278828
pulse infections[Iowa] = 0.0158597
-(0.0142737)
+(0.0174457)
0.304951
0.283916
genetic effect lifespan I = 0.956461 (dmnl)
-(0.860815)
+(1.05211)
0.292281
0.290187
base fawn mortality = 0.109029 (fraction/year)
-(0.0981261)
+(0.119932)
0.241224
0.237105
nonrange weight = 0.3104 (dmnl)
-(0.27936)
+(0.34144)
0.23681
0.233429
age eff contacts[yearling] = 1.41647
-(1.27482)
+(1.55812)
0.209305
0.203196
clinical excess mortality = 0 (fraction/year)
-(-0.1)
+(0.1)
0.195383
0.182929
age eff contacts[age5] = 1.01992
-(0.917928)
+(1.12191)
0.154899
0.152688
age eff contacts[old] = 1
-(0.9)
+(1.1)
0.150351
0.148495
age eff contacts[age3] = 1.22799
-(1.10519)
+(1.35079)
0.122491
0.120352
density dependence = 0.602576 (dmnl)
-(0.542318)
+(0.662834)
0.0854707
0.092939
Prion Half Life = 8.59407 (year)
-(7.84407)
+(9.34407)
0.080895
0.0701811
Birth CC sensitivity = 0.394736 (dmnl)
-(0.355262)
+(0.43421)
0.0714687
0.0692626
age eff contacts[age2] = 2.09056
-(1.8815)
+(2.29962)
0.0306509
0.0270076
init age adj[Iowa] = 0.0641574
-(0.0577417)
+(0.0705731)
0.0191816
0.0255497
envir density dependence = 0.554948 (dmnl)
-(0.499453)
+(0.610443)
0.0183495
0.0194617
Maternal transmission beta = 0.0499831 (fractic ...
-(0.0449848)
+(0.0549814)
0.0185405
0.0185533
mortality WSI sensitivity = 0.00100205 (dmnl)
-(0.000901845)
+(0.00110226)
0.000355922
0.000502216

b)

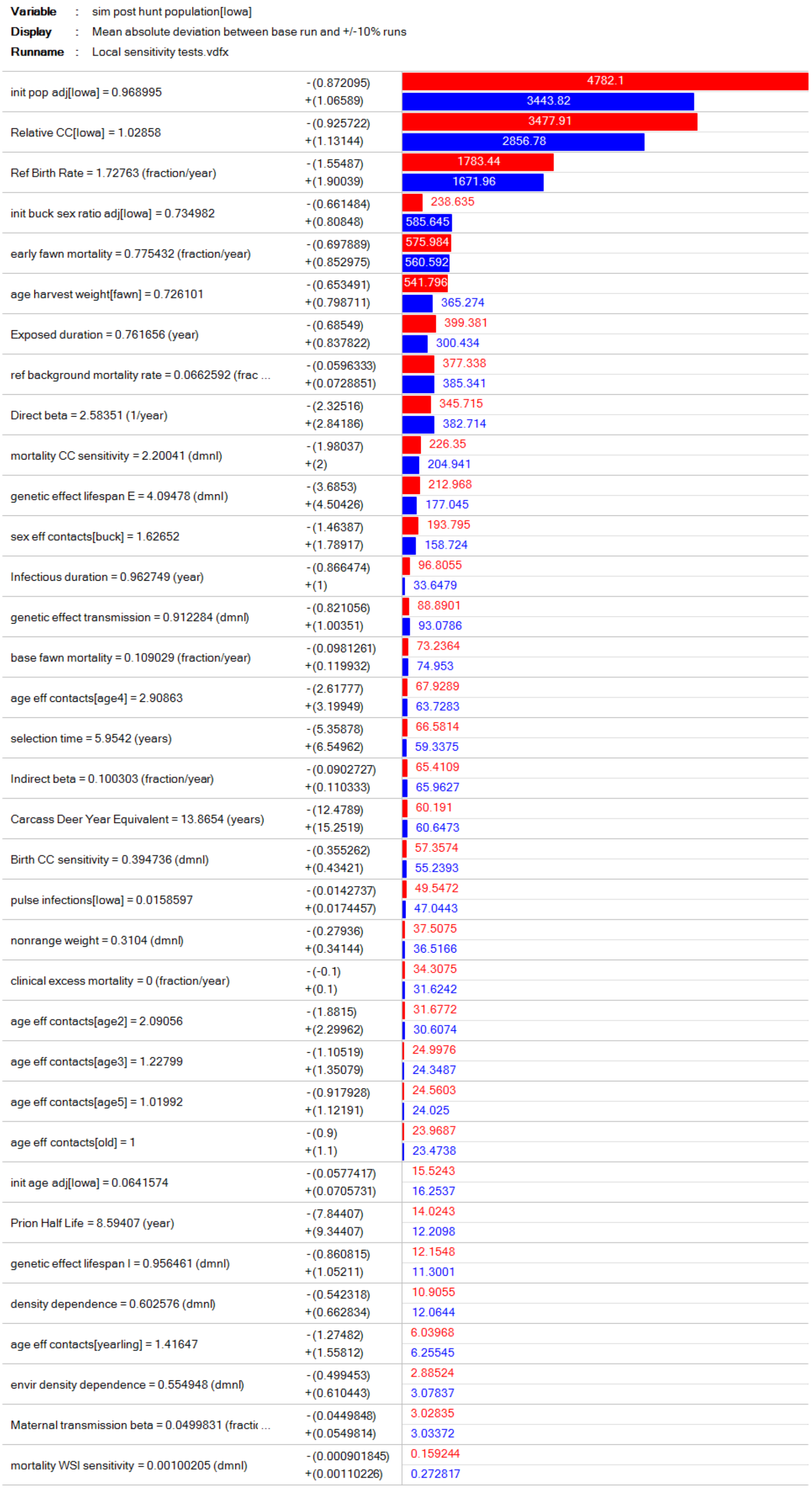
Variable : sim post hunt population[Iowa]
Display : Mean absolute deviation between base run and +/-10% runs
Runname : Local sensitivity tests.vdfx
init pop adj[Iowa] = 0.968995
-(0.872095)
+(1.06589)
4782.1
3443.82
Relative CC[Iowa] = 1.02858
-(0.925722)
+(1.13144)
3477.91
2856.78
Ref Birth Rate = 1.72763 (fraction/year)
-(1.55487)
+(1.90039)
1783.44
1671.96
init buck sex ratio adj[Iowa] = 0.734982
-(0.661484)
+(0.80848)
238.635
585.645
early fawn mortality = 0.775432 (fraction/year)
-(0.697889)
+(0.852975)
575.984
560.592
age harvest weight[fawn] = 0.726101
-(0.653491)
+(0.798711)
541.796
365.274
Exposed duration = 0.761656 (year)
-(0.68549)
+(0.837822)
399.381
300.434
ref background mortality rate = 0.0662592 (frac ...
-(0.0596333)
+(0.0728851)
377.338
385.341
Direct beta = 2.58351 (1/year)
-(2.32516)
+(2.84186)
345.715
382.714
mortality CC sensitivity = 2.20041 (dmnl)
-(1.98037)
+(2)
226.35
204.941
genetic effect lifespan E = 4.09478 (dmnl)
-(3.6853)
+(4.50426)
212.968
177.045
sex eff contacts[buck] = 1.62652
-(1.46387)
+(1.78917)
193.795
158.724
Infectious duration = 0.962749 (year)
-(0.866474)
+(1)
96.8055
33.6479
genetic effect transmission = 0.912284 (dmnl)
-(0.821056)
+(1.00351)
88.8901
93.0786
base fawn mortality = 0.109029 (fraction/year)
-(0.0981261)
+(0.119932)
73.2364
74.953
age eff contacts[age4] = 2.90863
-(2.61777)
+(3.19949)
67.9289
63.7283
selection time = 5.9542 (years)
-(5.35878)
+(6.54962)
66.5814
59.3375
Indirect beta = 0.100303 (fraction/year)
-(0.0902727)
+(0.110333)
65.4109
65.9627
Carcass Deer Year Equivalent = 13.8654 (years)
-(12.4789)
+(15.2519)
60.191
60.6473
Birth CC sensitivity = 0.394736 (dmnl)
-(0.355262)
+(0.43421)
57.3574
55.2393
pulse infections[Iowa] = 0.0158597
-(0.0142737)
+(0.0174457)
49.5472
47.0443
nonrange weight = 0.3104 (dmnl)
-(0.27936)
+(0.34144)
37.5075
36.5166
clinical excess mortality = 0 (fraction/year)
-(-0.1)
+(0.1)
34.3075
31.6242
age eff contacts[age2] = 2.09056
-(1.8815)
+(2.29962)
31.6772
30.6074
age eff contacts[age3] = 1.22799
-(1.10519)
+(1.35079)
24.9976
24.3487
age eff contacts[age5] = 1.01992
-(0.917928)
+(1.12191)
24.5603
24.025
age eff contacts[old] = 1
-(0.9)
+(1.1)
23.9687
23.4738
init age adj[Iowa] = 0.0641574
-(0.0577417)
+(0.0705731)
15.5243
16.2537
Prion Half Life = 8.59407 (year)
-(7.84407)
+(9.34407)
14.0243
12.2098
genetic effect lifespan I = 0.956461 (dmnl)
-(0.860815)
+(1.05211)
12.1548
11.3001
density dependence = 0.602576 (dmnl)
-(0.542318)
+(0.662834)
10.9055
12.0644
age eff contacts[yearling] = 1.41647
-(1.27482)
+(1.55812)
6.03968
6.25545
envir density dependence = 0.554948 (dmnl)
-(0.499453)
+(0.610443)
2.88524
3.07837
Maternal transmission beta = 0.0499831 (fractio ...
-(0.0449848)
+(0.0549814)
3.02835
3.03372
mortality WSI sensitivity = 0.00100205 (dmnl)
-(0.000901845)
+(0.00110226)
0.159244
0.272817

c)

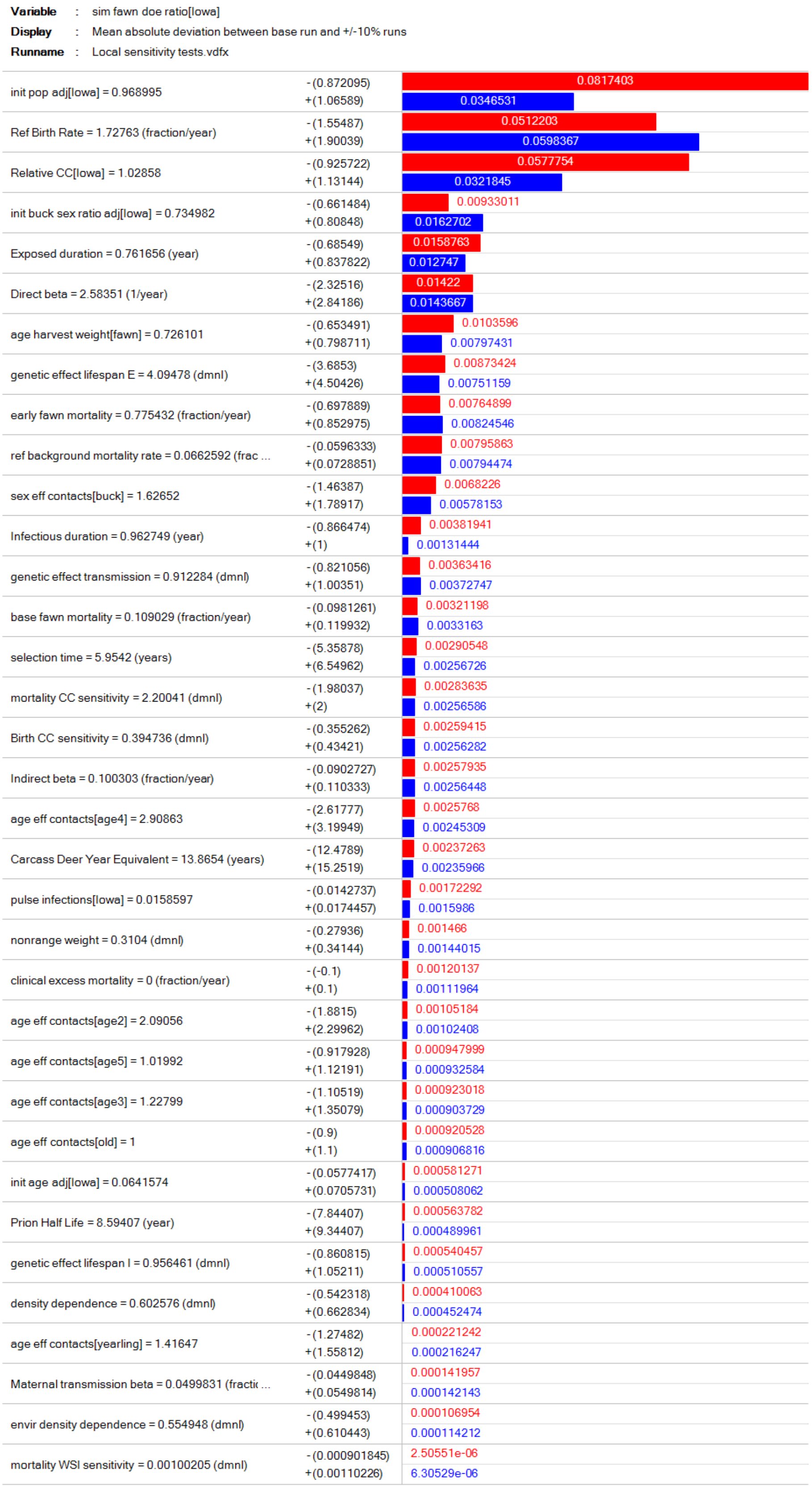

Variable : sim fawn doe ratio[Iowa]
Display : Mean absolute deviation between base run and +/-10% runs
Runname : Local sensitivity tests.vdfx
init pop adj[Iowa] = 0.968995
-(0.872095)
+(1.06589)
0.0817403
0.0346531
Ref Birth Rate = 1.72763 (fraction/year)
-(1.55487)
+(1.90039)
0.0512203
0.0598367
Relative CC[Iowa] = 1.02858
-(0.925722)
+(1.13144)
0.0577754
0.0321845
init buck sex ratio adj[Iowa] = 0.734982
-(0.661484)
+(0.80848)
0.00933011
0.0162702
Exposed duration = 0.761656 (year)
-(0.68549)
+(0.837822)
0.0158763
0.012747
Direct beta = 2.58351 (1/year)
-(2.32516)
+(2.84186)
0.01422
0.0143667
age harvest weight[fawn] = 0.726101
-(0.653491)
+(0.798711)
0.0103596
0.00797431
genetic effect lifespan E = 4.09478 (dmnl)
-(3.6853)
+(4.50426)
0.00873424
0.00751159
early fawn mortality = 0.775432 (fraction/year)
-(0.697889)
+(0.852975)
0.00764899
0.00824546
ref background mortality rate = 0.0662592 (frac ...
-(0.0596333)
+(0.0728851)
0.00795863
0.00794474
sex eff contacts[buck] = 1.62652
-(1.46387)
+(1.78917)
0.0068226
0.00578153
Infectious duration = 0.962749 (year)
-(0.866474)
+(1)
0.00381941
0.00131444
genetic effect transmission = 0.912284 (dmnl)
-(0.821056)
+(1.00351)
0.00363416
0.00372747
base fawn mortality = 0.109029 (fraction/year)
-(0.0981261)
+(0.119932)
0.00321198
0.0033163
selection time = 5.9542 (years)
-(5.35878)
+(6.54962)
0.00290548
0.00256726
mortality CC sensitivity = 2.20041 (dmnl)
-(1.98037)
+(2)
0.00283635
0.00256586
Birth CC sensitivity = 0.394736 (dmnl)
-(0.355262)
+(0.43421)
0.00259415
0.00256282
Indirect beta = 0.100303 (fraction/year)
-(0.0902727)
+(0.110333)
0.00257935
0.00256448
age eff contacts[age4] = 2.90863
-(2.61777)
+(3.19949)
0.0025768
0.00245309
Carcass Deer Year Equivalent = 13.8654 (years)
-(12.4789)
+(15.2519)
0.00237263
0.00235966
pulse infections[Iowa] = 0.0158597
-(0.0142737)
+(0.0174457)
0.00172292
0.0015986
nonrange weight = 0.3104 (dmnl)
-(0.27936)
+(0.34144)
0.001466
0.00144015
clinical excess mortality = 0 (fraction/year)
-(-0.1)
+(0.1)
0.00120137
0.00111964
age eff contacts[age2] = 2.09056
-(1.8815)
+(2.29962)
0.00105184
0.00102408
age eff contacts[age5] = 1.01992
-(0.917928)
+(1.12191)
0.000947999
0.000932584
age eff contacts[age3] = 1.22799
-(1.10519)
+(1.35079)
0.000923018
0.000903729
age eff contacts[old] = 1
-(0.9)
+(1.1)
0.000920528
0.000906816
init age adj[Iowa] = 0.0641574
-(0.0577417)
+(0.0705731)
0.000581271
0.000508062
Prion Half Life = 8.59407 (year)
-(7.84407)
+(9.34407)
0.000563782
0.000489961
genetic effect lifespan I = 0.956461 (dmnl)
-(0.860815)
+(1.05211)
0.000540457
0.000510557
density dependence = 0.602576 (dmnl)
-(0.542318)
+(0.662834)
0.000410063
0.000452474
age eff contacts[yearling] = 1.41647
-(1.27482)
+(1.55812)
0.000221242
0.000216247
Maternal transmission beta = 0.0499831 (fractic ...
-(0.0449848)
+(0.0549814)
0.000141957
0.000142143
envir density dependence = 0.554948 (dmnl)
-(0.499453)
+(0.610443)
0.000106954
0.000114212
mortality WSI sensitivity = 0.00100205 (dmnl)
-(0.000901845)
+(0.00110226)
2.50551e-06
6.30529e-06

d)

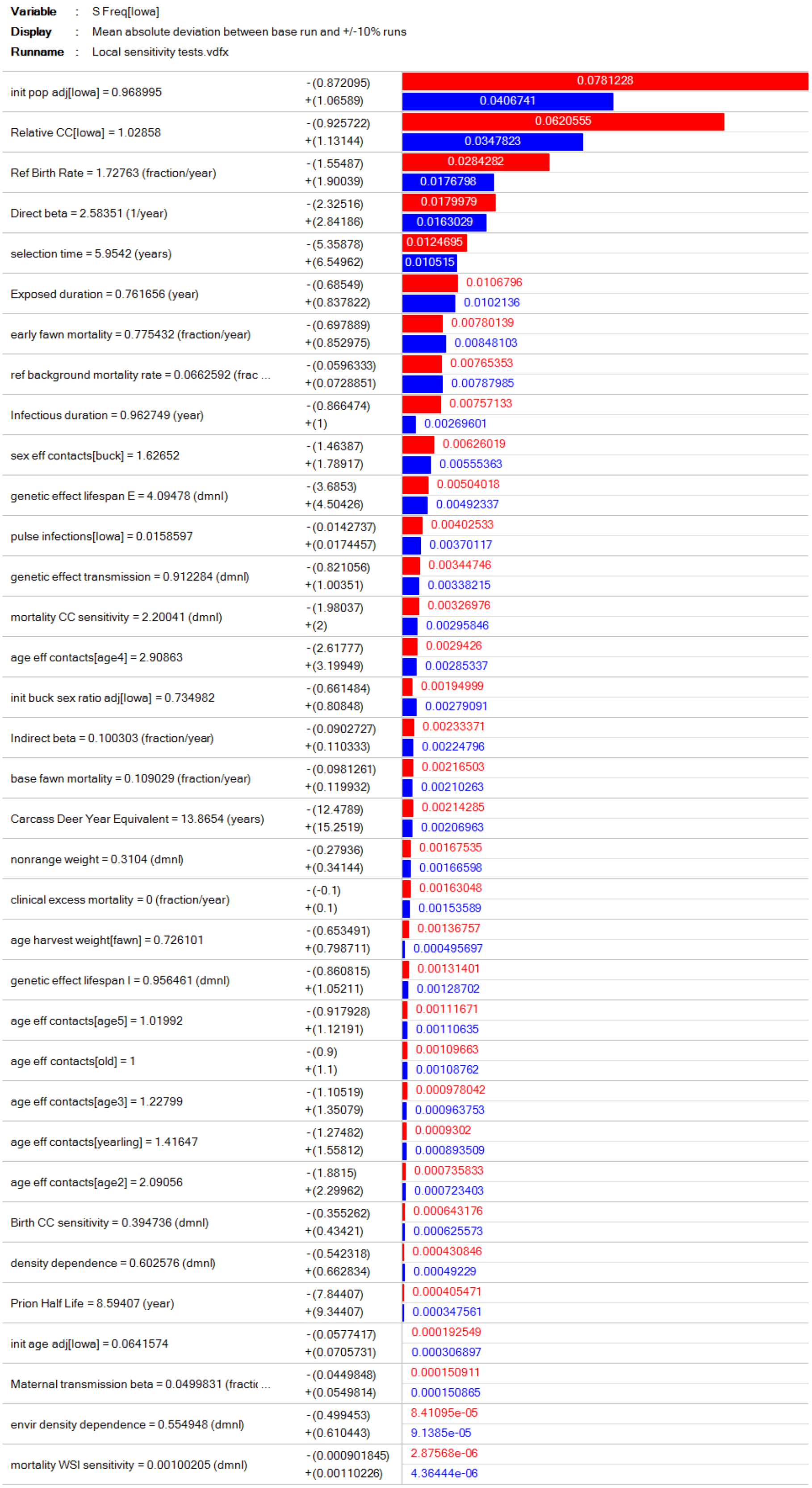

Variable : S Freq[Iowa]
Display : Mean absolute deviation between base run and +/-10% runs
Runname : Local sensitivity tests.vdfx
init pop adj[Iowa] = 0.968995
-(0.872095)
+(1.06589)
0.0781228
0.0406741
Relative CC[Iowa] = 1.02858
-(0.925722)
+(1.13144)
0.0620555
0.0347823
Ref Birth Rate = 1.72763 (fraction/year)
-(1.55487)
+(1.90039)
0.0284282
0.0176798
Direct beta = 2.58351 (1/year)
-(2.32516)
+(2.84186)
0.0179979
0.0163029
selection time = 5.9542 (years)
-(5.35878)
+(6.54962)
0.0124695
0.010515
Exposed duration = 0.761656 (year)
-(0.68549)
+(0.837822)
0.0106796
0.0102136
early fawn mortality = 0.775432 (fraction/year)
-(0.697889)
+(0.852975)
0.00780139
0.00848103
ref background mortality rate = 0.0662592 (frac ...
-(0.0596333)
+(0.0728851)
0.00765353
0.00787985
Infectious duration = 0.962749 (year)
-(0.866474)
+(1)
0.00757133
0.00269601
sex eff contacts[buck] = 1.62652
-(1.46387)
+(1.78917)
0.00626019
0.00555363
genetic effect lifespan E = 4.09478 (dmnl)
-(3.6853)
+(4.50426)
0.00504018
0.00492337
pulse infections[Iowa] = 0.0158597
-(0.0142737)
+(0.0174457)
0.00402533
0.00370117
genetic effect transmission = 0.912284 (dmnl)
-(0.821056)
+(1.00351)
0.00344746
0.00338215
mortality CC sensitivity = 2.20041 (dmnl)
-(1.98037)
+(2)
0.00326976
0.00295846
age eff contacts[age4] = 2.90863
-(2.61777)
+(3.19949)
0.0029426
0.00285337
init buck sex ratio adj[Iowa] = 0.734982
-(0.661484)
+(0.80848)
0.00194999
0.00279091
Indirect beta = 0.100303 (fraction/year)
-(0.0902727)
+(0.110333)
0.00233371
0.00224796
base fawn mortality = 0.109029 (fraction/year)
-(0.0981261)
+(0.119932)
0.00216503
0.00210263
Carcass Deer Year Equivalent = 13.8654 (years)
-(12.4789)
+(15.2519)
0.00214285
0.00206963
nonrange weight = 0.3104 (dmnl)
-(0.27936)
+(0.34144)
0.00167535
0.00166598
clinical excess mortality = 0 (fraction/year)
-(-0.1)
+(0.1)
0.00163048
0.00153589
age harvest weight[fawn] = 0.726101
-(0.653491)
+(0.798711)
0.00136757
0.000495697
genetic effect lifespan I = 0.956461 (dmnl)
-(0.860815)
+(1.05211)
0.00131401
0.00128702
age eff contacts[age5] = 1.01992
-(0.917928)
+(1.12191)
0.00111671
0.00110635
age eff contacts[old] = 1
-(0.9)
+(1.1)
0.00109663
0.00108762
age eff contacts[age3] = 1.22799
-(1.10519)
+(1.35079)
0.000978042
0.000963753
age eff contacts[yearling] = 1.41647
-(1.27482)
+(1.55812)
0.0009302
0.000893509
age eff contacts[age2] = 2.09056
-(1.8815)
+(2.29962)
0.000735833
0.000723403
Birth CC sensitivity = 0.394736 (dmnl)
-(0.355262)
+(0.43421)
0.000643176
0.000625573
density dependence = 0.602576 (dmnl)
-(0.542318)
+(0.662834)
0.000430846
0.00049229
Prion Half Life = 8.59407 (year)
-(7.84407)
+(9.34407)
0.000405471
0.000347561
init age adj[Iowa] = 0.0641574
-(0.0577417)
+(0.0705731)
0.000192549
0.000306897
Maternal transmission beta = 0.0499831 (fractio ...
-(0.0449848)
+(0.0549814)
0.000150911
0.000150865
envir density dependence = 0.554948 (dmnl)
-(0.499453)
+(0.610443)
8.41095e-05
9.1385e-05
mortality WSI sensitivity = 0.00100205 (dmnl)
-(0.000901845)
+(0.00110226)
2.87568e-06
4.36444e-06

e)

**Variable** : yearling harvest share[Iowa,buck]
**Display** : Mean absolute deviation between base run and +/-10% runs
**Runname** : Local sensitivity tests.vdfx

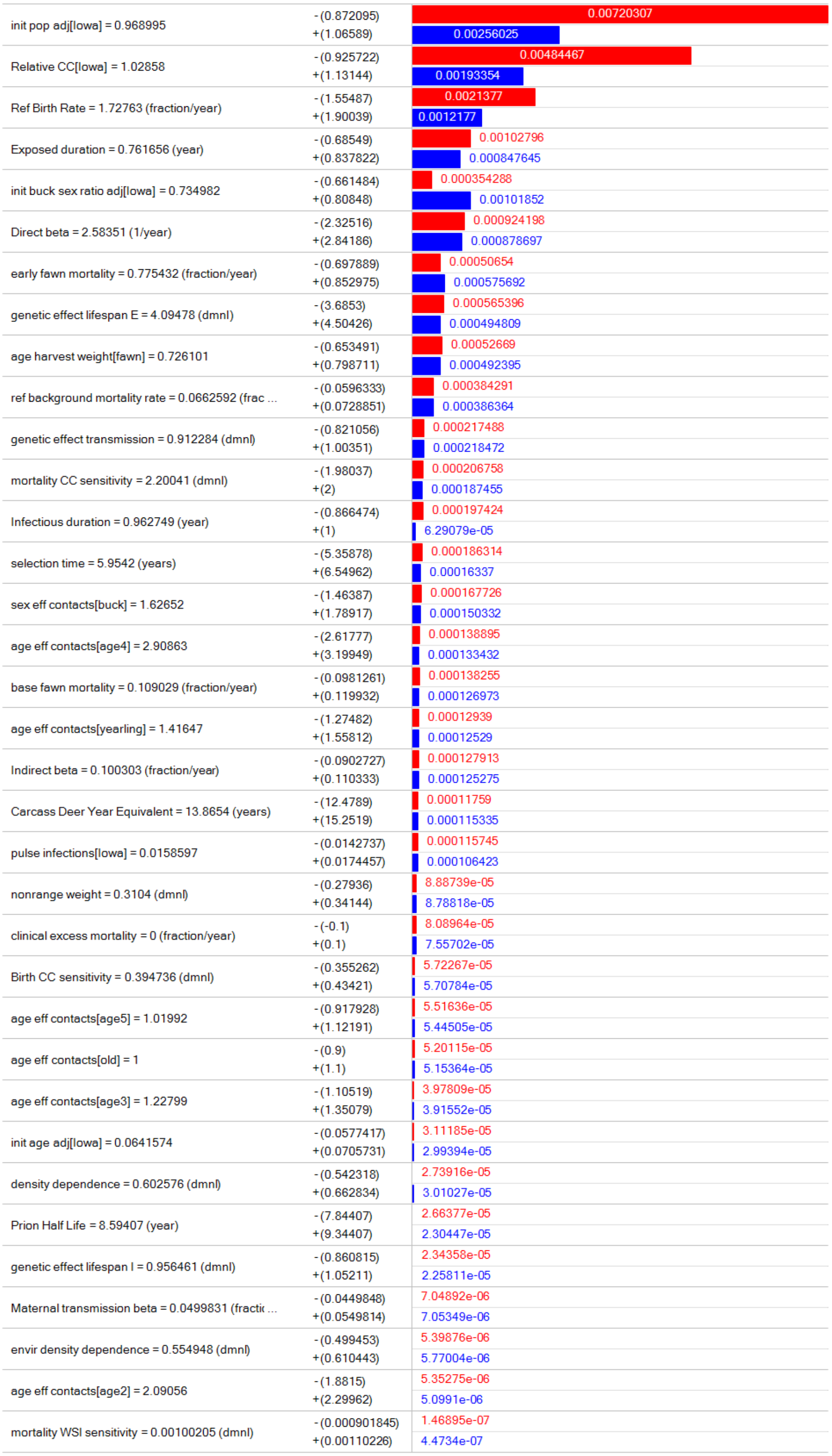

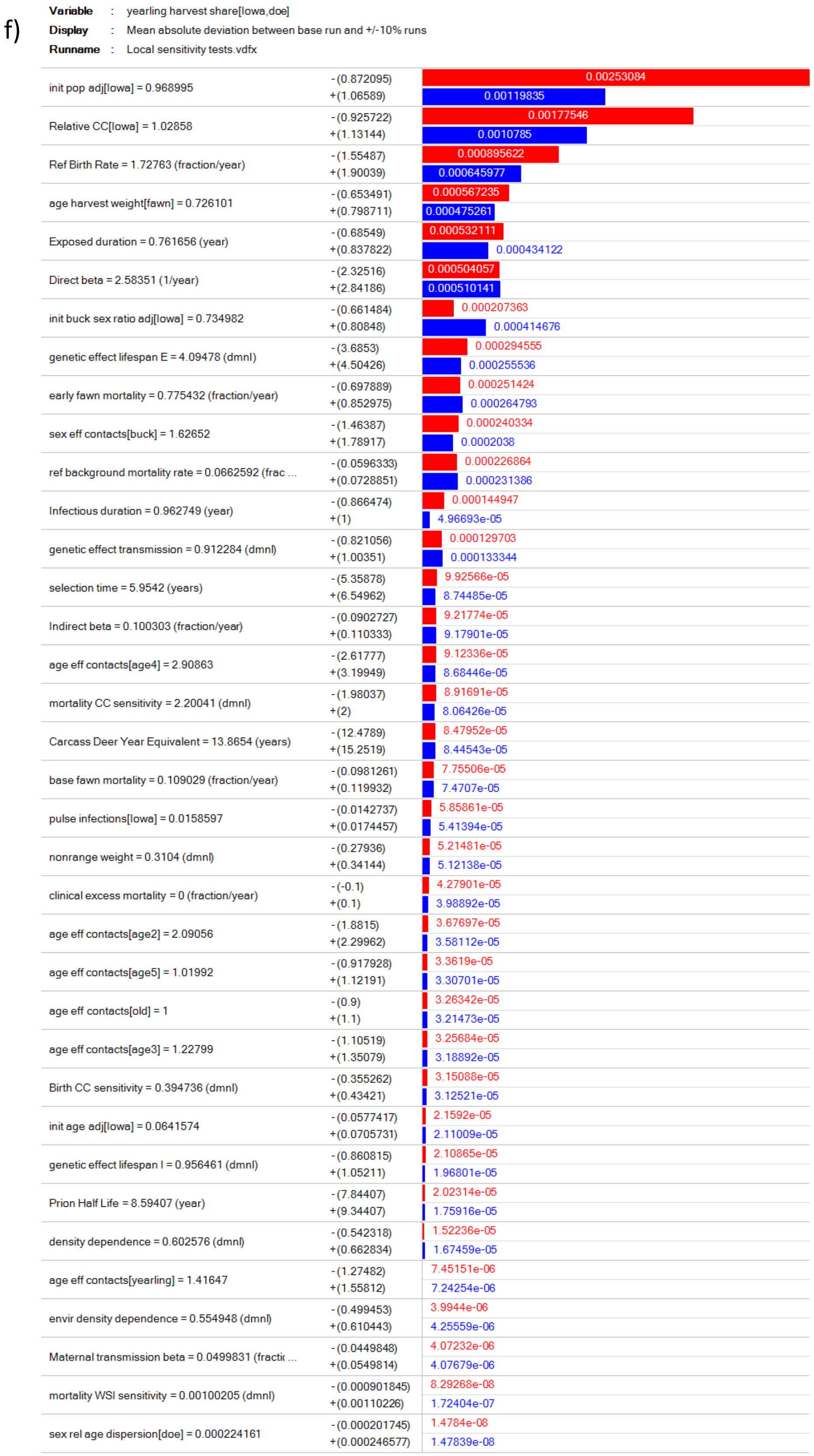


**Figure S16. The results from the *Sensitivy2All* local parameter sensitivity test, performed in Vensim, are**

**presented for each model simulated estimate of the historical time-series data from Wisconsin used to fit the Susceptible-Infected-Clinical (*SIC*) model.** For each parameter estimated during model calibration, the central value was independently increased and decreased by 10%. The mean absolute deviation between the baseline simulation—using all mean parameter estimates—and each perturbed run is reported (blue and red bars indicate decreased and increased parameters, respectively). Sensitivity results are shown for the following model predictions for Iowa county (units in brackets) in this order: (a) the post-hunt deer population estimate (deer), (b) the number of CWD positive deer samples (units of deer), (c) the fawn to doe ratio (dimensionless), (d) the frequency of the *S* allele (fraction), (e) ratio of yearling bucks to total harvested bucks (dimensionless), (f) ratio of yearling does to total harvested does (dimensionless). The results from increasing and decreasing parameters are shown in blue and red, respectively, while subscripts are shown in square brackets (e.g., Region = Iowa, sex= buck).

#### 4.2 Hunter submodule

##### 4.2.1 Estimation procedure

The hunter submodel operates at the county level (i.e., *Region*=County) using a continuous-value, discrete-time model that accommodates continuous hunter mortality and the discrete movement between states during the annual licensing event. A disaggregated inactive hunter state variable tracks the number of years hunters remain inactive (refer to *Hunter participation*). The model was fit to time-series data on active hunter numbers, hunter recruitment counts, and reactivation counts from inactive hunter stocks, all provided by the Wisconsin Department of Natural Resources (refer to *Hunter license data*). These data were used to estimate parameters governing hunter recruitment, retention, and reactivation transition probabilities (Table S18). A log-normal error distribution was assumed, as the reactivation count data were sufficiently large and better approximated by log-normal rather than Poisson distributions. The model was constrained by the county population estimates.

Initial values for the active hunter stock and all but the final inactive stage were set to the first observed values. Because mortality accumulates in the final inactivity stage, its initial observation is biased low. To address this, we calculated the proportion of inactive hunters in the final inactive stage at the last observation and assumed it remained constant over time. This proportion was used to estimate the initial number of hunters in the final inactive stage, given the observed counts in earlier inactive stages.

A hierarchical framework accounted for county-level heterogeneity not captured by the model structure, allowing variation in recruitment, retention, and reactivation transition probabilities (Table S18: *relative regional recruitment, relative regional loss,* and *relative regional reactivation*) and wildlife value orientation (Table S18: *relative regional WVO*) parameters, analogous to "random effects" in regression modeling.

We conducted two rounds of model fitting. A full fit, using the complete time series of all three datasets, and a restricted fit, using a dataset that excluded the final four years of the time series datasets (i.e., truncated at 2020) to test the model's generalizability and forecasting ability.

##### 4.2.2 Estimation results

###### *4.2.2.1 Parameters*

**Table S18. Summary of estimated recruitment, retention and reactivation transition probabilities and the wildlife value orientation parameters for Wisconsin estimated during calibration of the hunter submodel to**

**the full dataset.** Each variable is reported using its internal Vensim name for transparency and reproducibility. Priors were specified directly for each parameter where data or expert judgment permitted. To prevent the optimization algorithm from exploring biologically implausible regions of the parameter space (e.g., negative transition probabilities), we imposed hard bounds (minimum and maximum values) on each parameter. Where possible, citations for prior values informed by empirical literature are provided. Final parameter estimates are reported as the posterior mean ± standard deviation, derived from the joint posterior distribution sampled during the fitting process, along with their associated univariate Rubin/Brooks-Gelman potential scale reduction factor (PSRF; Brooks and Gelman, 1998). PSRF values approaching 1 indicate successful convergence, whereas values greater than 1.2 suggest non-convergence. Maximum PSRF values are reported for region specific parameters.

| Variable | Definition | Informative Prior [Min, Max] | Estimated Values | PSRF | Literature |
|---|---|---|---|---|---|
| recruitment | An instantaneous rate representing the global mean value for the recruitment of potential hunters across Wisconsin. | *Lognormal*(0.05, 0.05)<br>[1x$10^{-6}$, 1] | (2.91±0.0121)x$10^{-2}$ | 1.01 | (Enck et al., 1973; Riley et al., 2016; The Sporting Heritage Council 2020, 2020) |
| retention | An instantaneous rate representing the global mean value for the retention of active hunters across Wisconsin. | *Lognormal*(0.80, 0.1)<br>[1x$10^{-6}$, 1] | 0.868±4.34x$10^{-4}$ | 1.00 | (Hinrichs et al., 2020) |
| reactivation[Inactive] | An instantaneous rate representing the stage-specific global mean values for the reactivation of inactive hunters across Wisconsin. | *Lognormal*([0.25, 0.13, 0.09, 0.06, 0.05], 0.05)<br>[1x$10^{-6}$, 1] | 0.393±0.00143<br>0.193±0.00131<br>0.120±0.00108<br>0.0861±9.6634x$10^{-4}$<br>0.0213±3.82x$10^{-4}$ | 1.01<br>1.00<br>1.00<br>1.00<br>1.00 | (Hinrichs et al., 2020) |
| WI WVO | The estimated proportion of the population likely to ever consider hunting based on their wildlife value orientation across Wisconsin in 2018. | *Lognormal*(0.57, 0.05)<br>[1x$10^{-6}$, 1] | 0.571±7.30x$10^{-4}$ | 1.00 | (Dietsch et al., 2018) |
| cultural shift in WVO | The annual rate of change in the wildlife value orientation of the Wisconsin population. | *Lognormal*(0.0052, 0.00287)<br>[1x$10^{-6}$, 0.2] | 0.0108±8.20x$10^{-5}$ | 1.00 | (Manfredo et al., 2021a) |
| relative regional recruitment[Region] | A scaling factor applied to the global recruitment parameter to capture variation between counties. | *Uniform*(0.01, 3) | 1.21±0.284<br>(0.13-2.79) | ≤1.03 | The product of the multiplier and the global value is bounded between [1x$10^{-6}$, 1] |
| relative regional loss[Region] | A scaling factor for the global loss parameter -which is equal to one minus retention- to capture variation between counties. | *Uniform*(0.75, 1.5) | 0.970±0.116<br>(0.750-1.35) | ≤1.02 | The product of the multiplier and the global value is bounded between [1x$10^{-6}$, 1] |

| | | | | | |
|---|---|---|---|---|---|
| relative regional reactivation[Region ] | A scaling factor applied to the global reactivation parameters to capture variation between counties. | *Uniform*(0.5, 2) | 0.730±0.0595 (0.520-0.862) | ≤1.03 | The product of the multiplier and the global value is bounded between [1x10$^{-6}$, 1] |
| relative regional WVO[Region] | A scaling factor for the proportion of the WI population to ever consider hunting bade on their wildlife value orientation in 2018 to capture variation between counties. | *Uniform*(0.01, 3) | 1.21±0.284 (0.750-1.50) | ≤1.02 | The product of the multiplier and the global value is bounded between [1x10$^{-6}$, 1] |
| Ireacts data SD[Inactive] | A parameter estimating the variance in the time series of reactivation counts from each inactive hunter stock. | *Uniform*(0, 1) | 0.174±1.93x10$^{-3}$ 0.208±2.13x10$^{-3}$ 0.242±1.98x10$^{-3}$ 0.270±1.95x10$^{-3}$ 0.389±1.99x10$^{-3}$ | 1.02 | Full range of possibilities for SD around probability |

#### *4.2.2.2 Maps of region-specific parameter values*

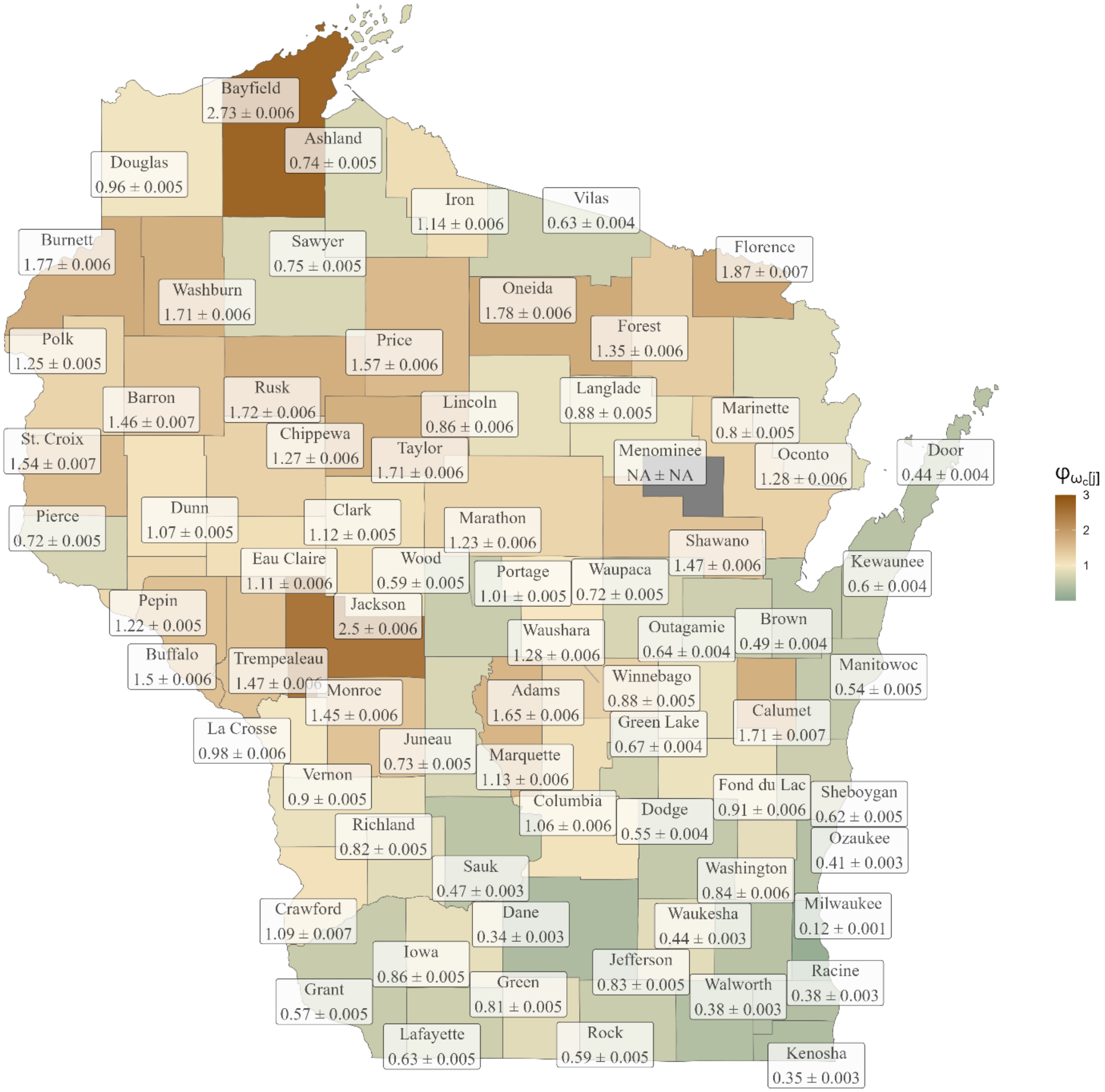


**Figure S17. County specific mean and ± 1 standard deviation of regional relative recruitment in Wisconsin estimated from the submodel of hunters, fit to the full dataset.** A value of one indicates that a county's recruitment matches the global mean exactly; values above and below one indicate higher and lower county-level recruitment county-level recruitment.

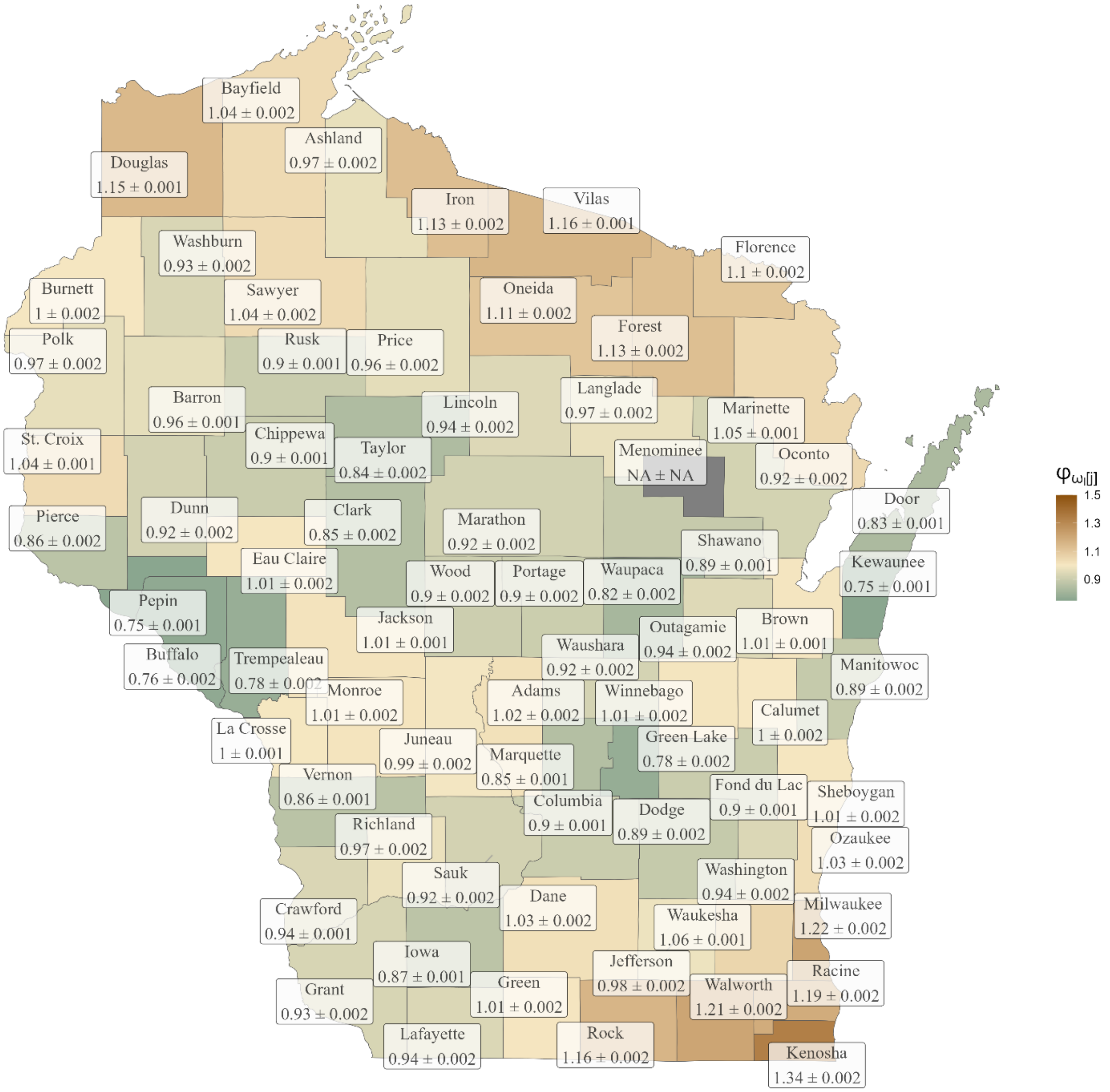


**Figure S18. County specific mean and ± 1 standard deviation of regional relative loss in Wisconsin estimated from the hunter submodel, fit to the full dataset.** A value of one indicates that a county's loss matches the global mean; values above and below indicate higher and lower county-level loss respectively.

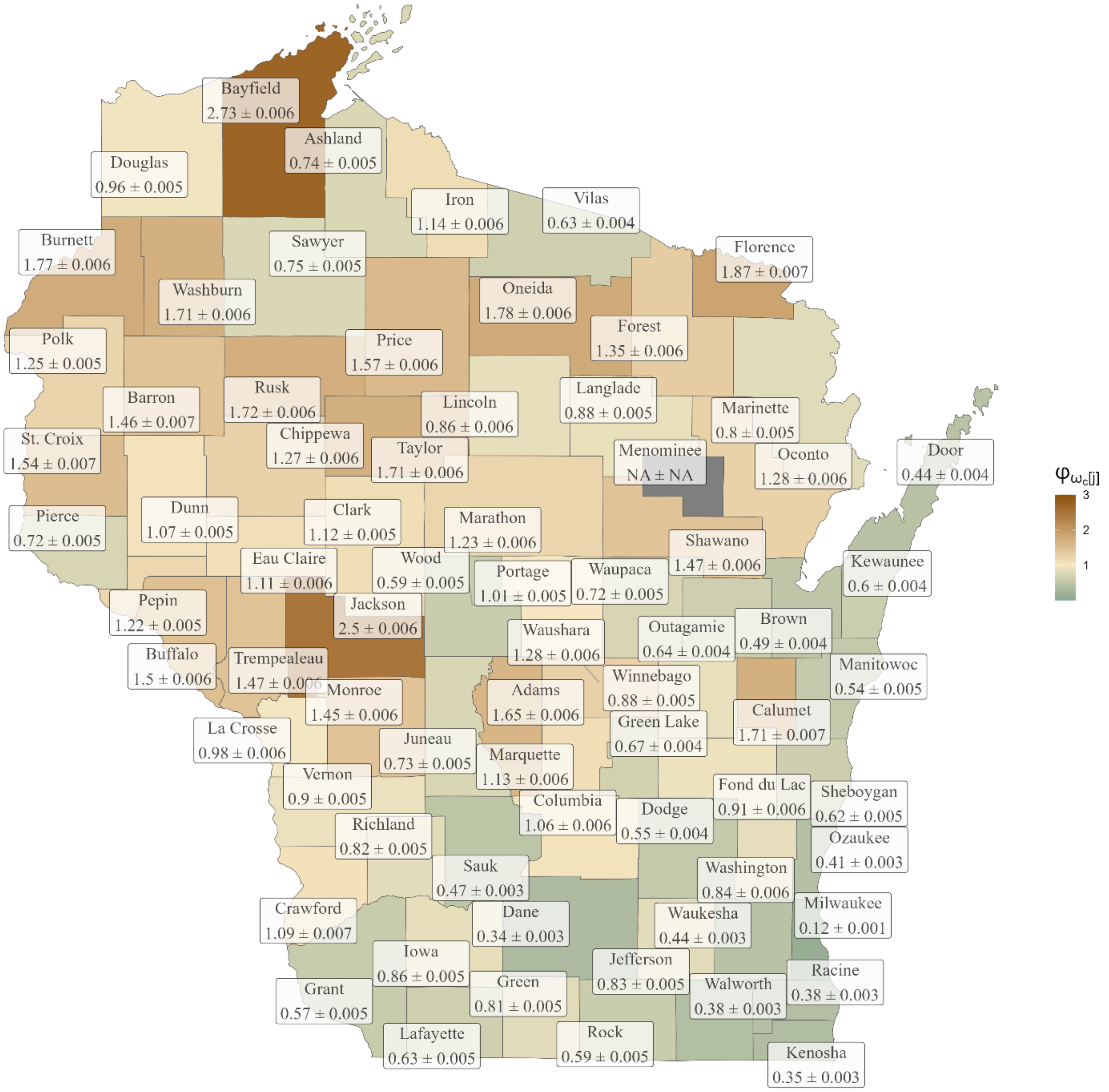


**Figure S19. County specific mean and ± 1 standard deviation of regional relative reactivation in Wisconsin estimated from the hunter submodel, fit to the full dataset.** A value of one indicates that a county's reactivation matches the global mean; values above and below indicate county-level reactivation is higher and lower respectively.

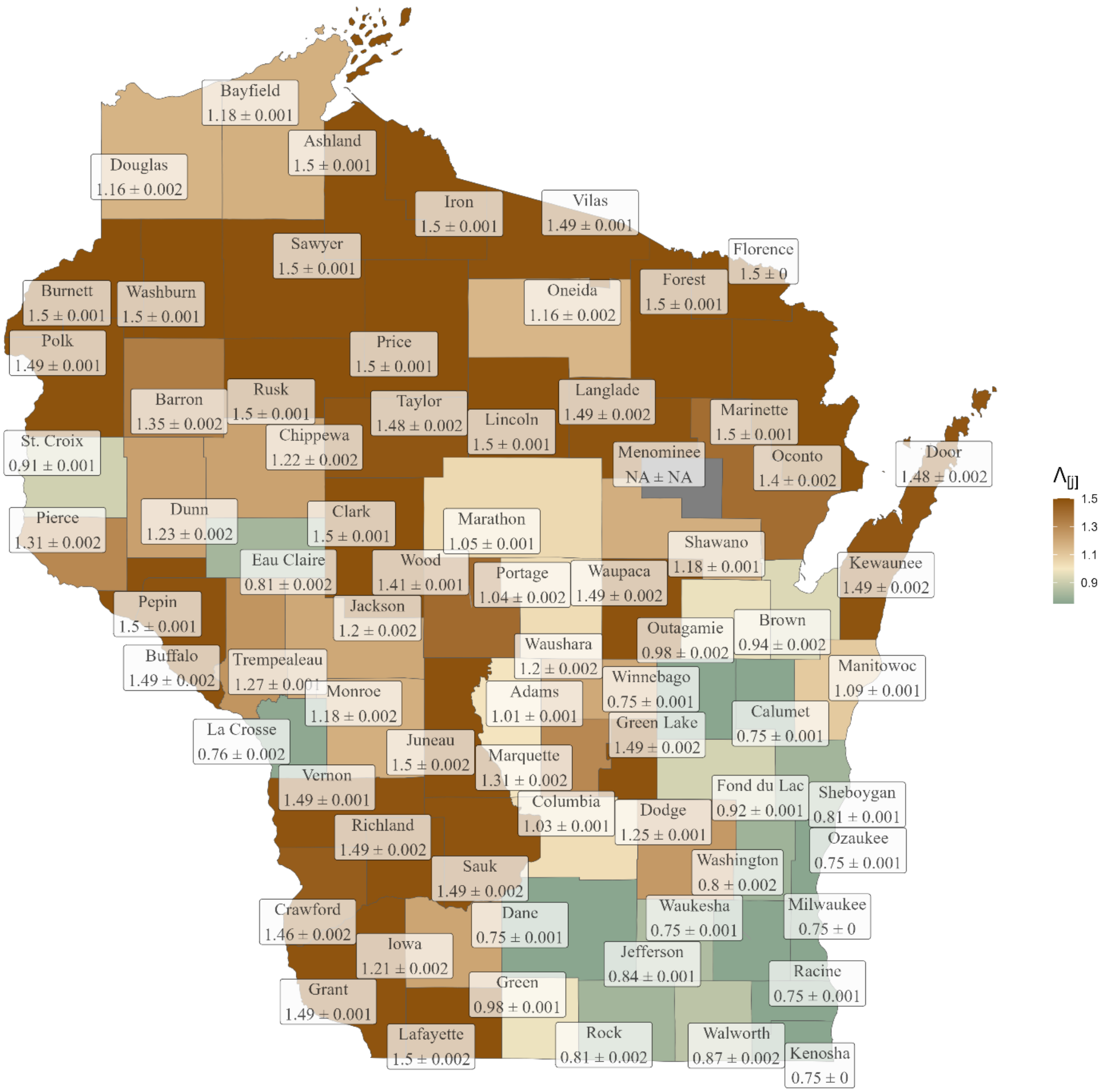


**Figure S20. County specific mean and ± 1 standard deviation of regional wildlife value orientation (WVO) in Wisconsin estimated from the hunter submodel, fit to the full dataset.** A value of one indicates that a county's WVO matches the global mean; values above and below indicate county-level WVO is higher and lower respectively.

##### 4.2.2.3 Model predictions

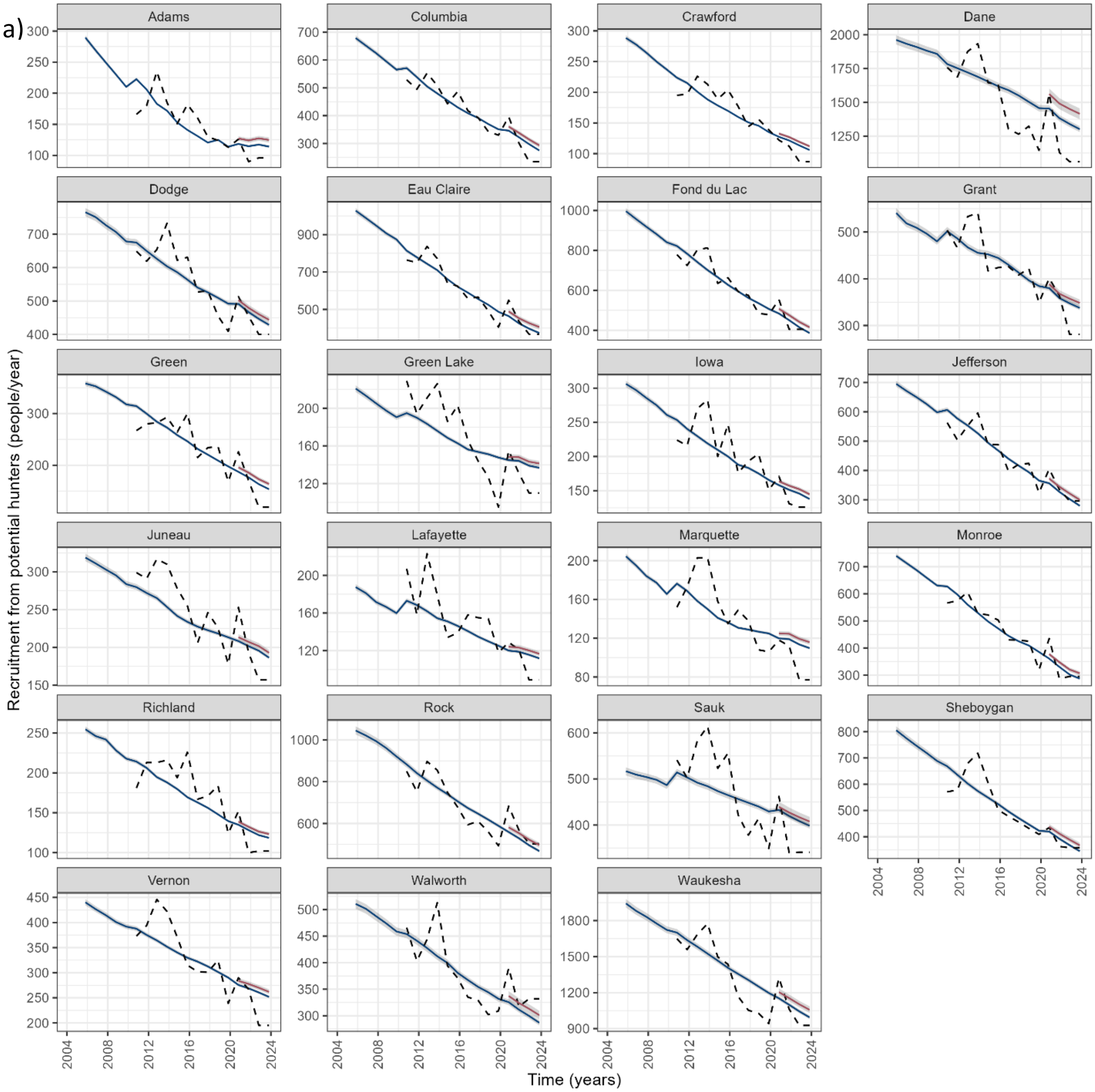

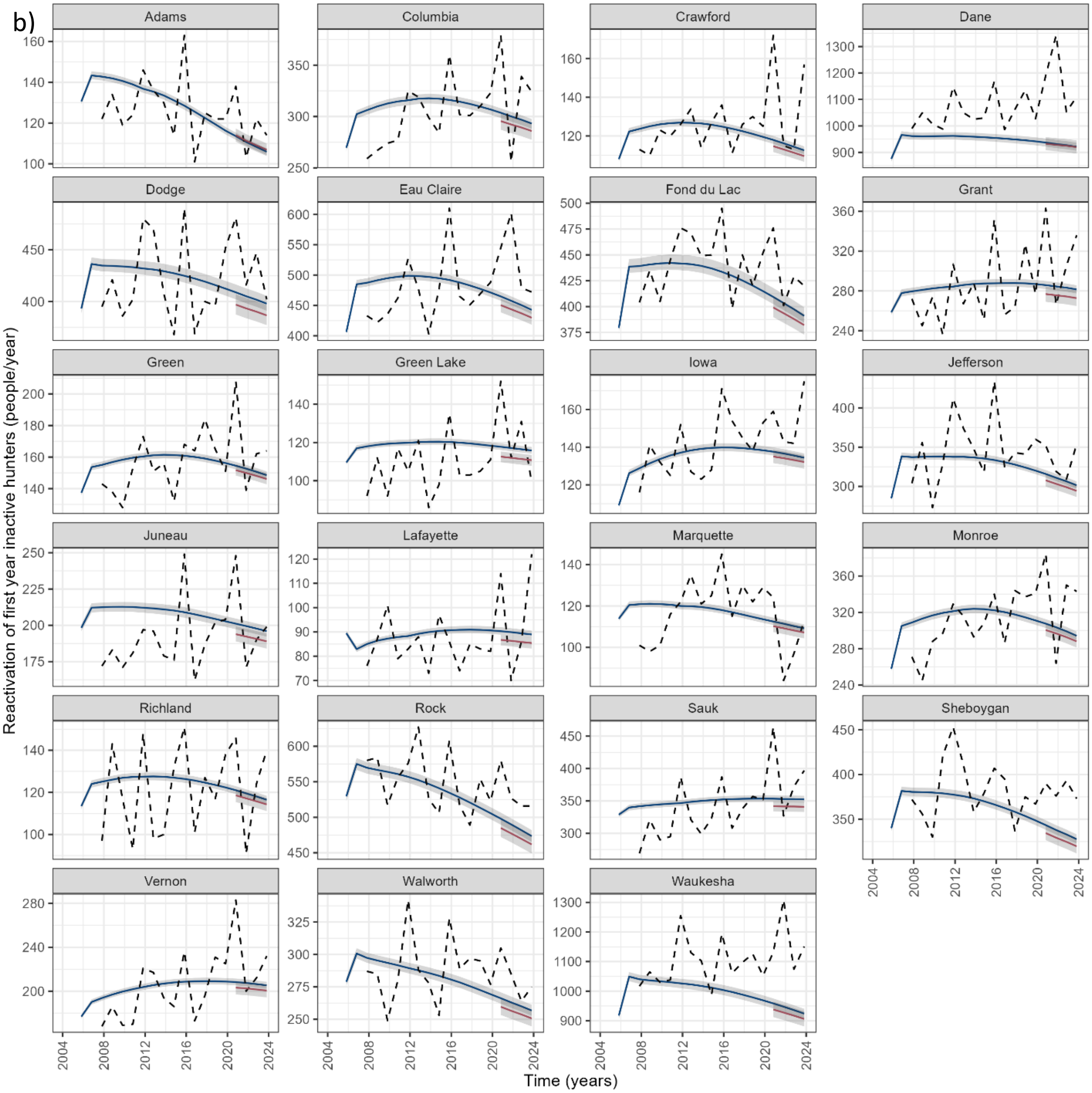
b)
Adams
Columbia
Crawford
Dane
Dodge
Eau Claire
Fond du Lac
Grant
Green
Green Lake
Iowa
Jefferson
Juneau
Lafayette
Marquette
Monroe
Richland
Rock
Sauk
Sheboygan
Vernon
Walworth
Waukesha
Reactivation of first year inactive hunters (people/year)
Time (years)

c)

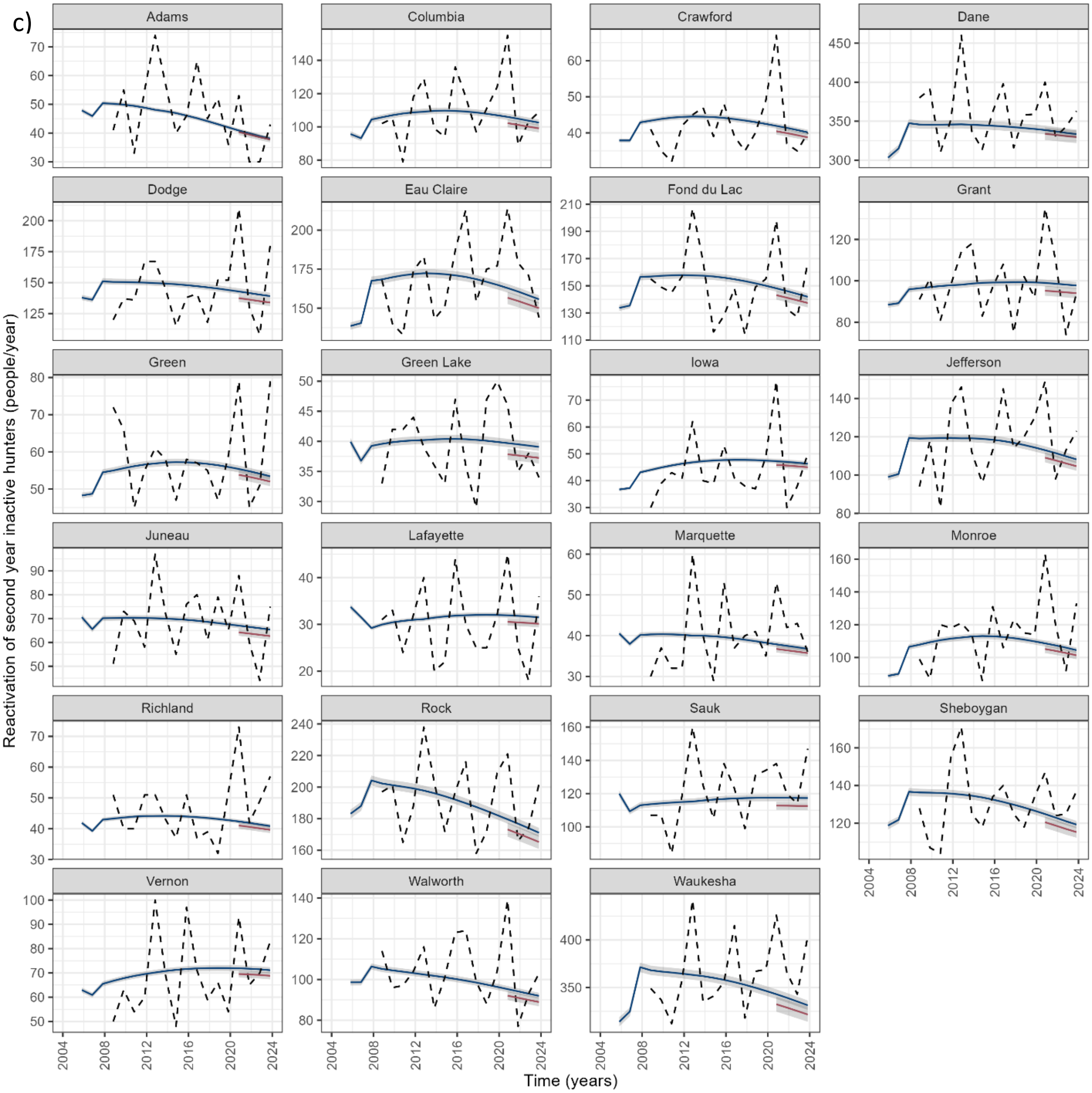

Adams
Columbia
Crawford
Dane
Dodge
Eau Claire
Fond du Lac
Grant
Green
Green Lake
Iowa
Jefferson
Juneau
Lafayette
Marquette
Monroe
Richland
Rock
Sauk
Sheboygan
Vernon
Walworth
Waukesha
Reactivation of second year inactive hunters (people/year)
Time (years)

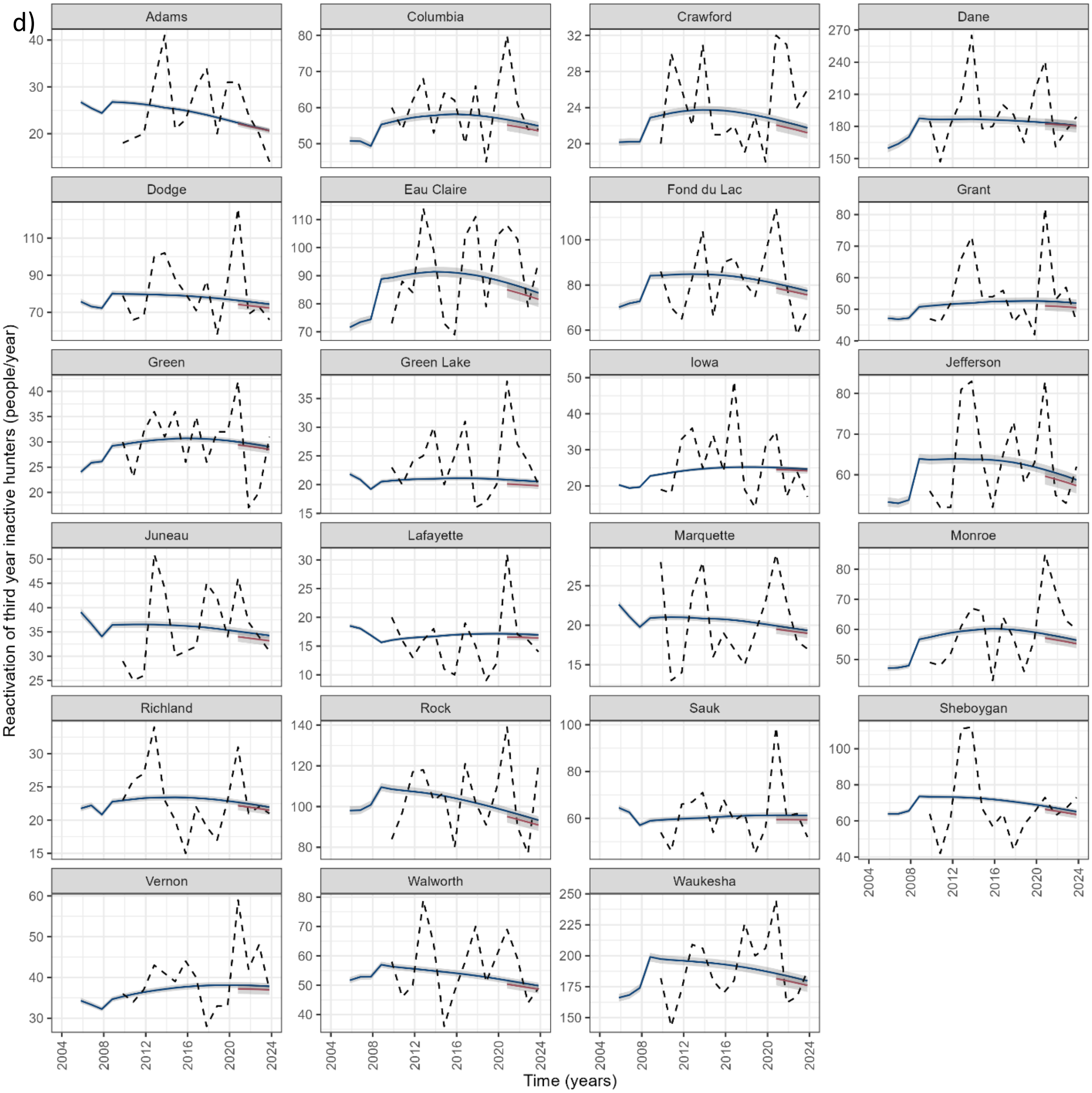
d)
Adams
Columbia
Crawford
Dane
Dodge
Eau Claire
Fond du Lac
Grant
Green
Green Lake
Iowa
Jefferson
Juneau
Lafayette
Marquette
Monroe
Richland
Rock
Sauk
Sheboygan
Vernon
Walworth
Waukesha
Reactivation of third year inactive hunters (people/year)
Time (years)

e)

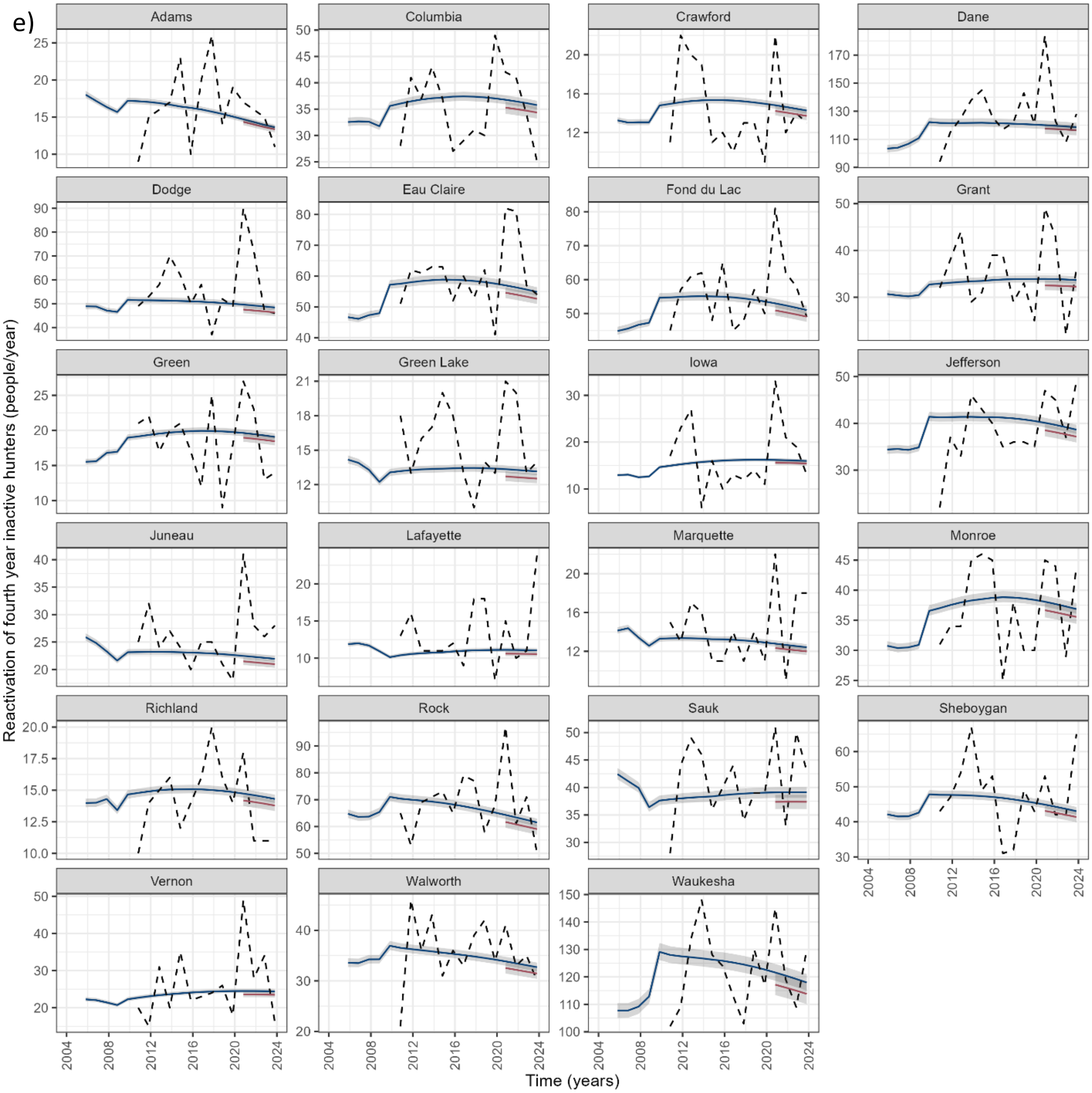
Adams
Columbia
Crawford
Dane
Dodge
Eau Claire
Fond du Lac
Grant
Green
Green Lake
Iowa
Jefferson
Juneau
Lafayette
Marquette
Monroe
Richland
Rock
Sauk
Sheboygan
Vernon
Walworth
Waukesha
Reactivation of fourth year inactive hunters (people/year)
Time (years)

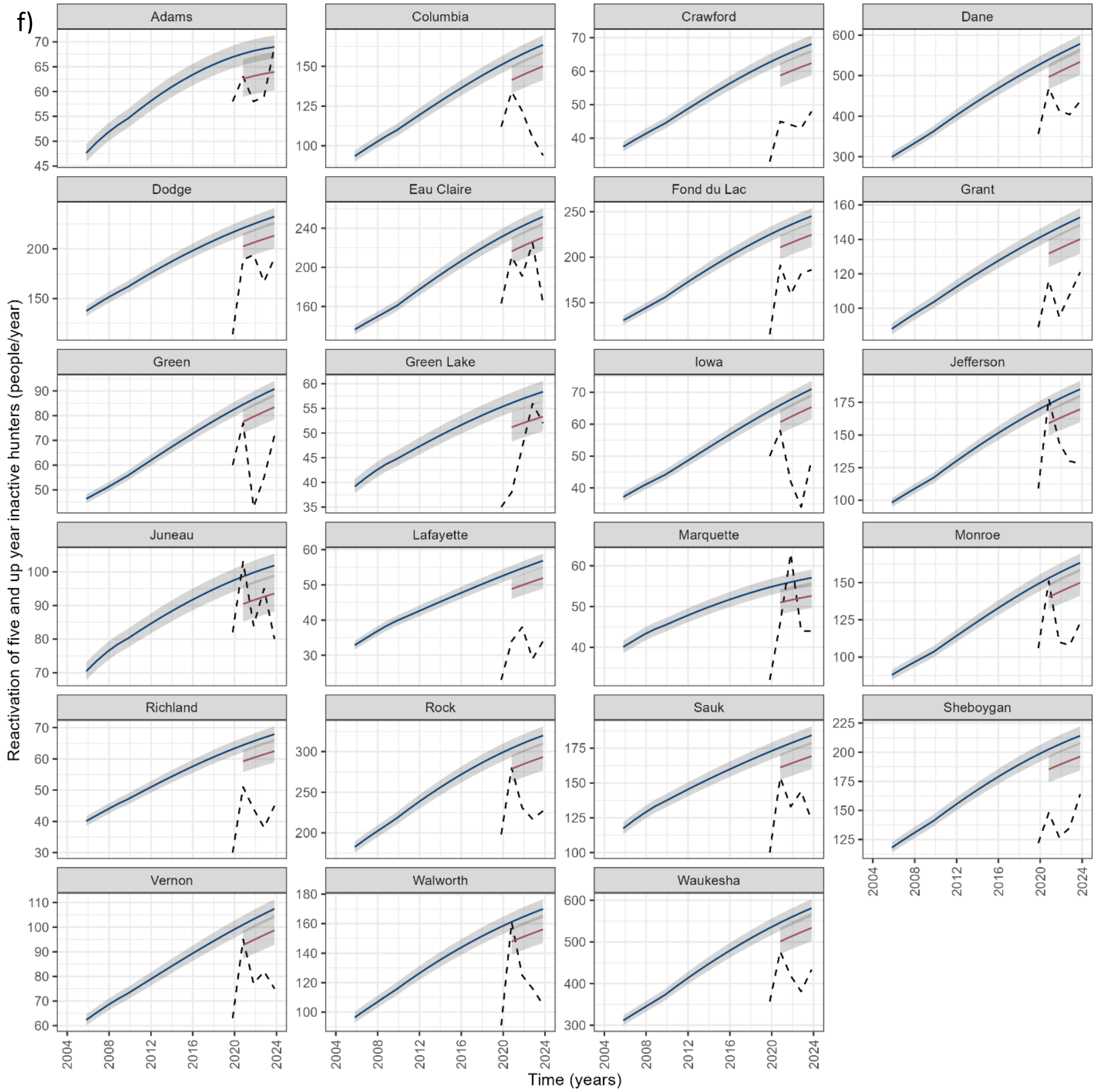


**Figure S21. Model predictions from the hunter submodel, fit to the full dataset (blue) and a restricted dataset when fitted to a restricted dataset truncated at 2019 (red), compared against historical time-series data (dashed) by county provided by the Wisconsin Department of Natural Resources.** Solid lines represent posterior mean predictions, with 95% credible intervals shaded in grey, calculated across 1,000 parameter sets sampled from the joint posterior to reflect both parameter uncertainty and data measurement error. Panels display: (a) annual number of new hunter recruits, (b) annual reactivations of hunters inactive for one year, (c) annual reactivations of hunters inactive for two years, (d) annual reactivations of hunters inactive for three years, (e) annual reactivations of hunters inactive for four years, and (f) annual reactivations of hunters inactive for five or more years.

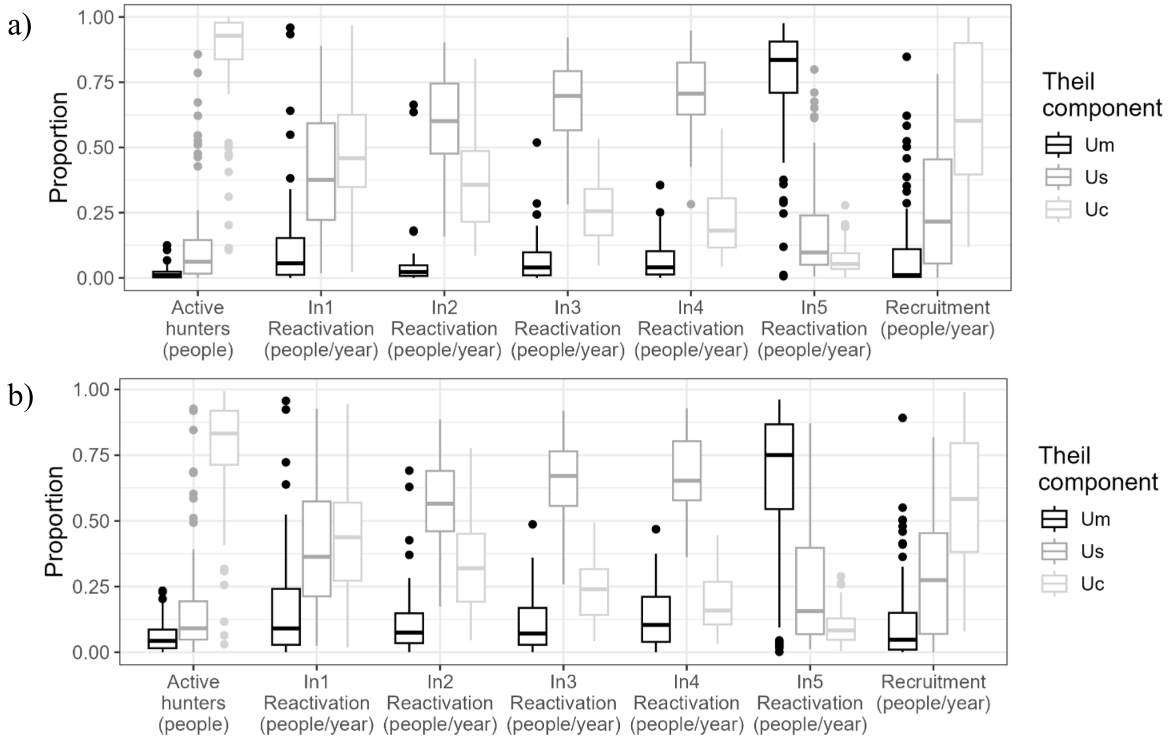


**Figure S22. Theil statistics comparing historical time-series data from the Wisconsin Department of Natural Resources to the simulated estimates from the hunter submodel fit to (a) the full dataset and (b) a restricted dataset truncated at 2019.** Theil statistics decompose the total error between model predictions and observed data into components attributed to: (Um) systematic bias in the mean, (Us) differences in variance, and (Uc) unexplained error (random noise). The three components sum to one. Boxplots represent Theil statistics across regions for seven key metrics: the total number of active hunters (Active hunters), total reactivation counts from inactive stages 1-5 (In1-In5 Reactivation), and the total recruitment counts from the potential hunter pool (Recruitment). Boxes denote the interquartile range (IQR), the line indicates the median, whiskers extend to 1.5×IQR, and points represent outliers.

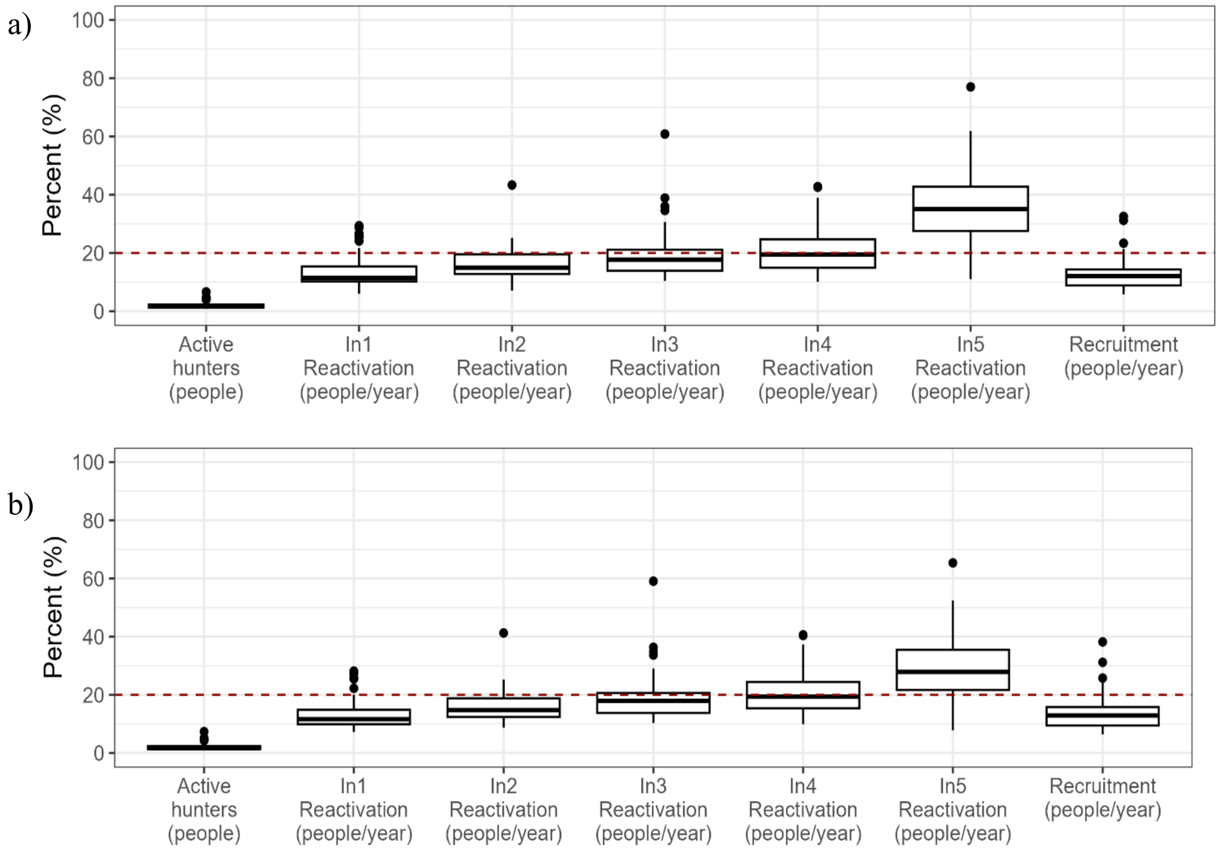


**Figure S23. Mean absolute percentage error (MAPE) comparing historical time-series data from the Wisconsin Department of Natural Resources to the simulated estimates from the hunter submodel fit to (a) the full dataset and (b) a restricted dataset truncated at 2019.** MAPE is a unitless measure of model accuracy, representing the proportion of variance in the data explained by the model. In the context of large-scale system dynamics models, MAPE values below 20% and 50% are typically considered indicative of good and reasonable forecasting performance, respectively. Boxplots display MAPE across regions for seven key metrics: the total number of active hunters (Active hunters), total reactivation counts from inactive stages 1-5 (In1-In5 Reactivation), and the total recruitment counts from the potential hunter pool (Recruitment). The box captures the interquartile range (IQR) of the values and the median is displayed as a line, the whiskers capture the furthest datapoints within 1.5xIQR while points indicate outliers.

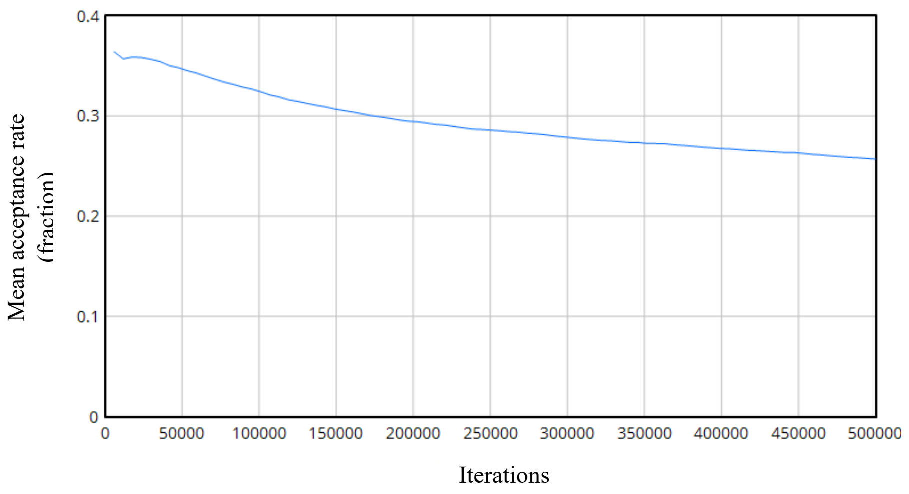


**Figure S24. Mean acceptance rate across Markov chain Monte Carlo chains (reflected as a proportion) from the posterior distribution of the hunter submodel when fit to the full dataset.** Acceptance rates reflect the efficiency of the sampling process and indicate good convergence when rates are neither too low (indicating poor mixing) nor too high (indicating overly conservative proposal steps).

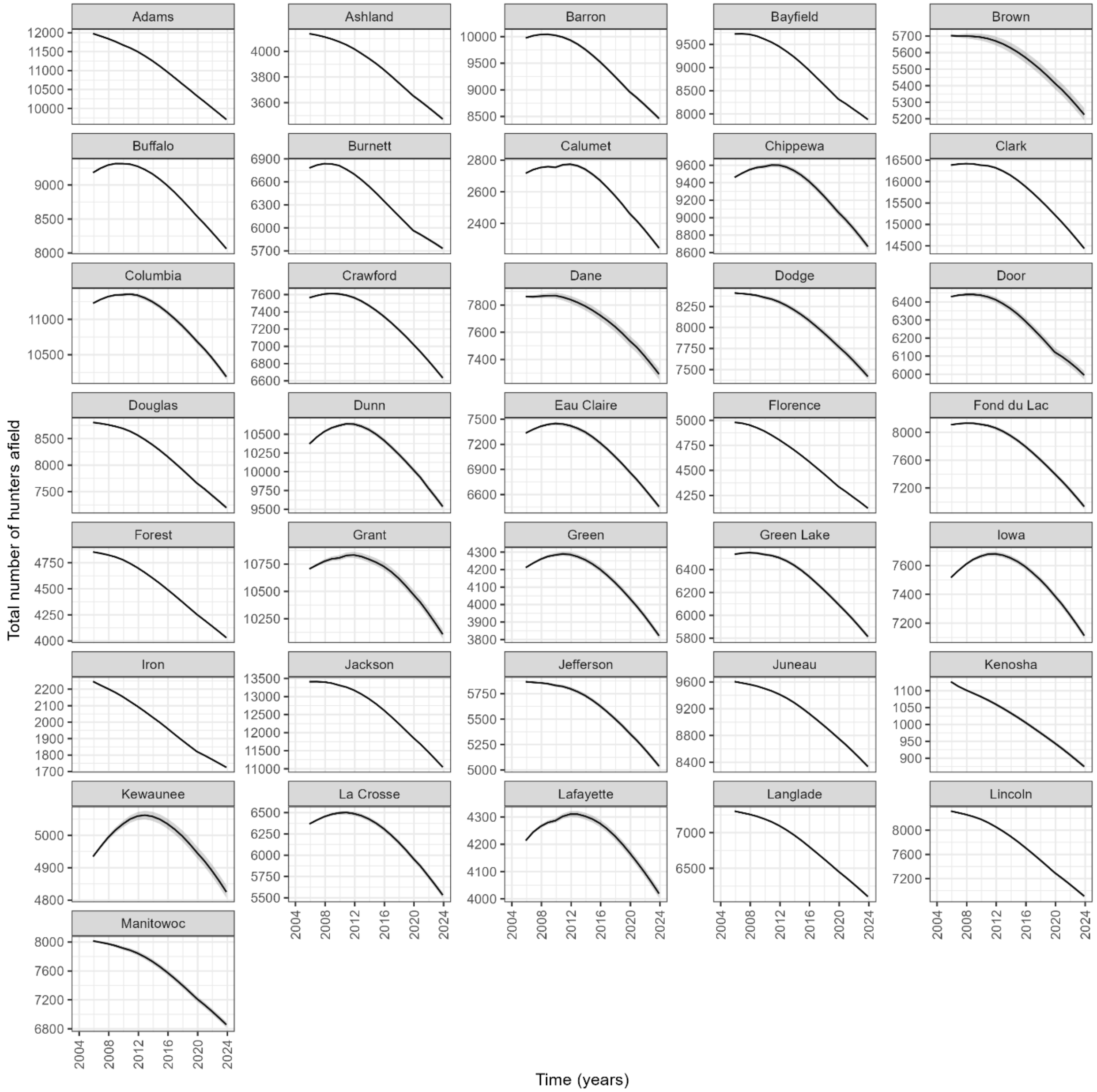


**Figure S25. The number of hunters afield attempting to harvest a deer in each county as predicted by the hunter submodel when fit to the full dataset.** Solid lines represent posterior mean predictions, with 95% credible intervals shaded in grey, calculated across 1,000 parameter sets sampled from the joint posterior to reflect both parameter uncertainty and data measurement error. This figure presents results for the first half of Wisconsin counties included in the model fitting.

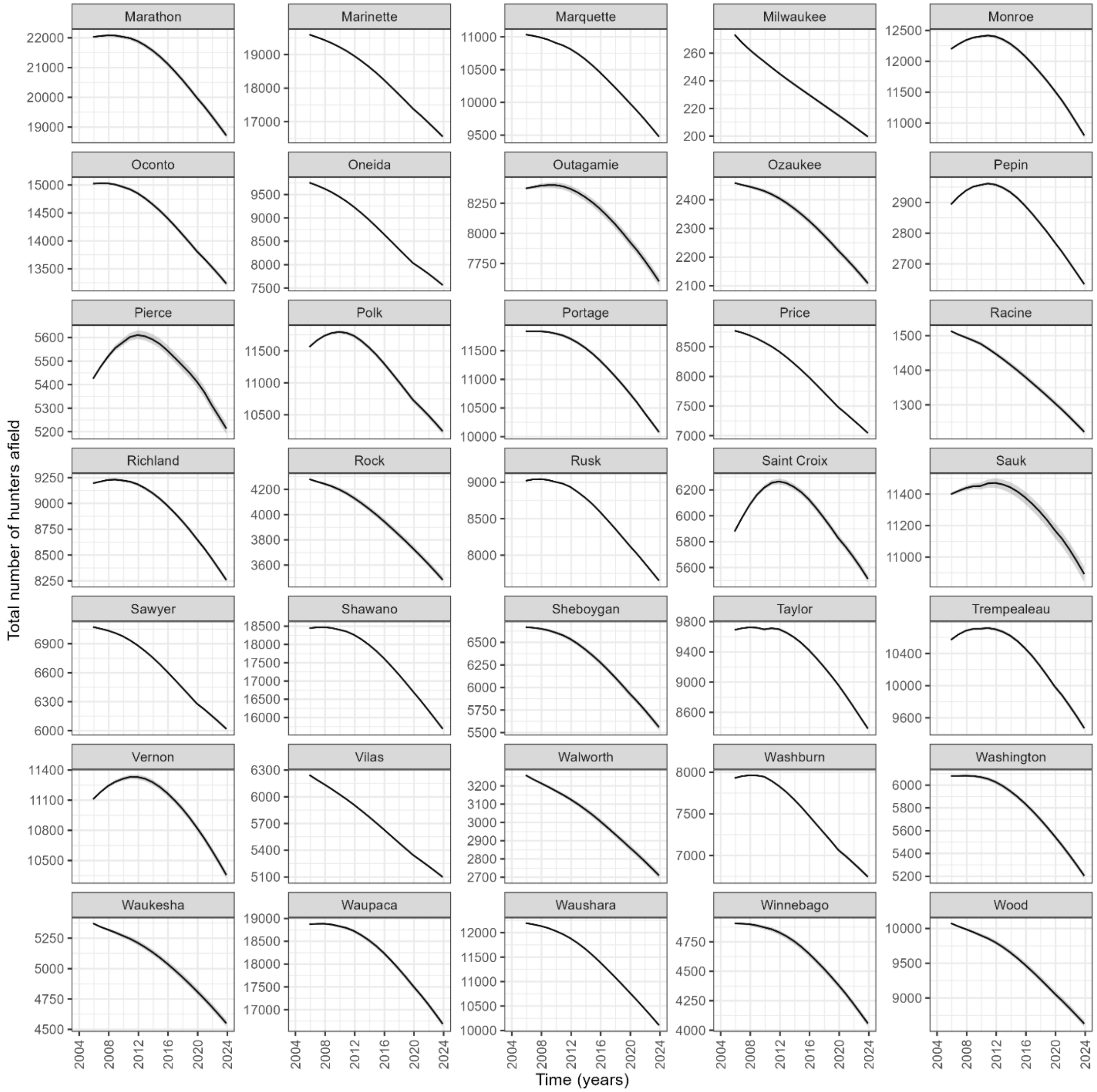


**Figure S26. The number of hunters afield attempting to harvest a deer in each county as predicted by the hunter submodel when fit to the full dataset.** Solid lines represent posterior mean predictions, with 95% credible intervals shaded in grey, calculated across 1,000 parameter sets sampled from the joint posterior to reflect both parameter uncertainty and data measurement error. This figure presents results for the second half of Wisconsin counties included in the model fitting.

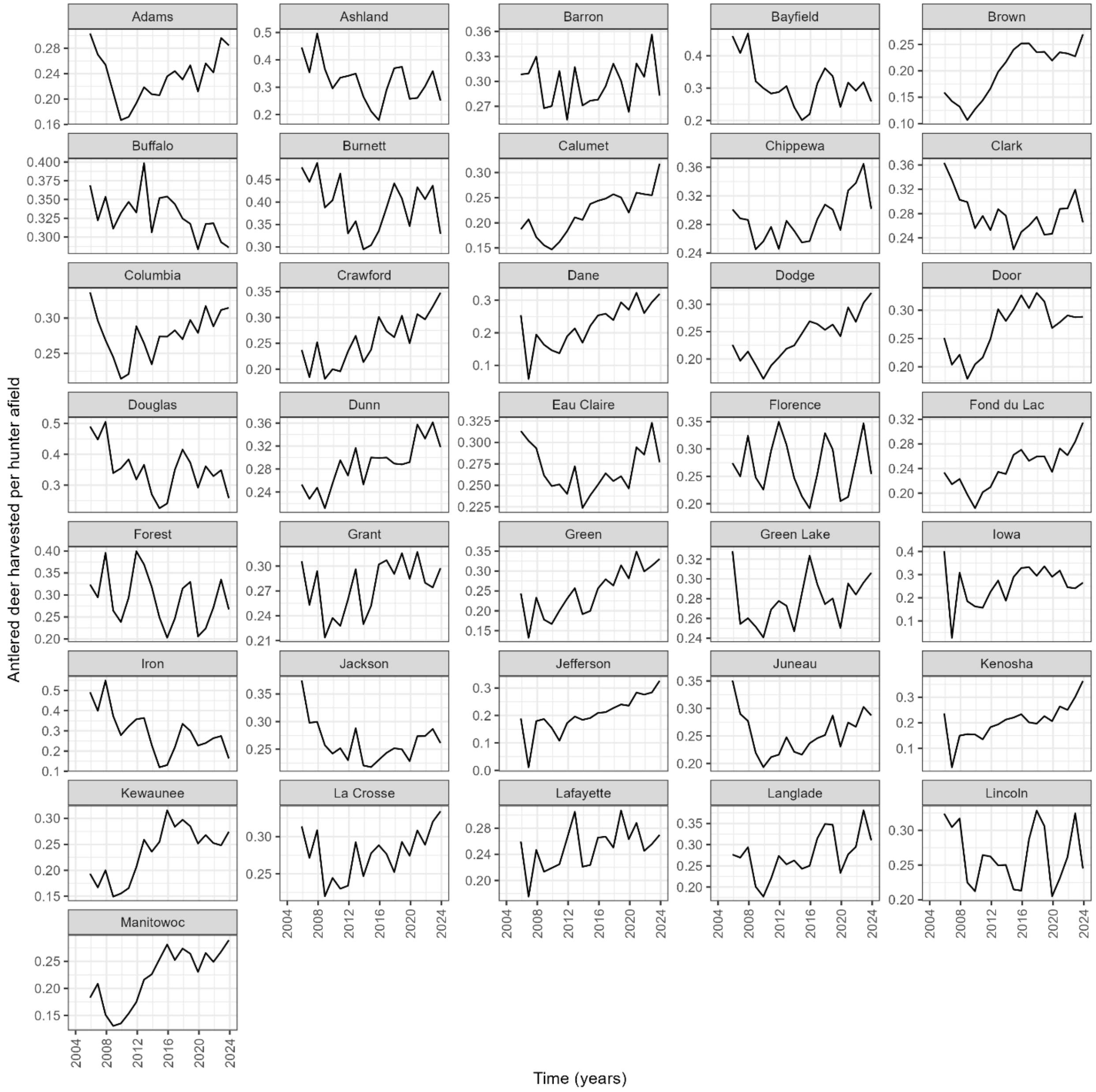


**Figure S27. The number of antlered deer harvested per hunter afield in each county as predicted by the hunter submodel when fit to the full dataset.** Solid lines represent posterior mean predictions, with 95% credible intervals shaded in grey, calculated across 1,000 parameter sets sampled from the joint posterior to reflect both parameter uncertainty and data measurement error. Note that the credible intervals are very small given the small variation in the number of hunters predicted. The estimates were calculated by dividing the historical number of antlered deer harvested (from Wisconsin Department of Natural Resources records) by the model-estimated total number of hunters afield in each county, thus including hunters who did not harvest any deer. This figure presents results for the first half of Wisconsin counties included in the model fitting.

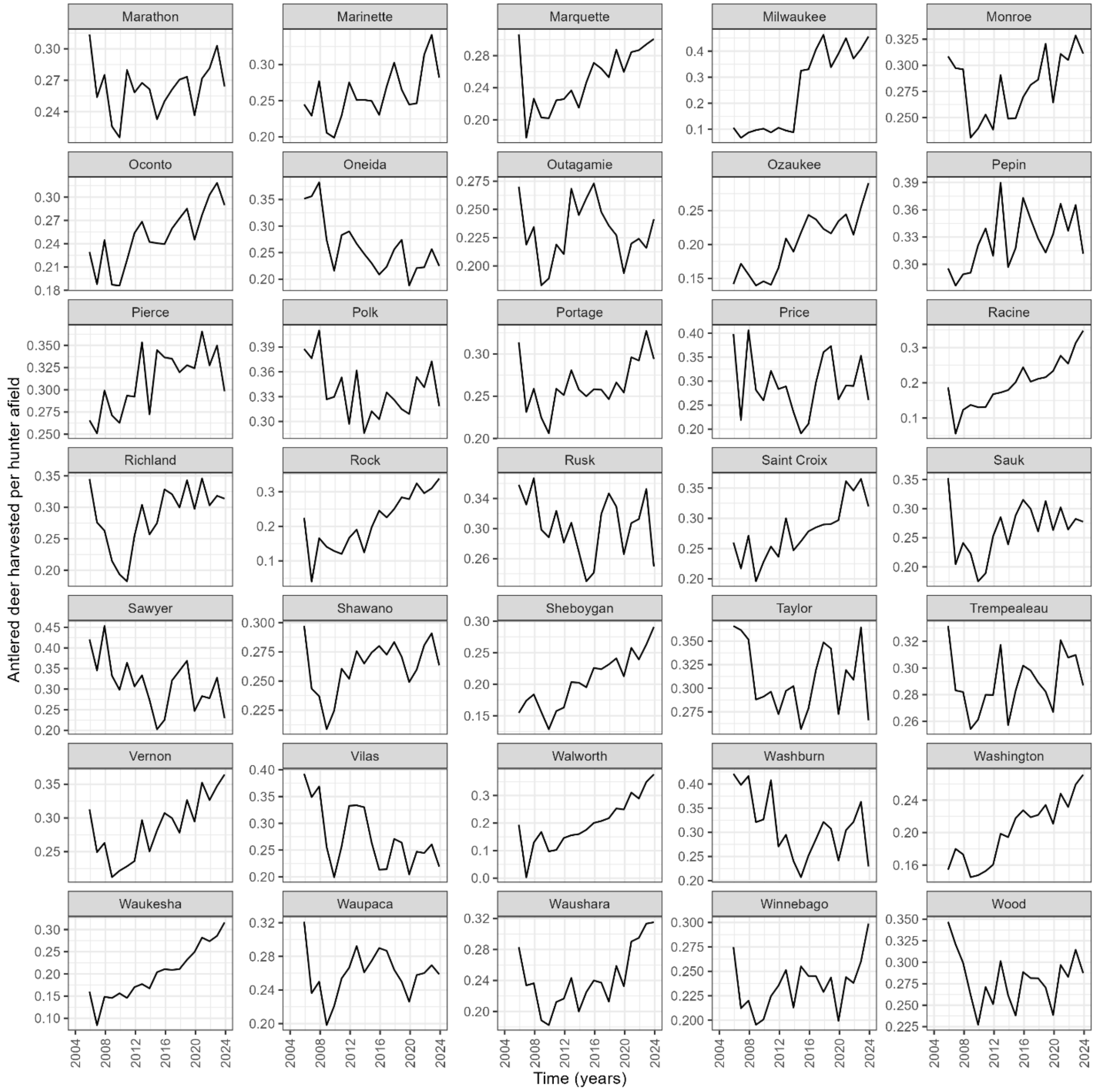


**Figure S28. The number of antlered deer harvested per hunter afield in each county as predicted by the hunter submodel when fit to the full dataset.** Solid lines represent posterior mean predictions, with 95% credible intervals shaded in grey, calculated across 1,000 parameter sets sampled from the joint posterior to reflect both parameter uncertainty and data measurement error. Note that the credible intervals are very small given the small variation in the number of hunters predicted. The estimates were calculated by dividing the historical number of antlered deer harvested (from Wisconsin Department of Natural Resources records) by the model-estimated total number of hunters afield in each county, thus including hunters who did not harvest any deer. This figure presents results for the second half of Wisconsin counties included in the model fitting.

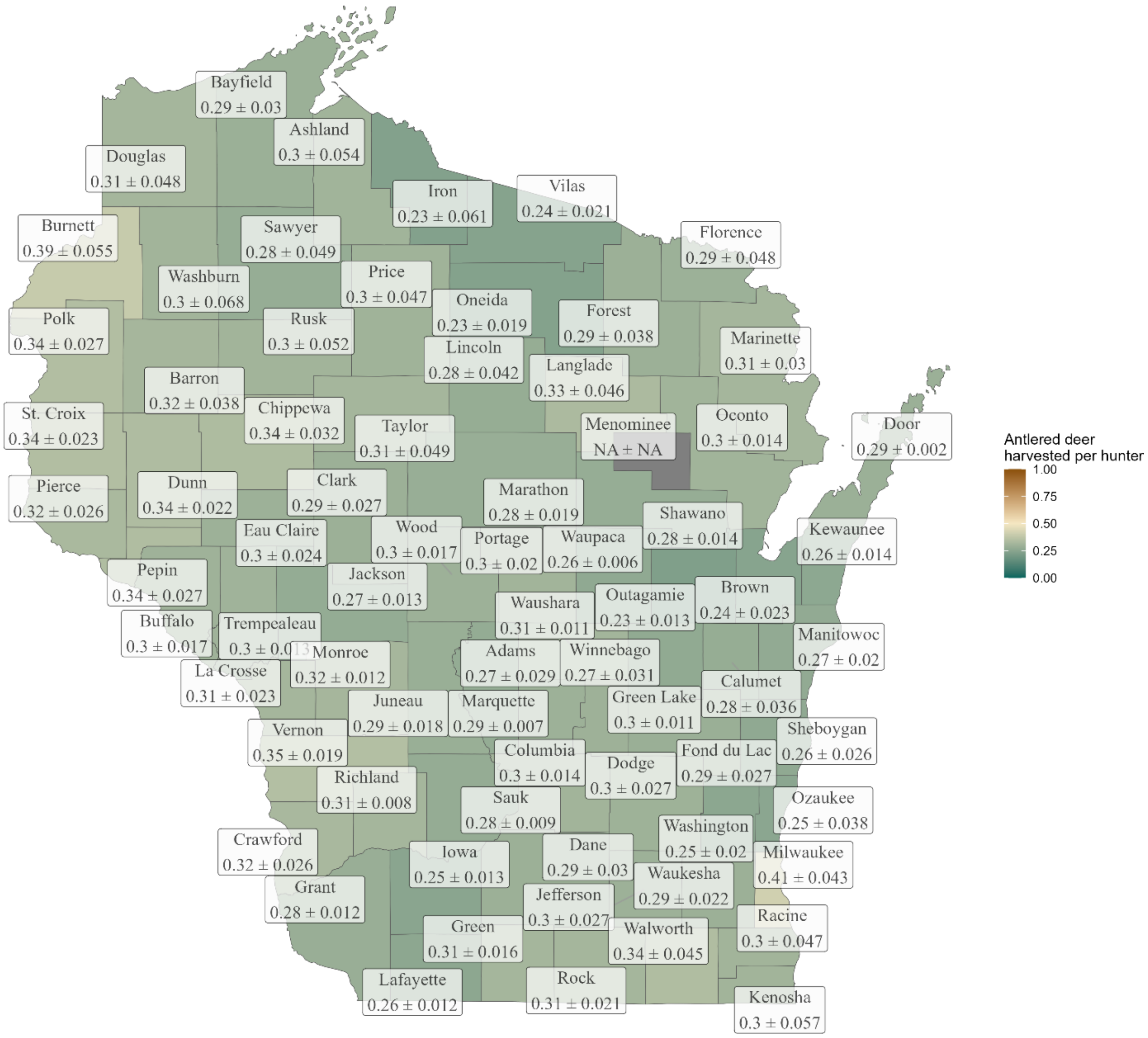


**Figure S29. The number of antlered deer harvested per hunter afield in each Wisconsin county, averaged over the final three years of data, as predicted by the hunter submodel, fit to full dataset.** The estimates were calculated by dividing the historical number of antlered deer harvested (from Wisconsin Department of Natural Resources records) by the model-estimated total number of hunters afield in each county, thus including hunters who did not harvest any deer.

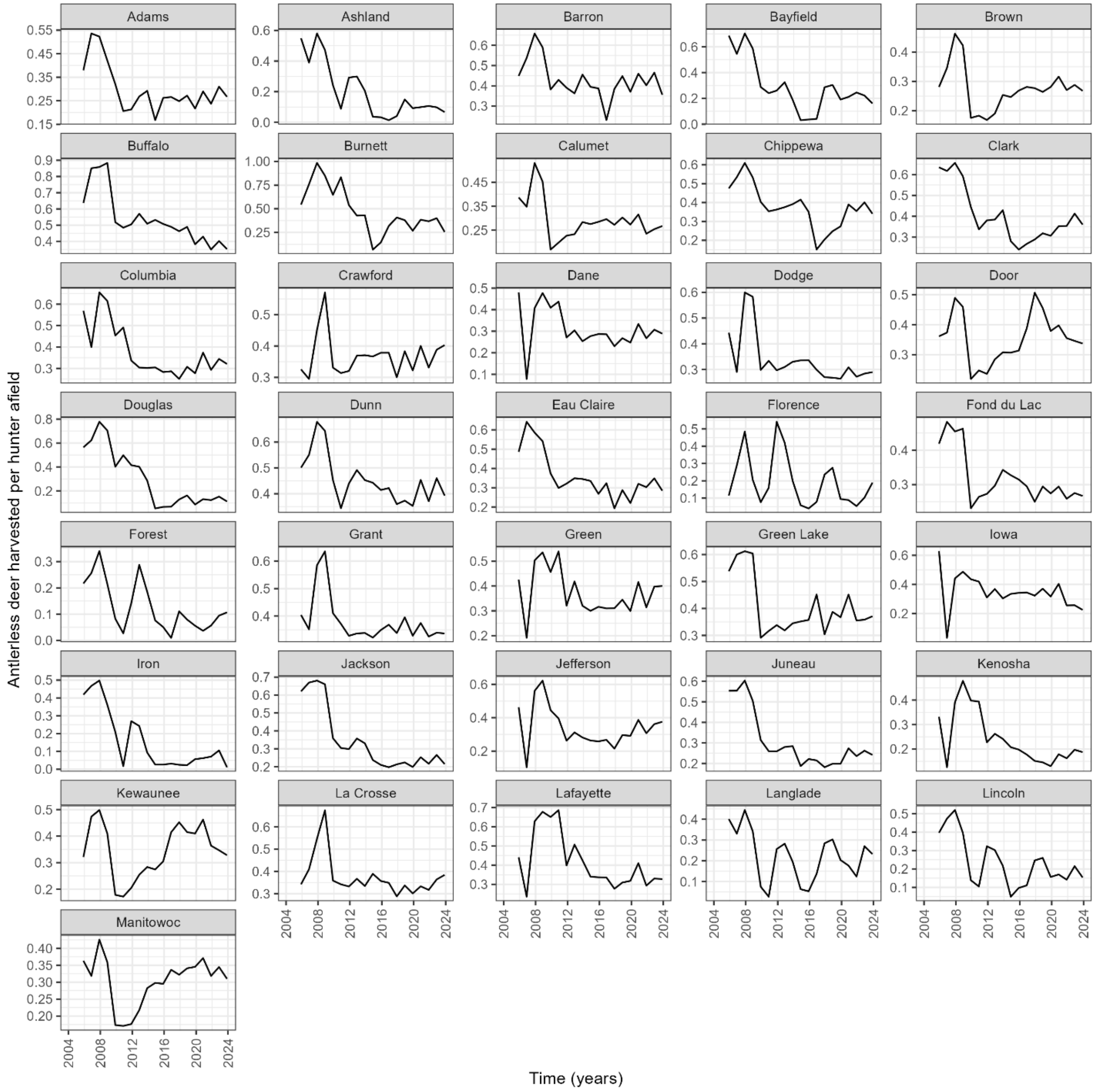


**Figure S30. The number of antlerless deer harvested per hunter afield in each county as predicted by the hunter submodel when fit to the full dataset.** Solid lines represent posterior mean predictions, with 95% credible intervals shaded in grey, calculated across 1,000 parameter sets sampled from the joint posterior to reflect both parameter uncertainty and data measurement error. Note that the credible intervals are very small given the small variation in the number of hunters predicted. The estimates were calculated by dividing the historical number of antlerless deer harvested (from Wisconsin Department of Natural Resources records) by the model-estimated total number of hunters afield in each county, thus including hunters who did not harvest any deer. This figure presents results for the first half of Wisconsin counties included in the model fitting.

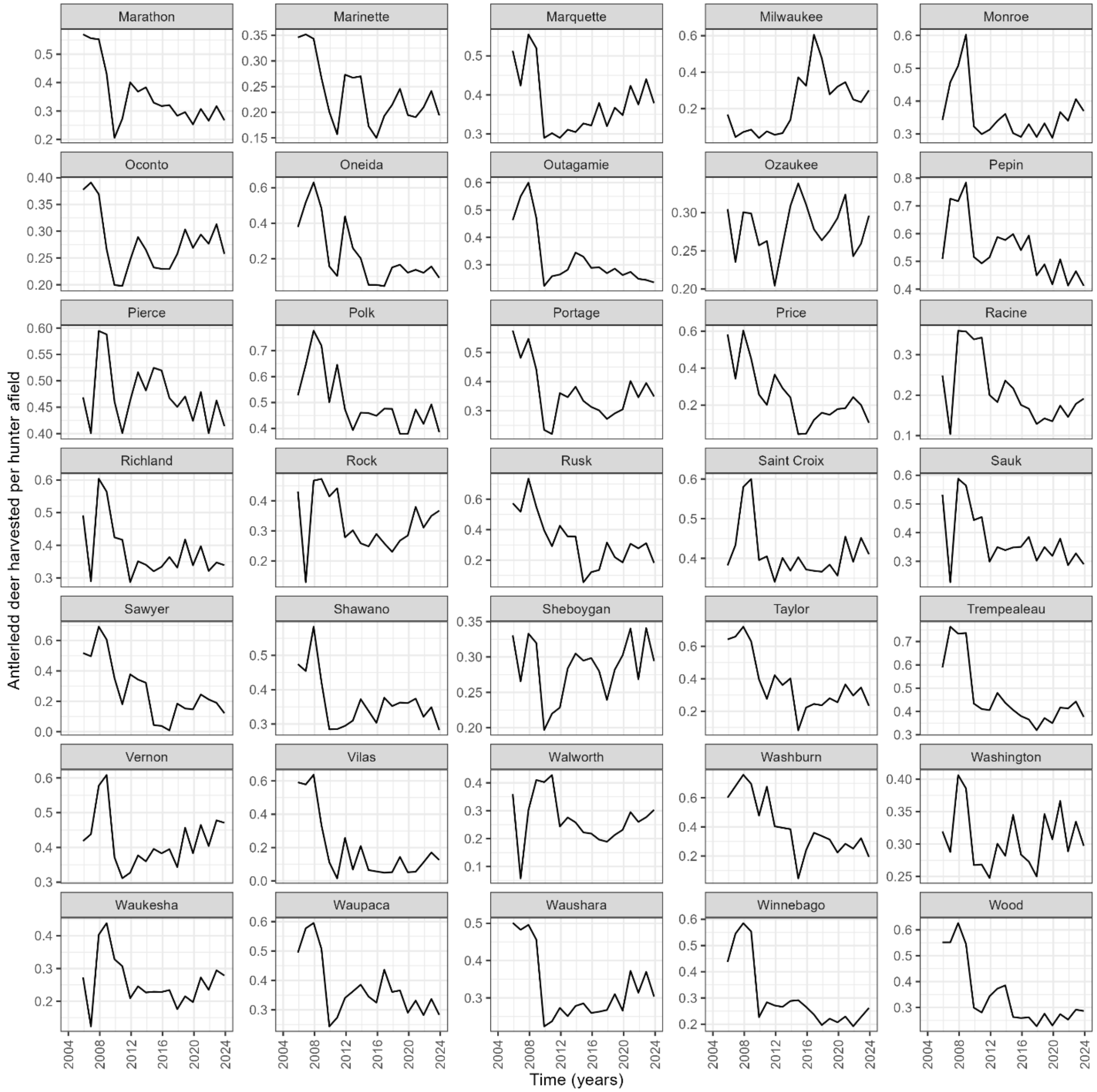


**Figure S31. The number of antlerless deer harvested per hunter afield in each county as predicted by the hunter submodel when fit to the full dataset.** Solid lines represent posterior mean predictions, with 95% credible intervals shaded in grey, calculated across 1,000 parameter sets sampled from the joint posterior to reflect both parameter uncertainty and data measurement error. Note that the credible intervals are very small given the small variation in the number of hunters predicted. The estimates were calculated by dividing the historical number of antlerless deer harvested (from Wisconsin Department of Natural Resources records) by the model-estimated total number of hunters afield in each county, thus including hunters who did not harvest any deer. This figure presents results for the second half of Wisconsin counties included in the model fitting.

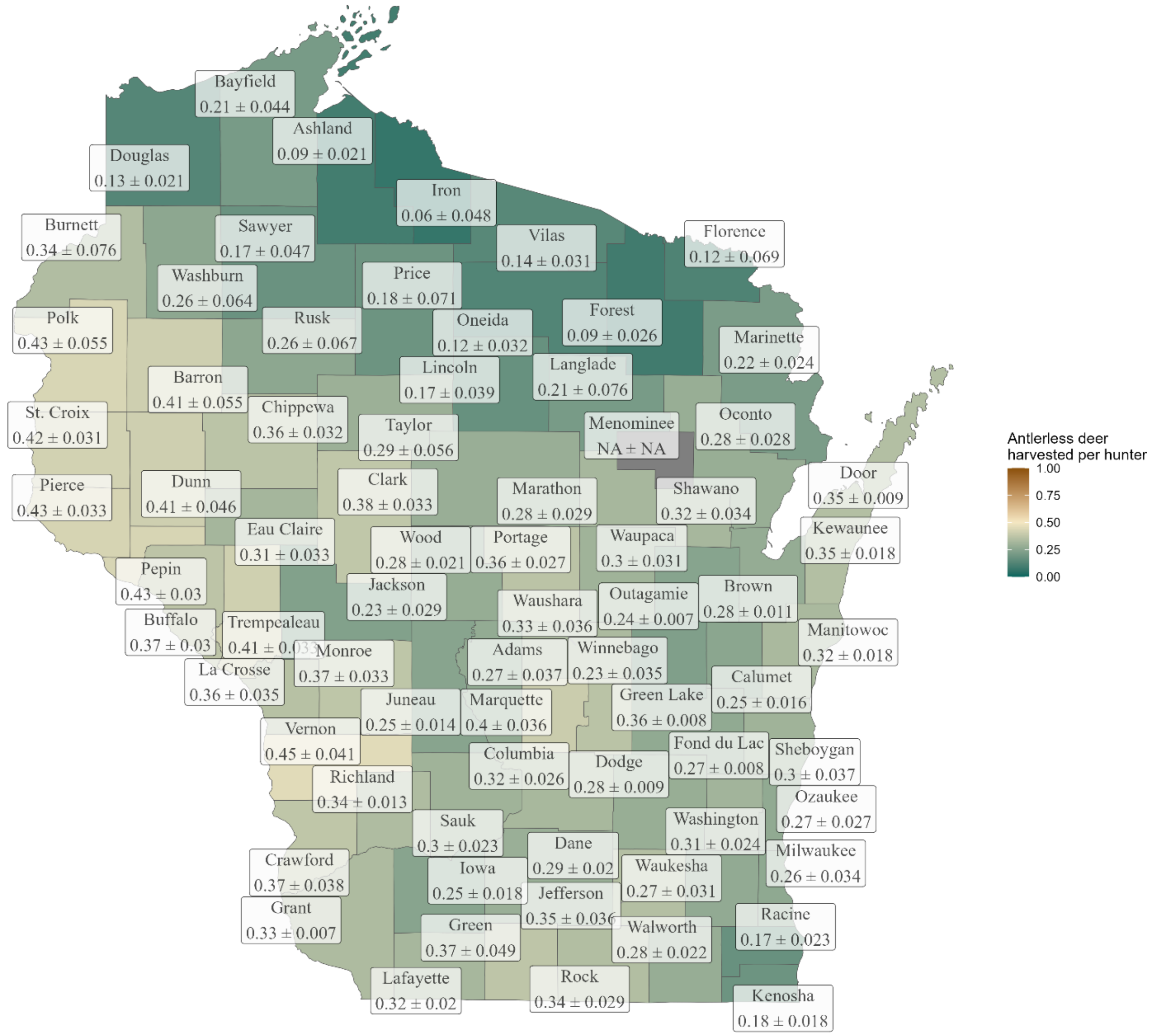


**Figure S32. The number of antlerless deer harvested per hunter afield in each Wisconsin county, averaged over the final three years of data, as predicted by the hunter submodel, fit to full dataset.** The estimates were calculated by dividing the historical number of antlerless deer harvested (from Wisconsin Department of Natural Resources records) by the model-estimated total number of hunters afield in each county, thus including hunters who did not harvest any deer.

To test trends in the average number of deer harvested per hunter over time, we compared the fit of a Bayesian multilevel model with both random intercepts and slopes by region to a simpler model with only random intercepts using leave-one-out (LOO) cross-validation. All Analyses were performed in the statistical program R (R Core Team, 2021), models were fit with the *brms()* package (Bürkner, 2021) and compared with the *loo()* package (Vehtari et al., 2024). We used a Gaussian likelihood with an identity link function to assess trends in the average value across time and regions as follows:

$H_{j,t} \sim N(\mu_{j,t}, \sigma^2)$ Data model

where the observed per hunter harvest rate ($H_{ij}$) for each region (*j*) at each timepoint (*t*) is drawn from a normal distribution with standard deviation ($\sigma^2$). The two process models were implemented as follows:

$\mu_{j,t} = \beta_0 + \mu_{0_j} + \beta_1 Time_t$ Random intercept process model

$\mu_{j,t} = \beta_0 + \mu_{0_j} + (\beta_1 + \mu_{1_j})Time_t$ Random intercept and slope process model

the global intercept ($\beta_0$) and random intercept ($\mu_{0_j}$) to account for regional variation are in each as well as a fixed effect of time ($\beta_1$). An additional term for random slope ($\mu_{1_{j,t}}$) is included to test for regional variation in the trend over time. Default priors from the *brm()* function were assigned for all model parameters.

The model including both random intercepts and slopes had significantly better predictive performance for both per hunter harvest of antlerless (ΔELPD = -173.5, SE = 18.4) and antlered deer (ΔELPD = -283.0, SE = 30.2).

For antlerless deer, the global time effect across all regions was negative ($\beta_1$= –0.0093, 95% credible interval (CI): –0.0116, –0.0071) and the 95% credible interval does not include zero, providing strong evidence for an overall decline in antlerless harvest per hunter over our time series. There was moderate variation across regions in both baseline harvest levels and temporal trends. The standard deviation of region-specific intercepts was estimated at 0.0785 (95% CI: 0.0656, 0.0939), while the standard deviation of regional slopes was 0.0088 (95% CI: 0.0072, 0.0108), suggesting some heterogeneity in how the trend varied across space. The mean region-specific slope for 63 out of 71 regions were negative. The mean region-specific slopes for Brown, Crawford, Door, Kewaunee, Manitowoc, Milwaukee, Saint Croix, and Vernon counties were positive but close to zero (0.000836 to 0.0107).

For antlered deer, the global time effect across all regions was positive and credibly different from zero ($\beta_1$= 0.0027, 95% CI: 0.0014, 0.0041), suggesting that the antlered per hunter harvest has slightly increased over our time series. There was moderate variability in baseline harvest levels and temporal trends across regions. The standard deviation of region-specific intercepts was 0.0395 (95% CI: 0.0331, 0.0471), while the standard deviation of regional slopes was 0.0057 (95% CI: 0.0048, 0.0069), suggesting some heterogeneity in how the trend varied across space. The mean region-specific slope for 51 out of 71 regions were positive. The mean region-specific slopes for Ashland, Bayfield, Buffalo, Burnett, Clark, Douglas, Florence, Forest, Iron, Jackson, Lincoln, Oneida, Polk, Price, Rusk, Sawyer, Taylor, Vilas, Washburn, and Wood counties were negative but close to zero (-0.01147 to $-4.507 \times 10^{-6}$).

The correlation between intercepts and slopes was positive but highly uncertain for antlerless deer (mean ρ = -0.0967, 95% CI: –0.1616 to 0.3472) and strongly negative for antlered deer (ρ= -0.6799, 95% CI: –0.8031 to -0.5167). These results indicate no strong evidence of a systematic relationship between baseline harvest levels and the rate of change over time in the per hunter harvest of antlerless deer. In comparison, results suggest that regions with higher initial per hunter antlered deer harvest rates tend to have slower growth or even to decline over time, while regions with lower starting values tended to experience more rapid increases.

There was no evidence of model non-convergence for either dataset ($\hat{R} \approx 1$ for all parameters), and effective sample sizes were sufficient for stable inference.

**Table S19. Parameter estimates from the Bayesian multilevel model including both random intercepts and slopes fit to the time series of the per hunter harvest of antlered and antlerless deer.** Models included a global intercept ($\beta_0$) fixed effect of time ($\beta_1$), and random region-level effects on the intercept ($\mu_{0_j}$) and slope ($\mu_{1_j}$) for each Wisconsin county for which the per hunter harvests were estimated. We also report the correlation between these regional level effects. Shown are posterior means and 95% credible intervals (CI) for each parameter, including the residual standard deviation (σ) and the standard deviation of the region-level intercepts. **Note:** All parameters are on the scale of the response variable (i.e., proportion or rate of antlerless harvest). SE = standard error; CI = credible interval; ESS = effective sample size.

| **Parameter** | **Dataset** | **Estimate** | **SE** | **95% CI** | **$\hat{R}$** | **Bulk ESS** | **Tail ESS** |
|---|---|---|---|---|---|---|---|
| *Global* | | | | | | | |
| Intercept | Antlered | 0.2659 | 0.0048 | 0.2567-0.2752 | 1.0056 | 567 | 1424 |
| | Antlerless | 0.3266 | 0.0097 | 0.3077-0.3456 | 1.0083 | 483 | 763 |
| Time (centered) | Antlered | 0.0027 | 0.0007 | 0.0014-0.0041 | 1.0043 | 861 | 2002 |
| | Antlerless | -0.0093 | 0.0011 | -(0.0116-0.0071) | 1.0014 | 1272 | 2522 |
| Residual SD | Antlered | 0.0400 | 0.0008 | 0.0385-0.0416 | 1.0002 | 9171 | 6248 |
| | Antlerless | 0.0807 | 0.0017 | 0.0775-0.0840 | 1.0001 | 7399 | 5787 |
| *Random Effects (Region)* | | | | | | | |
| SD (intercept) | Antlered | 0.0395 | 0.0036 | 0.0331-0.0471 | 1.0045 | 1170 | 2294 |
| | Antlerless | 0.0785 | 0.0072 | 0.0656-0.0939 | 1.0013 | 1080 | 2168 |
| SD (time slope) | Antlered | 0.0057 | 0.0005 | 0.0048-0.0069 | 1.0041 | 1327 | 3369 |
| | Antlerless | 0.0088 | 0.0009 | 0.0072-0.0108 | 1.0012 | 1942 | 3192 |
| Correlation (intercept/slope) | Antlered | -0.6799 | 0.0729 | -0.831-(-0.5167) | 1.0025 | 1590 | 3099 |
| | Antlerless | 0.0967 | 0.1307 | -0.1616-0.3472 | 1.0033 | 1330 | 2529 |

#### 4.2.3 Sensitivity analyses

##### *4.2.3.1 Local parameter sensitivity*

a)

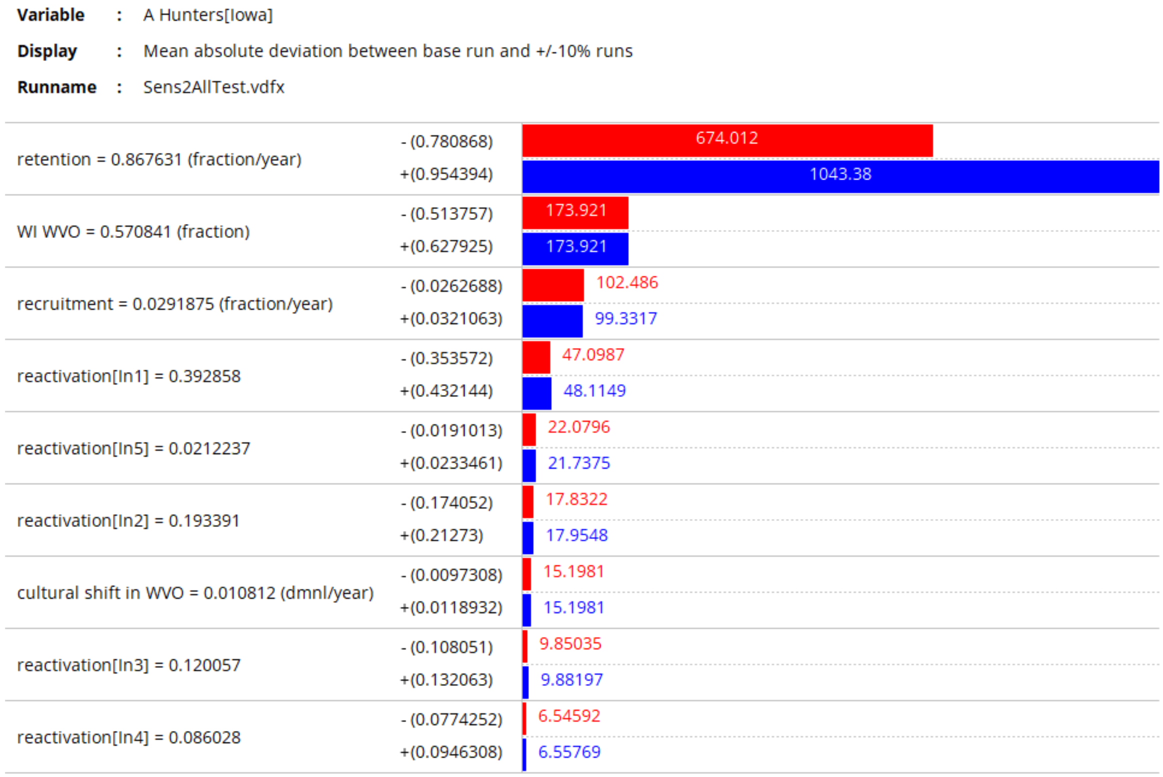


b)

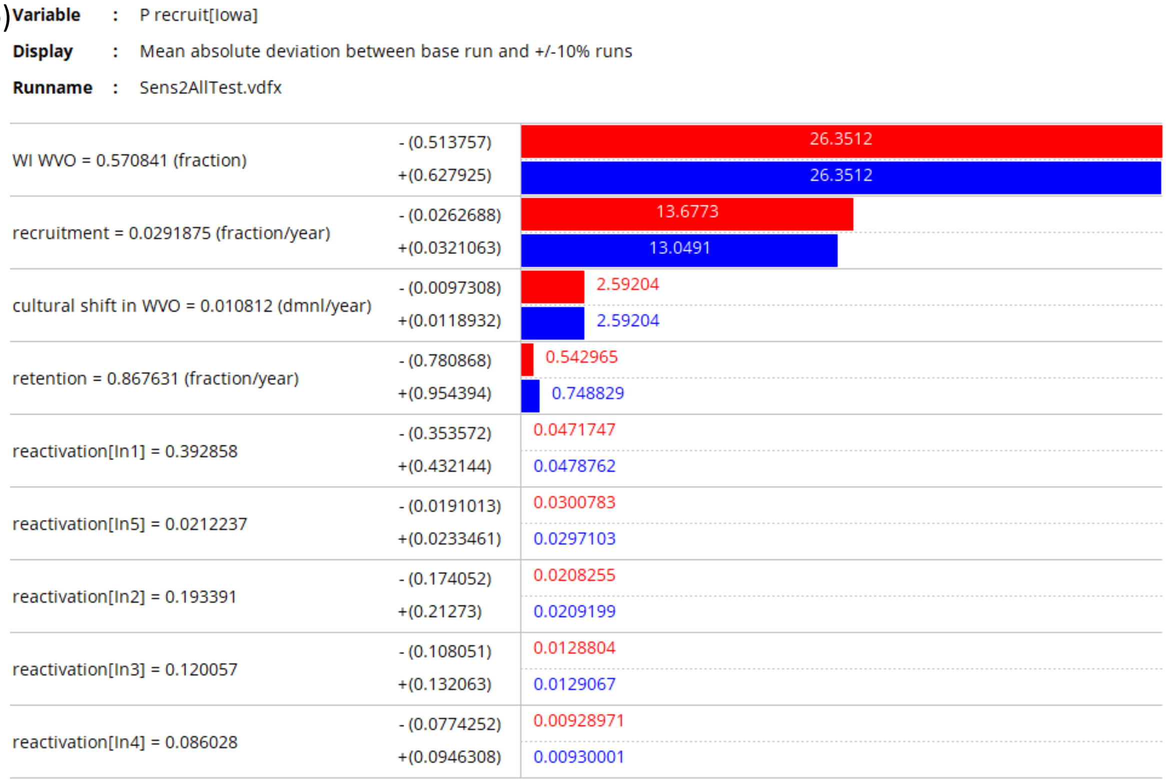

c)

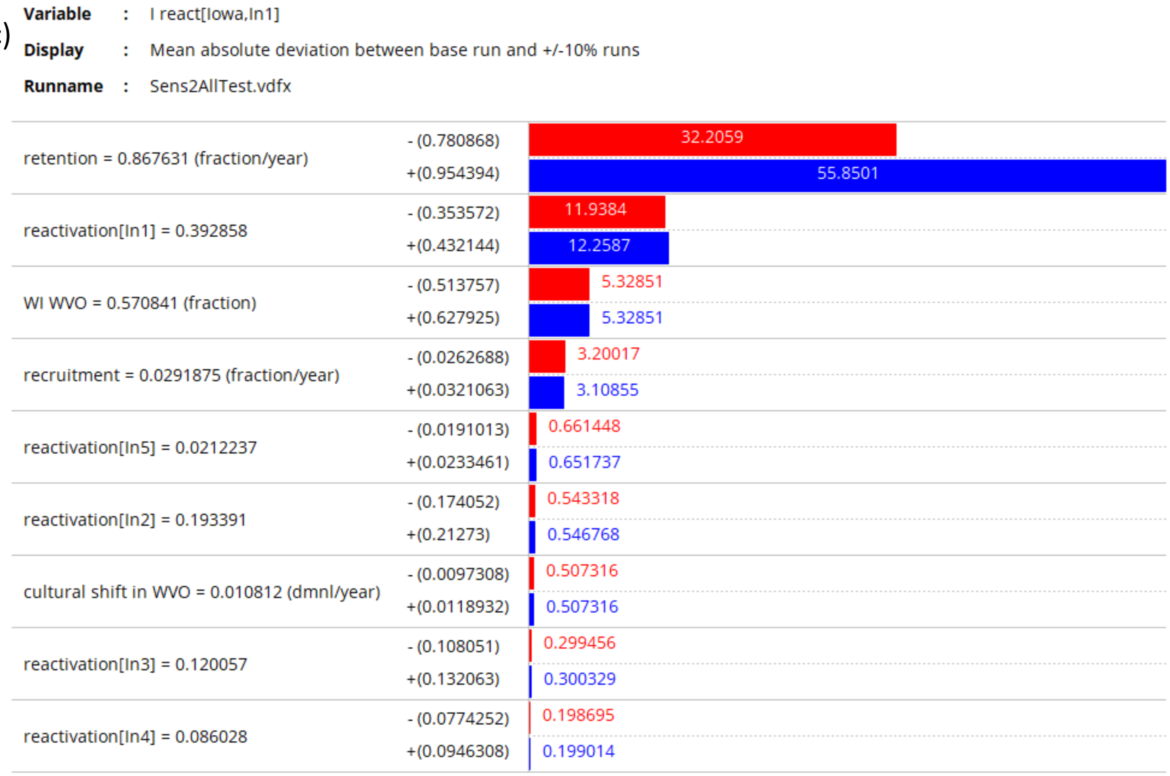
Variable : I react[Iowa,In1]
Display : Mean absolute deviation between base run and +/-10% runs
Runname : Sens2AllTest.vdfx
retention = 0.867631 (fraction/year)
- (0.780868)
32.2059
+(0.954394)
55.8501
reactivation[In1] = 0.392858
- (0.353572)
11.9384
+(0.432144)
12.2587
WI WVO = 0.570841 (fraction)
- (0.513757)
5.32851
+(0.627925)
5.32851
recruitment = 0.0291875 (fraction/year)
- (0.0262688)
3.20017
+(0.0321063)
3.10855
reactivation[In5] = 0.0212237
- (0.0191013)
0.661448
+(0.0233461)
0.651737
reactivation[In2] = 0.193391
- (0.174052)
0.543318
+(0.21273)
0.546768
cultural shift in WVO = 0.010812 (dmnl/year)
- (0.0097308)
0.507316
+(0.0118932)
0.507316
reactivation[In3] = 0.120057
- (0.108051)
0.299456
+(0.132063)
0.300329
reactivation[In4] = 0.086028
- (0.0774252)
0.198695
+(0.0946308)
0.199014

d)

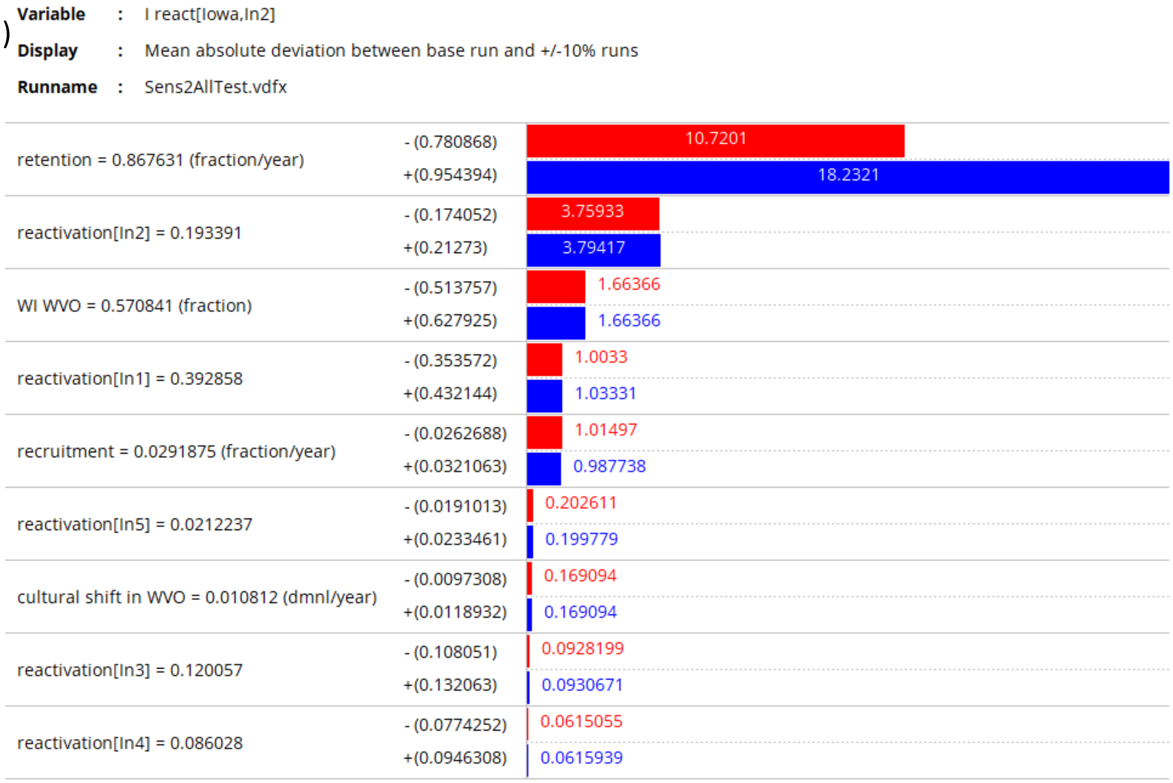
Variable : I react[Iowa,In2]
Display : Mean absolute deviation between base run and +/-10% runs
Runname : Sens2AllTest.vdfx
retention = 0.867631 (fraction/year)
- (0.780868)
10.7201
+(0.954394)
18.2321
reactivation[In2] = 0.193391
- (0.174052)
3.75933
+(0.21273)
3.79417
WI WVO = 0.570841 (fraction)
- (0.513757)
1.66366
+(0.627925)
1.66366
reactivation[In1] = 0.392858
- (0.353572)
1.0033
+(0.432144)
1.03331
recruitment = 0.0291875 (fraction/year)
- (0.0262688)
1.01497
+(0.0321063)
0.987738
reactivation[In5] = 0.0212237
- (0.0191013)
0.202611
+(0.0233461)
0.199779
cultural shift in WVO = 0.010812 (dmnl/year)
- (0.0097308)
0.169094
+(0.0118932)
0.169094
reactivation[In3] = 0.120057
- (0.108051)
0.0928199
+(0.132063)
0.0930671
reactivation[In4] = 0.086028
- (0.0774252)
0.0615055
+(0.0946308)
0.0615939

e)

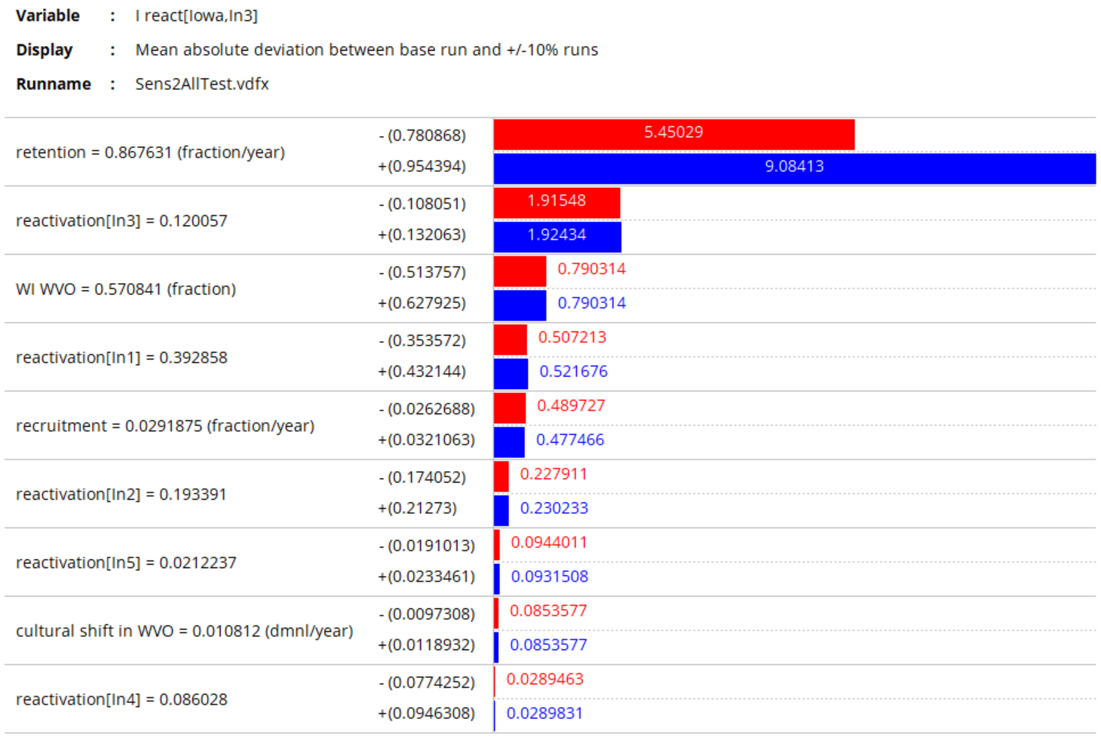
Variable : I react[Iowa,In3]
Display : Mean absolute deviation between base run and +/-10% runs
Runname : Sens2AllTest.vdfx
retention = 0.867631 (fraction/year)
- (0.780868)
+(0.954394)
5.45029
9.08413
reactivation[In3] = 0.120057
- (0.108051)
+(0.132063)
1.91548
1.92434
WI WVO = 0.570841 (fraction)
- (0.513757)
+(0.627925)
0.790314
0.790314
reactivation[In1] = 0.392858
- (0.353572)
+(0.432144)
0.507213
0.521676
recruitment = 0.0291875 (fraction/year)
- (0.0262688)
+(0.0321063)
0.489727
0.477466
reactivation[In2] = 0.193391
- (0.174052)
+(0.21273)
0.227911
0.230233
reactivation[In5] = 0.0212237
- (0.0191013)
+(0.0233461)
0.0944011
0.0931508
cultural shift in WVO = 0.010812 (dmnl/year)
- (0.0097308)
+(0.0118932)
0.0853577
0.0853577
reactivation[In4] = 0.086028
- (0.0774252)
+(0.0946308)
0.0289463
0.0289831

f)

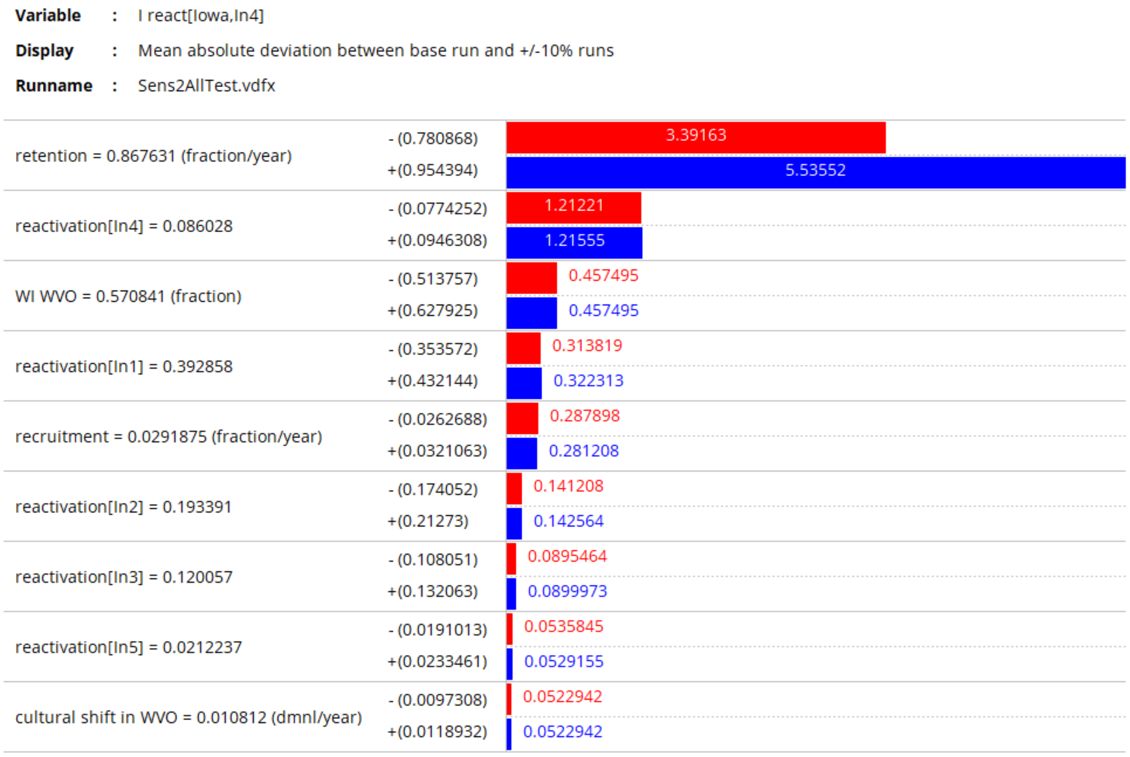
Variable : I react[Iowa,In4]
Display : Mean absolute deviation between base run and +/-10% runs
Runname : Sens2AllTest.vdfx
retention = 0.867631 (fraction/year)
- (0.780868)
+(0.954394)
3.39163
5.53552
reactivation[In4] = 0.086028
- (0.0774252)
+(0.0946308)
1.21221
1.21555
WI WVO = 0.570841 (fraction)
- (0.513757)
+(0.627925)
0.457495
0.457495
reactivation[In1] = 0.392858
- (0.353572)
+(0.432144)
0.313819
0.322313
recruitment = 0.0291875 (fraction/year)
- (0.0262688)
+(0.0321063)
0.287898
0.281208
reactivation[In2] = 0.193391
- (0.174052)
+(0.21273)
0.141208
0.142564
reactivation[In3] = 0.120057
- (0.108051)
+(0.132063)
0.0895464
0.0899973
reactivation[In5] = 0.0212237
- (0.0191013)
+(0.0233461)
0.0535845
0.0529155
cultural shift in WVO = 0.010812 (dmnl/year)
- (0.0097308)
+(0.0118932)
0.0522942
0.0522942

g)

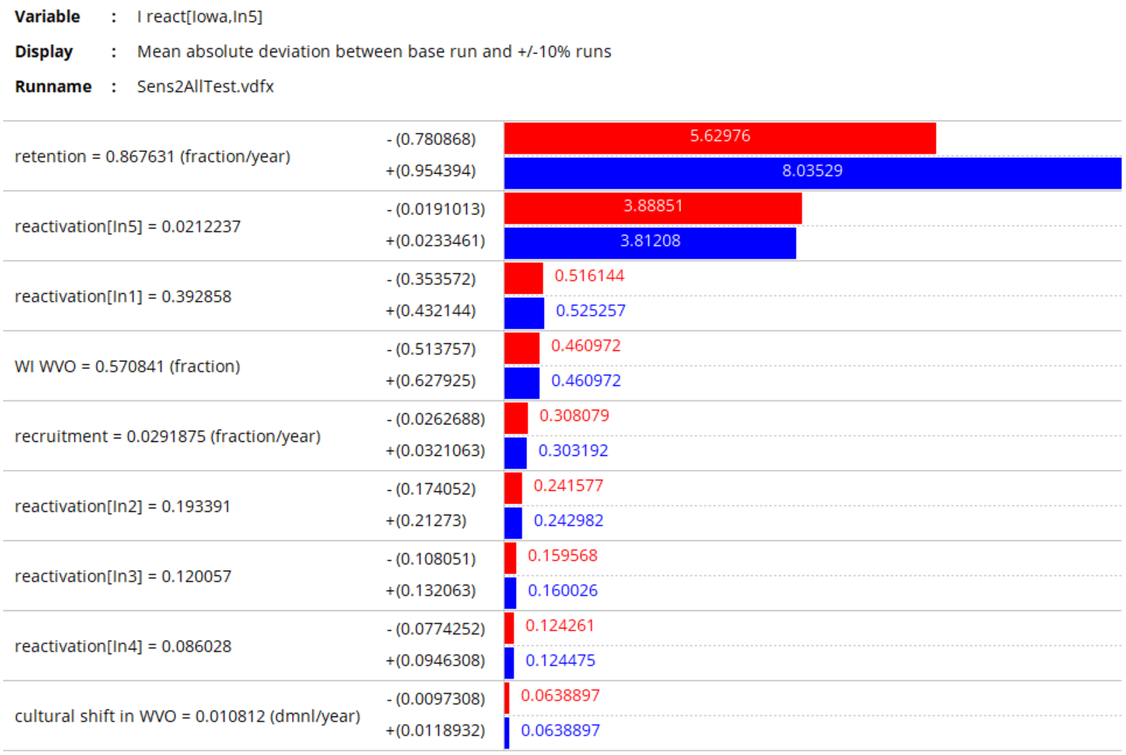


**Figure 33. The results from the *Sensitivy2All* local parameter sensitivity test, performed in Vensim, are presented for each model simulated estimate of the historical time-series data used to fit the hunter submodel model.** For each parameter estimated during model calibration, the mean value was independently increased and decreased by 10%. The mean absolute deviation between the baseline simulation—using all mean parameter estimates—and each perturbed run is reported (blue and red bars indicate decreased and increased parameters, respectively). Sensitivity results are shown for the following model predictions for Iowa county (units in brackets) in this order: (a) the number of active hunters (people), (b) the number of recruitments (people/year), (c-g) the number of reactivations from each stage of inactive hunters (people/year). The results from increasing and decreasing parameters are shown in blue and red respectively while subscripts are shown in square brackets (e.g., Inactive = In1).
